# Contrasting effects of forest fragmentation on the genetics and microbiomes of an endangered arboreal primate

**DOI:** 10.64898/2026.06.14.732134

**Authors:** Keren Klass, Sarie Van Belle, Eva C. Wikberg, Gwen Duytschaever, Maria Luisa Savo Sardaro, Adriana Morales-Guerrero, Ohad Peled, Keith D. Harris, Rachel M. Petersen, Maike Lynn Morrison, Michelle Soffer, Anthony Di Fiore, Katherine Ryan Amato, Amanda D. Melin, Julie A. Teichroeb, Gili Greenbaum

## Abstract

Landscape fragmentation, one of the leading drivers of biodiversity loss, can reshape both the genetics and microbiomes of wild populations. Although fragmentation is generally expected to limit gene flow, erode genetic diversity, and disrupt host-associated microbial communities, these responses arise via different pathways and may therefore diverge within the same population. To understand how fragmentation simultaneously shapes population genetics and gut microbiomes, we analyzed fecal-derived host genomic and microbiome data from endangered, arboreal black howler monkeys (*Alouatta pigra*) across a fragmentation gradient. We then integrated these data with measures of ecological connectivity, habitat quality, and demography to identify the drivers of genetic and microbiome variation and spatial structure. Multivariate analyses indicated that genetic patterns were shaped by both connectivity and habitat quality, whereas microbiome variation was driven mainly by habitat quality. Contrary to expectations, monkeys showed the strongest gene flow signal in the most fragmented region with the lowest connectivity, as well as higher genetic diversity and lower inbreeding compared to monkeys in continuous forest. Relatedness and isolation-by-distance patterns suggested that fragmentation has sex-specific effects on movement, disrupting the usual pattern of short-range male dispersal in the most fragmented region. Gut microbiomes, however, showed expected negative responses to fragmentation: individuals in highly fragmented habitat had lower microbial diversity and compositional shifts consistent with lower-quality diets and increased exposure to disturbed environments. These results show contrasting biological responses to fragmentation within a single population, with genetic patterns likely resulting from compensatory behavioral flexibility and microbiome patterns reflecting local habitat degradation. Our findings underscore the need for conservation assessments that integrate multiple dimensions of population health rather than relying on any single indicator of fragmentation impact.

## Introduction

Like much of global biodiversity, the order Primates is undergoing an extinction crisis^1,2^. Habitat loss and fragmentation are the most urgent threats to primate populations, with myriad simultaneous negative effects on behavior, ecology, health, and demography^2,3^. However, for primates and other species, the specific combinations and magnitudes of these effects depend on the landscape configuration and species characteristics, and fragmentation may affect different aspects of a population’s biology differently^4–6^ and at varying time scales^3^. Effective management and conservation require understanding the complexities of these responses to habitat loss and fragmentation^2^.

Classical population-genetic models predict that when fragmentation reduces realized connectivity among habitat patches, restricted gene flow should increase genetic drift and inbreeding, reduce local genetic variation, and strengthen landscape-scale population structure^7–9^. Such genetic signatures of reduced connectivity have been observed in many taxa, including in primates^10–18^, and are of conservation concern because they may lead to inbreeding depression and a reduced potential to adaptively respond to environmental changes^15,19–21^, heightening extinction risk^22^. However, these predictions rest on the ecological assumption that structural fragmentation translates into reduced functional connectivity. In behaviorally and ecologically flexible species, that assumption may not always hold, because fragmentation can also alter dispersal incentives, habitat quality, and sex-specific movement decisions^23–26^. As a result, genetic responses to fragmentation may depend not only on the spatial configuration of the landscape, but also on how individuals and populations respond behaviorally, ecologically, and demographically to the altered landscape (e.g.,^27,28^).

Whereas population genetic responses to fragmentation often depend on how structural fragmentation translates into realized movement and gene flow, gut microbiomes may reflect a different set of fragmentation-mediated processes, such as habitat degradation, dietary change, and exposure to disturbed environments. Fragmentation can, therefore, lead to spatial variation in gut microbiomes because microbial communities reflect variation in habitat quality and diet across the landscape^29,30^. For example, intraspecific comparisons of mammal populations living in continuous vs. anthropogenically fragmented habitats have shown increased microbiome heterogeneity between individuals in more disturbed habitat^31,32^. Dietary and behavioral characteristics also shape the microbiome, and populations in forest fragments can show either increased or decreased microbiota diversity compared to those in continuous forests^5^. For species with specialized diets (e.g., folivorous primates), lower microbiome diversity may reflect reduced dietary diversity in lower-quality forest fragments^33,34^. Additional factors affecting gut microbiomes, such as host physiological stress, microbial transmission within social groups, host demography, or host genetics^35–42^ can also be shaped by fragmentation^43^. Imbalanced and depleted microbiomes may lead to impaired immune defense, hormone imbalances, and behavioral disorders^21,31,44^, and these impacts on health and fitness have conservation implications for endangered species in fragmented landscapes. Additionally, the microbiome responds to environmental change at much shorter time scales than many other aspects of species’ behavior or ecology^45^, making it a potentially valuable proxy for estimating health-related fragmentation effects. However, in contrast to population genetic effects, there are fewer studies and no clear consensus on how fragmentation shapes wild animal gut microbiomes.

To evaluate how landscape fragmentation simultaneously shapes population genetics and gut microbiomes, we studied both types of data over a range of fragmentation intensities in a well-suited system: that of the endangered^46^, arboreal black howler monkey (*Alouatta pigra*) in and around Palenque National Park in Chiapas, Mexico. Although black howlers demonstrate considerable flexibility in their behavioral ecology^47–53^ and are generally considered able to persist in fragmented forests^48^, negative effects of fragmentation on black howlers have been observed^54–57^. In the unprotected landscape around Palenque National Park (Fig. 1B), anthropogenic forest loss and fragmentation began *∼* 100 years ago and have progressed unevenly^58^. This process creates a natural experiment for studying how the population genetics and gut microbiomes of howlers, originally from the same habitat and population, are shaped by varying levels of forest loss, degradation, and fragmentation, without the confounding effects of intrinsic variation in genetics, behavior, or ecology^59–61^.

**Figure 1:**
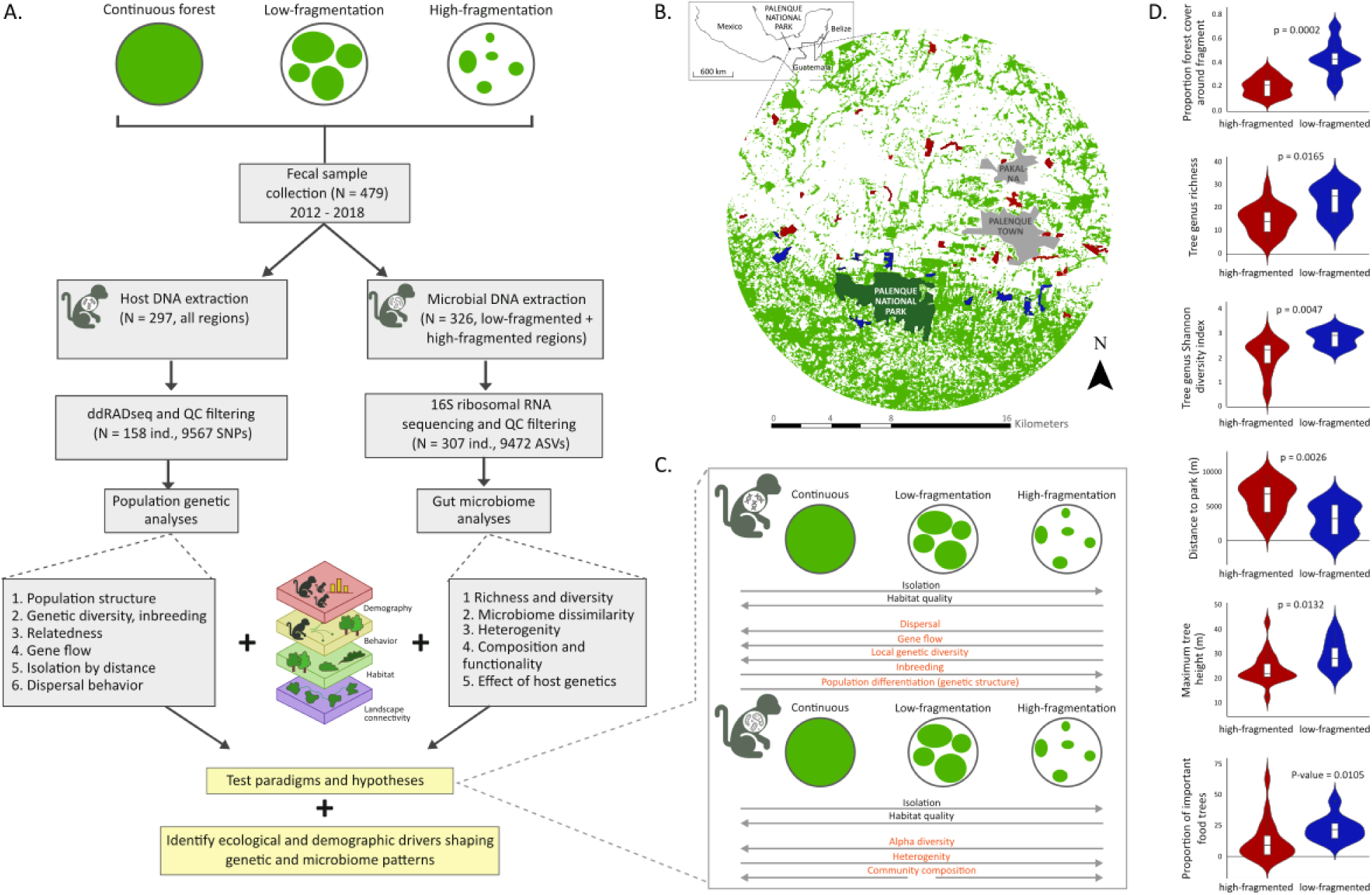
Study design. (A) Methods and workflow: we collected fecal samples from habitats with varying levels of fragmentation, extracted genetic and microbiome data, and integrated these data with ecological and behavioral data to test hypotheses and identify ecological drivers of genetic and microbiome patterns. (B) Map of study site: light green areas denote forest cover, white areas denote non-forest matrix, dark green denotes the continuous forest region (Palenque National Park), blue denotes sampled forest fragments in the low-fragmented region, red denotes sampled forest fragments in the high-fragmented region, gray denotes towns. *Inset* : location of study site in southeastern Mexico. (C) Hypotheses regarding the effects of fragmentation on population genetics and microbiomes that were examined in this study. At the top are hypotheses generated from classical population-genetic models assuming reduced realized connectivity with increasing fragmentation, and at the bottom hypotheses regarding microbiome responses to increasing fragmentation. (D) Differences in structural connectivity and habitat quality variables between the low (*N* = 10; blue) and high (*N* = 21; red) fragmentation regions (significance of differences shown for each variable).

We took advantage of this natural experiment to address three questions: (i) whether standard conservation-genetic expectations based on reduced connectivity^9,19^ predict genetic patterns in black howlers in a fragmented landscape, or whether these patterns are modified by habitat quality or species-specific characteristics; (ii) if howler gut microbiome richness, diversity, and homogeneity decrease with increasing fragmentation^33^; and (iii) how specific measures of habitat quality, connectivity, and demography shape both population genetic and gut microbiome patterns in the fragmented landscape (Fig. 1A). We assembled large datasets of fecal sample-derived host and microbial DNA, spanning hundreds of individuals and dozens of sampling sites, and built on an existing foundation of demographic, behavioral, habitat, and landscape data^33,47,51,53,58,62^. This integrative approach allowed us to evaluate not only whether fragmentation affects population genetics and gut microbiomes, which is informative in and of itself for assessing a species’ condition and likelihood of long-term persistence^22,63^, but also whether these biological dimensions respond to fragmentation similarly or through distinct ecological pathways. Our results show that different indicators of population health can respond to fragmentation in contrasting and potentially misleading ways if considered alone, underscoring the need for integrative assessments of fragmentation impacts using multiple complementary data types for endangered species.

## Results

Approximately 100 years ago, forest loss and fragmentation began progressing unevenly across the landscape around Palenque National Park in Chiapas, Mexico^58^ (Fig. 1B): the region to the north of the park is highly fragmented, with higher levels of isolation between fragments, while the southern region around Palenque National Park has maintained higher connectivity; the park itself is continuous primary and secondary rainforest that is 2–3 orders of magnitude larger than any other forest fragment in the region^51^. We examined how varying levels of forest fragmentation affected patterns in black howler genetics and microbiomes, evaluating our results in terms of our hypotheses (Fig. 1C), and conducted analyses to identify the specific drivers of genetic and microbiome variation across the landscape (Table S3). We collected fecal samples from howlers in unprotected forest fragments (henceforth ‘fragmented region’) around Palenque National Park, and extracted host DNA for population genetic analyses and microbial DNA for gut microbiome analyses (Fig. 1A; Tables S1, S2); host DNA from fecal samples was also collected in Palenque National Park^62^. We refer to the north as the ‘high-fragmented’ region, the southern region outside the park as the ‘low-fragmented’ region, and to the park itself as ‘continuous forest’ (Fig. 1). Importantly, the forest fragments in the high-fragmented region constitute lower-quality habitats for howlers than those in the low-fragmented region^58^ (Fig. 1D).

### Fragmentation is associated with increased gene flow across the non-forest matrix

Our genetic dataset included 158 individuals: 28 from the continuous forest, 64 from 11 sampling locations in the low-fragmented region, and 66 from 26 locations in the high-fragmented region (Table S1). Using ddRAD sequencing, we compiled a genomic dataset of 9567 SNPs (SI Methods).

Departing from the simple expectation that fragmentation reduces realized connectivity and therefore leads to reduced genetic diversity and increased inbreeding (Fig. 2E), we found that while observed heterozygosity indeed differed significantly among populations (Kruskal–Wallis *χ*^2^=27.34, df=2, *P <* 0.0001), it was significantly lower in the continuous forest (0.17 *±* 0.03) than in the high-fragmented population (0.21 *±* 0.02; Holm-adjusted Dunn comparisons, *P <* 0.0001) and the low-fragmented population (0.20 *±* 0.03; *P <* 0.0001; Fig. 2D, Table S4). Mean expected heterozygosity was 0.252, 0.247, and 0.240 in the high-fragmented, low-fragmented, and continuous forest populations, respectively. We also observed significantly lower inbreeding in the fragmented regions than in the continuous forest: 0.42 *±* 0.09, 0.43 *±* 0.09, and 0.61 *±* 0.04 individual inbreeding coefficients in the high-fragmented region, low-fragmented region, and continuous forest, respectively (Fig. 2D, Table S4, Fig. S1). These patterns held when subsampling the fragmented landscape to sample sizes and areas similar to that of the continuous forest (Table S5). Within the fragmented region, we did not observe substantial differences in genetic diversity and inbreeding between the high- and low-fragmentation levels (Fig. 2D).

**Figure 2:**
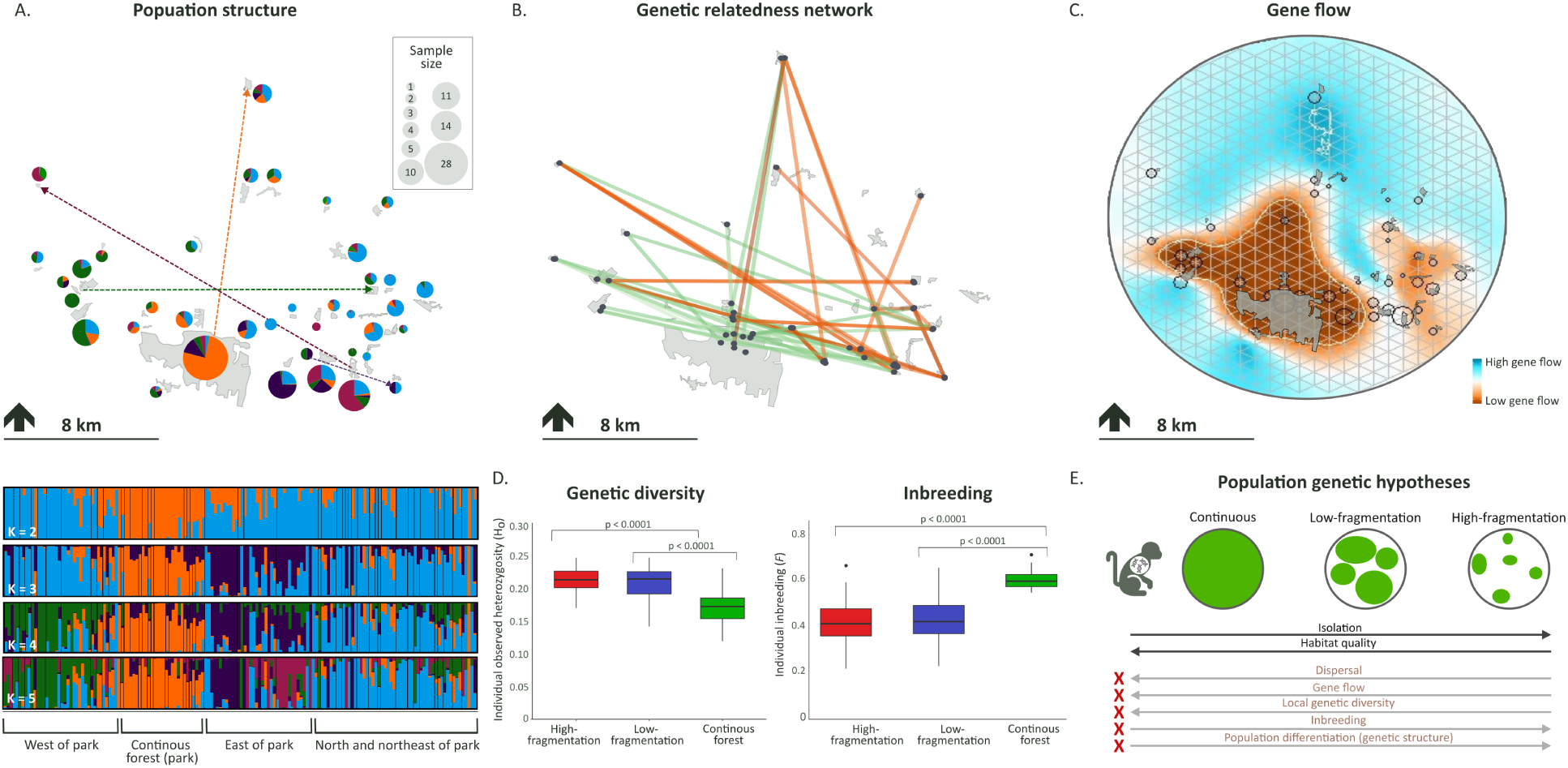
Population genetic patterns reflect altered realized connectivity across the fragmentation gradient. (A) Population structure analysis. Bottom panels show STRUCTURE results for different numbers of putative clusters (*K* = 2–5), with sampling areas noted below the plots. Top panel shows the results visualized spatially, with pie charts for each sampling location showing the proportions of all genetic clusters identified in that location for *K* = 5; the size of the circle corresponds to the number of individuals sampled at that location. Arrows denote potential long-distance dispersal events inferred from pairwise relatedness values, with arrow color corresponding to the dominant genetic cluster at the inferred point of origin and the dispersing individual’s assigned genetic cluster. (B) Pairwise relatedness network of individuals sampled in different locations. Orange lines denote the top 0.25% most highly related pairs, and green lines denote the bottom 0.25% least related pairs, with the magnitude of the estimated *r* denoted by increasingly dark line color. Black circles denote sampled individuals. (C) Migration surface map analysis (EEMS). Gene flow rates are visualized such that zero corresponds to the overall mean migration rate and effective migration that is significantly higher (dark blue) or lower (dark orange) than the overall average is emphasized. (D) Comparison of the distributions of observed individual heterozygosity (left) and inbreeding (right) for the three regions. (E) Summary of whether observed patterns followed classical conservation-genetic model expectations under reduced realized connectivity; patterns departing from these expectations are denoted with a red X.

We also found evidence for higher movement and gene flow in the high-fragmented region. Population genetic structure aligned with levels of fragmentation, with the continuous forest forming a distinct cluster separate from the fragmented region (Fig. 2A; Table S6). Within the fragmented region we observed additional fine-scale population structure, mainly in the low-fragmented region, with samples from fragments west of the park forming a distinct cluster and fragments immediately to the east forming two additional clusters (*K* = 3–5 in Fig. 2A). The remaining fragments, mainly in the high-fragmented region, were largely assigned to a single cluster with considerable admixture (blue cluster in Fig. 2A), indicating gene flow between those sampling locations (Table S6). These results were confirmed by STRUCTURE subsampling analyses to control for differences in sample sizes and sampling area (Fig. S2; Table S5). Migration surface maps reflected the population structure results, showing high gene flow rates in the highly fragmented region, whereas in the continuous forest and adjacent low-fragmented areas we observed higher genetic isolation (Fig. 2C).

The most highly related pairs (top 1%; mean *r* = 0.26 *±* 0.12, range = 0.16–0.76) were concentrated within the same social groups and sampling locations (Table S8), and this pattern was stronger in the continuous forest, with both within-group (*p* = 0.0003) and between-group relatedness (*p <* 0.00001) significantly higher than within fragments (Fig. S4). In the continuous forest, within-group relatedness was significantly higher than between-group relatedness, a pattern not found in fragments with multiple groups (Fig. S4). These results indicated greater group and population stability, and lower dispersal rates, in the continuous forest. We generated a network of extremely high- and low-related pairs sampled in different locations (top 0.25%: mean *r* = 0.30 *±* 0.12, range = 0.22–0.73, and bottom 0.25%: mean *r* = *−*0.09 *±* 0.005, range = -0.08–-0.11; Fig. 2B, Fig. S3; Table S7), which supported the finding that most gene flow across the matrix occurred between forest fragments rather than between fragments and the continuous forest. Most highly related between-fragment pairs (26 of 29) included individuals in the high-fragmented region, whereas most least-related pairs (26 of 29) included individuals in the continuous forest or low-fragmented region (Table S7). The 0.25% most highly related between-fragment pairs (29 pairs with *r ≥* 0.22) were on average sampled 8.6 km apart (range = 0.49 - 18.3 km), suggesting substantially larger dispersal distances in the fragmented regions compared to those previously reported in the continuous forest^53^. Contrasting these large dispersal distances with the relatively short distances to the nearest fragment (average 173±176 m, range: 8–549 m; Table S16), implies that finding suitable groups to join, or a habitat in which to establish a new group, obliges many dispersing individuals to travel well beyond the closest fragment. When considering isolation-by-distance (IBD) slopes, we found that genetic differentiation increase per kilometer was largest in the continuous forest (Fig. S7, Table S10), again supporting more limited movement there.

We also examined whether there were sex-specific effects of fragmentation on movement and gene flow. Studies in the continuous forest suggest that dispersal in black howlers is male-biased, where males disperse to nearby familiar groups. Females are more philopatric, but when they do disperse, they tend to travel longer distances and to unfamiliar areas^53^. Our relatedness results supported these observations in both the continuous and fragmented forest. Specifically, in the fragmented landscape adult male pairs sampled in different groups in the same fragment showed significantly higher relatedness than males sampled in different fragments, suggesting that males prefer to disperse to nearby groups. Adult females showed significantly higher relatedness within social groups, indicating that they were more philopatric than males (Fig. 3A). However, while these general sex-specific dispersal patterns held in the fragmented landscape, we found sex-specific responses of spatial genetic structure to forest fragmentation. Female isolation-by-distance was disrupted by fragmentation *per se*, being significantly steeper in the continuous forest than in both fragmented regions (Fig. 3B,C, Table S10), whereas male isolation-by-distance varied along a fragmentation gradient: males in the low-fragmented region showed significantly steeper isolation-by-distance than both females in the low-fragmented region and males in the high-fragmented region. We also found higher than expected relatedness between males at 500m in the low-fragmented region, but not in the high-fragmented region (Fig. S8, Table S11). These patterns suggest that the lack of short-range dispersal opportunities in the high-fragmented region obliged males to forgo their preference for short-distance dispersal, traveling longer distances than they would in continuous forest or in less fragmented regions.

**Figure 3:**
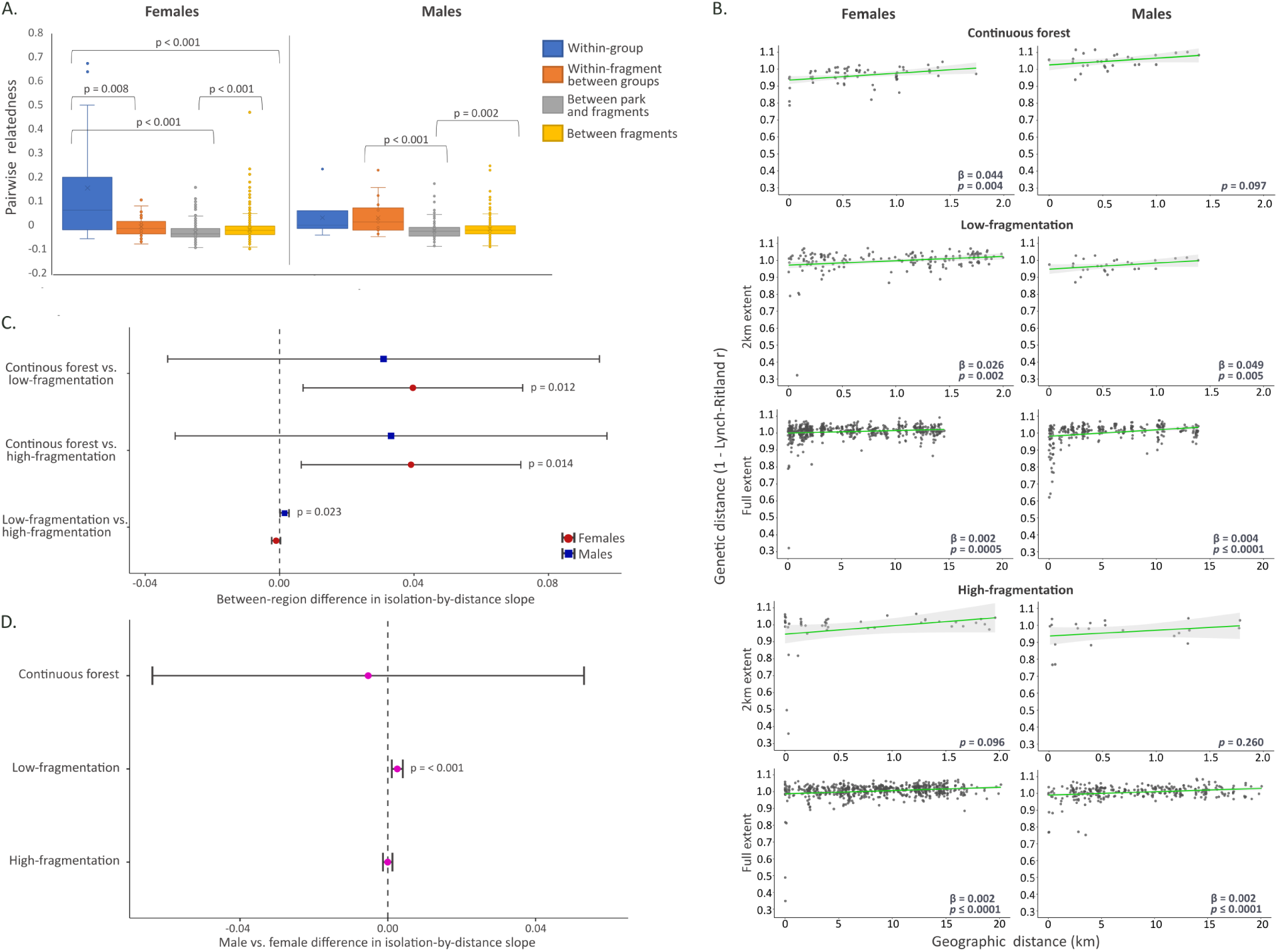
Differences in sex-specific dispersal patterns inferred from genetic analysis. (A) Male and female pairwise relatedness patterns in the fragmented landscape across socio-spatial categories: when both individuals were sampled in the same social group; in different social groups in the same fragment; in different fragments; and one individual in the continuous forest and one in the fragmented landscape. (B) Maximum-likelihood population effects (MLPE) models for IBD across levels of fragmentation and sexes; continuous, low-fragmented, and high-fragmented regions separately in rows, at 2km and full spatial extents; females and males in columns. Slope, 95% CI, and p-values noted on plots. (C) Comparison of IBD strength between regions, for males and females separately. (D) Comparison of male and female IBD strength within each region. For (C) and (D), points show estimated differences in IBD slopes for each pair; horizontal bars indicate 95% confidence intervals; significant p-values noted on plot; vertical dashed line at zero indicates no difference in IBD strength.

### Gut microbiomes show fragmentation-associated differences across the landscape

We compiled gut microbiome data for 307 individuals in the fragmented landscape (Tables S1, S2), consisting of 9472 amplicon sequence variants (ASVs). Individual samples were rarefied to 5000 sequences (Fig. S9). We identified many very rare ASVs, and a handful of very common ASVs (median frequency per ASV = 3, mean = 299, range: 1–156,625).

Individual gut microbiomes clustered clearly according to level of fragmentation (Fig. 4A), but not by age, sex, or sequencing batch (Fig. S10). The 50 samples from the high-fragmented region that overlapped with the low-fragmented region in the ordination space were from eight fragments bordering the low-fragmented region (yellow in Fig. 4A,B; Fig. S12), suggesting that spatial gut microbiome variation was very similar, but not identical to our connectivity-based partition of fragments into low- and high-fragmented regions. We therefore assigned these eight fragments to the low-fragmented region for microbiome analyses (yellow in Fig. 4; re-analyzed habitat variables in Fig. S11).

**Figure 4:**
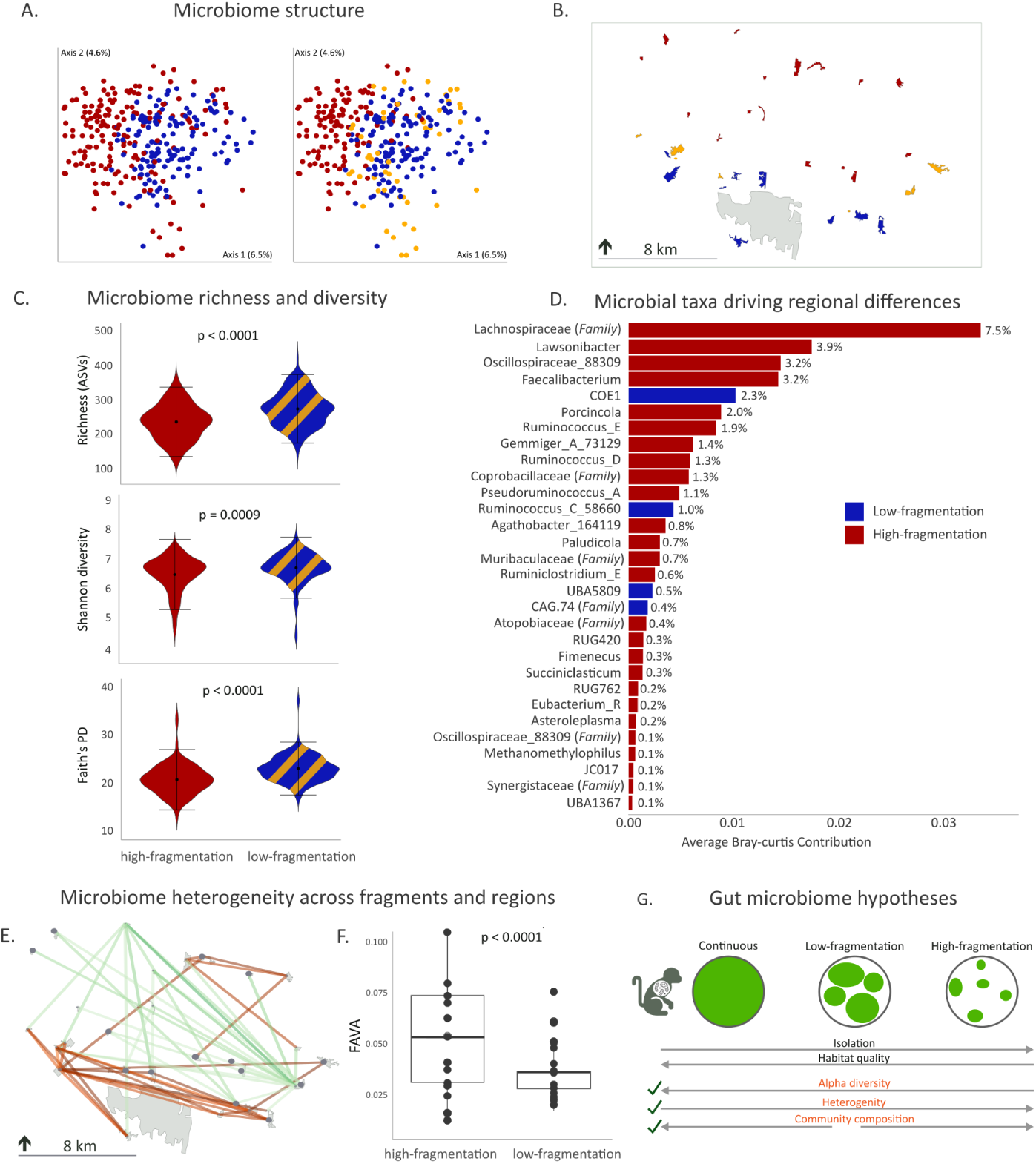
The effect of fragmentation intensity on microbiomes. (A) Bray-Curtis dissimilarity clustering (both panels). Colors denote region of sampling: high-fragmented in red, low-fragmented in blue, and in the right panel the southern high-fragmented sampling locations (adjacent to the low-fragmented region) are colored in yellow. (B) Map of study area with fragments colored corresponding to right panel in (A). (C) Microbiome diversity measures for the two regions compared in the microbiome analysis (N_low-fragmented_=18 fragments; N_high-fragmented_=16 fragments). (D) Top 30 taxa significantly driving differences in microbiome compositions of the two regions. Taxa more abundant in the high-fragmentation region in red, taxa more abundant in the low-fragmentation region in blue. Percentage values denote the total between-region Bray-Curtis dissimilarity index value accounted for by each taxon. X-axis denotes the average contribution of that taxon to the dissimilarity index across all pairs of individuals. See Table S11 for further details on taxonomic classifications. (E) Between-fragment microbiome dissimilarity network. Orange lines denote the top 5% of pairs of fragments with the most similar microbiomes, green lines denote the bottom 5% pairs with the most dissimilar microbiomes; color intensity indicates magnitude. Sampled fragments marked with black circles. (F) Differences in within-fragment microbiome heterogeneity in the two regions. Each point represents the FAVA index value for all individuals in a fragment. (G) Summary of whether results were consistent with expectations for fragmentation-associated microbiome change; consistent patterns are denoted with green check marks.

The taxa driving the divergence between the microbiomes of the high- and low-fragmented regions were indicative of dietary differences. The microbiomes from the high-fragmented region were enriched for taxa known for fiber digestion and short-chain fatty acid production, indicative of a more folivorous diet (e.g., Lachnospiraceae, the greatest contributor to dissimilarity between the regions, *Faecalibacterium*, Oscillospiraceae, *Ruminococcus*^5^). In contrast, the few taxa that were more abundant in the low-fragmented region were mostly associated with sugar digestion, indicative of a more frugivorous diet (e.g., *UBA5809*, *COE1* ; Fig. 4D, Table S12), in line with the higher proportions of known important food trees, and *Ficus* trees specifically, found in fragments in the low-fragmented region (Fig. S11).

Individuals inhabiting fragments in the low-fragmented region, with higher connectivity and better habitat quality, showed gut microbiomes that were overall richer, more diverse, and more homogeneous across individuals than individuals in the high-fragmented region. Individual gut microbiome richness, Shannon diversity, and Faith’s phylogenetic diversity were all significantly higher in the low-fragmented region (Fig. 4C), indicating greater and phylogenically broader taxonomic diversity. Microbiomes in the high-fragmented region were more heterogeneous across individuals than in the low-fragmented region, both within fragments (Fig. 4F), and across fragments (FAVA=0.067 vs. FAVA=0.045 in the high- and low-fragmented regions, respectively, *p* = 0.007; Fig. S13). Network analysis of averaged-by-fragment Bray-Curtis dissimilarity indices showed that the majority of the highest pairwise microbiome dissimilarity values were found in the high-fragmented region, while the most similar pairs were largely found in the low-fragmented region (Fig. 4E, Table S13).

Although fragmentation level was associated with observed differences in gut microbiomes, nested PERMANOVA analyses found that other factors—individual fragment identity, social group, age, and sex—all significantly shaped variation in individual gut microbiomes, with social group showing the largest effect size (Fig. S14, Table S14). These results indicated that local ecological and social variables were strong drivers of microbiome composition, but only after accounting for the overall level of fragmentation (Fig. S14). In contrast, we found no correlation between host genomic distance and microbiome dissimilarity for individuals from different social groups (*N* = 48), indicating that broad patterns of genomic similarity did not shape gut microbiome similarity. To further test the effect of host genetics vs. the current environment on microbiomes, we evaluated whether the microbiomes of likely long-distance dispersers (inferred from genetic data) were more similar to the microbiomes of their sampling location or their putative location of origin. The focal likely-dispersed individuals in 8/11 dyads had microbiomes that were significantly more similar to their current sampling location than their putative point of origin (Fig. S15, Table S15).

Thus, all of our predictions regarding the effects of increasing forest fragmentation on variation in black howler gut microbiomes were supported (Fig. 4G). However, the fact that microbiome patterns did not fully correspond to our connectivity-based partition of the landscape into high- and low-fragmented regions (Fig. 1B; Fig. 4A,B) raises interesting questions regarding the specific drivers of population genetic vs. gut microbiome variation in fragmented landscapes.

### Fragmentation affects population genetics and gut microbiomes via different pathways

Our results clearly indicated that while black howler population genetics and gut microbiomes were both substantially affected by forest fragmentation, they did not respond to increasing fragmentation in similar ways. For example, individual genetic diversity (*H_o_*) was higher in the more fragmented habitat while individual gut microbiome alpha diversity was lower, and monkeys in fragments in the low-fragmented region were more genetically isolated while their gut microbiomes were more similar to each other. To investigate the specific drivers shaping these contrasting patterns, we compared the effects of demography, connectivity and habitat quality variables (Table S3) on genetic and microbiome measures through multivariate analyses (Fig. 5; Figs. S18, S19, Tables S16, S17).

**Figure 5:**
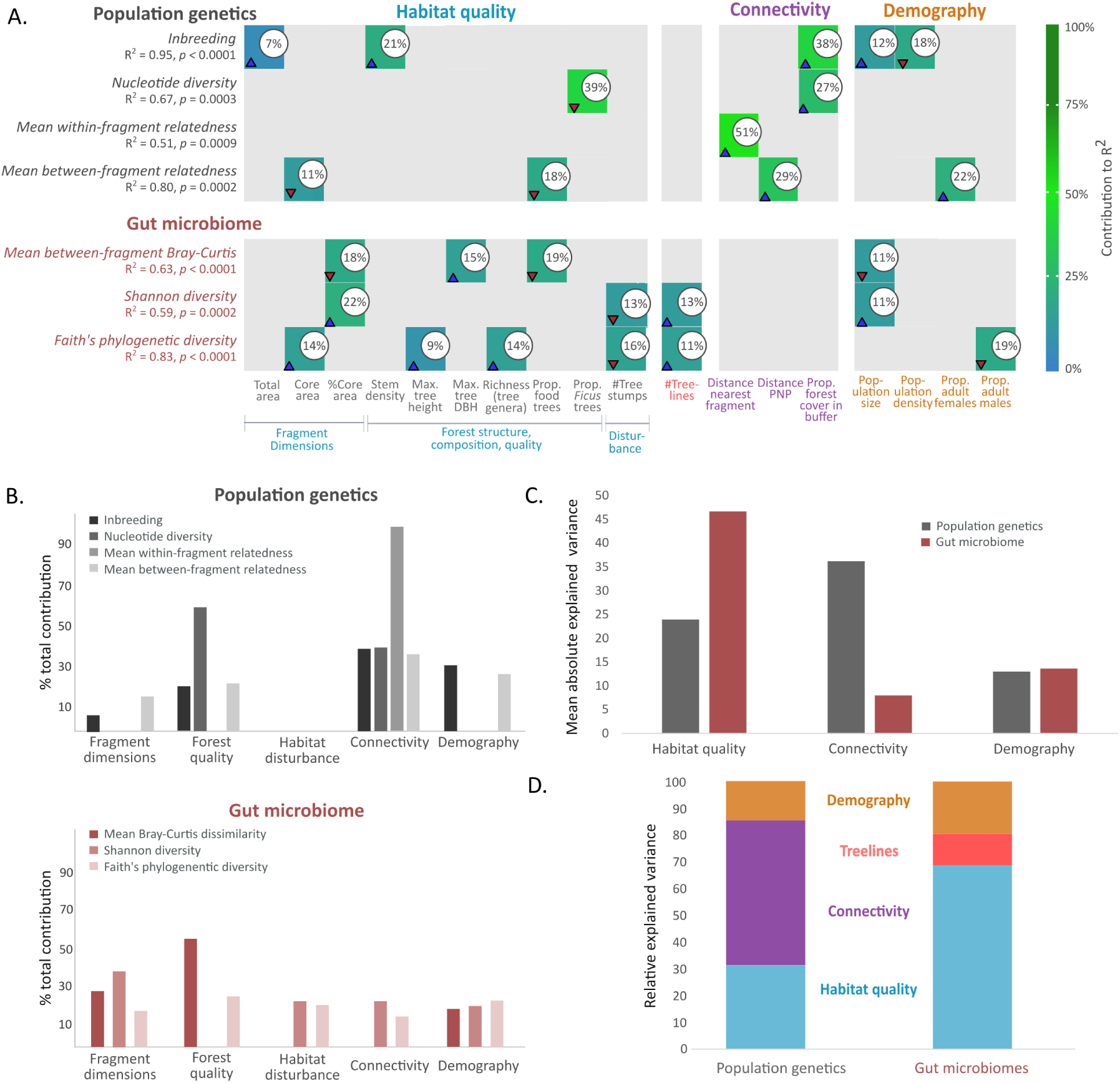
Factors explaining fragmentation effects on genetics and microbiomes. (A) Results of a multivariate analysis of the ecological and demographic variables shaping patterns in population genetics and gut microbiomes. Each line corresponds to a model for a genetic or microbiome metric (p-values and *R*^2^ shown for each model), and each column corresponds to an explanatory variable (only those variables included in at least one multiple regression model shown here). The explanatory variables that significantly contributed to each model are noted with their percentage contribution represented by cell color and text, and the direction of the effect noted with arrows (blue for positive, red for negative). The explanatory variables are assigned categories denoted by colors; ’# treelines’ is both a measure of connectivity and of habitat quality, but is included with connectivity measures in our analysis. (B) Summaries of the contribution of different categories of explanatory variables to variation in population genetic (top) and microbiome metrics (bottom). (C) Mean absolute variance explained by each variable type (habitat quality, connectivity, demography) across all population genetic variables, and across all microbiome variables. (D) Comparisons of the relative proportion of variance explained by each explanatory variable type for population genetic and microbiome metrics. Colors in the bars correspond to the colors of the explanatory variable types in (A); treelines, potentially reflecting both connectivity and habitat quality, is shown separately.

For population genetics, we evaluated several measures. Inbreeding (*F_IS_*) increased in larger fragments, in fragments with larger populations, with lower population density, and with increasing forest cover around the fragment (Fig. 5A), reflecting the high inbreeding coefficients in the continuous forest (Fig. 2D). While nucleotide diversity (*π*) increased with increasing forest cover around the fragment, it decreased with increasing proportions of *Ficus* trees in fragments, further supporting the unexpected association of lower genetic diversity with some measures of higher quality habitat, as *Ficus* are an important food source for howlers^50,64,65^. The relatedness within fragments was strongly positively associated (*R*^2^ = 0.51) with the distance to the nearest fragment (a measure of isolation), indicating that lack of connectivity does limit dispersal in the landscape, although mediated at times by habitat quality. The interplay of habitat connectivity and quality is seen in between-fragment relatedness, a measure of effective dispersal across the matrix, which increased with higher proportions of adult females in the fragment population, highlighting females as a main driver of gene flow across the matrix. Between-fragment relatedness also increased with decreasing proportions of important food trees and fragment core areas, indicating that low habitat quality and availability contributed to more dispersal across the matrix (Fig. 5A), and was positively associated (*R*^2^ = 0.29) with distance from the continuous forest, likely reflecting the increased movement across the matrix in the northern high-fragmented region (Fig. 2, Fig. 3). Thus, variation in genetic measures was largely explained both by connectivity and habitat quality metrics. (Fig. 5C,D).

In contrast, gut microbiome variation was mainly shaped by measures of habitat size and quality, and to a lesser extent demography (Fig. 5A,D). For example, microbiome diversity increased with total fragment population size, and with measures related to habitat size and availability, such as higher proportions of fragment core areas and the number of treelines extending from the fragment (treelines are both a measure of connectivity, and secondary habitat and resources^66,67^). Both microbiome Shannon diversity and phylogenetic diversity decreased with an increasing number of tree stumps, indicating a negative effect of increasing anthropogenic disturbance on howler microbiome diversity. Phylogenetic diversity also increased with increasing habitat quality measures, such as tree richness and larger core areas. Fragments with higher proportions of important food trees had, on average, microbiomes that were more similar to those of other fragments, indicative of less-disturbed diets shaped by known important primary forest food sources. Increasing core areas and fragment population sizes were also positively correlated with microbiome similarity between fragments (Fig. 5A).

We found interesting differences when comparing the effects of the three broad categories of explanatory variables (habitat quality, connectivity, demography) on population genetic vs. microbiome metrics (Fig. 5B,C,D). While demographic variables explained similar proportions of variance in both, they substantially differed in the variance explained by connectivity and habitat quality. Connectivity variables accounted for 54.3% of the variance explained for population genetic metrics, compared to 11.8% for microbiome metrics, suggesting that connectivity is a substantially stronger driver of variation in black howler population genetics than for microbiomes (Fig. 5D). Considering that the only connectivity variable that had a significant effect on microbiome variation was the number of treelines extending from fragments (Fig. 5D), which may reflect the role of treelines as a secondary food source^67,68^ rather than for facilitating movement, the difference in connectivity’s effect on genetics vs. microbiomes is striking. In contrast, while habitat quality drove a substantial proportion of the variation in population genetic patterns (31.2%), its relative contribution to explained gut microbiome variation was considerably larger (68.6%; Fig. 5D). These results suggest that microbiome variation is not shaped directly by reduced connectivity in the fragmented landscape, but rather indirectly, via changing habitat quality and demography in fragments, whereas forest fragmentation affects movement and gene flow in howlers directly, via changes to connectivity, and indirectly, via its effects on habitat quality and demography.

## Discussion

Taking advantage of a natural experiment created by a landscape in which forest fragmentation has progressed unevenly, we analyzed how the population genetics and gut microbiomes of black howler monkeys from the same original population responded to increasing fragmentation. Our results show that these two biological dimensions were both shaped by fragmentation, but through different pathways: population genetic patterns reflected changes in realized movement and gene flow mediated by connectivity, habitat quality, and behavior, whereas gut microbiome patterns primarily reflected habitat degradation and associated dietary and environmental changes.

### Fragmentation effects on gene flow were shaped by connectivity, habitat quality and behavioral responses

We observed that between-fragment connectivity strongly shaped population genetic patterns, through a pathway not captured by simple fragmentation-as-isolation expectations, with longer dispersal distances across the matrix and more gene flow in the most fragmented region. Our finding that population genetic patterns were also driven by habitat quality factors, combined with previous work on the dispersal and demography of these howlers^51,53^, suggests a mechanism. Higher-quality fragments likely support larger populations, shaping reproductive opportunities and dispersal decisions^6,43^, which can lead to reduced incentives for risky dispersal across the matrix. This is reflected in, for example, the negative association between a fragment’s proportion of important food trees and between-fragment relatedness (Fig. 5A). In the low-fragmented region, with better connectivity and higher habitat quality, males were more frequently able to disperse short distances within the fragment or to groups in nearby fragments, as occurs in the continuous forest^53^, leading to steeper male isolation-by-distance and fine-scale genetic structure (Fig. 2A; Fig. 3). In the high-fragmented region, where individuals likely contended with insufficient resources or conspecifics more frequently in more isolated fragments with poorer habitat quality^57^, black howlers’ behavioral flexibility allowed them to engage in risky dispersal across the matrix more often. Extending this reasoning, the continuous forest in the park may be thought of as a very large high-quality fragment, composed of primary and secondary rainforest and supporting a large population wherein dispersal occurs between many stable social groups. This dynamic creates an isolated ecological and genetic island with lower genetic diversity, higher inbreeding, and clear genetic differentiation from other fragments. The limited gene flow between the park and forest fragments also indicates an absence of source-sink dynamics between the park and the fragmented landscape^69^.

Higher levels of fragmentation seem to have been particularly disruptive to males’ regular short-range dispersal pattern^53^, which may explain previous findings of a female-biased adult sex ratio in the fragmented landscape^51^. As in the continuous forest, females in the fragmented regions were more philopatric but traveled farther when dispersing, while males preferentially dispersed to nearby groups. In the high-fragmented region, with fewer within-fragment dispersal opportunities, both sexes appear to have traveled farther and crossed the matrix more frequently. Increased adult male mortality during their more frequent dispersal across the matrix, as has been observed in other species^43^, could explain the female-biased sex ratio in the fragmented region. In this case, females may become a significant driver of gene flow across the matrix, as reflected by the increase in between-fragment relatedness with higher proportions of adult females in fragments (Fig. 5A). Thus, increased gene flow in the most fragmented region should not be interpreted as evidence that fragmentation was benign, but rather as a behavioral response to altered habitat quality and dispersal opportunities, with sex-specific outcomes.

### Gut microbiomes reflected habitat degradation more than reduced connectivity

We found that patterns in gut microbiomes were largely driven by variation in fragment habitat quality, not connectivity. This aligns with studies in other species (e.g., Tome’s spiny rat^32^) showing that gut microbiomes are not directly shaped by connectivity *per se*, but rather by other types of anthropogenic disturbance, causing lower microbiome diversity, altered community composition and increased heterogeneity^32^. Thus, black howlers exemplify the microbiome ’Anna Karenina Principle’^70^—healthy microbiomes are all alike but disturbed microbiomes are all different—where disturbance in this case is generated by fragmentation.

Many of the variables that we identified as significantly driving microbiome spatial structure were indicative of dietary diversity and the availability of high-quality foods. The low-fragmented region had higher proportions of important food trees (Fig. 1D, Fig. S8), and howlers in those fragments likely consumed diets more similar to each other and to howler diets in primary rainforest, contributing to greater microbiome diversity and consistency across individuals and across fragments (Fig. 5A). The microbial taxa shaping dissimilarities between the microbiomes of the low- and high-fragmented regions also suggested dietary differences as a main driver of microbiome variation with increasing fragmentation. Microbiomes of individuals in the high-fragmented region, with lower-quality habitat, were enriched for taxa associated with folivory, reflecting findings of increasing reliance on leaves with increasing habitat disturbance for black howlers at our study site^57^), and a study on black howlers in a different site in Mexico that showed that individuals in lower-quality fragments had higher abundances of many of those same folivory-associated fermenting bacteria (e.g., Ruminococcaceae, Lachnospiraceae, *Faecalibacterium*)^41^.

Many of the microbial taxa significantly driving differences between the low- and high-fragmented regions were rare, and were found only in microbiomes from the high-fragmented region (Table S12). These rare taxa included groups commonly found in soil, leaf litter, or freshwater (e.g., *Blastomonas*^71^,*Pedobacter* ^72^, *Reyranella*^73^) that are atypical of arboreal primate gut microbiomes^5^. This finding may reflect the increased traversal of non-forest matrix by black howlers living in the high-fragmented landscape, which often necessitates movement on the ground, potentially exposing individuals to microbial taxa from domesticated animals, humans, and invasive species^31,32,74^ that they would not normally encounter in the tree canopy.

While differences in methodology preclude detailed comparisons to previous studies examining the gut microbiomes of black howler groups in and around Palenque National Park^33,75,76^, we can discern some interesting trends that match our findings. Previous studies reported that black howler gut microbiomes were uniformly more diverse and consistent over time in the continuous forest than in fragments, which also showed distinct gut microbial community compositions^76^. Howlers occupying degraded habitats consumed less diverse diets and correspondingly had less diverse gut microbiomes, with the gut microbiome richness of groups in the continuous forest twice as high as that found in a group inhabiting a fragment 7 km away^33^. The ratio of fruit to leaf consumption drove differences in gut microbiomes across seasons in the park^75^, in line with our interpretation that regional microbiome differentiation stemmed from higher-quality, more frugivorous diets in the low-fragmented region and lower-quality, more folivorous diets in the high-fragmented region.

The fact that fragmentation mainly affected variation in microbiomes indirectly, via its effects on fragment habitat quality, may explain why microbiome spatial structure did not fully overlap with our original connectivity-based partition of the landscape (Fig. 4B). This mismatch may reflect the different timescales at which decreases in connectivity and reductions in habitat quality occurred as fragmentation progressed^3,77^. Population genetic patterns more closely reflected changing realized connectivity, whereas gut microbiome patterns appeared to reflect fragmentation-mediated habitat degradation, which may emerge as fragments become smaller, more isolated, and lower in quality.

### Conservation implications

Our study leveraged the advantages of fecal-derived DNA, particularly its non-invasiveness and ease of collection, to study an endangered species. However, with fecal samples, DNA quantity and quality may be low, with considerable sample dropout at different stages of the bioinformatic pipeline and large amounts of missing data, which can pose analytical challenges. We were able to address this issue by conducting control tests for the effects of missing data and uneven or small sample sizes throughout our analyses (SI Methods), highlighting the utility of non-invasive fecal samples for the assessment of wild populations.

Our results raise some interesting directions for future work. The higher movement and gene flow we observed in the most fragmented part of the landscape is a snapshot in time, captured from an ongoing process of landscape change. This snapshot indicated that, at least for some species, the effect of habitat fragmentation on dispersal and gene flow may be non-linear^78^. Black howlers’ behavioral and ecological flexibility may allow them to temporarily maintain gene flow despite increasing isolation and habitat degradation by increasing their movement rates and distances, at least up to a point. Some theoretical studies have suggested that in already fragmented landscapes, further habitat loss may induce higher dispersal rates, as the benefits of increased movement across the matrix begin to outweigh the additional costs^23^. Our study provides empirical evidence consistent with these models, and calls for wider recognition of the role of behavioral responses and flexibility in shaping gene flow and population genetics in fragmented landscapes (e.g.,^6,23,78–80^). Our results also highlight the importance of sex-specific effects of fragmentation on dispersal and gene flow. Indeed, the downstream genetic, demographic, and social consequences for species in which male and female dispersal patterns and mortality rates are differentially affected by fragmentation remain understudied^6,67^.

The study of genetic and microbiome effects in parallel enabled a more detailed and nuanced understanding of how fragmentation affects populations, demonstrating the importance and the potential of considering diverse effects when evaluating responses to environmental change. Both genetic and microbiome patterns and spatial structures were altered, but they reflected different direct and indirect pathways through which forest fragmentation affected the population. This contrast highlights the crucial role of species- and landscape-specific attributes in shaping the ultimate effects of fragmentation on wild populations. Here, black howlers’ behavioral flexibility^48,57^ seemed to compensate for increasing isolation by increasing movement and gene flow across the matrix. This alone might lead one to conclude that this species has been able to adapt to its anthropogenically fragmented habitat. However, this genetic signal should not be interpreted as evidence that fragmentation is inconsequential for black howlers, but rather as a compensatory response to habitat reduction and degradation that comes with ecological, physiological, and demographic costs^81,82^, such as a skewed adult sex ratio^51^ and altered and impoverished gut microbiomes. Indeed, considering our gut microbiome findings and additional studies on ongoing habitat loss and altered demography and ecology^47,51,57,58^, this appearance of stability may represent a transient phase in a general downward trend for this fragmented population, particularly if these negative effects continue to accumulate and interact^3^.

## Methods

### Study site and sample collection

Our study site encompassed 39 privately owned, unprotected forest fragments (0.2–36.2 ha, mean = 12 ha) and six opportunistic sampling sites within *∼* 10 km radius of Palenque National Park (PNP), for a total sampling area of 469.3 ha outside the park; we also collected samples from an area of 180 ha within the park (Fig. 1B; Table S1). PNP is a protected area of 1,771 ha, *∼* 900 ha of which is primary and secondary evergreen rainforest^62^. Forest fragments were composed of a mix of remnant primary rainforest, secondary growth forest, and plantations, with varying levels of anthropogenic disturbance. The non-forest matrix was mainly composed of cattle pasture and other forms of agriculture (e.g., corn fields, palm oil plantations; for further details, see^47,51,53,58,62,65^).

In 2015 and 2017–2018 we collected georeferenced (Garmin GPSMAP 78s) fecal samples (*∼* 2 grams material per sample) in the fragmented landscape from individually identified monkeys. Samples were preserved in *∼* 2.5 ml of RNA*later*. The samples from the continuous forest were collected in 2010–2012 using a similar protocol^53,62^.

### Processing and analyses of genetic data

#### Pre-processing and filtering

We extracted host DNA from 297 fecal samples with QIAamp DNA Stool Mini Kits (Qiagen) and Macherey-Nagel Nucleospin Tissue Mini Kits with stool support protocols (Macherey-Nagel), following the manufacturers’ instructions with slight modifications (Table S2; SI Methods). We enriched host DNA following the FecalSeq protocol^83^ with minor modifications, using the NEBNext Microbiome DNA Enrichment Kit (New England BioLabs E2612S). Next, we performed double digest restriction-site associated DNA sequencing (ddRADseq), which combines whole-genome reduced representation sequencing by digestion with two restriction enzymes (SphI and MluCI) and next-generation sequencing for SNP genotype calling^84,85^. The final libraries were sequenced in three batches on an Illumina NovaSeq platform at the University of Calgary’s Center for Health Genomics and Informatics.

Quality control, filtering, alignment, and SNP calling were conducted with FastQC^86^, SAM-tools^87^, BWA^88^, and Stacks^89^ (code for the genetic data processing pipeline currently available upon request). In Stacks, the raw read data were demultiplexed with the *process radtags* module using default parameters and filters and the “–disable rad check” flag, which disables checking if the RAD cutsite is intact, allowing us to retain *>* 95% of reads (SI Methods).

We used the default parameters in BWA-MEM^90^ to align the data to an *Alouatta palliata* reference genome^91^, and performed SNP calling with Stacks^89^ resulting in 13.5% aligned reads and a mean read depth of 32.5±46.8x (range: 5.6x–301.2x). We filtered out individuals with *>* 75% missing data^92^ to maximize the number of loci and minimize missing data per SNP. We then set the following filters in the Stacks *populations* module: maximum 60% missing data per SNP; minor allele count = 3 (i.e., all minor alleles were found in at least two individuals; recommended over minor allele frequency when the dataset has high or variable amounts of missing data^93^); the “write single SNP” filter, which retains only the first SNP per locus; and we included only variant sites found in at least 25% of both continuous forest and fragmented region samples, to ensure that results were not skewed because of insufficient overlap of SNPs in these two areas. Our final genetic dataset included 28 samples from 15 groups in the continuous forest, and 130 samples from 37 locations in the fragmented landscape (Table S1), with a mean of 48.5 *±* 15.0% (range = 21.9–74.3%) missing data per individual.

### Genetic diversity and inbreeding coefficients

We computed population-level genetic diversity and inbreeding coefficients using the Stacks *populations* module for the continuous forest, the fragmented landscape as a whole and for the high- and low-fragmented regions separately. We calculated individual observed heterozygosity (H_o_) with the R package *adegenet* ^94^. For each individual, H_o_ was calculated as the proportion of successfully genotyped SNPs that were heterozygous; missing genotypes were excluded from the individual-specific denominator. Sample-size-corrected expected heterozygosity was calculated separately for each population and SNP using the ’basic.stats’ function in the *hierfstat* package^95^. The resulting within-population gene-diversity estimates (H_S_, reported as corrected H_E_) incorporate a finite-sample correction following Nei^96^ and were averaged without weighting across all SNP loci. Loci that were monomorphic within-population were retained and contributed values of zero to the population-level average.

Individual inbreeding coefficients were estimated using the Lynch–Ritland method-of-moments estimator implemented in COANCESTRY^97,98^. The estimator evaluates each individual’s multilocus homozygosity relative to allele frequencies estimated from the entire pooled dataset of 158 individuals; therefore, individual estimates represent inbreeding relative to the allele-frequency distribution of the complete study sample rather than relative to each region separately. We compared the individual inbreeding and H_o_ values among the three regions (high-fragmented, low-fragmented, and continuous forest) with a Kruskal-Wallis test followed by a *post hoc* Dunn’s test with Holm adjustment for multiple comparisons.

### Population structure

We analyzed population genetic structure with the program STRUCTURE^99^ for *K* = 2–5, using 100,000 burn-in + 500,000 iterations x 20 runs per *K*, with separate alphas estimated from the data for each genetic cluster to account for uneven sample sizes^14^. Results shown here are the consensus solution identified using CLUMPAK^100^ from the synthesis of 20 iterations for each *K* value. To determine if results were being driven by small or uneven sample sizes and sampling locations, or by high levels of missing data, we created and analyzed several subsets of the data in addition to the whole-dataset analyses: (i) a subset produced by more stringent missing data thresholds; (ii) subsets of the fragmented population to be more directly comparable to PNP, including a comparable number of samples, a comparable sampling area, or both. Heterozygosity, inbreeding, and STRUCTURE results with these smaller datasets were very similar to those found for the full dataset (Table S5; Fig. S2).

### Relatedness

We calculated pairwise relatedness coefficients with the Lynch-Ritland relatedness estimator^97^ using the software COANCESTRY^98^ (Fig. S3). A pairwise relatedness value of zero signifies that the two individuals share no more alleles than would be expected by random chance given the allele frequencies in the population. We validated these relatedness values with pedigree information^101^ for five pairs of known closely related individuals (full siblings or parent-offspring) in the well-studied population of the continuous forest^53^. All of these known highly related pairs fell within the top 124 (1%) of pairwise relatedness values for the whole dataset (*r* = 0.17–0.29), providing verification that the Lynch-Ritland relatedness estimates successfully identified them as unusually closely related within the study sample and reflected biological reality, and were not unduly skewed by missing data. Their absolute estimates were lower than the theoretical expectation of *r* = 0.5, reflecting the relative nature of the estimator, the use of pooled allele frequencies across regions, and potential effects of missing genotypes. We quantified differences in patterns of relatedness between regions, socio-spatial categories (within-group, between-group in same fragment/PNP, and between-fragments or between fragments and PNP), and between sexes with two-tailed Mann-Whitney U tests and Kruskal-Wallis tests followed by *post hoc* Dunn’s tests to determine which groups differed significantly in their relatedness (see Table S8 for sample sizes). We combined STRUCTURE results with demographic data^51^ and relatedness coefficients to identify likely interfragment dispersers that were assigned to different STRUCTURE clusters from the major cluster of their sampling location^24^ and were highly related to individuals in a different sampling location with the same major STRUCTURE cluster assignment.

### Gene flow

To evaluate gene flow across the landscape, we conducted an analysis using EEMS (Estimated Effective Migration Surfaces)^102^. EEMS assumes isolation-by-distance as a null model, and visualizes a migration surface showing deviation from this null (regions with higher than expected gene flow rates in blue, lower than expected in brown). We ran the model three times each for 100, 200, 300, 400, 500, 600, 700, and 800 demes, for a total of 24 runs, with an MCMC length of 5,000,000, burn-in of 2,000,000, and thinning rate of 9,999. Results presented in the main text are the merged composite of the three runs at a resolution of 600 demes, which showed both clear MCMC chain convergence and a good model fit (Fig. S5). Results were visualized using the R package *reemsplots2* (https://github.com/dipetkov/reemsplots2).

To visualize gene flow across the landscape as represented by likely individual dispersal events, we used the COANCESTRY Lynch-Ritland pairwise relatedness values as input to create a network of genetic relatedness across the landscape with the python library *networkX* ^103^. To highlight patterns of very high or low relatedness across the landscape rather than within fragments or social groups, we considered only between-fragment pairs and included only the highest (in orange) and lowest (in green) 0.25% of pairwise relatedness values (N_Total_=58; Table S7). Increasing edge color intensity denoted increasing absolute values of relatedness.

### Isolation-by-distance

To quantify isolation-by-distance (IBD) within each region (Palenque National Park, low-fragmented, high-fragmented) and for males and females, we modeled pairwise genetic distance as a function of geographic distance using maximum-likelihood population effects (MLPE) models implemented as linear mixed-effects models with the *lme4* package in R^104^. We calculated genetic distance as (1 *− r*) (Lynch–Ritland relatedness) for all dyads and regressed this against pairwise Euclidean distance (km), while accounting for repeated sampling of individuals (as each individual appears in multiple dyads). We tested IBD significance by comparing a distance model (genetic distance *∼* geographic distance) to a null model without distance using a likelihood ratio test. The fixed-effect slope for geographic distance represents the rate of genetic distance increase per kilometer, and its sign and magnitude quantify the strength of spatial genetic structure; associated 95% confidence intervals and p-values indicate whether the slope differs from zero. For more direct comparisons with the park, we applied this approach to subsets of the data from the high- and low-fragmented regions, including only pairs within the park’s smaller spatial extent (*∼* 2 km; Table S9).

To compare the strength of IBD between regions and sexes, we extended the MLPE framework to include group effects and interactions with geographic distance, using *lme4* ^104^, *emmeans*^105^, and *lmerTest* ^106^ in R. Differences in IBD strength were tested using likelihood ratio tests comparing models with and without the interaction term; a significant interaction indicated that the slope of genetic distance on geographic distance differed among groups (i.e., different IBD strength). Landscape- or sex-specific slopes were then estimated using marginal trend contrasts, and pairwise differences between slopes were evaluated with Tukey-adjusted comparisons. The associated standard error, 95% confidence interval, and adjusted p-value quantified estimate precision and whether slopes differed significantly between groups.

We conducted finer-scale spatial autocorrelation tests with the Lynch-Ritland relatedness coefficient in SPAGeDi (Spatial Pattern Analysis of Genetic Diversity)^107^. We ran iterations on several subsets of the full dataset (e.g., by region or sex; SI Methods), which introduced variation in the Lynch-Ritland coefficients, the number of permutations (range: 40-499), and the number of SNPs (range: 2230-8454; Table S11). We used identical distance classes across all datasets in Palenque National Park, and identical distance classes across all datasets in the fragmented landscape. In all iterations, the first distance class was defined as the social group.

### Processing and analyses of microbiome data

#### Pre-processing and filtering

We amplified and sequenced the v4-v5 16S region of the ribosomal RNA gene from bacterial DNA extracted from fecal samples. We assessed the gut microbiome to the amplicon sequence variant (ASV) level for 326 individuals from the fragmented landscape, in two batches (see Tables S1, S2, SI Methods for further details on sample sizes, sequencing batches, and the final combined dataset).

We extracted DNA using the commercial DNeasy PowerLyzer Powersoil Kit (Qiagen, Germantown, MD, USA) with modifications (SI Methods). A two-step PCR was used to amplify the V4–V5 region of the 16S rRNA gene, utilizing the 515 forward and 926 reverse Earth Microbiome Project primers (www.earthmicrobiome.org), as described previously^108,109^. Sequencing of barcoded amplicons was performed on an Illumina MiSeq V4 platform by the Rush University Genomics and Microbiome Core Facility.

After each batch of samples was sequenced, we performed quality control, processing, filtering, and ASV identification steps in Qiime2^110^ to ensure the two batches were of comparably high quality, and to assess possible batch effects (code for the microbiome data processing pipeline currently available upon request). PCoA plots showed no sorting by sequencing batch (Fig. S10); thus, we pooled together the raw sequencing data and proceeded with the combined dataset of 312 unique samples, repeating quality control, filtering, and processing steps in Qiime2. We generated 13,034,743 sequences with an average of 41,778 sequences per sample (range = 15,731–70,322 sequences per sample).

With DADA2^111^ in Qiime2, we trimmed forward reads to 273 bp and reverse reads to 220 bp, using a base call quality score of 25 as the cutoff. We applied the the Qiime2 classifier to the Greengenes2^112^ database to identify ASVs. We detected 11,315 unique ASVs (total frequency = 3,382,023; average per sample = 10,840, range per sample = 3,528–19,171). Based on per-sample rarefaction plots for richness, Shannon diversity, and Faith’s phylogenetic diversity (Fig. S9), we rarefied the data to 5000 sequences per sample, excluding five additional samples with *<* 5000 sequences. Our final dataset included 307 unique individuals from 34 locations in the fragmented landscape only (Tables S1, S2), and 9472 ASVs. All analyses were conducted at the taxonomic level of ASV unless noted otherwise.

### Richness, diversity, and heterogeneity

For this rarefied dataset, we used Qiime2 to calculate per-sample richness, Shannon diversity, and Faith’s phylogenetic diversity (PD). A higher Faith’s PD indicates greater evolutionary diversity, while lower values suggest more taxonomically uniform communities. In Qiime2, we created by-individual PCoA plots for four beta diversity indices: Jaccard, accounting for composition only; Bray-Curtis, accounting for composition and abundance; unweighted Unifrac, accounting for composition and phylogenetic distances among microbial taxa; and weighted Unifrac, accounting for composition, abundance, and phylogenetic distances. We used the summary table output from Qiime2 to create an *a priori* averaged by-fragment microbiome composition summary table and used *picante*^113^ in R to calculate averaged by-fragment Faith’s PD, and *phyloseq* ^114^ in R to calculate averaged by-fragment richness and Shannon diversity, and to calculate the same four beta diversity indices for the averaged by-fragment dataset.

Similar to the genetic relatedness network (see “Genetic data processing and analyses”), we used the averaged by-fragment Bray-Curtis dissimilarity index to visualize the network of microbiome similarity with the python library *networkX* ^103^. To highlight patterns of high or low microbiome similarity across the fragmented landscape, we set an inclusion threshold of the highest and lowest 5% of Bray-Curtis values (N_Total_=54). We excluded fragment 28, as it emerged as an outlier with a highly dissimilar microbiome from most other fragments (Table S13). To facilitate comparisons between this and the genetic network, here high-value edges were colored in green (denoting lower similarity between fragment microbiomes) and low-value edges were colored in orange (denoting higher similarity). Increasing edge color intensity denoted increasing absolute values.

We used the FAVA (F_ST_-based Assessment of Variability across vectors of relative Abundances) index^115^ to quantify the compositional variability across a group of microbiome samples, where higher values imply more microbiome heterogeneity across samples. We compared the heterogeneity of microbiomes of individual fragments (aggregating the microbiomes of all individuals sampled in each fragment; SI Methods), and we compared the heterogeneity of high-fragmented (*N* = 138) vs. low-fragmented (*N* = 169) microbiomes by pooling all individuals in each region. We used the FAVA R package^115^ to test the significance of differences in regional FAVA values by drawing 1,000 bootstrap replicates of the microbiome samples from the two regions under the null hypothesis that there is no true difference between them.

### Drivers of between-region microbiome variation

To determine which microbial taxa were driving differences between microbiomes in the high- and low-fragmented regions, we ran a SIMPER (“similarity percentages”) analysis with the *vegan* package^116^ in R. SIMPER takes the Bray-Curtis dissimilarity indices between each high-fragmented -low-fragmented pair of individuals and calculates each microbial taxon’s average abundance in each region, average contribution to the microbiome dissimilarity across all between-region pairs of individuals, and the statistical significance of each taxon’s contribution to dissimilarity between the regions. We ran this analysis for all 307 individuals, at the microbial taxa level of genus.

We conducted nested PERMANOVA analyses with the ’adonis2’ command in the *vegan* package^116^ in R to determine the specific contributions of region (low- and high-fragmented), fragment ID, social group, age category (adult, sub-adult, juvenile, infant), and sex to variation in individual microbiomes. Nested PERMANOVAs account for the variables’ hierarchical structure, such that each lower-level variable is evaluated within the context of the higher-level variables. The response variable was a microbiome dissimilarity matrix, and we ran nested PERMANOVAs for all four dissimilarity indices for the entire dataset together, and then for each region separately, with 999 permutations to determine significance.

We had both genetic and gut microbiome data for 92 individuals, sampled from 24 fragments and two opportunistic sampling locations (Tables S1, S2). We conducted Mantel correlation tests with the *vegan* package^116^ in R to assess the relationship between individual microbiome dissimilarity and genetic distance, calculated as (1 *− r*) (Lynch–Ritland relatedness), with a subset of the data including only one individual from each social group to control for the confounding effects of horizontal transmission and similar diets. We selected the adult with the lowest proportion of missing genetic data from each group for inclusion in this subset of 48 individuals, representing 45 groups and three solitary individuals from 24 sampling locations. We used the Spearman correlation coefficient and 9,999 permutations to determine significance.

To further examine the effects of host genetics on gut microbiomes, we conducted a distance-to-centroid analysis with permutation tests (N=999) with the *vegan* package^116^ in R. Specifically, we tested whether the gut microbiomes of inferred dispersers were significantly more similar to gut microbiomes in their sampling location or in their putative location of origin, as computed with the Bray-Curtis dissimilarity index; putative locations of origin were inferred by combining the results of our population genetic structure and pairwise genetic relatedness analyses with demographic data. From the dataset of 92 individuals with both genetic and microbiome data, we included pairs of individuals sampled in different fragments, with Lynch-Ritland relatedness values *>* 0.1, and for which both sampling locations and putative locations of origin included at least three individuals (excluding the focal individual, i.e., the inferred disperser), to allow for the calculation of a centroid. Our final dataset included 11 pairs of a likely disperser and a related individual from the inferred location of origin (Table S15). We plotted Bray-Curtis dissimilarity values in a two-dimensional PCoA plot, and computed the centroids and centroid coordinates of each group (excluding the focal individual from its sampling location’s group centroid). We calculated and tested the significance of the distance from the focal individual to each of the two group centroids with permutation tests.

### Identifying habitat and demographic drivers of genetic and microbiome patterns

Beyond identifying and describing patterns in population genetics and gut microbiomes of black howlers across levels of fragmentation in the study landscape, we also aimed to determine the specific habitat quality, isolation, and demographic variables that shaped these patterns. We focused on the fragmented landscape only, and analyses were done at the fragment, not individual level, to match the scale of the explanatory variables (Tables S3, S16).

### Population genetic response variables

We included four population genetic variables: (i) per-fragment average inbreeding, as calculated in the *populations* module of Stacks^89^; (ii) per fragment nucleotide diversity (*π*), as calculated in the *populations* module of Stacks^89^; (iii) mean within-fragment relatedness, derived from pairwise Lynch-Ritland relatedness coefficients^97^; and (iv) mean relatedness to other fragments, also derived from pairwise Lynch-Ritland relatedness coefficients^97^. This dataset included the 18 fragments with habitat quality data and genetic data from at least two individuals (mean number of samples per fragment=4.8±3.9, range=2-14; number of sampled groups per fragment = 1-6), for a total of 86 samples (Tables S1, S16; see SI Methods and Methods subsection ’LASSO variable selection and multiple regression’ for details on sensitivity analyses to address small and variable per-fragment sample sizes).

### Gut microbiome response variables

We included three gut microbiome variables: (i) per-fragment Shannon diversity, calculated with the *phyloseq* package^114^ in R; (ii) per-fragment Faith’s PD, calculated with the *picante* package^113^ in R; and (iii) mean Bray-Curtis microbiome dissimilarity index for that fragment with all other fragments, calculated with the *phyloseq* package^114^ in R. This dataset included the 28 fragments that had habitat quality data and microbiome data from at least two individuals (mean number of samples per fragment=9.9±7.1, range=2-31; number of sampled groups per fragment = 1-6), for a total of 276 samples (Tables S1, S16).

### Explanatory variables

We tested the effects of the same set of 26 variables on both the genetic and the microbiome response variables, including six variables describing fragment demography (see^51^ for details on demographic data collection). The demographic variables included total fragment population size, which also acted as a proxy for per-fragment sample size, as they were significantly, highly positively correlated (Fig. S18; Tables S3, S16 for full details of explanatory variables).

We calculated 20 variables that quantified forest fragment dimensions, isolation, and within-fragment habitat quality (Table S3; for detailed descriptions, see^47,58^). Briefly, for each surveyed fragment (N=39), we quantified fragment area, shape, isolation distance from the nearest fragment and from the continuous forest, and the number of treelines extending from the fragment, by examining high-resolution (1 m^2^) satellite images in Google Earth© (2018), combined with on-the-ground validation during fragment surveys. We calculated an additional area-based isolation measure, the proportion of forest habitat present within a buffer zone of a 2.4 km radius around fragments^58^. We used FRAGSTATS^117^ with the “PatchStat” function in the *SDMTools*^118^ package in R to calculate fragment core area (the area within each fragment unaffected by edges) and fragment core area index (core area / total patch area), based on the forest cover layer we used to calculate proportions of forest cover around fragments^119^ (30 m^2^ resolution; Table S3). From December 2017 – April 2018, we collected data on forest structure and composition (e.g., mean tree height, stem density, tree diversity) in 29 fragments using a modified Gentry protocol^47,58^. From these transect data, we also calculated three additional variables: the percentage of important food trees per fragment, derived from comparing the genera observed in each fragment to the 12 tree genera that each accounted for *>* 1% of time spent feeding in black howlers in the continuous forest^65^; the proportion of *Ficus* trees per fragment, as *Ficus* are known to be a particularly important food source for howlers,^50^ and for black howlers in Palenque National Park specifically^64^; and the Bray-Curtis dissimilarity index for tree genus composition for each pair of fragments with the “vegdist” command in *vegan*^116^ (Table S3).

### LASSO variable selection and multiple regression

We performed LASSO+AICc variable reduction and selection followed by multiple regression analyses to identify the drivers of variation across the landscape in black howler population genetic and gut microbiome patterns. Because the analyses were based on 18 or 28 forest fragments (for population genetic and gut microbiome variables, respectively; Table S1), and included a comparatively large set of potentially correlated fragment-level habitat, connectivity, and demographic explanatory variables, we incorporated robustness assessments to determine whether variable selection was disproportionately influenced by particular fragments, cross-validation partitions, or plausible measurement error in the explanatory variables. The following variable selection procedure was conducted separately for each population genetic or gut microbiome response variable.

First, we performed an exhaustive fragment-deletion sensitivity analysis by removing every possible pair and every possible trio of fragments. For the population genetic variables, this generated 153 pair-deletion datasets containing 16 fragments and 816 trio-deletion datasets containing 15 fragments, for a total of 969 reduced datasets; for microbiome variables, this generated 378 pair-deletion datasets containing 26 fragments and 3,276 trio-deletion datasets containing 25 fragments, for a total of 3,654 reduced datasets. Within each reduced dataset, predictors were standardized using the ’scale’ function in the R package *base*^120^. Variable selection was performed using the least absolute shrinkage and selection operator regression method (‘LASSO’^121^), implemented using the ’cv.glmnet’ function of the *glmnet* R package^122,123^, combined with the Akaike Information Criterion (AICc), following^124^. LASSO penalizes redundant predictors while minimizing the penalized regression error, reducing the influence of multicollinearity by favoring a subset of informative variables and shrinking redundant coefficients toward zero^121^. The LASSO penalty was selected using five-fold cross-validation and the value of *λ* that minimized cross-validation error.

For each reduced dataset, we performed 100 paired analyses. In each pair, one analysis used the original predictor values and the other used independently perturbed predictor values, with multiplicative error of up to 5%. The same cross-validation fold assignment was used for both members of each pair, allowing differences between them to reflect the introduced predictor error rather than differences in cross-validation partitioning. When the cross-validated LASSO selected more than seven predictors, we moved to the nearest stronger penalty along the LASSO path that retained no more than seven predictors. These predictors were entered into a linear-model candidate set, and all subsets were ranked using the small-sample corrected AICc with the ’dredge’ function of the *MuMIn* R package^125^; the lowest-AICc model was retained.

For each reduced dataset, predictor variables were considered robust if they were retained in more than a third of the best–fitting models both with and without introduced error in predictor variables. We then calculated, separately for the original and error-perturbed data, the proportion of reduced datasets in which each predictor met this criterion. A predictor variable was classified as consistently selected when it was retained in over 33% of the reduced datasets, i.e., *≥* 323 reduced datasets for population genetics variables and *≥* 1, 218 reduced datasets for microbiome variables (although in practice, 21/26 selected variables were retained in over 50% of the reduced datasets; Fig. S19). These selection frequencies were interpreted as measures of robustness.

These robust predictors were then included in the subsequent multiple regression model for that response variable. We estimated the contribution of each predictor to the model’s explained variance by multiplying its normalized contribution (its standardized regression coefficient divided by the sum of the absolute values of all coefficients) by the *R*^2^ value. Lastly, we evaluated whether different categories of explanatory variables (habitat quality, connectivity, and demography; Fig. 5, Tables S3, S16) had substantially different magnitudes of effect on population genetic and microbiome metrics. We compared (i) the mean absolute variance explained by habitat quality, connectivity, and demography across the four population genetic and the three microbiome metrics, respectively, and (ii) the relative contribution of each explanatory variable type to the total explained variation for population genetics and gut microbiomes, such that the relative contributions sum to 100.

## Data Availability

Raw sequencing data generated in this study have been deposited in the NCBI Sequence Read Archive (SRA) under BioProject PRJNA1526035. The 16S rRNA gut microbiome data are deposited under SRA accessions: SAMN63026212 - SAMN63026518, and the host ddRADSeq data under SRA accessions: SAMN63192716 - SAMN63192873. These datasets will be released publicly upon publication of this article. Code used for data processing, statistical analyses, and generation of the results and figures presented in this study will be available here upon publication: https://github.com/Greenbaum-Lab/Black-howler-forest-fragmentation-genetics-and-microbiomes.

## Supporting information

All supplementary information (figures, tables, and expanded Methods section)

## Acknowledgments

We thank A. Lazar, A. Cardenas, D. Levey, S. Mellon, C. Alley, T. Pinho, E. Barrios, and D.A. Campos Villanueva for their assistance in the field. KK expresses her deepest thanks to A. Lazar for countless other forms of support during this research. We most particularly thank the local landowners and Mayan communities who provided access to the study fragments. We thank S. Lehman and A. Estrada for logistical and research support and for helpful discussions and comments during the early stages of this research, F. Mercado Malabet for assistance with GIS, and H. Hamou for assistance with the genetics lab work. We thank the Mexican government (CONANP and INAH) for granting research permission to work in and around PNP. This work was supported by the National Autonomous University of Mexico’s Institute of Biology, the Natural Sciences and Engineering Research Council of Canada’s Vanier Canada Graduate Scholarship, the University of Toronto School of Graduate Studies, the University of Toronto School of the Environment’s Beatrice and Arthur Minden Research Fellowship, the University of Toronto Scarborough Research Competitiveness Fund, the Azrieli International Postdoctoral Fellowship, the Minerva Center for the Study of Population Fragmentation (MCPF), and Bi-National Science Foundation (BSF) Grant No. 2021244. KRA is supported as a fellow in CIFAR’s ’Health Microbiomes in a Changing World’ Program.

## Author contributions

Conceptualization: KK, GG; Study design: KK, SVB, JAT, KRA, GG; Extractions and sequencing: KK, ECW, GD, MLSS, AD, KRA, ADM; Coding: KK, KDH, RP, KRA, AMG, OP; Analyses: KK, RP, OP, AMG, MLM, MS, KRA, KDH, GG; Writing of Original Draft: KK; Review and Editing: KK, SVB, JAT, ECW, GD, MLSS, KDH, OP, RP, AMG, MLM, MS, AD, KRA, ADM, GG; Funding Acquisition: KK, GG, JAT, SVB, ADM, KRA, AD.

