## Supplementary material for "Contrasting effects of forest fragmentation on the genetics and microbiomes of an endangered arboreal primate": All supplementary information (figures, tables, and expanded Methods section)

### Expanded Methods

#### **Population genetics**

##### Sample processing, library prep, sequencing, and SNP calling

We extracted and sequenced host DNA from the fecal samples of 297 individuals in three batches: batch #1 included 88 adult and subadult samples from 35 locations in the fragmented landscape, collected throughout the sampling period; batch #2 included a total of 126 adult, subadult, and juveniles from 41 sampling locations in the fragmented landscape, collected throughout the sampling period (15 of these samples were re-sequenced after failing sequencing in batch #1); and batch #3 included 98 adult, subadult, juvenile, and infant samples from 18 groups in PNP (Table S2).

We extracted DNA with QIAamp DNA Stool Mini Kits (Qiagen) and Macherey-Nagel Nucleospin Tissue Mini Kits with stool support protocols (Macherey-Nagel), following the manufacturers' instructions with slight modifications. We ran PCRs on ~50% of sample extractions for the microsatellite locus LL1115, isolated from *Lagothrix* (Di Fiore and Fleischer, 2004), together with positive and negative controls, to confirm the presence of primate host DNA. We measured final DNA concentrations with a Qubit 2.0 fluorometer and Qubit dsDNA HS Assay Kits (Invitrogen) and we also performed droplet digital PCR (ddPCR) with the Evagreen Master Mix (BioRad #1864034). We used primers targeting the mammalian sequence (C-myc). The C-myc forward primer (5' GCCAGAGGAGGAACGAGCT 3') and reverse primer (5' GGGCCTTTTCATTGTTTTCCA 3') amplified a region of the host cell DNA. We generated droplets using a ddPCR system (BioRad QX200), followed by amplification with the following cycling conditions: initial denaturation at 95°C for five minutes, 40 cycles of denaturation at 95°C for 30 seconds, annealing at 59°C for one minute, extension at 72°C for 30 seconds, then signal stabilization at 4°C for five minutes and at 90°C for five minutes. We used the ddPCR system to quantify the DNA concentration by analyzing the fluorescence from individual droplets.

We enriched the host DNA using the NEBNext Microbiome DNA Enrichment Kit (New England BioLabs E2612S). We followed the FecalSeq (Chiou and Bergey, 2018) protocol with minor modifications, namely, reducing the 1× BW buffer wash volume from 1 ml to 200 µl to conserve reagents, using a 1.8× SPRI bead (Cytiva, Fisher scientific #09981123; these beads were used throughout the whole protocol) purification ratio, and increasing the ethanol wash volume from 100 µl to 200 µl to ensure complete coverage of the bead pellet. This enrichment procedure captures eukaryotic DNA by using a methylated CpG-specific binding domain protein fused to the Fc fragment of human IgG (MBD2-Fc) to selectively target sequences with high CpG methylation density. We prepared MBD2-Fc-bound magnetic beads according to the manufacturer's instructions. To elute the hDNA, we added 100 µl of 2M NaCl to each tube off the magnetic rack and gently pipetted. We incubated the mixture in a rotating mixer for three minutes at room temperature. After 2-5 minutes on the magnetic rack, we carefully transferred the supernatant into a new tube. We then performed x1.8 bead clean-up on the eluted hDNA. Another ddPCR was performed on the eluted DNA after the enrichment to double check the concentration of the hDNA using the same settings as before.

Next, we performed double digest restriction-site associated DNA sequencing (ddRADseq), which combines whole-genome reduced representation sequencing by digestion with two restriction enzymes and next-generation sequencing (NGS). This approach allowed us to obtain thousands of single nucleotide polymorphisms (SNPs) and genotype dozens of samples simultaneously (Aguirre et al., 2023; Peterson et al., 2012). For double digestion, we diluted the 1x stock of restriction enzymes (SphI and MluCI) to 1/10 using the appropriate digestion buffers. We diluted SphI with Diluent B and MluCI with Diluent A. We prepared a total reaction volume of 30 µL by mixing 24 µL of DNA with the restriction enzymes. We incubated the reaction at 37°C for three hours and then held it at 4°C. After incubation, we allowed the reaction to cool to room temperature and then performed x1.5 bead purification. The ligation of the adapter barcodes was done by using the NEB T4 DNA ligase (# M0202), where the reaction was incubated for 30 minutes at 23°C, heat-inactivated it for 10 minutes at 65°C, and then held it at 4°C. We combined equal amounts of ligated DNA from each sample to create a pool of individuals with unique barcodes, where the first barcode was five bases and the second barcode was 6 bases; we used 12 different first barcodes and nine different second barcodes, for a total of 108 unique barcodes across all three sequencing batches. We cleaned the double digest with 1.5x SPRI beads and eluted it in 35 µL of H2O.

We then performed a PippinPrep for size selection to ensure the DNA fragments were within the desired size range. We combined approximately 20 ng of the size-selected sample with PCR primers P1 and P2 at a final concentration of 2 µM each, along with the recommended amount of 5X-HF buffer, dNTPs, water, and Phusion polymerase in a standard 200 µL PCR tube. After completing the PCR, we pooled the reactions and cleaned them with 1.5x SPRI beads, eluting the DNA in 35 µL of H2O. The final libraries were sequenced at 2 x 151 bp with 6 bp single indexing on the Illumina NovaSeq platform at the University of Calgary's Center for Health Genomics and Informatics.

Quality control, filtering, alignment, and SNP calling were conducted for all three batches with FastQC (Andrews, 2010), SAMtools (Danecek et al., 2021), BWA (Li and Durbin, 2009), and Stacks (Catchen et al., 2013; code for full genetic data processing pipeline can be found at: https://github.com/Greenbaum-Lab/Black-howler-forest-fragmentation-genetics-and-microbiomes). First, the raw read data were demultiplexed with the *process_radtags* module in Stacks using default parameters and filters to remove reads with an uncalled base, discard reads with low quality scores, and rescue reads with barcodes or RADtags with an error in a single base. In all three batches, we found that while few reads were discarded due to low quality (<0.2% in each batch) or problems identifying barcodes (<5% in each batch), a large proportion of reads were dropped because the RAD cutsite was not found (batch 1: 67.5%; batch 2: 77.5%; batch 3: 62.5%), leading to an overall very low proportion of initial sequenced reads being retained (20-35%). We therefore processed all batches using the "--disable_rad_check" flag, which disables checking if the RAD cutsite is intact, allowing us to retain 95% of reads in batch 1 and 97% of reads in batches 2 and 3.

We used the default parameters in BWA-MEM (Li, 2013) to align the data to an *Alouatta palliata* reference genome (*A. palliata* Genome, 2019), and performed SNP calling with the *gstacks* module in Stacks. The percentage of reads aligned and effective per-sample coverage were similar across the full datasets for all three batches (batch #1: 10.2% of reads aligned; 321,211 loci genotyped; effective per-sample coverage: mean=12.9x, SD=9.9x, min=1.7x, max=53.3x; batch #2: 11.4% of reads aligned; 474,762 loci genotyped; effective per-sample coverage: mean=13.3x, SD=9.7x, min=2.2x, max=68.4x; batch #3: 10.2% of reads aligned; 216,828 loci genotyped, effective per-sample coverage: mean=16.0x, SD=9.8x, min=1.6x, max=69.8x). Given this similarity in data quality, we pooled all samples into a single dataset at the post-alignment phase, and conducted SNP calling in Stacks with the *gstacks* and *populations* modules for this combined dataset, which had 13.5% aligned reads and a mean read depth of 32.5±46.8x (range: 5.6x-301.2x). It should also be noted that, given that the PNP samples (batch #3) had similar, if not better, sequencing quality and coverage as compared to batches #1 and #2, the lower number of polymorphic sites found in PNP in downstream analyses (see Results section) almost certainly reflects lower genetic diversity, as opposed to sequencing batch effects. We filtered out individuals with > 75% missing data (Cerca et al.,

2021) to maximize the number of retained loci and minimize missing data per SNP. We then set the following filters in the Stacks *populations* module: maximum 60% missing data per SNP; minor allele count = 3 (i.e., all minor alleles were found in at least two individuals); the "write single SNP" filter, which retains only the first SNP per locus; and we included only variant sites found in at least 25% of both continuous forest and fragmented region samples, to ensure that results were not skewed because of insufficient overlap of SNPs in these two areas. Our final genetic dataset included 28 samples from 15 groups in the continuous forest, and 130 samples from 37 locations in the fragmented landscape (Table S1), with a mean of 48.5±15.0% (range = 21.9-74.3%) missing data per individual.

Population genetic analyses

We created several subsets of the data to test the potential effects of uneven/small sample sizes on population genetic structure, genetic diversity, and inbreeding results. First, we created a subset produced by more stringent missing data thresholds: maximum 55% missing data per individual and 40% missing per SNP. Second, to test the possible effect of uneven sample sizes and sampling areas on population-level heterozygosity and inbreeding results, we also ran several iterations with subsets of the fragmented population, either with a comparable number of samples, a comparable sampling area, or both. Summary population genetic statistics and STRUCTURE (Pritchard et al., 2000) results with these smaller datasets were very similar to those found for the full dataset (Table S5; Fig. S2).

We computed population-level genetic diversity and inbreeding coefficients using the Stacks *populations* module for the continuous forest, the fragmented landscape as whole and for the high- and low-fragmented regions separately. We calculated individual observed heterozygosity ($H_o$) with the R package *adegenet* (Jombart 2008), as the number of successfully typed SNP loci at which the individual was heterozygous divided by the total number of successfully typed SNP loci for that individual. Consequently, each individual's estimate was based on that individual's available genotype calls rather than on the total nominal number of SNPs in the dataset. For each population, individual $H_o$ values were summarized using the mean, standard deviation, median, interquartile range, minimum, and maximum. The individual, rather than the SNP locus, was treated as the independent sampling unit for comparisons among populations. Because individual $H_o$ values are bounded proportions, population sample sizes were unequal, and a normal distribution was not assumed, the overall comparison of individual $H_o$ among the three populations was performed using the Kruskal–Wallis rank-sum test. Following the overall comparison, Dunn's rank-based procedure (Dunn, 1964) was used for all three pairwise population comparisons, and pairwise P-values were adjusted using Holm's (Holm, 1979) sequentially rejective procedure to control the family-wise type I error rate across the three comparisons. All statistical tests were two-sided, and adjusted $P < 0.05$ was considered statistically significant.

We calculated population-level expected heterozygosity ($H_E$) with the 'basic.stats' function in the R package *hierfstat* (Goudet, 2005). The $H_S$ statistic produced by 'basic.stats' represents within-population gene diversity, commonly reported as expected heterozygosity. The implementation follows Nei's gene-diversity framework (Nei, 1987) and applies a finite-sample correction based on the number of genotyped individuals and the observed genotype frequencies at each locus. This correction was used because the three populations differed substantially in sample size. Corrected $H_E$ was initially calculated separately for every population-by-SNP combination. Population-level corrected $H_E$ was then obtained as the unweighted mean of the locus-specific estimates across all 9567 SNPs. Monomorphic loci were included and contributed an expected heterozygosity of zero. Loci with missing genotypes were calculated using the individuals successfully genotyped at that locus within the relevant population. Corrected $H_E$ was reported descriptively by population, because it is a population-level estimate derived from allele frequencies rather than an independently measured individual characteristic.

Individual inbreeding coefficients were estimated using the Lynch–Ritland method-of-moments estimator (Lynch and Ritland, 1999) implemented in COANCESTRY (Wang, 2011). This marker-based estimator quantifies whether an individual exhibits greater or lower multi-locus homozygosity than expected from the allele frequencies of a defined reference population. At each informative locus, the individual's genotype is evaluated relative to the frequencies of its constituent alleles, and locus-specific information is combined across the SNP panel using weights related to estimator precision. Positive estimates indicate greater homozygosity than expected from the reference allele frequencies, values near zero indicate homozygosity close to the reference expectation, and negative estimates indicate greater heterozygosity than expected.

Allele frequencies were estimated from all 158 individuals pooled across the high-fragmented, low-fragmented, and continuous-forest regions. Thus, the complete study sample constituted the reference population for every individual, and the resulting coefficients measure relative inbreeding against the pooled allele-frequency distribution. Because the reference combines individuals from three geographic regions, the estimates may reflect not only recent parental relatedness but also regional allele-frequency differentiation or population substructure. They should therefore be interpreted as relative marker-based inbreeding coefficients rather than direct estimates of pedigree inbreeding.

We analyzed population genetic structure with the program STRUCTURE for K=2-5, 100,000 burn-in + 500,000 iterations x 20 runs per K, with separate alphas. The alpha parameter controls the prior distribution of admixture proportions, i.e., the average degree of admixture expected in populations. By default, STRUCTURE uses a single alpha value for all populations, but when populations have uneven sample sizes, an identical alpha can skew results, biasing individual assignment probabilities (Wang 2016). Allowing each population to have its own alpha enables STRUCTURE to adjust their priors independently, thereby accounting for differences in sample size and admixture levels between populations, preventing overestimation of admixture in smaller populations, and improving the reliability of population assignments, particularly when sample sizes are uneven (Wang 2016). We merged results of STRUCTURE iterations with CLUMPAK (Kopelman et al. 2015). Results shown here are the consensus solution identified by CLUMPAK from the synthesis of 20 iterations for each value of K.

Relatedness patterns within social groups, between social groups within the same habitat fragment, between individuals sampled in different fragments, and patterns that differ between males and females can provide considerable information regarding movement and gene flow, and how these might vary across different levels of fragmentation in the landscape. We calculated pairwise relatedness coefficients and individual inbreeding coefficients with the Lynch-Ritland relatedness estimator (Lynch and Ritland, 1999) in COANCESTRY (Wang 2010). The Lynch-Ritland relatedness estimator quantifies genetic relatedness based on shared alleles across all loci, calculating the probability that two individuals share alleles identical by descent (IBD) by comparing their observed genotypes to the population allele frequencies (Lynch and Ritland, 1999). A pairwise relatedness value of zero signifies that the two individuals share no more alleles than would be expected by random chance given the allele frequencies in the population, whereas negative relatedness values indicate that pairs are less related than would be expected from random mating.

We validated these relatedness values with pedigree information (Hauser et al., 2022) for five pairs of closely related individuals (full siblings or parent-offspring) in PNP (Van Belle and Di Fiore 2021). The Lynch-Ritland pairwise relatedness estimates for all of these known highly related pairs fell within the 124 highest pairwise relatedness values for the whole dataset, or the top 1%, providing a strong indication that the pairwise relatedness estimates calculated with this method successfully detected unusually closely related pairs within the study sample, reflecting biological reality, and were not unduly skewed by missing data. However, their absolute estimates were lower than the theoretical expectation of $r$=0.5, likely reflecting the relative nature of the estimator, the use of pooled allele frequencies across regions, and potential effects of missing genotypes.

We quantified differences in patterns of relatedness between different regions, sexes, and socio-spatial categories (within-group, between-group in same fragment/PNP, and between-fragments or between fragments and PNP) with two-tailed Mann-Whitney U tests and Kruskal-Wallis tests followed by *post hoc* Dunn's tests to determine which groups differed significantly in their relatedness (see Table S8 for sample sizes). We combined STRUCTURE results with demographic (Klass et al., 2020A) and relatedness data to identify likely inter-fragment dispersers. These individuals were assigned to different STRUCTURE clusters from the major cluster of their sampling location (Bertrand et al., 2017), and were highly related to individuals in a different sampling location (the putative location of origin) with the same major STRUCTURE cluster assignment.

To quantify isolation-by-distance (IBD) within each region (PNP, LF, HF) and for males and females, we modeled pairwise genetic distance as a function of geographic distance using maximum-likelihood population effects (MLPE) models implemented as linear mixed-effects models with the *lme4* package in R (Bates et al., 2015). We calculated genetic distance as $1-r$ (Lynch–Ritland relatedness) for all dyads and regressed this against pairwise Euclidean distance (km). Because pairwise observations are non-independent (each individual appears in multiple dyads), standard linear regression would violate independence assumptions. MLPE models address this by including crossed random intercepts for both individuals in each pair ($1|id_1$ and $1|id_2$), which appropriately accounts for repeated sampling of individuals and controls Type I error while retaining full pairwise information. We tested IBD significance by comparing a distance model (genetic distance ~ geographic distance) to a null model without distance using a likelihood ratio test. The fixed-effect slope for geographic distance represents the rate at which genetic distance increases per kilometer, and its sign and magnitude quantify the strength of spatial genetic structure; associated 95% confidence intervals and p-values indicate whether the slope differs from zero.

To facilitate more direct comparisons of the patterns seen in PNP to those in the fragmented landscape, we applied this approach to subsets of the data from the HF and LF regions, including only pairs within the same finer-scale spatial extent covered by our data for the population in PNP (~2km; Table S9). For subsets with small sample sizes where random-effect variances approached zero and mixed models became singular, we simplified the model by removing random effects and fitting an ordinary least-squares regression (genetic distance ~ geographic distance). In these cases, inference focused on the same slope parameter and its confidence interval, providing a conservative and numerically stable test of within-region IBD while acknowledging reduced power. This approach allowed consistent estimation of IBD strength across datasets while accommodating differences in sample size and dyadic structure.

To compare the strength of isolation-by-distance among regions and between sexes, we extended the MLPE framework to include group effects and interactions with geographic distance, using *lme4* (Bates et al., 2015), *emmeans* (Lenth and Piaskowski, 2025), and *lmerTest* (Kuznetsova et al., 2017) in R. Specifically, we fit linear mixed-effects models of the form genetic distance ~ geographic distance × group (where group represented landscape, sex, or landscape × sex), again including crossed random intercepts for both individuals in each dyad to account for non-independence of pairwise observations. Differences in IBD strength were tested using likelihood ratio tests comparing models with and without the interaction term; a significant interaction indicates that the slope of genetic distance on geographic distance differs among groups (i.e., different IBD strength). Landscape- or sex-specific slopes were then estimated using marginal trend contrasts, and pairwise differences between slopes were evaluated with Tukey-adjusted comparisons. For each group, the estimated slope represents the rate of increase in genetic distance per kilometer, while the associated standard error, 95% confidence interval, and adjusted p-value quantify the precision of the estimate and whether slopes differ significantly between groups. This approach enables direct statistical comparison of IBD magnitude across regions and sexes while appropriately accounting for the shared individuals among dyads.

We conducted finer-scale spatial autocorrelation tests with SPAGeDi (Spatial Pattern Analysis of Genetic Diversity; Hardy and Vekemans, 2002). SPAGeDi takes raw genetic data and individual spatial coordinates as input, and can compute several relatedness coefficients, including Lynch-Ritland, which we used for this analysis as well. As we ran iterations of tests in SPAGeDi on subsets of the full dataset (i.e., separately for PNP and the fragmented landscape, for the HF and LF regions, and by sex), a different subset of individuals and SNPs were used for each iteration, making the LR coefficients slightly different each time; similarly, the number of permutations ranged from 40-499, and the number of SNPs from 2230-8454 (Table S11). We used identical distance classes across all datasets in PNP, and identical (but different from PNP, given the different spatial extent) distance classes across all datasets in the fragmented landscape, to facilitate comparisons. In all iterations the first distance class was defined as the social group.

To visualize gene flow across the landscape as represented by likely individual dispersal events, we used the Lynch-Ritland pairwise relatedness values as input to create a network of genetic relatedness across the landscape. We used the python library *networkX* (Hagberg et al., 2008) to create a network in which nodes represented individuals and the edges connecting them represented the pairwise relatedness value between those individuals. To highlight patterns of very high or low relatedness across the landscape rather than within fragments or social groups, we considered only between-fragment pairs and set an inclusion threshold of the highest and lowest 0.25% of pairwise relatedness values ($N_{Total}$=58; Table S7). Orange and green edges denoted pairs with relatedness values in the top and bottom 0.25% of the dataset, respectively, and increasing edge color intensity denoted increasing absolute values of relatedness.

We also quantified and visualized gene flow across the landscape with the program EEMS (Estimated Effective Migration Surfaces; Petkova et al., 2016). EEMS assumes IBD as the null model and produces visualizations that deviate from IBD either with high effective migration rates (blue areas on the map, signifying areas of high gene flow) or low effective migration rates (brown areas on the map, signifying areas of low gene flow, indicating genetic isolation and/or the presence of barriers to movement and gene flow). Using PLINK (Purcell et al., 2007), we created a .bed file from the .ped and .map files generated by *populations* in STACKS. Then, we used *bed2diffs* from the EEMS package (Petkova et al., 2016) to create a dissimilarity matrix. EEMS requires that this input matrix of genetic dissimilarity be a valid full-rank distance matrix with no missing genotypes. We therefore imputed missing genotypes with the observed mean genotype at the corresponding SNP, by multiplying that SNP's observed allele frequency by two. The extent of the area to be analyzed (i.e., the outer coordinate file), was a circular shape encompassing all of the sampling sites in our landscape with an additional buffer of ~1-5km beyond sampling sites. We ran the model three times each for 100, 200, 300, 400, 500, 600, 700, and 800 demes, for a total of 24 runs.

For each number of demes, we first optimized the variance parameters by tweaking parameter values to achieve the recommended proposal acceptance rates of 20-30% (Petkova et al., 2016). We then ran the additional two chains for each number of demes, with random seeds, with an MCMC length of 5,000,000, burn-in of 2,000,000, and thinning rate of 9,999. Results presented in the main text are the merged composite of three chains at a resolution of 600 demes, which showed both clear MCMC chain convergence and a good model fit (Fig. S15). We visualized results in R using *reemsplots2* (https://github.com/dipetkov/reemsplots2), with the following plots: (i) mrates02 (posterior probability contours for the given probability level): as migration rates are visualized on the log10 scale after mean centering, in this contour plot, mapped onto the study site, zero corresponds to the overall mean migration rate and effective migration that is significantly higher or lower than the overall average is emphasized; (ii) qrates02: similar to mrates02 but applied to the effective diversity rates, not migration; (iii) rdist01: scatter plot of the observed vs the fitted between-deme component of genetic dissimilarity, where one point represents a pair of sampled demes and singleton demes are excluded from the plot; (iv) rdist02: scatter plot of the observed vs the fitted within-deme component of genetic dissimilarity, where one point represents a sampled deme and singleton demes are excluded; and (v) posterior trace plot: this plot visualized the convergence of different MCMC chains (Petkova et al., 2016). After determining that the MCMC chains converged with the posterior trace plot, we used rdist01 and rdist02 to determine the fit of the EEMS models; a strong linear relationship between the observed and fitted values in both scatter plots indicates that the model fit the data well. We merged and visualized the results of all chains for all deme numbers with the *reemsplots* package (https://github.com/dipetkov/eems/tree/master/plotting; see Fig. S16 for plotted results of merging all chains for 100-800 demes).

#### Gut microbiomes

##### Sample processing, sequencing, and ASV identification

We amplified and sequenced the v4-v5 16S ribosomal region of the RNA gene from bacterial DNA extracted from fecal samples to assess the gut microbiome to the amplicon sequence variant (ASV) level for 326 individuals (code for microbiome processing pipeline available at: https://github.com/Greenbaum-Lab/Black-howler-forest-fragmentation-genetics-and-microbiomes). Samples were sequenced in two batches containing similar representations of sexes, sampling locations, sample collectors, and sampling periods, i.e., the sample pools differed only in sequencing batch and a higher proportion of both adults and infants in batch #1. Batch #1 consisted of 242 samples and batch #2 consisted of 140 samples (see Tables S1, S2 for details on the per-location demography and sample sizes in each sequencing batch and the final combined dataset).

We extracted DNA from all samples using the commercial DNeasy PowerLyzer Powersoil Kit (Qiagen, Germantown, MD, USA) with modifications. Briefly, after adding solution C1 and the beads provided, the samples were incubated at 65 °C for 10 min before vortexing for 10 min. The samples in solution C2 and C3 were placed at 4°C for 5 minutes. Additionally in the elution step, we warmed solution C6 at 65°C before adding it. All samples were quantified by using the Eppendorf spectrophotometer and a total amount of 50ng of DNA were used in the PCR reaction. A two-step PCR was used to amplify the V4–V5 region of the 16S rRNA gene, utilizing the 515 forward and 926 reverse Earth Microbiome Project primers (www.earthmicrobiome.org), as described previously (Walters et al. 2016; Mallott and Amato 2018). PCR products were purified and normalized using SequalPrep Normalization. Sequencing of barcoded amplicons was performed on an Illumina MiSeq V4 platform by the Rush University Genomics and Microbiome Core Facility with a depth of at least 20,000 sequences per sample. Negative controls from DNA extractions and PCRs were included in the initial data set and used as negative controls for contamination.

After each batch of samples was sequenced, we performed quality control, processing, filtering, and ASV identification steps in Qiime2 (Bolyen et al., 2016) to ensure the two batches were of comparable and high quality, and that results were not being shaped by batch effects. While batch #1 contained more data overall (even after accounting for the larger sample size) and per sample than batch #2 (batch #1: total number of sequenced features = 2,716,363, per-sample median = 12,088; batch #2: total number of sequenced features = 888,189, per-sample median = 6639), the two datasets were of very similar quality: similar percentages of reads per-sample passed initial quality control filters (batch 1: mean percentage of reads retained = 76.5 ± 2.2%, range = 66.7-81.8%; batch 2: mean percentage of reads retained = 75.0 ± 3.1%, range = 66.1-80.2%), and following denoising and chimera filtering steps, the final percentages of reads retained per sample was also very similar (batch 1: mean percentage of reads retained = 25.2 ± 3.7%, range = 17.3-33.9%; batch 2: mean percentage of reads retained = 27.7 ± 5.6%, range = 16.9-52.4%). Additionally, PCoA plots showed no sorting by sequencing batch (Fig. S10). Thus, after removing from each batch negative controls, samples that failed sequencing (two in batch #1 and one in batch #2), samples with uncertain IDs, and duplicates of other high-quality samples, we pooled together the raw sequencing data for the remaining 196 samples from batch #1 and 116 samples from batch #2 and proceeded with this combined dataset of 312 unique samples, repeating quality control, filtering, and processing steps in Qiime2. We generated 13,034,743 sequences with an average of 41,778 sequences per sample (range = 15,731-70,322 sequences per sample).

With DADA2 (Callahan et al., 2016) in Qiime2, we trimmed the forward reads to 273 bp and the reverse reads to 220 bp, using a base call quality score of 25 as the cutoff. We applied the Qiime2 classifier to the Greengenes2 (McDonald et al., 2024) database to identify ASVs. The total number of unique ASVs detected was 11,315 (total frequency = 3,382,023; average per sample = 10,840, range per sample = 3,528- 19,171). We produced per-sample rarefaction plots for richness, Shannon diversity, and Faith's phylogenetic diversity (Fig. S9), and based on these plots we rarefied the data to 5000 ASVs per sample. We thereby excluded an additional five samples with <5000 ASVs, for a final dataset of 307 unique individuals and a total of 9472 ASVs that we used in all subsequent analyses. All analyses were conducted at the taxonomic level of ASV except where noted otherwise.

Gut microbiome analyses

For this rarefied dataset, we used Qiime2 to calculate per-sample alpha diversity with richness, Shannon diversity, and Faith's PD. Faith's PD quantifies the biodiversity of microbial communities by considering the evolutionary relationships between species; it sums the branch lengths of a phylogenetic tree that connects all the species in a community, providing a measure of the total evolutionary history represented by that community. Thus, a higher Faith's PD indicates greater evolutionary diversity, while lower values suggest more taxonomically uniform communities. We also used Qiime2 to create by-individual PCoA plots for four beta diversity indices: Jaccard, which accounts for composition only; Bray-Curtis, which accounts for composition and abundance; unweighted Unifrac, which accounts for composition and phylogenetic distances among microbial taxa; and the weighted Unifrac, which accounts for composition, abundance, and phylogenetic distances. We used the summary table output from Qiime2 to create an *a priori* averaged-by-fragment microbiome composition summary table and used *picante* (Kembel et al., 2010) in R to calculate averaged by-fragment Faith's PD, and *phyloseq* (McMurdie and Holmes, 2013) in R to calculate averaged by-fragment richness and Shannon diversity, and to calculate the same four beta diversity indices for the averaged by-fragment dataset.

Similar to the genetic relatedness network (see "Genetic data processing and analyses"), we used the averaged by-fragment Bray-Curtis dissimilarity index as input to visualize the network of microbiome similarity across the fragmented landscape with the python library *networkX* (Hagberg et al., 2008). Here, nodes represented fragments and the edges connecting them represented the Bray-Curtis microbiome dissimilarity value between those fragments. For clarity, and to highlight patterns of high or low microbiome similarity across the landscape, we set an inclusion threshold of the highest and lowest 5% of Bray-Curtis values ($N_{Total}$=54). We excluded fragment 28 from this visualization, as it emerged as an outlier with a highly dissimilar microbiome from most other fragments (Table S13). Unlike in the genetic relatedness network, where high values denote higher relatedness and low values denote lower relatedness, with the Bray-Curtis dissimilarity index, high values denote less similar microbiomes and low values denote more similar microbiomes. Therefore, to facilitate comparisons between the networks, in this network high-value edges were colored in green (denoting lower similarity) and low-value edges were colored in orange (denoting higher similarity). Increasing edge color intensity denoted increasing absolute values (either high or low).

We used the FAVA ($F_{ST}$-based Assessment of Variability across vectors of relative Abundances) index (Morrison et al., 2025) to quantify the compositional variability across a group of gut microbiome samples. FAVA is a statistic that measures the variability in microbiome composition across a group of two or more microbiome samples, equaling 0 if all samples have identical composition and equaling 1 if each sample is comprised entirely of a single taxon. Higher values of FAVA imply more heterogeneity in microbiome composition across samples. We compared the heterogeneity of microbiomes of individual fragments, aggregating the microbiomes of all individuals sampled in each fragment. The FAVA index does not explicitly control for sample size; while smaller sample sizes may make FAVA noisier, they do not systematically bias the index in a particular direction, because it works with relative abundances and does not depend on the number of taxa or the number of samples one computes across.

We also pooled all individuals in each region to compare the heterogeneity of microbiomes in the HF region (N=138) vs. the LF region (N=169). We used the FAVA R package (Morrison et al. 2025) to test the significance of the differences in FAVA values between regions by drawing 1,000 bootstrap replicates of the microbiome samples from the HF or LF regions under the null hypothesis that there is no true difference between the two categories.

To determine which microbial taxa were driving the differences between the microbiomes in the HF and LF regions, we ran a SIMPER ("similarity percentages") analysis with the *vegan* package in R (Oksanen et al., 2016). SIMPER takes the Bray-Curtis dissimilarity indices between each HF-LF pair of individuals and calculates each microbial taxon's average abundance in each region, average contribution to the microbiome dissimilarity across all between-region pairs of individuals, and the statistical significance of each taxon's contribution to dissimilarity between the regions. We ran this analysis for all 307 individuals, at the microbial taxa level of genus, not ASV.

We conducted nested PERMANOVA analyses with the adonis2 command in the *vegan* (Oksanen et al., 2016) package in R to determine the specific contributions of region (LF/HF), fragment, social group, age, and sex to variation in individual black howler gut microbiomes. In a nested PERMANOVA, the hierarchical structure of the variables is accounted for, such that the variation is partitioned out sequentially according to the non-independent levels of the explanatory variables. Each lower-level variable is evaluated within the context of the higher-level variables, i.e., they are not independent of the higher-level variables. For example, the effect of social group is dependent on and evaluated within the context of the specific fragment, which is evaluated within the context of the region. Because of this dependence, the marginal effect for the whole nested structure is evaluated together. This approach does not provide individual variance for each variable because it assumes the lower-level variables (like group and age/sex) only have meaning within the context of the higher-level variables. The response variable was a microbiome dissimilarity matrix, and we ran nested PERMANOVAs for all four dissimilarity indices for the entire dataset together, and then for each region separately, with 999 permutations to determine significance.

We conducted Mantel correlation tests with the *vegan* (Oksanen et al., 2016) package in R to assess the relationship between individual microbiome dissimilarity and (i) Euclidean distance; and (ii) genetic distance. All Mantel tests were conducted with the Spearman correlation coefficient and 9,999 permutations to determine significance. To test the relationship between microbiome dissimilarity and distance, we created the pairwise Euclidean distance matrix with the *geosphere* package in R (https://github.com/rspatial/geosphere), based on each individual sample's collection location coordinates. We ran Mantel tests for the full dataset, as well as for a subset of the data (N=68) that included only one individual from each social group and all solitary individuals, to control for the possible confounding effects of horizontal transmission, very similar diets, and higher genetic relatedness within groups (Fig. S16). We also tested the relationship between microbiome dissimilarity and Euclidean distance for the HF (N=119) and LF (N=188) regions separately, and for the one individual per group subset of the data in each region (HF = 25 individuals; LF = 43 individuals; Fig. S16). Lastly, we conducted a Mantel test to assess the relationship between averaged-by-fragment microbiome dissimilarity and the Bray-Curtis dissimilarity index for fragment forest composition, and ran additional iterations of this test for each region separately (HF=13 fragments; LF=15 fragments; Fig. S17).

To test the relationship between microbiome dissimilarity and genetic distance, we calculated pairwise genetic distance as 1-(pairwise Lynch-Ritland relatedness coefficient), for the dataset of 92 individuals with both microbiome and genetic data, who represented 45 groups from 24 sampling locations (Tables S1, S2). We ran a second iteration of this latter Mantel test with a subset of the data, including only one individual from each social group to control for the possible confounding effects of horizontal transmission and very similar diets. We selected the adult with the lowest proportion of missing genetic data from each group for inclusion in this subset, which included 48 individuals representing 45 groups and three solitary individuals.

To further test the effects of host genetics on gut microbiomes, we conducted a distance-to-centroid analysis with permutation tests (N=999) for significance, with the *vegan* package in R (Oksanen et al., 2016). We wanted to assess if the gut microbiomes of putative dispersers were significantly more similar to gut microbiomes in their sampling location, or in their putative location of origin, as computed with the Bray-Curtis dissimilarity index; putative locations of origin were inferred by combining the results of our population genetic structure and pairwise genetic analyses with demographic data. Of the 92 individuals for which we had both genetic and microbiome data, here we included pairs of individuals sampled in different fragments, with Lynch-Ritland relatedness values >0.1, and for which both sampling locations and putative locations of origin included at least three individuals (excluding the focal individual, i.e., the inferred disperser whose microbiome was being compared to the two locations), to allow for the calculation of a centroid. This left us with 11 pairs, where each pair included a likely disperser and an individual from the inferred location of origin (Table S15). We plotted Bray-Curtis dissimilarity values in a two-dimensional PCoA plot, and computed the centroids and centroid coordinates of each group (excluding the focal individual from the group where it was sampled when computing that group's centroid). We then calculated the distance in the ordination space from the focal individual to each of the two group centroids; if the distance was positive, the focal individual's microbiome was more similar to the sampling group, if negative it was more similar to the microbiome at the putative location of origin. We then tested the significance of the focal individual's distance to each group centroid with permutation tests.

#### **Identifying habitat and demographic drivers of genetic and microbiome patterns**

Beyond identifying and describing patterns in population genetics and gut microbiomes of black howlers across levels of fragmentation in the study landscape, we also aimed to determine the specific forest fragment habitat quality, isolation, and demographic variables that were shaping changes in each of these aspects of black howler biology. For these analyses, we focused on a handful of genetic and gut microbiome response variables, and all analyses were done at the fragment, not individual level, to match the scale of the explanatory variables (see Methods, main text; Table S3 for detailed description of all explanatory variables; Table S16 for full dataset).

##### LASSO variable selection and multiple regression

We performed LASSO+AICc variable reduction and selection followed by multiple regression analyses to identify the drivers of variation across the landscape in black howler population genetic and gut microbiome patterns. Because the analyses were based on only 18 or 28 forest fragments (for population genetic and gut microbiome variables, respectively; Table S1), an apparent association could potentially be driven by a small number of fragments with unusual habitat characteristics, demographic composition, population genetic values, or gut microbiome profiles. Additionally, we aimed to test a comparatively large set of potentially correlated fragment-level habitat, connectivity, and demographic explanatory variables. We therefore developed a combined fragment-deletion, cross-validation, and predictor-perturbation sensitivity analysis. Its purpose was to identify predictors that were selected consistently across alternative compositions of the fragment dataset and under modest uncertainty in the explanatory variables. This procedure allowed us to distinguish associations that were broadly supported across fragments from associations dependent on particular fragments, particular cross-validation partitions, or small changes in the measured explanatory variables. The following variable selection procedure was conducted separately for each fragment-level population genetic or gut microbiome response variable.

First, we performed an exhaustive fragment-deletion sensitivity analysis by removing every possible pair and every possible trio of fragments. For the population genetic variables, this generated 153 pair-deletion datasets containing 16 fragments and 816 trio-deletion datasets containing 15 fragments, for a total of 969 reduced datasets; for microbiome variables, this generated 378 pair-deletion datasets containing 26 fragments and 3,276 trio-deletion datasets containing 25 fragments, for a total of 3,654 reduced datasets. The pipeline verified that all removal combinations were unique and that the expected numbers of pair and trio deletions were generated. Removing all combinations of two and three fragments provided a more stringent evaluation than a conventional leave-one-out analysis. In particular, it allowed us to identify results that remained stable when multiple potentially influential fragments were absent simultaneously, including situations in which two or more fragments might exert joint leverage on the fitted relationship.

Within each reduced dataset, each ecological and demographic predictor variable was standardized using the 'scale' function in the R package *base* (R Core Team, 2021). Variable selection was performed using the least absolute shrinkage and selection operator regression method (‘LASSO’; Tibshirani 1996), implemented using the 'cv.glmnet' function of the *glmnet* R package (Friedman et al., 2021; Tay et al., 2023), combined with the Akaike Information Criterion (AICc), following (Marami Milani e al., 2016). LASSO penalizes redundant predictors while minimizing the penalized regression error, reducing the influence of multicollinearity by favoring a subset of informative variables and shrinking redundant coefficients toward zero (Tibshirani 1996). The LASSO penalty was selected using five-fold cross-validation and the value of $\lambda$ that minimized cross-validation error.

For each reduced dataset, we performed 100 paired analyses. Given the variability of forest fragment habitat even within small spatial scales and the potential for human error when collecting demographic data on wild animals, we wanted to account for estimation errors to evaluate the robustness of our results against potential inaccuracies. Thus, in each paired analysis, one analysis used the original predictor values and the other used independently perturbed predictor values, where random multiplicative error of up to 5% was introduced independently into every explanatory-variable value, while the response variable was left unchanged. Thus, the direction and magnitude of the perturbation varied independently among fragments and variables, with a value-specific maximum error bound between 1% and 5% and a realized absolute perturbation between 0% and 5%. This procedure was intended to represent modest uncertainty in fragment-level ecological and demographic measurements rather than to simulate a specific empirically estimated error distribution.

The same cross-validation fold assignment was used for both members of each pair, allowing differences between them to reflect the introduced predictor error rather than differences in cross-validation partitioning. At the beginning of each iteration, fragments were assigned randomly to five balanced cross-validation folds. Pair-deletion datasets contained three or four fragments per fold, whereas trio-deletion datasets contained three fragments in each fold. The fold assignment changed among iterations but was shared between the original-data and error-perturbed analyses within each iteration. Because exhaustive all-subsets model comparison increases exponentially with the number of predictors, and because only 15/16 or 25/26 fragments remained after deletion, when the cross-validated LASSO selected more than seven predictors, we moved to the nearest stronger penalty along the LASSO path that retained no more than seven predictors. These predictors were entered into a linear-model candidate set, and all subsets were ranked using the small-sample corrected Akaike information criterion, AICc, with the 'dredge' function of the *MuMIn* R package (Barton, 2025); the lowest-AICc model was retained. This sensitivity analysis was implemented using a custom function in R developed for analyses aimed at identifying drivers of variation between populations in the genomic signatures of MHC and the whole genome.

For each reduced dataset, predictor variables were considered robust if they were retained in more than a third (33/100 iterations) of the best–fitting models both with and without introduced error in predictor variables. We then calculated, separately for the original and error-perturbed data, the proportion of reduced datasets in which each predictor met this criterion. A predictor variable was classified as consistently selected when it was retained in over 33% of the reduced datasets, i.e., ≥323 reduced datasets for population genetics variables and ≥1,218 reduced datasets for microbiome variables (although in practice, in all but 5/26 cases selected variables were retained in over 50% of the reduced datasets; Fig. S19). The final result for each predictor variable was therefore a count ranging from 0 to 969 for population genetic response variables and 0 to 3654 for gut microbiome variables, representing the number of alternative pair- or trio-deletion datasets in which that variable displayed a minimum level of repeated selection. Large values indicated that the variable was retained across many alternative fragment compositions, whereas low values indicated that its selection was sensitive to the inclusion of particular fragments, cross-validation partitions, predictor perturbations, or combinations of these factors. These frequencies were interpreted as descriptive measures of predictor variable selection robustness, not as posterior probabilities, conventional confidence levels, or formal tests of statistical significance.

These robust predictors were then included in the subsequent multiple regression model for that response variable. To calculate the proportional contribution of each predictor to the overall slope, we first standardized the predictor variables and fit a linear model in which the response variable was regressed to the standardized predictors. We then extracted the coefficients and normalized each by the absolute sum of the slopes. Furthermore, we calculated the percentage contribution of each variable to $R^2$ by multiplying its normalized contribution by the value of $R^2$. We performed linear regression analyses for each variable retained in at least 33% of the reduced dataset iterations and that were also found to be significant in the multiple regression analyses to clarify the relationship between each explanatory variable and the respective response variable individually, to isolate the strength and direction of each variable's effect (Fig. S20).

Lastly, we evaluated whether different categories of explanatory variables (habitat quality, connectivity, and demography; Fig 5, Tables S3, S16) had substantially different magnitudes of effect on population genetic and microbiome metrics. We did not conduct significance testing; given the small number of response variables (four for population genetics and three for gut microbiomes), permutation tests and bootstrapping would be underpowered and sensitive to individual values. We compared (i) the mean absolute variance explained by habitat quality, connectivity, and demography across the four population genetic and the three microbiome metrics, respectively, and (ii) the relative contribution of each explanatory variable type to the total explained variation, such that the relative contributions sum to 100 for population genetics and gut microbiomes, respectively.

### Summaries of datasets

**Table S1. Summary data for sampling locations**. Fragment size, regional classification, and basic demographic information regarding each sample location in the fragmented landscape; and basic demographic information for each group in PNP, as well as information on sample sizes per sampling location for genetic and microbiome analyses and inclusion in LASSO + multiple regression analyses. Acronyms: HF – high fragmented; LF – low fragmented; MB – microbiome; GEN – genetic; OSL – opportunistic sampling location.

| | sampling location/group ID | general description | | | | | | genetics dataset | | microbiome dataset | | in both datasets | | vegetation transects | Included in LASSO + multiple regression analyses |
|---|---|---|---|---|---|---|---|---|---|---|---|---|---|---|---|
| | | size (ha) | # groups surveyed | # solitary ind. | total group/ pop. size | Region | Revised region for microbiome | # groups | # ind. | # groups | # ind. | # groups | # ind. | | |
| Fragmented landscape | 1 | 28.7 | 3 | | 18 | LF | LF | 3 | 4 | 3 | 12 | 3 | 4 | Y | GEN, MB |
| | 2 | 8.1 | 1 | | 7 | LF | LF | \ | \ | 1 | 7 | \ | \ | Y | MB |
| | 3 | 2.61 | 1 | | 8 | LF | LF | 1 | 2 | 1 | 6 | 1 | 2 | Y | GEN, MB |
| | 4 | 3.67 | 1 | | 3 | HF | LF | 1 | 2 | 1 | 2 | 1 | 2 | Y | GEN, MB |
| | 6 | 3.59 | 1 | | 5 | HF | HF | \ | \ | 1 | 4 | \ | \ | N | |
| | 9 | 8.26 | 1 | | 7 | HF | HF | \ | \ | 1 | 7 | \ | \ | Y | MB |
| | 10 | 12.4 | 2 | | 8 | HF | HF | 1 | 1 | 2 | 8 | 1 | 1 | Y | MB |
| | 12 | 13.3 | 2 | | 15 | HF | HF | 2 | 3 | 2 | 14 | 2 | 3 | Y | GEN, MB |
| | 13 | 26.1 | 3 | 1 | 17 | HF | HF | 2 | 3 | 3 | 12 | 2 | 3 | Y | GEN, MB |
| | 14 | 4.54 | 1 | | 5 | HF | HF | 1 | 1 | 1 | 3 | 1 | 1 | Y | MB |
| | 16 | 12.1 | 4 | | 24 | HF | HF | 2 | 2 | 4 | 21 | 2 | 2 | Y | GEN, MB |
| | 18 | 5.47 | 1 | | 8 | HF | HF | 1 | 2 | 1 | 4 | 1 | 2 | Y | GEN, MB |
| | 19 | 27 | 2 | 1 | 18 | HF | LF | 2 | 4 | 2 | 10 | 2 | 4 | Y | GEN, MB |
| | 20 | 6.98 | 1 | | 8 | HF | LF | 1 | 1 | 1 | 6 | 1 | 1 | Y | MB |
| | 21 | 11.7 | 2 | | 22 | HF | HF | \ | \ | 2 | 10 | \ | \ | Y | MB |
| | 23 | NA | 1 | | 8 | HF | HF | 1 | 3 | 1 | 4 | 1 | 3 | OSL | |
| | 24 | 3.6 | 1 | | 3 | HF | HF | \ | \ | 1 | 3 | \ | \ | Y | MB |
| | 25 | 8.88 | 1 | | 11 | HF | LF | 1 | 5 | 1 | 9 | 1 | 5 | Y | GEN, MB |
| | 26 | 3.17 | 1 | | 7 | HF | HF | \ | \ | 1 | 7 | \ | \ | Y | MB |
| | 28 | 3.33 | 1 | | 5 | HF | HF | \ | \ | 1 | 2 | \ | \ | Y | MB |
| | 29 | 4.99 | 1 | | 6 | HF | HF | 1 | 2 | 1 | 6 | 1 | 2 | Y | GEN, MB |
| | 35 | 21.2 | 6 | 2 | 27 | LF | LF | 6 | 14 | 6 | 25 | 6 | 14 | Y | GEN, MB |
| | 37 | 27.5 | 4 | | 28 | LF | LF | 4 | 11 | 4 | 23 | 4 | 9 | Y | GEN, MB |
| | 38 | 10.6 | 2 | 1 | 20 | LF | LF | 2 | 11 | 2 | 10 | 2 | 6 | Y | GEN, MB |
| | 39 | 2.24 | 1 | 1 | 5 | LF | LF | \ | 1 | 1 | 4 | \ | 1 | Y | MB |
| | 41 | 3.29 | 1 | | 7 | HF | HF | 1 | 2 | 1 | 7 | 1 | 2 | Y | GEN, MB |
| | 43 | 4.12 | 2 | | 14 | HF | LF | 1 | 2 | 2 | 10 | 1 | 2 | Y | GEN, MB |
| | 45 | 5.26 | 2 | | 14 | LF | LF | 1 | 2 | 2 | 10 | 1 | 2 | Y | GEN, MB |
| | 46 | 29.2 | 1 | | 8 | HF | LF | 1 | 5 | 1 | 7 | 1 | 5 | Y | GEN, MB |
| | 48 | 36.2 | 6 | | 38 | LF | LF | 5 | 10 | 6 | 31 | 5 | 10 | Y | GEN, MB |
| | 49 | 0.19 | 1 | | 8 | HF | LF | 1 | 3 | 1 | 4 | 1 | 2 | N | |
| | 50 | 21 | 2 | | 11 | LF | LF | 2 | 2 | 2 | 10 | 2 | 2 | N | |
| | 38NB | NA | 1 | | 5 | HF | LF | 1 | 2 | 1 | 2 | 1 | 2 | OSL | |
| | MDA | NA | 1 | | 8 | HF | HF | \ | \ | 1 | 7 | \ | \ | OSL | |
| | Artisanias | 13.8 | 6 | 1 | 37 | HF | | 1 | 5 | | | | | N | |
| | Camino Real | 6.8 | 1 | | 5 | HF | | 1 | 2 | | | | | N | |
| | CBTA | 26 | 3 | | 21 | HF | | 3 | 5 | | | | | N | |
| | Chacamax | 11.7 | 1 | | 7 | HF | | 1 | 4 | | | | | N | |
| | Chan Kha | NA | 1 | | 5 | LF | | 1 | 2 | | | | | OSL | |
| | El Panchan | NA | 1 | 1 | 7 | LF | | 1 | 5 | | | | | OSL | |
| | La Mision | 15.4 | 1 | | 11 | HF | | 1 | 2 | | | | | N | |
| | Leon Brindis | 5.8 | 1 | | 10 | HF | | 1 | 2 | | | | | N | |
| | Nututun | 25.2 | 3 | | 21 | HF | | 1 | 1 | | | | | N | |
| | Quiloma | 5.3 | 1 | | 5 | HF | | 1 | 1 | | | | | N | |
| | Unitaria | NA | \ | 1 | 1 | HF | | \ | 1 | | | | | OSL | |
| PNP groups | BANO REINA | | | | 4 | PNP | | | 1 | | | | | | |
| | BOLAS | | | | 7 | PNP | | | 2 | | | | | | |
| | CALAVERA | | | | 9 | PNP | | | 3 | | | | | | |
| | EDGIES/NAHA | | | | 7 | PNP | | | 1 | | | | | | |
| | GROUP C2 | | | | 3 | PNP | | | 3 | | | | | | |
| | JAGUAR1 | | | | 4 | PNP | | | 1 | | | | | | |
| | MIGHTY DUC | | | | 4 | PNP | | | 1 | | | | | | |
| | MOTIEPA | 180 ha | | | 8 | PNP | | | 2 | | | | | | |
| | MUSEUM | | | | 3 | PNP | | | 1 | | | | | | |
| | PAKAL | | | | 9 | PNP | | | 1 | | | | | | |
| | PIGRITOS | | | | 6 | PNP | | | 1 | | | | | | |
| | T NORTE | | | | 10 | PNP | | | 4 | | | | | | |
| | TOLVIES | | | | 6 | PNP | | | 1 | | | | | | |
| | UNITES | | | | 9 | PNP | | | 5 | | | | | | |
| | WUPSIES | | | | 5 | PNP | | | 1 | | | | | | |
| | Total fragments | 469.3 | 81 | 8 | 536 | | | 57 | 130 | 62 | 307 | 45 | 92 | | |
| | Total PNP | 180 | 15 | \ | 94 | | | 15 | 28 | \ | \ | \ | \ | | |

**Table S2. By-sample summary of inclusion in final datasets.** Acronyms: MB – microbiome; Seq. batch – sequencing batch; LF – low-fragmentation; HF – high-fragmentation; SA – subadults; A – adult; J – juvenile; INF – infant; M – male; F – female; group names beginning with "L" indicate the sample is from a lone (solitary) individual.

| | | | | | | | Genetics | | | Microbiome | |
|---|---|---|---|---|---|---|---|---|---|---|---|
| **Sample ID** | **Fragment** | **Group name** | **Region frag-stats** | **Region MB** | **Age** | **Sex** | **Seq. batch** | **In final 158 IND dataset** | **% missing data in 158 IND dataset** | **Seq. batch** | **In final 307 IND dataset** |
| KK60 | 1 | 1A | LF | LF | SA | M | 1 | Y | 0.67 | 2 | Y |
| AC40 | 1 | 1A | LF | LF | A | M | \ | N | \ | 1 | Y |
| KK61 | 1 | 1A | LF | LF | A | F | \ | N | \ | 1 | Y |
| DL93 | 1 | 1A | LF | LF | A | F | 1, 2 | N | \ | 1 | Y |
| KK59 | 1 | 1A | LF | LF | J | M | \ | N | \ | 1 | N |
| KK74 | 1 | 1B | LF | LF | A | F | 1 | Y | 0.64 | 1 | Y |
| SM2 | 1 | 1B | LF | LF | A | F | \ | N | \ | 1 | Y |
| KK73 | 1 | 1B | LF | LF | A | M | \ | N | \ | 2 | Y |
| SM9 | 1 | 1C | LF | LF | A | M | 1 | Y | 0.30 | 1 | Y |
| KK76 | 1 | 1C | LF | LF | A | F | 2 | Y | 0.46 | 1 | Y |
| SM6 | 1 | 1C | LF | LF | A | F | 1 | N | \ | 1 | Y |
| KK77 | 1 | 1C | LF | LF | A | F | 2 | N | \ | 2 | Y |
| SM10 | 1 | 1C | LF | LF | J | M | \ | N | \ | 2 | Y |
| KK79 | 2 | 2A | LF | LF | A | F | 1 | N | \ | 1 | Y |
| KK80 | 2 | 2A | LF | LF | A | F | 1 | N | \ | 1 | Y |
| SM11 | 2 | 2A | LF | LF | SA | M | 2 | N | \ | 1 | Y |
| SM14 | 2 | 2A | LF | LF | A | M | 2 | N | \ | 1 | Y |
| SM12 | 2 | 2A | LF | LF | A | F | \ | N | \ | 1 | Y |
| SM15 | 2 | 2A | LF | LF | A | M | \ | N | \ | 1 | Y |
| SM13 | 2 | 2A | LF | LF | J | F | \ | N | \ | 2 | Y |
| KK78 | 2 | 2A | LF | LF | A | F | \ | N | \ | 1 | N |
| DL9 | 3 | 3A | LF | LF | A | M | 1, 2 | Y | 0.48 | 1 | Y |
| DL7 | 3 | 3A | LF | LF | SA | M | 2 | Y | 0.43 | 2 | Y |
| DL8 | 3 | 3A | LF | LF | A | F | \ | N | \ | 1 | Y |
| LE1 | 3 | 3A | LF | LF | J | M | \ | N | \ | 1 | Y |
| LE2 | 3 | 3A | LF | LF | J | M | \ | N | \ | 1 | Y |
| LE3 | 3 | 3A | LF | LF | A | F | \ | N | \ | 2 | Y |
| SM44 | 4 | 4A | HF | LF | A | M | 1 | Y | 0.34 | 1 | Y |
| SM43 | 4 | 4A | HF | LF | A | F | 1 | Y | 0.46 | 1 | Y |
| KK6 | 4 | 4A | HF | LF | A | M | \ | N | \ | 1 | N |
| KK72 | 6 | 6A | HF | HF | A | M | 1 | N | \ | 1 | Y |
| KK71 | 6 | 6A | HF | HF | A | F | 2 | N | \ | 1 | Y |
| DL106 | 6 | 6A | HF | HF | INF | F | \ | N | \ | 1 | Y |
| KK70 | 6 | 6A | HF | HF | J | F | \ | N | \ | 2 | Y |

| | | | | | | | | | | | |
|---|---|---|---|---|---|---|---|---|---|---|---|
| DL107 | 6 | 6A | HF | HF | UID | UID | \ | N | \ | 1 | N |
| DL6 | 9 | 9A | HF | HF | A | M | 1 | N | \ | 1 | Y |
| AC3 | 9 | 9A | HF | HF | A | F | 2 | N | \ | 1 | Y |
| KK4 | 9 | 9A | HF | HF | J | M | \ | N | \ | 1 | Y |
| KK5 | 9 | 9A | HF | HF | INF | M | \ | N | \ | 1 | Y |
| DL5 | 9 | 9A | HF | HF | A | F | 2 | N | \ | 2 | Y |
| AC2 | 9 | 9A | HF | HF | SA | M | \ | N | \ | 2 | Y |
| DL4 | 9 | 9A | HF | HF | J | M | \ | N | \ | 2 | Y |
| DL1 | 10 | 10A | HF | HF | A | M | \ | N | \ | 1 | Y |
| KK1 | 10 | 10A | HF | HF | A | F | \ | N | \ | 1 | Y |
| AC1 | 10 | 10A | HF | HF | A | F | \ | N | \ | 2 | Y |
| KK3 | 10 | 10B | HF | HF | A | F | 2 | Y | 0.56 | 1 | Y |
| DL2 | 10 | 10B | HF | HF | A | F | 2 | N | \ | 1 | Y |
| DL3 | 10 | 10B | HF | HF | A | F | \ | N | \ | 1 | Y |
| KK2 | 10 | 10B | HF | HF | A | F | 1, 2 | N | \ | 1 | Y |
| RC1 | 10 | 10B | HF | HF | J | M | \ | N | \ | 2 | Y |
| RC2 | 10 | 10B | HF | HF | A | F | \ | N | \ | 1 | N |
| DL38 | 12 | 12A | HF | HF | A | F | 1 | Y | 0.64 | 1 | Y |
| DL40 | 12 | 12A | HF | HF | A | M | 1 | Y | 0.59 | 1 | Y |
| DL39 | 12 | 12A | HF | HF | J | M | \ | N | \ | 1 | Y |
| DL42 | 12 | 12A | HF | HF | A | F | \ | N | \ | 1 | Y |
| LE17 | 12 | 12A | HF | HF | J | F | \ | N | \ | 1 | Y |
| LE18 | 12 | 12A | HF | HF | A | F | \ | N | \ | 1 | Y |
| DL41 | 12 | 12A | HF | HF | J | F | \ | N | \ | 2 | Y |
| DL47 | 12 | 12B | HF | HF | A | M | 1 | Y | 0.57 | 1 | Y |
| DL44 | 12 | 12B | HF | HF | A | M | \ | N | \ | 1 | Y |
| DL46 | 12 | 12B | HF | HF | A | F | \ | N | \ | 1 | Y |
| LE19 | 12 | 12B | HF | HF | J | F | \ | N | \ | 1 | Y |
| LE20 | 12 | 12B | HF | HF | A | F | \ | N | \ | 1 | Y |
| DL45 | 12 | 12B | HF | HF | J | M | \ | N | \ | 2 | Y |
| LE21 | 12 | 12B | HF | HF | SA | M | \ | N | \ | 2 | Y |
| KK32 | 13 | 13A | HF | HF | A | F | 1 | Y | 0.38 | 2 | Y |
| AC18 | 13 | 13A | HF | HF | SA | M | 1 | Y | 0.69 | 2 | Y |
| AC19 | 13 | 13A | HF | HF | J | F | \ | N | \ | 1 | Y |
| KK29 | 13 | 13A | HF | HF | J | M | \ | N | \ | 2 | Y |
| KK31 | 13 | 13A | HF | HF | J | F | \ | N | \ | 2 | Y |
| AC22 | 13 | 13B | HF | HF | A | F | 2 | Y | 0.38 | 1 | Y |
| DL49 | 13 | 13B | HF | HF | SA | M | 2 | N | \ | 2 | Y |
| AC20 | 13 | 13B | HF | HF | A | M | \ | N | \ | 2 | Y |
| AC21 | 13 | 13B | HF | HF | A | F | \ | N | \ | 2 | Y |
| DL48 | 13 | 13B | HF | HF | J | M | \ | N | \ | 2 | Y |
| DL50 | 13 | 13B | HF | HF | A | M | 1 | N | \ | \ | N |
| DL51 | 13 | 13C | HF | HF | J | F | \ | N | \ | 2 | Y |

| | | | | | | | | | | | |
|---|---|---|---|---|---|---|---|---|---|---|---|
| KK33 | 13 | 13LF | HF | HF | A | F | \ | N | \ | 2 | Y |
| KK65 | 14 | 14A | HF | HF | A | F | 1 | Y | 0.57 | 2 | Y |
| DL98 | 14 | 14A | HF | HF | A | F | 2 | N | \ | 1 | Y |
| AC45 | 14 | 14A | HF | HF | A | M | \ | N | \ | 1 | Y |
| KK66 | 14 | 14A | HF | HF | J | M | \ | N | \ | 2 | N |
| DL23 | 16 | 16A | HF | HF | A | F | 1 | Y | 0.27 | 1 | Y |
| KK19 | 16 | 16A | HF | HF | A | M | \ | N | \ | 1 | Y |
| DL22 | 16 | 16A | HF | HF | A | F | \ | N | \ | 1 | Y |
| KK20 | 16 | 16A | HF | HF | J | F | \ | N | \ | 1 | Y |
| KK21 | 16 | 16A | HF | HF | J | M | \ | N | \ | 1 | Y |
| AC10 | 16 | 16B | HF | HF | INF | F | \ | N | \ | 1 | Y |
| DL10 | 16 | 16B | HF | HF | A | F | \ | N | \ | 1 | Y |
| KK8 | 16 | 16B | HF | HF | A | F | 2 | N | \ | 2 | Y |
| KK7 | 16 | 16B | HF | HF | A | M | \ | N | \ | 2 | Y |
| AC4 | 16 | 16B | HF | HF | J | F | \ | N | \ | 2 | Y |
| KK22 | 16 | 16C | HF | HF | A | M | 1 | N | \ | 1 | Y |
| DL25 | 16 | 16C | HF | HF | A | F | \ | N | \ | 1 | Y |
| DL24 | 16 | 16C | HF | HF | INF | M | \ | N | \ | 1 | Y |
| DL26 | 16 | 16C | HF | HF | INF | F | \ | N | \ | 1 | Y |
| DL27 | 16 | 16C | HF | HF | J | M | \ | N | \ | 2 | Y |
| DL29 | 16 | 16C | HF | HF | A | F | \ | N | \ | 2 | Y |
| KK24 | 16 | 16D | HF | HF | A | F | 1 | Y | 0.31 | 1 | Y |
| LE11 | 16 | 16D | HF | HF | J | F | \ | N | \ | 1 | Y |
| AC11 | 16 | 16D | HF | HF | A | M | \ | N | \ | 1 | Y |
| LE9 | 16 | 16D | HF | HF | A | F | \ | N | \ | 2 | Y |
| DL28 | 16 | 16D | HF | HF | J | F | \ | N | \ | 2 | Y |
| LE10 | 16 | 16D | HF | HF | A | F | \ | N | \ | 1 | N |
| KK108 | 16 | 16E? | HF | HF | A | M | 1 | N | \ | \ | N |
| AC14 | 18 | 18A | HF | HF | A | F | 2 | Y | 0.50 | 1 | Y |
| KK25 | 18 | 18A | HF | HF | A | F | 1, 2 | Y | 0.48 | 1 | Y |
| AC12 | 18 | 18A | HF | HF | A | F | \ | N | \ | 1 | Y |
| KK26 | 18 | 18A | HF | HF | J | M | \ | N | \ | 2 | Y |
| AC15 | 18 | 18A | HF | HF | A | M | 1, 2 | N | \ | \ | N |
| CA1 | 19 | 19A | HF | LF | A | F | 1 | Y | 0.67 | 2 | Y |
| KK81 | 19 | 19A | HF | LF | A | F | \ | N | \ | 1 | Y |
| SM16 | 19 | 19A | HF | LF | A | F | \ | N | \ | 1 | Y |
| CA2 | 19 | 19A | HF | LF | A | M | \ | N | \ | 1 | N |
| CA3 | 19 | 19B | HF | LF | A | F | 2 | Y | 0.55 | 1 | Y |
| SM19 | 19 | 19B | HF | LF | SA | M | 1 | Y | 0.59 | 2 | Y |
| SM18 | 19 | 19B | HF | LF | J | M | 2 | Y | 0.60 | 2 | Y |
| KK82 | 19 | 19B | HF | LF | A | M | \ | N | \ | 1 | Y |
| CA4 | 19 | 19B | HF | LF | J | M | \ | N | \ | 1 | Y |
| SM17 | 19 | 19B | HF | LF | J | F | \ | N | \ | 2 | Y |

| | | | | | | | | | | | |
|---|---|---|---|---|---|---|---|---|---|---|---|
| SM41 | 19 | 19LM | HF | LF | A | M | 1 | N | \ | 1 | Y |
| DL35 | 20 | 20A | HF | LF | A | F | 1 | Y | 0.60 | 1 | Y |
| DL33 | 20 | 20A | HF | LF | J | F | \ | N | \ | 1 | Y |
| DL34 | 20 | 20A | HF | LF | INF | F | \ | N | \ | 1 | Y |
| LE12 | 20 | 20A | HF | LF | A | F | \ | N | \ | 1 | Y |
| LE13 | 20 | 20A | HF | LF | INF | F | \ | N | \ | 1 | Y |
| DL30 | 20 | 20A | HF | LF | A | M | 1, 2 | N | \ | 1 | Y |
| DL31 | 20 | 20A | HF | LF | J | F | \ | N | \ | 1 | N |
| CA23 | 21 | 21A | HF | HF | A | M | \ | N | \ | 1 | Y |
| KK89 | 21 | 21A | HF | HF | A | F | \ | N | \ | 1 | Y |
| KK90 | 21 | 21A | HF | HF | J | M | \ | N | \ | 1 | Y |
| CA28 | 21 | 21A | HF | HF | J | F | \ | N | \ | 2 | Y |
| KK103 | 21 | 21A | HF | HF | J | F | \ | N | \ | 2 | Y |
| TP5 | 21 | 21A | HF | HF | J | M | \ | N | \ | 2 | Y |
| KK102 | 21 | 21A | HF | HF | A | F | 1 | N | \ | 1 | N |
| CA24 | 21 | 21B | HF | HF | A | M | 2 | N | \ | 1 | Y |
| SM47 | 21 | 21B | HF | HF | A | F | \ | N | \ | 1 | Y |
| KK105 | 21 | 21B | HF | HF | A | F | \ | N | \ | 1 | Y |
| KK104 | 21 | 21B | HF | HF | J | F | \ | N | \ | 2 | Y |
| CA26 | 23 | 23A | HF | HF | A | F | 1 | Y | 0.34 | 1 | Y |
| CA25 | 23 | 23A | HF | HF | SA | M | 1 | Y | 0.67 | 2 | Y |
| KK107 | 23 | 23A | HF | HF | J | F | 2 | Y | 0.28 | 2 | Y |
| KK106 | 23 | 23A | HF | HF | A | M | \ | N | \ | 1 | Y |
| DL71 | 24 | 24A | HF | HF | A | M | 1 | N | \ | 1 | Y |
| KK47 | 24 | 24A | HF | HF | INF | F | \ | N | \ | 1 | Y |
| DL72 | 24 | 24A | HF | HF | A | F | 1 | N | \ | 2 | Y |
| DL36 | 25 | 25A | HF | LF | A | M | 1 | Y | 0.67 | 1 | Y |
| DL37 | 25 | 25A | HF | LF | A | M | 2 | Y | 0.24 | 1 | Y |
| LE15 | 25 | 25A | HF | LF | A | F | 2 | Y | 0.67 | 1 | Y |
| LE16 | 25 | 25A | HF | LF | A | M | 2 | Y | 0.66 | 1 | Y |
| KK27 | 25 | 25A | HF | LF | A | F | 1, 2 | Y | 0.63 | 1 | Y |
| AC17 | 25 | 25A | HF | LF | SA | M | 2 | N | \ | 2 | Y |
| AC16 | 25 | 25A | HF | LF | SA | F | \ | N | \ | 2 | Y |
| KK28 | 25 | 25A | HF | LF | J | F | \ | N | \ | 2 | Y |
| LE14 | 25 | 25A | HF | LF | J | M | \ | N | \ | 2 | Y |
| KK45 | 26 | 26A | HF | HF | A | F | 2 | N | \ | 1 | Y |
| DL69 | 26 | 26A | HF | HF | A | M | \ | N | \ | 1 | Y |
| DL70 | 26 | 26A | HF | HF | A | M | \ | N | \ | 1 | Y |
| KK46 | 26 | 26A | HF | HF | A | F | \ | N | \ | 1 | Y |
| KK44 | 26 | 26A | HF | HF | A | F | \ | N | \ | 2 | Y |
| DL67 | 26 | 26A | HF | HF | J | M | \ | N | \ | 2 | Y |
| DL68 | 26 | 26A | HF | HF | J | M | \ | N | \ | 2 | Y |
| SM46 | 28 | 28A | HF | HF | A | M | 2 | N | \ | 1 | Y |

| | | | | | | | | | | | |
|---|---|---|---|---|---|---|---|---|---|---|---|
| SM45 | 28 | 28A | HF | HF | A | F | \ | N | \ | 1 | Y |
| KK101 | 28 | 28A | HF | HF | A | F | 1 | N | \ | 2 | N |
| KK86 | 29 | 29A | HF | HF | A | F | 2 | Y | 0.52 | 1 | Y |
| TP3 | 29 | 29A | HF | HF | J | M | 2 | Y | 0.50 | 2 | Y |
| SM20 | 29 | 29A | HF | HF | A | F | \ | N | \ | 1 | Y |
| CA7 | 29 | 29A | HF | HF | A | M | 1 | N | \ | 2 | Y |
| SM22 | 29 | 29A | HF | HF | J | F | \ | N | \ | 2 | Y |
| KK87 | 29 | 29A | HF | HF | A | M | \ | N | \ | 2 | Y |
| KK94 | 35 | 35A | LF | LF | A | M | 2 | Y | 0.69 | 1 | Y |
| SM30 | 35 | 35A | LF | LF | A | F | \ | N | \ | 1 | Y |
| SM31 | 35 | 35A | LF | LF | A | F | 1 | N | \ | 2 | Y |
| SM32 | 35 | 35A | LF | LF | A | F | 2 | N | \ | 2 | Y |
| TP9 | 35 | 35A | LF | LF | J | F | 2 | N | \ | 2 | Y |
| KK96 | 35 | 35B | LF | LF | A | M | 1 | Y | 0.54 | 1 | Y |
| CA17 | 35 | 35B | LF | LF | A | F | 2 | Y | 0.73 | 1 | Y |
| CA14 | 35 | 35B | LF | LF | A | F | 2 | Y | 0.48 | 2 | Y |
| CA15 | 35 | 35B | LF | LF | SA | M | 2 | N | \ | 2 | Y |
| KK95 | 35 | 35B | LF | LF | A | F | \ | N | \ | 1 | N |
| CA16 | 35 | 35B | LF | LF | A | F | 2 | N | \ | \ | N |
| SM34 | 35 | 35C | LF | LF | A | F | 1 | Y | 0.59 | 2 | Y |
| SM35 | 35 | 35C | LF | LF | J | F | 2 | Y | 0.56 | 2 | Y |
| SM33 | 35 | 35C | LF | LF | A | M | \ | N | \ | 1 | Y |
| TP12 | 35 | 35C | LF | LF | A | F | \ | N | \ | 2 | Y |
| KK98 | 35 | 35D | LF | LF | A | M | 1, 2 | Y | 0.72 | 1 | Y |
| CA19 | 35 | 35D | LF | LF | A | F | 2 | N | \ | 1 | Y |
| SM36 | 35 | 35D | LF | LF | SA | F | 2 | N | \ | 2 | Y |
| SM37 | 35 | 35E | LF | LF | A | M | 1 | Y | 0.31 | 1 | Y |
| SM38 | 35 | 35E | LF | LF | A | F | 2 | Y | 0.61 | 1 | Y |
| AL6 | 35 | 35E | LF | LF | A | F | 2 | N | \ | 1 | Y |
| CA20 | 35 | 35F | LF | LF | A | M | 1 | Y | 0.55 | 1 | Y |
| SM39 | 35 | 35F | LF | LF | A | F | 2 | Y | 0.74 | 1 | Y |
| CA21 | 35 | 35F | LF | LF | SA | M | 2 | Y | 0.67 | 2 | Y |
| TP13 | 35 | 35F | LF | LF | A | F | 2 | N | \ | 1 | Y |
| KK97 | 35 | 35LF | LF | LF | SA | F | 1 | Y | 0.57 | 2 | Y |
| CA18 | 35 | 35LM | LF | LF | SA | M | 1 | Y | 0.39 | 2 | Y |
| DL12 | 37 | 37A | LF | LF | A | M | 1 | Y | 0.50 | 1 | Y |
| LE5 | 37 | 37A | LF | LF | A | F | 2 | Y | 0.24 | 1 | Y |
| LE4 | 37 | 37A | LF | LF | SA | F | 2 | Y | 0.68 | 2 | Y |
| DL11 | 37 | 37A | LF | LF | INF | F | \ | N | \ | 1 | Y |
| KK9 | 37 | 37A | LF | LF | A | M | \ | N | \ | 1 | Y |
| KK13 | 37 | 37B | LF | LF | A | F | 1 | Y | 0.32 | 1 | Y |
| LE6 | 37 | 37B | LF | LF | A | F | 2 | Y | 0.22 | 1 | Y |
| KK10 | 37 | 37B | LF | LF | A | M | 1, 2 | Y | 0.46 | 2 | Y |

| | | | | | | | | | | | |
|---|---|---|---|---|---|---|---|---|---|---|---|
| AC5 | 37 | 37B | LF | LF | J | M | \ | N | \ | 1 | Y |
| LE7 | 37 | 37B | LF | LF | INF | F | \ | N | \ | 1 | Y |
| KK12 | 37 | 37B | LF | LF | J | M | \ | N | \ | 2 | Y |
| KK11 | 37 | 37B | LF | LF | A | F | \ | N | \ | 1 | N |
| LE8 | 37 | 37C | LF | LF | A | F | 1 | Y | 0.33 | 1 | Y |
| KK18 | 37 | 37C | LF | LF | A | F | 2 | Y | 0.29 | 2 | Y |
| KK16 | 37 | 37C | LF | LF | SA | M | 2 | Y | 0.25 | \ | N |
| KK15 | 37 | 37C | LF | LF | A | M | 1 | N | \ | 1 | Y |
| DL18 | 37 | 37C | LF | LF | J | M | \ | N | \ | 1 | Y |
| DL19 | 37 | 37C | LF | LF | J | F | \ | N | \ | 1 | Y |
| KK17 | 37 | 37C | LF | LF | SA | M | \ | N | \ | 2 | Y |
| DL21 | 37 | 37C | LF | LF | J | F | \ | N | \ | 2 | Y |
| AC9 | 37 | 37C | LF | LF | A | UID | \ | N | \ | 1 | N |
| DL14 | 37 | 37D | LF | LF | A | F | 2 | Y | 0.43 | 1 | Y |
| AC7 | 37 | 37D | LF | LF | A | F | 1 | Y | 0.62 | \ | N |
| AC6 | 37 | 37D | LF | LF | A | F | 2 | N | \ | 1 | Y |
| DL13 | 37 | 37D | LF | LF | A | M | \ | N | \ | 1 | Y |
| DL16 | 37 | 37D | LF | LF | INF | M | \ | N | \ | 1 | Y |
| AC8 | 37 | 37D | LF | LF | J | F | \ | N | \ | 2 | Y |
| SM24 | 38 | 38A | LF | LF | A | M | 1 | Y | 0.44 | 1 | Y |
| SM29 | 38 | 38A | LF | LF | A | F | 1, 2 | Y | 0.31 | 1 | Y |
| CA9 | 38 | 38A | LF | LF | SA | M | 2 | Y | 0.30 | \ | N |
| KK93 | 38 | 38A | LF | LF | A | F | 2 | Y | 0.34 | \ | N |
| SM30.1 | 38 | 38A | LF | LF | ? | F | 2 | Y | 0.25 | \ | N |
| CA12 | 38 | 38A | LF | LF | A | M | 2 | Y | 0.58 | \ | N |
| AL5 | 38 | 38A | LF | LF | A | F | \ | N | \ | 1 | Y |
| TP6 | 38 | 38A | LF | LF | J | F | \ | N | \ | 1 | Y |
| SM28 | 38 | 38A | LF | LF | J | F | \ | N | \ | 2 | Y |
| SM26 | 38 | 38C | LF | LF | A | M | 1 | Y | 0.56 | 1 | Y |
| CA11 | 38 | 38C | LF | LF | A | F | 2 | Y | 0.49 | 1 | Y |
| SM25 | 38 | 38C | LF | LF | SA | M | 2 | Y | 0.41 | 2 | Y |
| SM27 | 38 | 38C | LF | LF | SA | F | 1, 2 | Y | 0.26 | 2 | Y |
| CA10 | 38 | 38C | LF | LF | A | M | 2 | Y | 0.47 | \ | N |
| KK92 | 38 | 38LM | LF | LF | A | M | \ | N | \ | 1 | Y |
| KK83 | 39 | 39A | LF | LF | A | M | \ | N | \ | 1 | Y |
| KK85 | 39 | 39A | LF | LF | A | F | \ | N | \ | 1 | Y |
| KK84 | 39 | 39A | LF | LF | J | F | \ | N | \ | 2 | Y |
| TP1 | 39 | 39A | LF | LF | A | M | \ | N | \ | 1 | N |
| SM42 | 39 | 39LSAF | LF | LF | SA | F | 1 | Y | 0.29 | 2 | Y |
| DL75 | 41 | 41A | HF | HF | A | M | 1 | Y | 0.30 | 1 | Y |
| AC30 | 41 | 41A | HF | HF | A | F | 1 | Y | 0.58 | 2 | Y |
| AC31 | 41 | 41A | HF | HF | J | M | \ | N | \ | 1 | Y |
| DL74 | 41 | 41A | HF | HF | A | F | \ | N | \ | 1 | Y |

| | | | | | | | | | | | |
|---|---|---|---|---|---|---|---|---|---|---|---|
| AC29 | 41 | 41A | HF | HF | J | M | \ | N | \ | 2 | Y |
| DL73 | 41 | 41A | HF | HF | A | M | \ | N | \ | 2 | Y |
| KK48 | 41 | 41A | HF | HF | J | F | \ | N | \ | 2 | Y |
| KK62 | 43 | 43A | HF | LF | A | M | 2 | N | \ | 1 | Y |
| DL94 | 43 | 43A | HF | LF | A | F | \ | N | \ | 1 | Y |
| DL95 | 43 | 43A | HF | LF | A | F | 1 | N | \ | 2 | Y |
| AC41 | 43 | 43A | HF | LF | J | F | \ | N | \ | 2 | Y |
| AC44 | 43 | 43B | HF | LF | A | M | 1 | Y | 0.66 | 1 | Y |
| KK64 | 43 | 43B | HF | LF | SA | M | 2 | Y | 0.72 | 2 | Y |
| AC42 | 43 | 43B | HF | LF | A | F | \ | N | \ | 1 | Y |
| AC43 | 43 | 43B | HF | LF | J | M | \ | N | \ | 1 | Y |
| AL3 | 43 | 43B | HF | LF | J | M | \ | N | \ | 1 | Y |
| DL97 | 43 | 43B | HF | LF | A | F | \ | N | \ | 1 | Y |
| DL96 | 43 | 43B | HF | LF | J | M | \ | N | \ | 2 | Y |
| KK39 | 45 | 45A | LF | LF | A | M | 1 | N | \ | 1 | Y |
| AC27 | 45 | 45A | LF | LF | A | F | 1, 2 | N | \ | 1 | Y |
| KK40 | 45 | 45A | LF | LF | J | M | \ | N | \ | 2 | N |
| DL57 | 45 | 45B | LF | LF | A | F | 2 | Y | 0.68 | 1 | Y |
| DL60 | 45 | 45B | LF | LF | A | F | 2 | Y | 0.27 | 2 | Y |
| DL58 | 45 | 45B | LF | LF | J | F | \ | N | \ | 1 | Y |
| DL59 | 45 | 45B | LF | LF | J | M | \ | N | \ | 1 | Y |
| LE22 | 45 | 45B | LF | LF | A | F | \ | N | \ | 1 | Y |
| LE24 | 45 | 45B | LF | LF | A | M | \ | N | \ | 1 | Y |
| LE23 | 45 | 45B | LF | LF | A | F | 1 | N | \ | 2 | Y |
| AC28 | 45 | 45B | LF | LF | J | F | \ | N | \ | 2 | Y |
| DL61 | 45 | 45B | LF | LF | J | M | \ | N | \ | 2 | N |
| KK35 | 46 | 46A | HF | LF | A | F | 1 | Y | 0.43 | 1 | Y |
| DL55 | 46 | 46A | HF | LF | A | F | 1 | Y | 0.28 | 1 | Y |
| KK37 | 46 | 46A | HF | LF | A | F | 2 | Y | 0.46 | 1 | Y |
| KK36 | 46 | 46A | HF | LF | A | M | 1 | Y | 0.73 | 2 | Y |
| DL53 | 46 | 46A | HF | LF | SA | M | 2 | Y | 0.38 | 2 | Y |
| AC24 | 46 | 46A | HF | LF | INF | F | \ | N | \ | 1 | Y |
| DL54 | 46 | 46A | HF | LF | J | M | \ | N | \ | 1 | Y |
| DL76 | 48 | 48A | LF | LF | A | M | \ | N | \ | 1 | Y |
| DL77 | 48 | 48A | LF | LF | A | F | \ | N | \ | 1 | Y |
| KK49 | 48 | 48A | LF | LF | A | F | \ | N | \ | 1 | Y |
| KK51 | 48 | 48B | LF | LF | A | M | 1 | Y | 0.51 | 1 | Y |
| DL78 | 48 | 48B | LF | LF | A | F | 1, 2 | Y | 0.67 | 1 | Y |
| AC34 | 48 | 48B | LF | LF | A | F | \ | N | \ | 1 | Y |
| AC33 | 48 | 48B | LF | LF | J | M | \ | N | \ | 1 | Y |
| KK50 | 48 | 48B | LF | LF | A | M | \ | N | \ | 1 | Y |
| AC32 | 48 | 48B | LF | LF | J | F | \ | N | \ | 2 | Y |
| DL79 | 48 | 48C | LF | LF | A | M | 2 | Y | 0.32 | 1 | Y |

| | | | | | | | | | | | |
|---|---|---|---|---|---|---|---|---|---|---|---|
| AL1 | 48 | 48C | LF | LF | SA | M | 1 | Y | 0.62 | 2 | Y |
| AC35 | 48 | 48C | LF | LF | A | F | \ | N | \ | 1 | Y |
| DL80 | 48 | 48C | LF | LF | J | F | \ | N | \ | 1 | Y |
| DL82 | 48 | 48C | LF | LF | SA | F | \ | N | \ | 2 | Y |
| DL81 | 48 | 48C | LF | LF | J | F | \ | N | \ | 1 | N |
| DL88 | 48 | 48D | LF | LF | A | F | 1 | Y | 0.33 | 1 | Y |
| DL87 | 48 | 48D | LF | LF | A | F | 2 | Y | 0.69 | 2 | Y |
| DL85 | 48 | 48D | LF | LF | J | M | 2 | Y | 0.74 | 2 | Y |
| DL86 | 48 | 48D | LF | LF | INF | M | \ | N | \ | 1 | Y |
| KK52 | 48 | 48D | LF | LF | A | M | \ | N | \ | 2 | Y |
| DL89 | 48 | 48E | LF | LF | A | F | 2 | Y | 0.68 | 1 | Y |
| AC37 | 48 | 48E | LF | LF | A | M | 1 | N | \ | 1 | Y |
| AC36 | 48 | 48E | LF | LF | A | F | \ | N | \ | 1 | Y |
| DL90 | 48 | 48E | LF | LF | A | F | \ | N | \ | 1 | Y |
| AL2 | 48 | 48E | LF | LF | J | F | \ | N | \ | 2 | Y |
| DL91 | 48 | 48F | LF | LF | A | F | 2 | Y | 0.56 | 1 | Y |
| AC38 | 48 | 48F | LF | LF | SA | M | 2 | Y | 0.40 | 2 | Y |
| KK56 | 48 | 48F | LF | LF | A | M | 1 | N | \ | 1 | Y |
| AC39 | 48 | 48F | LF | LF | A | F | \ | N | \ | 1 | Y |
| KK58 | 48 | 48F | LF | LF | J | M | \ | N | \ | 1 | Y |
| DL92 | 48 | 48F | LF | LF | A | F | 1 | N | \ | 2 | Y |
| KK57 | 48 | 48F | LF | LF | J | F | \ | N | \ | 2 | Y |
| KK38 | 49 | 49A | HF | LF | A | F | 1 | Y | 0.53 | 1 | Y |
| SM40 | 49 | 49A | HF | LF | A | M | 1 | Y | 0.41 | 1 | Y |
| DL56 | 49 | 49A | HF | LF | A | F | 1 | Y | 0.32 | \ | N |
| AC26 | 49 | 49A | HF | LF | A | F | \ | N | \ | 1 | Y |
| AC25 | 49 | 49A | HF | LF | J | M | \ | N | \ | 2 | Y |
| KK34 | 49 | 49A | HF | LF | UID | UID | 2 | N | \ | \ | N |
| DL105 | 50 | LMA | LF | LF | A | F | 1 | Y | 0.52 | 1 | Y |
| AL4 | 50 | LMA | LF | LF | A | M | \ | N | \ | 1 | Y |
| DL104 | 50 | LMA | LF | LF | A | F | 1, 2 | N | \ | 1 | Y |
| KK68 | 50 | LMRA | LF | LF | A | F | 1 | Y | 0.28 | 2 | Y |
| DL100 | 50 | LMRA | LF | LF | A | M | 2 | N | \ | 1 | Y |
| KK67 | 50 | LMRA | LF | LF | A | F | \ | N | \ | 1 | Y |
| DL101 | 50 | LMRA | LF | LF | J | F | \ | N | \ | 2 | Y |
| AC46 | 50 | LMRA | LF | LF | J | F | \ | N | \ | 2 | Y |
| DL102 | 50 | LMRA | LF | LF | J | F | \ | N | \ | 2 | Y |
| DL99 | 50 | LMRA | LF | LF | J | M | \ | N | \ | 2 | Y |
| KK100 | 38NB | 38NB | HF | LF | J | M | 2 | Y | 0.28 | 2 | Y |
| CA22 | 38NB | 38NB | HF | LF | A | M | 1 | Y | 0.30 | 1 | N |
| K22 | Artisanias | A4 | HF | HF | A | F | 1 | Y | 0.55 | \ | N |
| K34 | Artisanias | LSAF | HF | HF | SA | F | 2 | Y | 0.46 | \ | N |
| A9 | Artisanias | A1 | HF | HF | SA | M | 2 | N | \ | \ | N |

| | | | | | | | | | | | |
|---|---|---|---|---|---|---|---|---|---|---|---|
| A11 | Artisanias | A3 | HF | HF | A | M | 2 | N | \ | \ | N |
| K18 | Artisanias | A3 | HF | HF | A | F | 2 | N | \ | \ | N |
| A16 | Artisanias | A4 | HF | HF | UID | F | 2 | Y | 0.72 | \ | N |
| K23 | Artisanias | A4 | HF | HF | A | F | 2 | Y | 0.30 | \ | N |
| A13 | Artisanias | A4 | HF | HF | SA | F | 2 | Y | 0.50 | \ | N |
| A39 | By House | BH | HF | HF | A | F | 2 | N | \ | \ | N |
| K7 | Camino Real | CR | HF | HF | A | F | 2 | Y | 0.26 | \ | N |
| K6 | Camino Real | CR | HF | HF | A | M | 2 | Y | 0.43 | \ | N |
| K8 | Camino Real | CR | HF | HF | A | M | 2 | N | \ | \ | N |
| K26 | CBTA | CBTA1 | HF | HF | A | F | 2 | Y | 0.29 | \ | N |
| A19 | CBTA | CBTA1 | HF | HF | A | M | 2 | Y | 0.51 | \ | N |
| K24 | CBTA | CBTA1 | HF | HF | A | M | 2 | Y | 0.59 | \ | N |
| A23 | CBTA | CBTA2 | HF | HF | A | M | 2 | Y | 0.30 | \ | N |
| A25 | CBTA | CBTA2 | HF | HF | A | F | 2 | N | \ | \ | N |
| A27 | CBTA | CBTA3 | HF | HF | A | M | 2 | Y | 0.34 | \ | N |
| A28 | CBTA | CBTA3 | HF | HF | A | F | 2 | N | \ | \ | N |
| A37 | Chacamax | CM | HF | HF | A | F | 2 | Y | 0.34 | \ | N |
| A38 | Chacamax | CM | HF | HF | A | M | 2 | Y | 0.35 | \ | N |
| A35 | Chacamax | CM | HF | HF | J | M | 2 | Y | 0.26 | \ | N |
| A36 | Chacamax | CM | HF | HF | J | M | 2 | Y | 0.35 | \ | N |
| K47 | Chacamax | CM | HF | HF | A | F | 2 | N | \ | \ | N |
| A5 | Chan Kha | CK | LF | LF | A | F | 2 | Y | 0.61 | \ | N |
| A8 | Chan Kha | CK | LF | LF | A | F | 2 | Y | 0.37 | \ | N |
| A6 | Chan Kha | CK | LF | LF | A | M | 2 | N | \ | \ | N |
| A30 | Chiapaneca | CP | HF | HF | A | F | 2 | N | \ | \ | N |
| A40 | El Panchan | LJM | LF | LF | J | M | 2 | Y | 0.64 | \ | N |
| K59 | El Panchan | Park Gate | LF | LF | A | F | 2 | Y | 0.62 | \ | N |
| K60 | El Panchan | Park Gate | LF | LF | A | M | 2 | Y | 0.26 | \ | N |
| K56 | El Panchan | Park Gate | LF | LF | J | M | 2 | Y | 0.51 | \ | N |
| K58 | El Panchan | Park Gate | LF | LF | J | M | 2 | Y | 0.68 | \ | N |
| K31 | La Mision | LM | HF | HF | A | M | 2 | Y | 0.46 | \ | N |
| A32 | La Mision | LM | HF | HF | J | F | 2 | Y | 0.48 | \ | N |
| K4 | La Quiloma | LQ | HF | HF | A | M | 2 | Y | 0.61 | \ | N |
| K43 | Leon Brindis | LB | HF | HF | A | F | 2 | Y | 0.32 | \ | N |
| K45 | Leon Brindis | LB | HF | HF | A | M | 2 | Y | 0.37 | \ | N |
| A33 | Leon Brindis | LB | HF | HF | SA | M | 2 | N | \ | \ | N |
| DL62 | MD | MDA | HF | HF | A | F | 1 | N | \ | 1 | Y |
| KK42 | MD | MDA | HF | HF | A | F | \ | N | \ | 1 | Y |
| DL63 | MD | MDA | HF | HF | A | F | \ | N | \ | 1 | Y |
| KK43 | MD | MDA | HF | HF | A | F | 1, 2 | N | \ | 1 | Y |
| KK41 | MD | MDA | HF | HF | SA | M | 1 | N | \ | 2 | Y |
| DL65 | MD | MDA | HF | HF | A | M | \ | N | \ | 2 | Y |
| DL64 | MD | MDA | HF | HF | J | M | \ | N | \ | 2 | Y |

| | | | | | | | | | | | |
|---|---|---|---|---|---|---|---|---|---|---|---|
| K49 | Nututun | NT1 | HF | HF | A | F | 2 | Y | 0.38 | \ | N |
| K50 | Nututun | NT2 | HF | HF | A | F | 2 | N | \ | \ | N |
| A34 | Unitaria | LAF | HF | HF | A | F | 1 | Y | 0.49 | \ | N |
| SVB-58 | PNP | Balam | PNP | PNP | J | M | 3 | N | \ | \ | N |
| SVB-59 | PNP | Balam | PNP | PNP | A | F | 3 | N | \ | \ | N |
| SVB-60 | PNP | Balam | PNP | PNP | A | M | 3 | N | \ | \ | N |
| SVB-78 | PNP | Balam | PNP | PNP | A | F | 3 | N | \ | \ | N |
| SVB-79 | PNP | Balam | PNP | PNP | A | M | 3 | N | \ | \ | N |
| SVB-151 | PNP | BanoReina | PNP | PNP | J | F | 3 | Y | 0.70 | \ | N |
| SVB-104 | PNP | BanoReina | PNP | PNP | A | M | 3 | N | \ | \ | N |
| SVB-105 | PNP | BanoReina | PNP | PNP | A | F | 3 | N | \ | \ | N |
| SVB-152 | PNP | BanoReina | PNP | PNP | J | F | 3 | N | \ | \ | N |
| SVB-198 | PNP | Bolas | PNP | PNP | A | F | 3 | Y | 0.42 | \ | N |
| SVB-196 | PNP | Bolas | PNP | PNP | A | M | 3 | Y | 0.51 | \ | N |
| SVB-181 | PNP | Bolas | PNP | PNP | J | M | 3 | N | \ | \ | N |
| SVB-194 | PNP | Bolas | PNP | PNP | A | F | 3 | N | \ | \ | N |
| SVB-195 | PNP | Bolas | PNP | PNP | A | F | 3 | N | \ | \ | N |
| SVB-197 | PNP | Bolas | PNP | PNP | SA | M | 3 | N | \ | \ | N |
| SVB-121 | PNP | Calavera | PNP | PNP | A | F | 3 | Y | 0.31 | \ | N |
| SVB-117 | PNP | Calavera | PNP | PNP | SA | M | 3 | Y | 0.57 | \ | N |
| SVB-119 | PNP | Calavera | PNP | PNP | J | M | 3 | Y | 0.63 | \ | N |
| SVB-114 | PNP | Calavera | PNP | PNP | A | M | 3 | N | \ | \ | N |
| SVB-115 | PNP | Calavera | PNP | PNP | A | M | 3 | N | \ | \ | N |
| SVB-118 | PNP | Calavera | PNP | PNP | SA | M | 3 | N | \ | \ | N |
| SVB-120 | PNP | Calavera | PNP | PNP | INF | M | 3 | N | \ | \ | N |
| SVB-122 | PNP | Calavera | PNP | PNP | A | F | 3 | N | \ | \ | N |
| SVB-166 | PNP | Edgies/Naha | PNP | PNP | J | F | 3 | Y | 0.57 | \ | N |
| SVB-161 | PNP | Edgies/Naha | PNP | PNP | A | M | 3 | N | \ | \ | N |
| SVB-162 | PNP | Edgies/Naha | PNP | PNP | A | M | 3 | N | \ | \ | N |
| SVB-164 | PNP | Edgies/Naha | PNP | PNP | A | F | 3 | N | \ | \ | N |
| SVB-165 | PNP | Edgies/Naha | PNP | PNP | A | F | 3 | N | \ | \ | N |
| SVB-189 | PNP | Group C2 | PNP | PNP | A | F | 3 | Y | 0.28 | \ | N |
| SVB-190 | PNP | Group C2 | PNP | PNP | A | F | 3 | Y | 0.45 | \ | N |
| SVB-188 | PNP | Group C2 | PNP | PNP | A | M | 3 | Y | 0.46 | \ | N |
| SVB-103 | PNP | Jaguar1 | PNP | PNP | J | M | 3 | Y | 0.62 | \ | N |
| SVB-100 | PNP | Jaguar1 | PNP | PNP | A | F | 3 | N | \ | \ | N |
| SVB-101 | PNP | Jaguar1 | PNP | PNP | A | F | 3 | N | \ | \ | N |
| SVB-97 | PNP | Jaguar1 | PNP | PNP | A | M | 3 | N | \ | \ | N |
| SVB-98 | PNP | Jaguar1 | PNP | PNP | A | M | 3 | N | \ | \ | N |
| SVB-99 | PNP | Jaguar1 | PNP | PNP | SA | M | 3 | N | \ | \ | N |
| SVB-128 | PNP | Jaguar2 | PNP | PNP | A | M | 3 | N | \ | \ | N |
| SVB-129 | PNP | Jaguar2 | PNP | PNP | A | F | 3 | N | \ | \ | N |
| SVB-130 | PNP | Jaguar2 | PNP | PNP | A | F | 3 | N | \ | \ | N |

| | | | | | | | | | | | |
|---|---|---|---|---|---|---|---|---|---|---|---|
| SVB-131 | PNP | Jaguar2 | PNP | PNP | J | M | 3 | N | \ | \ | N |
| SVB-170 | PNP | Mayabell | PNP | PNP | A | F | 3 | N | \ | \ | N |
| SVB-171 | PNP | Mayabell | PNP | PNP | A | F | 3 | N | \ | \ | N |
| SVB-173 | PNP | Mayabell | PNP | PNP | A | M | 3 | N | \ | \ | N |
| SVB-174 | PNP | Mayabell | PNP | PNP | A | M | 3 | N | \ | \ | N |
| SVB-176 | PNP | Mayabell | PNP | PNP | SA | M | 3 | N | \ | \ | N |
| SVB-178 | PNP | Mayabell | PNP | PNP | SA | F | 3 | N | \ | \ | N |
| SVB-187 | PNP | Mighty Duc | PNP | PNP | SA | F | 3 | Y | 0.53 | \ | N |
| SVB-179 | PNP | Mighty Duc | PNP | PNP | SA | M | 3 | N | \ | \ | N |
| SVB-185 | PNP | Mighty Duc | PNP | PNP | A | M | 3 | N | \ | \ | N |
| SVB-186 | PNP | Mighty Duc | PNP | PNP | A | F | 3 | N | \ | \ | N |
| SVB-62 | PNP | Motiepa | PNP | PNP | A | F | 3 | Y | 0.42 | \ | N |
| SVB-73 | PNP | Motiepa | PNP | PNP | J | F | 3 | Y | 0.74 | \ | N |
| SVB-180 | PNP | Motiepa | PNP | PNP | INF | M | 3 | N | \ | \ | N |
| SVB-48 | PNP | Motiepa | PNP | PNP | SA | M | 3 | N | \ | \ | N |
| SVB-61 | PNP | Motiepa | PNP | PNP | A | M | 3 | N | \ | \ | N |
| SVB-63 | PNP | Motiepa | PNP | PNP | J | M | 3 | N | \ | \ | N |
| SVB-75 | PNP | Motiepa | PNP | PNP | A | F | 3 | N | \ | \ | N |
| SVB-76 | PNP | Motiepa | PNP | PNP | A | M | 3 | N | \ | \ | N |
| SVB-80 | PNP | Motiepa | PNP | PNP | J | F | 3 | N | \ | \ | N |
| SVB-134 | PNP | Museum | PNP | PNP | J | F | 3 | Y | 0.72 | \ | N |
| SVB-132 | PNP | Museum | PNP | PNP | A | M | 3 | N | \ | \ | N |
| SVB-133 | PNP | Museum | PNP | PNP | A | F | 3 | N | \ | \ | N |
| SVB-44 | PNP | Pakal | PNP | PNP | A | M | 3 | Y | 0.58 | \ | N |
| SVB-137 | PNP | Pakal | PNP | PNP | INF | UID | 3 | N | \ | \ | N |
| SVB-143 | PNP | Pakal | PNP | PNP | INF | UID | 3 | N | \ | \ | N |
| SVB-42 | PNP | Pakal | PNP | PNP | A | M | 3 | N | \ | \ | N |
| SVB-65 | PNP | Pakal | PNP | PNP | A | F | 3 | N | \ | \ | N |
| SVB-66 | PNP | Pakal | PNP | PNP | A | F | 3 | N | \ | \ | N |
| SVB-69 | PNP | Pakal | PNP | PNP | A | F | 3 | N | \ | \ | N |
| SVB-70 | PNP | Pakal | PNP | PNP | SA | F | 3 | N | \ | \ | N |
| SVB-71 | PNP | Pakal | PNP | PNP | A | F | 3 | N | \ | \ | N |
| SVB-112 | PNP | Pigritos | PNP | PNP | A | F | 3 | Y | 0.72 | \ | N |
| SVB-110 | PNP | Pigritos | PNP | PNP | A | M | 3 | N | \ | \ | N |
| SVB-111 | PNP | Pigritos | PNP | PNP | A | M | 3 | N | \ | \ | N |
| SVB-113 | PNP | Pigritos | PNP | PNP | A | F | 3 | N | \ | \ | N |
| SVB-156 | PNP | T Norte | PNP | PNP | A | F | 3 | Y | 0.50 | \ | N |
| SVB-157 | PNP | T Norte | PNP | PNP | A | F | 3 | Y | 0.48 | \ | N |
| SVB-154 | PNP | T Norte | PNP | PNP | A | M | 3 | Y | 0.63 | \ | N |
| SVB-158 | PNP | T Norte | PNP | PNP | J | F | 3 | Y | 0.32 | \ | N |
| SVB-153 | PNP | T Norte | PNP | PNP | A | M | 3 | N | \ | \ | N |
| SVB-155 | PNP | T Norte | PNP | PNP | A | M | 3 | N | \ | \ | N |
| SVB-160 | PNP | T Norte | PNP | PNP | J | F | 3 | N | \ | \ | N |

| | | | | | | | | | | | |
|---|---|---|---|---|---|---|---|---|---|---|---|
| SVB-126 | PNP | Tolvies | PNP | PNP | J | M | 3 | Y | 0.68 | \ | N |
| SVB-123 | PNP | Tolvies | PNP | PNP | A | M | 3 | N | \ | \ | N |
| SVB-124 | PNP | Tolvies | PNP | PNP | A | F | 3 | N | \ | \ | N |
| SVB-125 | PNP | Tolvies | PNP | PNP | A | F | 3 | N | \ | \ | N |
| SVB-199 | PNP | Unites | PNP | PNP | A | F | 3 | Y | 0.52 | \ | N |
| SVB-205 | PNP | Unites | PNP | PNP | A | F | 3 | Y | 0.32 | \ | N |
| SVB-200 | PNP | Unites | PNP | PNP | SA | F | 3 | Y | 0.43 | \ | N |
| SVB-202 | PNP | Unites | PNP | PNP | A | M | 3 | Y | 0.29 | \ | N |
| SVB-203 | PNP | Unites | PNP | PNP | A | M | 3 | Y | 0.71 | \ | N |
| SVB-148 | PNP | Unites | PNP | PNP | A | F | 3 | N | \ | \ | N |
| SVB-149 | PNP | Unites | PNP | PNP | J | M | 3 | N | \ | \ | N |
| SVB-204 | PNP | Unites | PNP | PNP | J | M | 3 | N | \ | \ | N |
| SVB-107 | PNP | Wupsies | PNP | PNP | A | M | 3 | Y | 0.51 | \ | N |
| SVB-108 | PNP | Wupsies | PNP | PNP | A | F | 3 | N | \ | \ | N |
| SVB-109 | PNP | Wupsies | PNP | PNP | J | M | 3 | N | \ | \ | N |

**Table S3. Description of all habitat quality, connectivity, and demographic variables** analyzed throughout this study, primarily in the LASSO + multiple regression analyses (adapted from Klass et al., 2020a, Klass 2024).

| | Variable | Calculation method | Description and meaning | Data source/s |
|---|---|---|---|---|
| **Habitat quality** | **area (ha)** | Area tool in Google Earth© | fragment size | high-resolution (1 $m^2$) Google Earth© (2018) satellite images combined with on-the-ground validation during fragment surveys |
| | **shape index** | Shape index = fragment perimeter / (√(fragment area) × π) | The higher the index, the more irregular the fragment shape; a shape index = 1 indicates a perfect circle. Higher shape indices may indicate more edge habitat. | high-resolution (1 $m^2$) Google Earth© (2018) satellite images combined with on-the-ground validation during fragment surveys |
| | **core area** | Calculated with the “PatchStat” function in the SDMTools package in R (VanDerWal et al., 2014) | The area within each fragment unaffected by edges, as identified by FRAGSTATS (McGarigal et al., 2012). Environmental variables (e.g., temperature, humidity) may change with increasing edge habitat, which can cause changes in the physical structure and tree species composition of the forest. | University of Maryland’s open access Global Forest Change Dataset (Hansen et al., 2013) and high-resolution (1 $m^2$) Google Earth© (2018) satellite images combined with on-the-ground validation during fragment surveys |
| | **core area index** | Calculated with the “PatchStat” function in the SDMTools package in R (VanDerWal et al., 2014) | The percentage of the core area from the total patch area as identified by FRAGSTATS (McGarigal et al., 2012). Environmental variables (e.g., temperature, humidity) may change with increasing edge habitat, which can cause changes in the physical structure and tree species composition of the forest. | University of Maryland’s open access Global Forest Change Dataset (Hansen et al., 2013) and high-resolution (1 $m^2$) Google Earth© (2018) satellite images combined with on-the-ground validation during fragment surveys |
| | **stem density** | Stem density = (number of trees sampled of DBH ≥ 10 cm)/ (area sampled in transects (1,000 $m^2$)) | lower stem density is expected in mature, primary forest | Transect data |
| | **mean tree DBH (cm)** | Mean tree diameter for all trees with DBH ≥ 10 cm sampled in fragment transects | indicative of size/maturity of fragment tree community | Transect data |
| | **maximum tree DBH (cm)** | Largest diameter of tree with DBH ≥ 10 cm sampled in fragment transects | indicative of size/maturity of fragment tree community | Transect data |
| | **mean tree height (m)** | Mean tree height for all trees with DBH ≥ 10 cm sampled in fragment transects | indicative of size/maturity of fragment tree community | Transect data |

| | | | | |
|---|---|---|---|---|
| | **maximum tree height (m)** | Largest tree height measured in fragment transects of those with DBH ≥ 10 cm | indicative of size/maturity of fragment tree community | Transect data |
| | **genus richness** | the number of tree genera identified in fragment transects | indicative of forest quality; fragments with very low richness may provide howlers with uniform, nutritionally poor diets | Transect data |
| | **Shannon diversity index (H')** | $H' = \sum_{i=1}^{s} - (\pi * \ln(\pi))$<br><br>π = fraction of population made up of type i; S = total numbers of types | indicative of forest quality; fragments with low tree diversity may provide howlers with uniform, nutritionally poor diets | Transect data |
| | **Simpson's Evenness Index** | $E=(1/\Sigma\pi^2)/S$<br>π = the proportion of individuals belonging to genus i; S = total number of genera sampled in fragment (Morris et al. 2014) | indicative of forest quality; fragments with low evenness are dominated by one or a few species, which may lead to a clumped resource distribution | Transect data |
| | **proportion known food trees** | # trees of genera found to be important food sources for black howlers in PNP identified in fragment transects/total # of identified trees in fragment transects | indicative of habitat quality for black howlers | Transect data, SVB unpublished data |
| | **proportion of *Ficus* trees** | # trees from the genus *Ficus* identified in fragment transects/total # of identified trees in transects | indicative of habitat quality for howlers; *Ficus* are a particularly important food source for howler monkeys in general, and for black howlers in PNP specifically | Transect data |
| | **mean Bray-Curtis dissimilarity index - trees** | The fragment's average Bray-Curtis index with all other fragments.<br><br>BCij = 1 – (2*Cij) / (Si + Sj)<br><br>Cij: The sum of the lesser values for the species found in each site<br>Si: The total number of specimens counted at site i<br>Sj: The total number of specimens counted at site j | Measures how similar two sites are in terms of species composition and abundance. The lower the value, the greater the similarity. The per-fragment mean Bray-Curtis index is indicative of how distinct and different that fragment's forest is compared to other study fragments. | Transect data |
| | **number of tree stumps** | # of tree stumps with DBH ≥ 10 cm in fragment transects | indicative of anthropogenic or natural forest disturbance such as tree mortality or logging | Transect data |

| | | | | |
|---|---|---|---|---|
| **Habitat connectivity** | **number of treelines** | the number of treelines (living fences, narrow secondary growth) extending from the fragment edge into the matrix, including those connecting the fragment to other forest fragments | indicative of connectivity - black howlers in the study landscape have been observed to use treelines for travel. Trees in treelines may also be used as an additional food source | high-resolution (1 $m^2$) Google Earth© (2018) satellite images combined with on-the-ground validation during fragment surveys |
| | **proportion forest cover in buffer around fragment** | proportion forested area in buffer/total buffer area<br><br>buffer around fragment = a circle with a radius of median dispersal distance, which can be estimated based on the species' home range size. Using the mean home range size (9.3 ha) of five groups studied long term in PNP (de Guinea et al., 2019), we calculated an estimated median dispersal distance, and buffer radius, of 2,400m with the formula (Bowman et al., 2002):<br>dispersal distance = 7 × (diameter of home range size)<br><br>forest was defined as 30$m^2$ pixels with ≥70% canopy cover | the proportion of forest habitat present within a pre-defined buffer zone around each fragment, excluding the area of the fragment itself; an area-based measure of connectivity, which may be more appropriate for arboreal species that are able, but reluctant, to travel on the ground | University of Maryland's open access Global Forest Change Dataset (Hansen et al., 2013), high-resolution (1 $m^2$) Google Earth© (2018) satellite images combined with on-the-ground validation during fragment surveys |
| | **distance to the nearest fragment (m)** | straight-line distance to nearest fragment ≥1 ha in size, regardless of fragment occupancy. Measured with the distance tool in Google Earth©. | indicative of connectivity; the distance between the study fragment and the nearest patch of forest | high-resolution (1 $m^2$) Google Earth© (2018) satellite images |
| | **distance to Palenque National Park (m)** | straight-line distance to edge of PNP. Measured with the distance tool in Google Earth©. | indicative of connectivity; the distance between the fragment and the nearest large block of continuous, primary forest | high-resolution (1 $m^2$) Google Earth© (2018) satellite images |
| **Demography** | **total fragment population size** | total number of unique individuals observed in fragment surveys | | fragment surveys (Klass et al. 2020A) |

| | | | | |
|---|---|---|---|---|
| | **population density** | total number of individuals/fragment area | | fragment surveys (Klass et al. 2020A), high-resolution (1 $m^2$) Google Earth© (2018) satellite images combined with on-the-ground validation during fragment surveys |
| | **number groups in fragment** | total number of unique social groups observed in fragment surveys | the number of social units with distinct territories in the fragment sharing limited space and resources | fragment surveys (Klass et al. 2020A) |
| | **proportion of AM** | total number of adult males observed in fragment / total fragment population | may be indicative of male dispersal patterns or other sex-specific responses to changing habitat quality and connectivity (e.g., increased mortality) | fragment surveys (Klass et al. 2020A) |
| | **proportion of AF** | total number of adult females observed in fragment / total fragment population | may be indicative of female dispersal patterns or other sex-specific responses to changing habitat quality and connectivity (e.g., increased mortality) | fragment surveys (Klass et al. 2020A) |
| | **ratio of INF:AF** | the total number of infants observed in the fragment / total number of adult females observed in the fragment | indicative of reproduction and population growth rates, which may in turn be indicative of stress or resource competition | fragment surveys (Klass et al. 2020A) |

### Genetic pipeline and analyses

#### Genetic diversity and inbreeding

**Table S4. Genetic data:** A) population-level summaries and B) individual values.

**(A)** Population-level genetic diversity and inbreeding measures for Palenque National Park (PNP) and fragmented landscape (Fragments – whole fragmented landscape; LF – low-fragmented area; HF – high-fragmented area), generated in Stacks.

| **2 REGIONS: PNP, ALL FRAGMENTS COMBINED** | | | | | | | |
|---|---|---|---|---|---|---|---|
| Variant positions | | | | | | | |
| Pop ID | | Obs_Het | Obs_Hom | Exp_Het | Exp_Hom | Pi | Fis |
| FRAGMENTS | | 0.20868 | 0.79132 | 0.24771 | 0.75229 | 0.24965 | 0.40494* |
| PNP | | 0.18886 | 0.81114 | 0.22775 | 0.77225 | 0.23844 | 0.28122 |
| All positions (variant and fixed) | | | | | | | |
| Pop ID | % Polymorphic Loci | Obs_Het | Obs_Hom | Exp_Het | Exp_Hom | Pi | Fis |
| FRAGMENTS | 0.18552 | 0.00039 | 0.99961 | 0.00046 | 0.99954 | 0.00047 | 0.00076 |
| PNP | 0.12252 | 0.00035 | 0.99965 | 0.00043 | 0.99957 | 0.00045 | 0.00053 |
| **3 REGIONS: PNP, LF, HF** | | | | | | | |
| Variant positions | | | | | | | |
| Pop ID | | Obs_Het | Obs_Hom | Exp_Het | Exp_Hom | Pi | Fis |
| HF | | 0.21071 | 0.78929 | 0.24767 | 0.75233 | 0.25143 | 0.34996* |
| LF | | 0.20655 | 0.79345 | 0.24215 | 0.75785 | 0.24621 | 0.35031* |
| PNP | | 0.18896 | 0.81104 | 0.2279 | 0.7721 | 0.2386 | 0.28133 |
| All positions (variant and fixed) | | | | | | | |
| Pop ID | % Polymorphic Loci | Obs_Het | Obs_Hom | Exp_Het | Exp_Hom | Pi | Fis |
| HF | 0.25488 | 0.00057 | 0.99943 | 0.00067 | 0.99933 | 0.00068 | 0.00095 |
| LF | 0.24785 | 0.00056 | 0.99944 | 0.00066 | 0.99934 | 0.00067 | 0.00095 |
| PNP | 0.17831 | 0.00051 | 0.99949 | 0.00062 | 0.99938 | 0.00065 | 0.00077 |

* The average population inbreeding values ($F_{is}$) found here for the fragmented landscape are higher than the values for PNP, which contradicts the pattern found in the by-individual inbreeding values reported in the main text. We attribute these high inbreeding values calculated for the fragmented landscape population as a whole to the Wahlund effect (Garnier-Géré and Chikhi, 2013), as a result of aggregating individuals from a genetically structured population into a single group (i.e., combining individuals from different fragments that form distinct genetic clusters into one large group), which can lead to the observation of an excess of homozygotes.

**(B)** Individual observed heterozygosity (calculated in *adegenet* in R) and inbreeding coefficients (calculated in COANCESTRY with Lynch-Ritland method).

| Individual | Region | Ho | LR inbreeding | Individual | Region | Ho | LR inbreeding |
|---|---|---|---|---|---|---|---|
| A13 | fragments | 0.219 | 0.368 | KK107 | fragments | 0.162 | 0.397 |
| A16 | fragments | 0.203 | 0.512 | KK13 | fragments | 0.181 | 0.465 |
| A19 | fragments | 0.213 | 0.453 | KK16 | fragments | 0.183 | 0.290 |
| A23 | fragments | 0.226 | 0.284 | KK18 | fragments | 0.202 | 0.368 |
| A27 | fragments | 0.209 | 0.391 | KK24 | fragments | 0.201 | 0.372 |
| A32 | fragments | 0.211 | 0.360 | KK25 | fragments | 0.216 | 0.291 |
| A34 | fragments | 0.201 | 0.492 | KK27 | fragments | 0.222 | 0.393 |
| A35 | fragments | 0.191 | 0.332 | KK3 | fragments | 0.232 | 0.411 |
| A36 | fragments | 0.215 | 0.403 | KK32 | fragments | 0.184 | 0.500 |
| A37 | fragments | 0.224 | 0.416 | KK35 | fragments | 0.176 | 0.528 |
| A38 | fragments | 0.205 | 0.442 | KK36 | fragments | 0.215 | 0.471 |
| A40 | fragments | 0.244 | 0.462 | KK37 | fragments | 0.222 | 0.325 |
| A5 | fragments | 0.222 | 0.413 | KK38 | fragments | 0.209 | 0.480 |
| A8 | fragments | 0.232 | 0.299 | KK51 | fragments | 0.190 | 0.492 |
| AC14 | fragments | 0.234 | 0.308 | KK60 | fragments | 0.218 | 0.458 |
| AC18 | fragments | 0.231 | 0.374 | KK64 | fragments | 0.223 | 0.414 |
| AC22 | fragments | 0.207 | 0.415 | KK65 | fragments | 0.207 | 0.445 |
| AC30 | fragments | 0.224 | 0.480 | KK68 | fragments | 0.185 | 0.384 |
| AC38 | fragments | 0.190 | 0.512 | KK74 | fragments | 0.224 | 0.496 |
| AC44 | fragments | 0.215 | 0.526 | KK76 | fragments | 0.226 | 0.386 |
| AC7 | fragments | 0.217 | 0.435 | KK86 | fragments | 0.201 | 0.580 |
| AL1 | fragments | 0.220 | 0.403 | KK93 | fragments | 0.211 | 0.419 |
| CA1 | fragments | 0.224 | 0.392 | KK94 | fragments | 0.224 | 0.317 |
| CA10 | fragments | 0.207 | 0.528 | KK96 | fragments | 0.218 | 0.444 |
| CA11 | fragments | 0.226 | 0.424 | KK97 | fragments | 0.230 | 0.434 |
| CA12 | fragments | 0.182 | 0.591 | KK98 | fragments | 0.237 | 0.441 |
| CA14 | fragments | 0.232 | 0.260 | LE15 | fragments | 0.244 | 0.362 |
| CA17 | fragments | 0.198 | 0.575 | LE16 | fragments | 0.247 | 0.448 |
| CA18 | fragments | 0.193 | 0.374 | LE4 | fragments | 0.210 | 0.431 |
| CA20 | fragments | 0.196 | 0.460 | LE5 | fragments | 0.171 | 0.356 |
| CA21 | fragments | 0.238 | 0.403 | LE6 | fragments | 0.221 | 0.228 |
| CA22 | fragments | 0.135 | 0.422 | LE8 | fragments | 0.168 | 0.549 |
| CA25 | fragments | 0.227 | 0.539 | SM18 | fragments | 0.208 | 0.459 |
| CA26 | fragments | 0.160 | 0.562 | SM19 | fragments | 0.220 | 0.423 |
| CA3 | fragments | 0.234 | 0.245 | SM24 | fragments | 0.220 | 0.338 |
| CA9 | fragments | 0.138 | 0.342 | SM25 | fragments | 0.221 | 0.314 |
| DL105 | fragments | 0.216 | 0.399 | SM26 | fragments | 0.247 | 0.317 |
| DL12 | fragments | 0.194 | 0.519 | SM27 | fragments | 0.196 | 0.394 |
| DL14 | fragments | 0.223 | 0.270 | SM29 | fragments | 0.107 | 0.535 |

| | | | | | | | |
|---|---|---|---|---|---|---|---|
| DL23 | fragments | 0.170 | 0.469 | SM30.1 | fragments | 0.210 | 0.353 |
| DL35 | fragments | 0.204 | 0.537 | SM34 | fragments | 0.228 | 0.350 |
| DL36 | fragments | 0.236 | 0.449 | SM35 | fragments | 0.195 | 0.405 |
| DL37 | fragments | 0.208 | 0.218 | SM37 | fragments | 0.155 | 0.511 |
| DL38 | fragments | 0.228 | 0.373 | SM38 | fragments | 0.225 | 0.387 |
| DL40 | fragments | 0.212 | 0.406 | SM39 | fragments | 0.246 | 0.502 |
| DL47 | fragments | 0.230 | 0.358 | SM40 | fragments | 0.175 | 0.506 |
| DL53 | fragments | 0.229 | 0.295 | SM42 | fragments | 0.142 | 0.515 |
| DL55 | fragments | 0.156 | 0.412 | SM43 | fragments | 0.186 | 0.517 |
| DL56 | fragments | 0.207 | 0.317 | SM44 | fragments | 0.151 | 0.529 |
| DL57 | fragments | 0.199 | 0.485 | SM9 | fragments | 0.151 | 0.520 |
| DL60 | fragments | 0.180 | 0.415 | TP3 | fragments | 0.225 | 0.404 |
| DL7 | fragments | 0.227 | 0.381 | 44 | PNP | 0.177 | 0.569 |
| DL75 | fragments | 0.180 | 0.497 | 62 | PNP | 0.155 | 0.681 |
| DL78 | fragments | 0.222 | 0.412 | 73 | PNP | 0.211 | 0.550 |
| DL79 | fragments | 0.117 | 0.565 | 103 | PNP | 0.168 | 0.597 |
| DL85 | fragments | 0.225 | 0.494 | 107 | PNP | 0.181 | 0.588 |
| DL87 | fragments | 0.235 | 0.405 | 112 | PNP | 0.178 | 0.609 |
| DL88 | fragments | 0.112 | 0.659 | 117 | PNP | 0.157 | 0.635 |
| DL89 | fragments | 0.232 | 0.481 | 119 | PNP | 0.210 | 0.591 |
| DL9 | fragments | 0.232 | 0.338 | 121 | PNP | 0.129 | 0.577 |
| DL91 | fragments | 0.229 | 0.414 | 126 | PNP | 0.223 | 0.575 |
| K22 | fragments | 0.180 | 0.595 | 134 | PNP | 0.141 | 0.715 |
| K23 | fragments | 0.228 | 0.279 | 151 | PNP | 0.204 | 0.609 |
| K24 | fragments | 0.245 | 0.422 | 154 | PNP | 0.184 | 0.633 |
| K26 | fragments | 0.238 | 0.252 | 156 | PNP | 0.162 | 0.626 |
| K31 | fragments | 0.153 | 0.668 | 157 | PNP | 0.150 | 0.631 |
| K34 | fragments | 0.235 | 0.327 | 158 | PNP | 0.119 | 0.675 |
| K4 | fragments | 0.218 | 0.424 | 166 | PNP | 0.165 | 0.650 |
| K43 | fragments | 0.207 | 0.283 | 187 | PNP | 0.177 | 0.622 |
| K45 | fragments | 0.196 | 0.479 | 188 | PNP | 0.176 | 0.619 |
| K49 | fragments | 0.232 | 0.303 | 189 | PNP | 0.141 | 0.575 |
| K56 | fragments | 0.203 | 0.480 | 190 | PNP | 0.188 | 0.585 |
| K58 | fragments | 0.213 | 0.426 | 196 | PNP | 0.169 | 0.577 |
| K59 | fragments | 0.211 | 0.534 | 198 | PNP | 0.139 | 0.577 |
| K6 | fragments | 0.211 | 0.388 | 199 | PNP | 0.198 | 0.562 |
| K60 | fragments | 0.215 | 0.274 | 200 | PNP | 0.176 | 0.628 |
| K7 | fragments | 0.247 | 0.229 | 202 | PNP | 0.140 | 0.554 |
| KK10 | fragments | 0.214 | 0.494 | 203 | PNP | 0.231 | 0.559 |
| KK100 | fragments | 0.193 | 0.469 | 205 | PNP | 0.166 | 0.602 |

**Fig. S1. Correlation of individual $H_o$ and inbreeding, in PNP and in fragments,** demonstrating differing in PNP vs. the fragmented landscape.

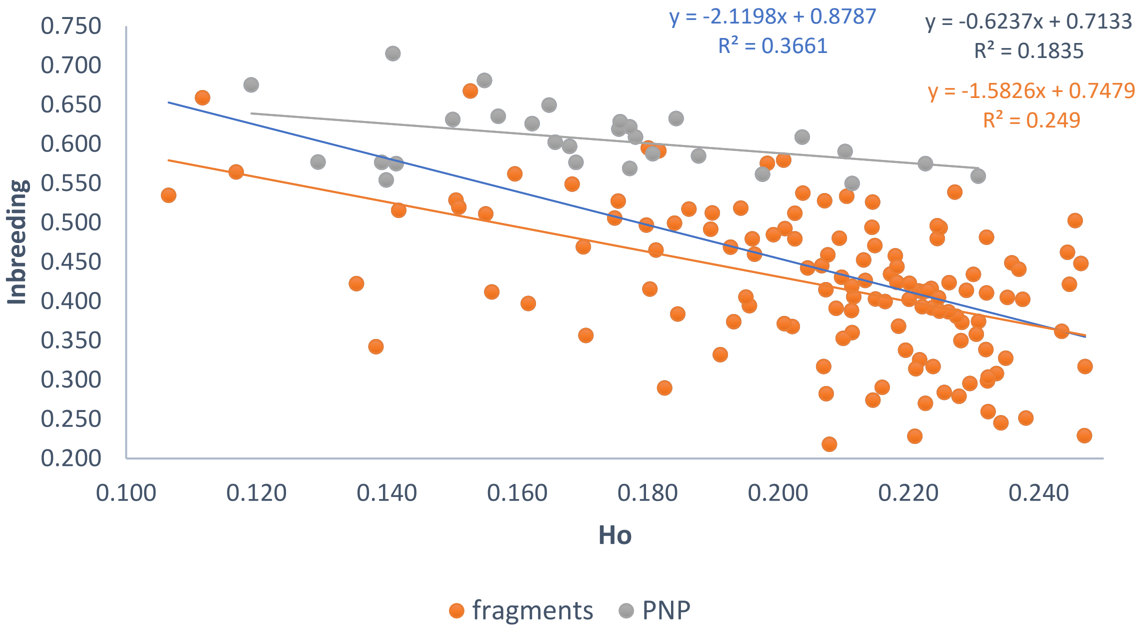

PNP individuals have consistently high inbreeding coefficients across a narrower range of heterozygosity. This suggests population-wide elevated relatedness and reduced genetic diversity, consistent with a small or long-isolated population where most individuals share similar ancestry. Because nearly everyone is similarly inbred, variation in Ho does not strongly translate into variation in inbreeding, so the correlation is weak. Fragment individuals span a wider range of Ho and inbreeding, producing the expected negative relationship. This indicates heterogeneous relatedness among individuals—some are relatively outbred while others are more inbred. This pattern is typical of recently fragmented populations or populations with mixed ancestry / variable mating among relatives.

In short, PNP appears more genetically uniform and chronically inbred, while the fragmented region retains more variation in relatedness among individuals. Given that allele frequencies were estimated from the entire dataset (PNP + fragments), the pattern likely reflects real differences in genetic structure between the populations, not an estimator artifact (blue trendline and equation on chart – whole dataset combined).

#### Population structure analyses

**Fig. S2. STRUCTURE results for subsets of data.** A) – D) similar sampling areas and/or sample sizes in PNP and the fragmented landscapes; E) more stringent missing data thresholds. See Table S5 for further details

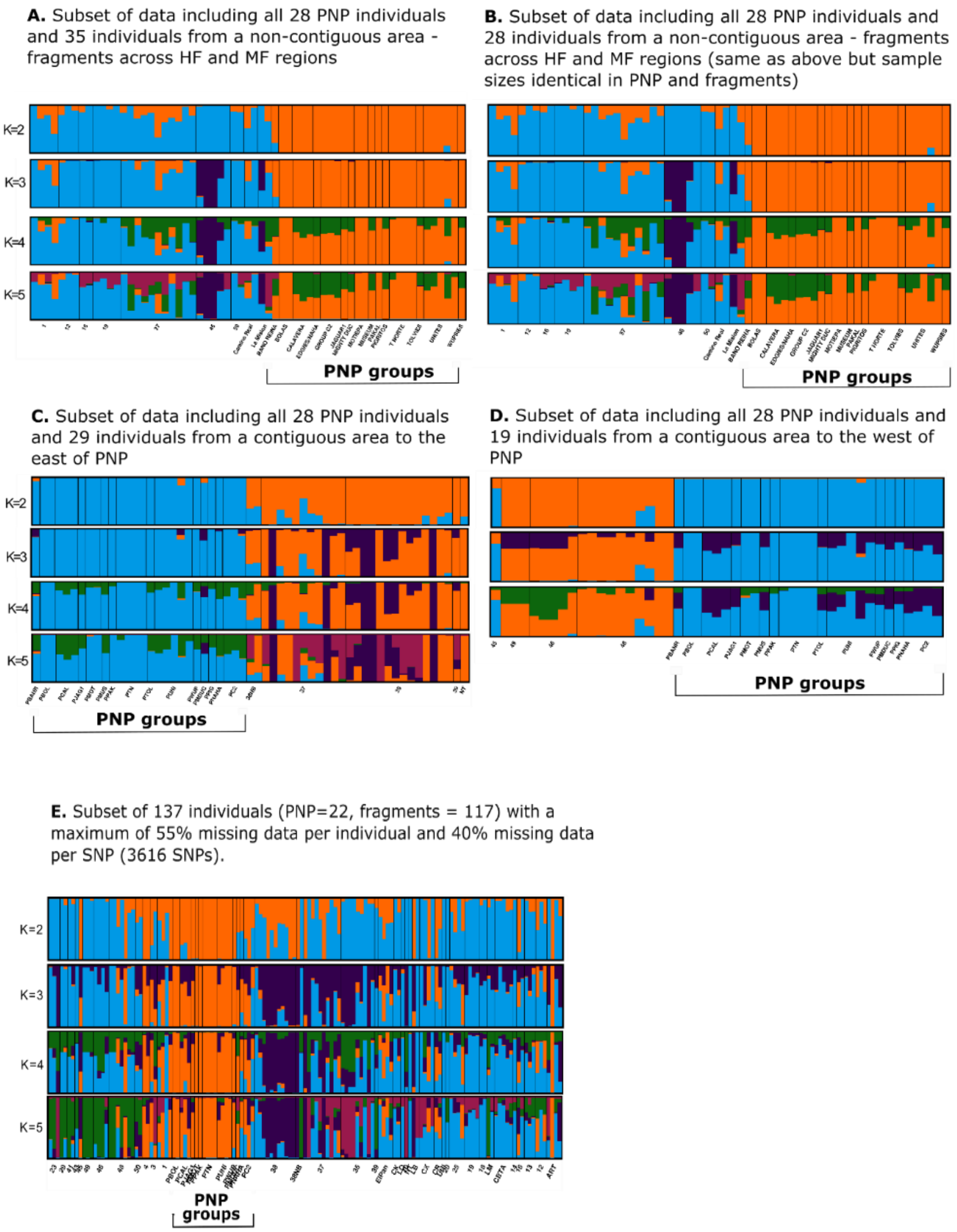

**Table S5. Additional details of subsetted datasets analyzed in Fig. S2.**

A) Description of the datasets used in subsetted STRUCTURE analyses.

| **Subset of data** | **fragmented landscape** | | | | **PNP** | | | **# SNPs** |
|---|---|---|---|---|---|---|---|---|
| | **#samples** | **#sampling locations** | **#sampled groups** | **sampling area (ha)** | **#samples** | **#sampled groups** | **sampling area (ha)** | |
| Including all 28 PNP individuals and 35 individuals from a non-contiguous area - fragments across HF and LF regions | 35 | 9 | 18 | 181 | 28 | 15 | 180 | 8810 |
| Including all 28 PNP individuals and 28 individuals from a non-contiguous area - fragments across HF and LF regions (sample sizes identical in PNP and fragments) | 28 | 9 | 18 | 181 | 28 | 15 | 180 | 8617 |
| Including all 28 PNP individuals and 29 individuals from a contiguous area to the east of PNP | 29 | 5 | 12 (+3 solitary ind.) | 77 | 28 | 15 | 180 | 8928 |
| Including all 28 PNP individuals and 19 individuals from a contiguous area to the west of PNP | 19 | 4 | 8 | 71 | 28 | 15 | 180 | 7808 |
| Including 137 individuals with a maximum of 55% missing data per individual and 40% missing data per SNP | 117 | 37 | 56 | 469 | 22 | 15 | 180 | 3616 |

B) Population-level (PNP vs. fragmented landscape) diversity and inbreeding statistics

| **Subset of data** | **% Poly-morphic loci** | **Obs. hetero-zygosity** | **Exp. hetero-zygosity** | **Obs. homo-zygosity** | **Exp. homo-zygosity** | **Nucleotide diversity (Pi)** | **Inbreeding (Fis)** |
|---|---|---|---|---|---|---|---|
| 28 PNP individuals and 35 individuals from a non-contiguous area - fragments across HF and LF regions | | | | | | | |
| **FRAGMENTS** | 0.215 | 0.270 | 0.305 | 0.730 | 0.695 | 0.314 | 0.269 |
| **PNP** | 0.162 | 0.205 | 0.258 | 0.795 | 0.742 | 0.269 | 0.319 |
| 28 PNP individuals and 28 individuals from a non-contiguous area - fragments across HF and LF regions (sample sizes identical in PNP and fragments) | | | | | | | |
| **FRAGMENTS** | 0.204 | 0.294 | 0.706 | 0.317 | 0.683 | 0.329 | 0.229 |
| **PNP** | 0.162 | 0.218 | 0.782 | 0.269 | 0.731 | 0.280 | 0.314 |
| 28 PNP individuals and 29 individuals from a contiguous area to the east of PNP | | | | | | | |
| **FRAGMENTS** | 0.201 | 0.297 | 0.703 | 0.316 | 0.684 | 0.328 | 0.224 |
| **PNP** | 0.164 | 0.235 | 0.765 | 0.277 | 0.723 | 0.289 | 0.295 |
| 28 PNP individuals and 19 individuals from a contiguous area to the west of PNP | | | | | | | |
| **FRAGMENTS** | 0.176 | 0.312 | 0.688 | 0.332 | 0.668 | 0.355 | 0.215 |
| **PNP** | 0.170 | 0.258 | 0.742 | 0.310 | 0.690 | 0.322 | 0.329 |
| 137 individuals with a maximum of 55% missing data per individual and 40% missing data per SNP. | | | | | | | |
| **FRAGMENTS** | 0.179 | 0.153 | 0.847 | 0.218 | 0.782 | 0.220 | 0.523 |
| **PNP** | 0.109 | 0.134 | 0.866 | 0.203 | 0.797 | 0.214 | 0.337 |

**Table S6. Consensus results from CLUMPAK synthesis of STRUCTURE runs for K=5, by individual.**

**A)** Each individual's proportional assignment (q value) to each of five clusters; when an individual had a q value > 0.6, the individual's row is colored in the color of its major cluster assignment (where q>0.6). If there is no major cluster assignment, the row remains white.

| IND | % missing | Fragment | region | dark purple | light purple | orange | blue | Green |
|---|---|---|---|---|---|---|---|---|
| 1 | 64 | 12 | HF | 0.311 | 0.018 | 0.519 | 0.118 | 0.034 |
| 2 | 58 | 12 | HF | 0.353 | 0.009 | 0.009 | 0.548 | 0.081 |
| 3 | 57 | 12 | HF | 0.246 | 0.004 | 0.359 | 0.339 | 0.052 |
| 4 | 68 | 13 | HF | 0.418 | 0.095 | 0.014 | 0.296 | 0.177 |
| 5 | 37 | 13 | HF | 0.001 | 0.171 | 0.002 | 0.799 | 0.027 |
| 6 | 37 | 13 | HF | 0.283 | 0.008 | 0.155 | 0.477 | 0.077 |
| 7 | 56 | 14 | HF | 0.342 | 0.041 | 0.085 | 0.455 | 0.077 |
| 8 | 27 | 16 | HF | 0.325 | 0.051 | 0.349 | 0.190 | 0.084 |
| 9 | 31 | 16 | HF | 0.256 | 0.072 | 0.004 | 0.594 | 0.074 |
| 10 | 50 | 18 | HF | 0.000 | 0.015 | 0.001 | 0.897 | 0.088 |
| 11 | 47 | 18 | HF | 0.001 | 0.044 | 0.001 | 0.884 | 0.071 |
| 12 | 66 | 19 | HF | 0.174 | 0.044 | 0.003 | 0.636 | 0.144 |
| 13 | 55 | 19 | HF | 0.101 | 0.018 | 0.002 | 0.775 | 0.105 |
| 14 | 59 | 19 | HF | 0.001 | 0.027 | 0.001 | 0.887 | 0.085 |
| 15 | 58 | 19 | HF | 0.021 | 0.046 | 0.004 | 0.799 | 0.130 |
| 16 | 59 | 20 | HF | 0.365 | 0.185 | 0.007 | 0.373 | 0.070 |
| 17 | 66 | 25 | HF | 0.001 | 0.005 | 0.155 | 0.685 | 0.155 |
| 18 | 24 | 25 | HF | 0.001 | 0.002 | 0.213 | 0.665 | 0.120 |
| 19 | 63 | 25 | HF | 0.002 | 0.040 | 0.001 | 0.804 | 0.153 |
| 20 | 67 | 25 | HF | 0.005 | 0.016 | 0.397 | 0.544 | 0.038 |
| 21 | 66 | 25 | HF | 0.001 | 0.037 | 0.549 | 0.413 | 0.001 |
| 22 | 49 | Artisanias | HF | 0.001 | 0.664 | 0.001 | 0.332 | 0.002 |
| 23 | 72 | Artisanias | HF | 0.194 | 0.001 | 0.002 | 0.763 | 0.040 |
| 24 | 54 | Artisanias | HF | 0.000 | 0.000 | 0.999 | 0.000 | 0.000 |
| 25 | 29 | Artisanias | HF | 0.116 | 0.007 | 0.001 | 0.852 | 0.024 |
| 26 | 45 | Artisanias | HF | 0.001 | 0.154 | 0.016 | 0.396 | 0.433 |
| 27 | 49 | Unitaria | HF | 0.019 | 0.066 | 0.001 | 0.819 | 0.096 |
| 28 | 42 | Camino Real | HF | 0.001 | 0.001 | 0.370 | 0.545 | 0.083 |
| 29 | 26 | Camino Real | HF | 0.056 | 0.175 | 0.259 | 0.489 | 0.022 |
| 30 | 51 | CBTA | HF | 0.021 | 0.161 | 0.002 | 0.783 | 0.033 |
| 31 | 30 | CBTA | HF | 0.141 | 0.007 | 0.003 | 0.775 | 0.074 |
| 32 | 34 | CBTA | HF | 0.000 | 0.004 | 0.001 | 0.937 | 0.058 |
| 33 | 59 | CBTA | HF | 0.005 | 0.313 | 0.043 | 0.602 | 0.037 |
| 34 | 29 | CBTA | HF | 0.195 | 0.111 | 0.002 | 0.649 | 0.044 |
| 35 | 25 | Chacamax | HF | 0.000 | 0.000 | 0.000 | 0.700 | 0.299 |

| | | | | | | | | |
|---|---|---|---|---|---|---|---|---|
| 36 | 35 | Chacamax | HF | 0.001 | 0.269 | 0.001 | 0.727 | 0.002 |
| 37 | 34 | Chacamax | HF | 0.001 | 0.001 | 0.404 | 0.515 | 0.081 |
| 38 | 35 | Chacamax | HF | 0.000 | 0.003 | 0.000 | 0.715 | 0.282 |
| 39 | 61 | Chan Kha | LF | 0.001 | 0.003 | 0.001 | 0.930 | 0.066 |
| 40 | 36 | Chan Kha | LF | 0.045 | 0.007 | 0.003 | 0.909 | 0.037 |
| 41 | 64 | El Panchan | LF | 0.036 | 0.010 | 0.284 | 0.665 | 0.004 |
| 42 | 50 | El Panchan | LF | 0.002 | 0.006 | 0.261 | 0.624 | 0.107 |
| 43 | 68 | El Panchan | LF | 0.001 | 0.156 | 0.002 | 0.198 | 0.643 |
| 44 | 62 | El Panchan | LF | 0.002 | 0.003 | 0.157 | 0.831 | 0.008 |
| 45 | 25 | El Panchan | LF | 0.069 | 0.074 | 0.004 | 0.741 | 0.112 |
| 46 | 47 | La Mision | HF | 0.843 | 0.004 | 0.001 | 0.117 | 0.035 |
| 47 | 45 | La Mision | HF | 0.000 | 0.271 | 0.001 | 0.706 | 0.023 |
| 48 | 60 | Quiloma | HF | 0.001 | 0.001 | 0.002 | 0.941 | 0.055 |
| 49 | 31 | Leon Brindis | HF | 0.000 | 0.213 | 0.067 | 0.074 | 0.646 |
| 50 | 36 | Leon Brindis | HF | 0.000 | 0.000 | 0.000 | 0.700 | 0.300 |
| 51 | 37 | Nututun | HF | 0.062 | 0.002 | 0.001 | 0.869 | 0.067 |
| 52 | 48 | 35 | LF | 0.001 | 0.763 | 0.001 | 0.233 | 0.002 |
| 53 | 72 | 35 | LF | 0.027 | 0.850 | 0.002 | 0.119 | 0.002 |
| 54 | 39 | 35 | LF | 0.002 | 0.980 | 0.001 | 0.001 | 0.017 |
| 55 | 55 | 35 | LF | 0.042 | 0.933 | 0.000 | 0.002 | 0.023 |
| 56 | 67 | 35 | LF | 0.432 | 0.292 | 0.003 | 0.215 | 0.059 |
| 57 | 69 | 35 | LF | 0.002 | 0.885 | 0.001 | 0.111 | 0.002 |
| 58 | 53 | 35 | LF | 0.053 | 0.923 | 0.001 | 0.009 | 0.015 |
| 59 | 57 | 35 | LF | 0.093 | 0.336 | 0.162 | 0.216 | 0.193 |
| 60 | 72 | 35 | LF | 0.418 | 0.316 | 0.027 | 0.198 | 0.040 |
| 61 | 58 | 35 | LF | 0.295 | 0.608 | 0.004 | 0.055 | 0.039 |
| 62 | 56 | 35 | LF | 0.120 | 0.001 | 0.001 | 0.861 | 0.018 |
| 63 | 30 | 35 | LF | 0.000 | 0.999 | 0.000 | 0.000 | 0.000 |
| 64 | 60 | 35 | LF | 0.002 | 0.007 | 0.002 | 0.948 | 0.042 |
| 65 | 73 | 35 | LF | 0.064 | 0.139 | 0.069 | 0.677 | 0.051 |
| 66 | 61 | 37 | LF | 0.029 | 0.248 | 0.058 | 0.020 | 0.645 |
| 67 | 50 | 37 | LF | 0.001 | 0.993 | 0.001 | 0.000 | 0.005 |
| 68 | 42 | 37 | LF | 0.139 | 0.003 | 0.006 | 0.773 | 0.079 |
| 69 | 45 | 37 | LF | 0.001 | 0.144 | 0.003 | 0.833 | 0.020 |
| 70 | 31 | 37 | LF | 0.193 | 0.448 | 0.100 | 0.063 | 0.197 |
| 71 | 25 | 37 | LF | 0.000 | 0.279 | 0.065 | 0.008 | 0.648 |
| 72 | 28 | 37 | LF | 0.080 | 0.412 | 0.007 | 0.484 | 0.016 |
| 73 | 68 | 37 | LF | 0.020 | 0.012 | 0.538 | 0.413 | 0.017 |
| 74 | 24 | 37 | LF | 0.000 | 0.999 | 0.000 | 0.000 | 0.000 |
| 75 | 21 | 37 | LF | 0.119 | 0.125 | 0.030 | 0.520 | 0.206 |
| 76 | 32 | 37 | LF | 0.159 | 0.416 | 0.264 | 0.039 | 0.122 |
| 77 | 47 | 38 | LF | 0.003 | 0.001 | 0.001 | 0.941 | 0.055 |
| 78 | 48 | 38 | LF | 0.000 | 0.004 | 0.007 | 0.987 | 0.002 |

| | | | | | | | | |
|---|---|---|---|---|---|---|---|---|
| 79 | 57 | 38 | LF | 0.001 | 0.236 | 0.008 | 0.336 | 0.420 |
| 80 | 30 | 38 | LF | 0.001 | 0.352 | 0.002 | 0.001 | 0.645 |
| 81 | 33 | 38 | LF | 0.000 | 0.285 | 0.001 | 0.087 | 0.627 |
| 82 | 44 | 38 | LF | 0.195 | 0.252 | 0.044 | 0.002 | 0.508 |
| 83 | 40 | 38 | LF | 0.002 | 0.153 | 0.023 | 0.297 | 0.526 |
| 84 | 56 | 38 | LF | 0.005 | 0.158 | 0.091 | 0.222 | 0.524 |
| 85 | 25 | 38 | LF | 0.000 | 0.234 | 0.009 | 0.110 | 0.647 |
| 86 | 31 | 38 | LF | 0.000 | 0.350 | 0.000 | 0.000 | 0.650 |
| 87 | 24 | 38 | LF | 0.000 | 0.345 | 0.005 | 0.001 | 0.649 |
| 88 | 28 | 39 | LF | 0.925 | 0.042 | 0.001 | 0.001 | 0.032 |
| 89 | 55 | 10 | HF | 0.003 | 0.900 | 0.001 | 0.093 | 0.004 |
| 90 | 66 | 23 | HF | 0.420 | 0.369 | 0.015 | 0.146 | 0.049 |
| 91 | 33 | 23 | HF | 0.603 | 0.317 | 0.005 | 0.006 | 0.070 |
| 92 | 28 | 23 | HF | 0.000 | 0.994 | 0.000 | 0.003 | 0.003 |
| 93 | 51 | 29 | HF | 0.769 | 0.001 | 0.001 | 0.214 | 0.015 |
| 94 | 49 | 29 | HF | 0.431 | 0.011 | 0.002 | 0.538 | 0.019 |
| 95 | 57 | 41 | HF | 0.946 | 0.021 | 0.002 | 0.001 | 0.029 |
| 96 | 30 | 41 | HF | 0.525 | 0.288 | 0.117 | 0.032 | 0.038 |
| 97 | 65 | 43 | HF | 0.619 | 0.097 | 0.017 | 0.223 | 0.044 |
| 98 | 72 | 43 | HF | 0.047 | 0.114 | 0.007 | 0.813 | 0.020 |
| 99 | 67 | 45 | LF | 0.994 | 0.002 | 0.000 | 0.004 | 0.000 |
| 100 | 26 | 45 | LF | 0.000 | 0.173 | 0.413 | 0.001 | 0.413 |
| 101 | 37 | 46 | HF | 0.747 | 0.001 | 0.001 | 0.219 | 0.033 |
| 102 | 28 | 46 | HF | 0.949 | 0.000 | 0.000 | 0.000 | 0.050 |
| 103 | 42 | 46 | HF | 0.949 | 0.000 | 0.000 | 0.000 | 0.050 |
| 104 | 73 | 46 | HF | 0.913 | 0.007 | 0.016 | 0.010 | 0.055 |
| 105 | 46 | 46 | HF | 0.020 | 0.218 | 0.006 | 0.727 | 0.029 |
| 106 | 40 | 48 | LF | 0.879 | 0.007 | 0.013 | 0.086 | 0.016 |
| 107 | 62 | 48 | LF | 0.851 | 0.013 | 0.011 | 0.079 | 0.045 |
| 108 | 67 | 48 | LF | 0.313 | 0.023 | 0.112 | 0.483 | 0.069 |
| 109 | 31 | 48 | LF | 0.997 | 0.002 | 0.001 | 0.001 | 0.000 |
| 110 | 73 | 48 | LF | 0.150 | 0.010 | 0.321 | 0.512 | 0.007 |
| 111 | 68 | 48 | LF | 0.285 | 0.006 | 0.217 | 0.490 | 0.002 |
| 112 | 33 | 48 | LF | 0.401 | 0.001 | 0.598 | 0.000 | 0.000 |
| 113 | 68 | 48 | LF | 0.080 | 0.003 | 0.234 | 0.682 | 0.001 |
| 114 | 55 | 48 | LF | 0.472 | 0.002 | 0.003 | 0.515 | 0.008 |
| 115 | 50 | 48 | LF | 0.916 | 0.021 | 0.054 | 0.002 | 0.007 |
| 116 | 32 | 49 | HF | 0.989 | 0.003 | 0.001 | 0.001 | 0.006 |
| 117 | 53 | 49 | HF | 0.996 | 0.001 | 0.001 | 0.001 | 0.002 |
| 118 | 40 | 49 | HF | 0.997 | 0.001 | 0.001 | 0.000 | 0.000 |
| 119 | 51 | 50 | LF | 0.303 | 0.104 | 0.172 | 0.209 | 0.212 |
| 120 | 28 | 50 | LF | 0.565 | 0.250 | 0.009 | 0.051 | 0.126 |
| 121 | 29 | 38NB | HF | 0.000 | 0.322 | 0.029 | 0.000 | 0.649 |

| | | | | | | | | |
|---|---|---|---|---|---|---|---|---|
| 122 | 27 | 38NB | HF | 0.998 | 0.001 | 0.001 | 0.001 | 0.000 |
| 123 | 66 | 1 | LF | 0.207 | 0.031 | 0.318 | 0.375 | 0.069 |
| 124 | 64 | 1 | LF | 0.007 | 0.169 | 0.490 | 0.286 | 0.048 |
| 125 | 46 | 1 | LF | 0.000 | 0.001 | 0.003 | 0.819 | 0.177 |
| 126 | 30 | 1 | LF | 0.001 | 0.025 | 0.973 | 0.000 | 0.001 |
| 127 | 43 | 3 | LF | 0.004 | 0.002 | 0.519 | 0.474 | 0.002 |
| 128 | 48 | 3 | LF | 0.082 | 0.659 | 0.191 | 0.046 | 0.023 |
| 129 | 46 | 4 | HF | 0.313 | 0.004 | 0.682 | 0.001 | 0.000 |
| 130 | 33 | 4 | HF | 0.251 | 0.081 | 0.642 | 0.002 | 0.024 |
| 131 | 69 | BANO REINA | PNP | 0.456 | 0.033 | 0.496 | 0.002 | 0.013 |
| 132 | 50 | BOLAS | PNP | 0.000 | 0.000 | 0.999 | 0.000 | 0.000 |
| 133 | 42 | BOLAS | PNP | 0.000 | 0.000 | 0.999 | 0.000 | 0.000 |
| 134 | 57 | CALAVERA | PNP | 0.000 | 0.101 | 0.662 | 0.001 | 0.236 |
| 135 | 62 | CALAVERA | PNP | 0.015 | 0.135 | 0.687 | 0.002 | 0.161 |
| 136 | 31 | CALAVERA | PNP | 0.001 | 0.105 | 0.680 | 0.001 | 0.214 |
| 137 | 56 | EDGIES/NAHA | PNP | 0.003 | 0.114 | 0.505 | 0.248 | 0.131 |
| 138 | 45 | GROUP C2 | PNP | 0.194 | 0.115 | 0.488 | 0.048 | 0.157 |
| 139 | 27 | GROUP C2 | PNP | 0.001 | 0.161 | 0.649 | 0.001 | 0.189 |
| 140 | 44 | GROUP C2 | PNP | 0.015 | 0.127 | 0.398 | 0.237 | 0.223 |
| 141 | 61 | JAGUAR1 | PNP | 0.002 | 0.152 | 0.829 | 0.001 | 0.016 |
| 142 | 52 | MIGHTY DUC | PNP | 0.016 | 0.457 | 0.510 | 0.001 | 0.016 |
| 143 | 41 | MOTIEPA | PNP | 0.022 | 0.001 | 0.974 | 0.001 | 0.003 |
| 144 | 74 | MOTIEPA | PNP | 0.000 | 0.183 | 0.813 | 0.002 | 0.003 |
| 145 | 72 | MUSEUM | PNP | 0.314 | 0.026 | 0.629 | 0.003 | 0.027 |
| 146 | 58 | PAKAL | PNP | 0.000 | 0.005 | 0.992 | 0.000 | 0.003 |
| 147 | 72 | PIGRITOS | PNP | 0.002 | 0.160 | 0.549 | 0.004 | 0.286 |
| 148 | 62 | T NORTE | PNP | 0.000 | 0.001 | 0.990 | 0.001 | 0.008 |
| 149 | 49 | T NORTE | PNP | 0.000 | 0.000 | 0.999 | 0.000 | 0.000 |
| 150 | 48 | T NORTE | PNP | 0.000 | 0.000 | 0.999 | 0.000 | 0.000 |
| 151 | 32 | T NORTE | PNP | 0.000 | 0.000 | 0.999 | 0.000 | 0.000 |
| 152 | 67 | TOLVIES | PNP | 0.010 | 0.100 | 0.724 | 0.024 | 0.143 |
| 153 | 52 | UNITES | PNP | 0.004 | 0.139 | 0.645 | 0.001 | 0.210 |
| 154 | 42 | UNITES | PNP | 0.000 | 0.057 | 0.881 | 0.001 | 0.062 |
| 155 | 28 | UNITES | PNP | 0.000 | 0.000 | 0.999 | 0.000 | 0.000 |
| 156 | 71 | UNITES | PNP | 0.225 | 0.080 | 0.530 | 0.132 | 0.033 |
| 157 | 31 | UNITES | PNP | 0.000 | 0.000 | 0.999 | 0.000 | 0.000 |
| 158 | 50 | WUPSIES | PNP | 0.013 | 0.161 | 0.674 | 0.004 | 0.148 |
| | | | **Region** | dark purple | light purple | orange | blue | green |
| | | | **PNP** | 0.05 | 0.09 | 0.76 | 0.03 | 0.08 |
| | | | **LF** | 0.34 | 0.18 | 0.12 | 0.22 | 0.14 |
| | | | **HF** | 0.10 | 0.18 | 0.09 | 0.53 | 0.10 |

**B)** Summary by cluster - proportion of each STRUCTURE cluster found in each region; left - including only individuals with a very clear major cluster assignment (q>0.75); right - including individuals with a lower threshold for clear major cluster assignment (q>0.6)

**breakdown of clusters by region (IND with q >75%)**

| | HF | LF | PNP |
|---|---|---|---|
| dark purple | 0.65 | 0.35 | 0 |
| light purple | 0.18 | 0.82 | 0 |
| orange | 0.07 | 0.07 | 0.87 |
| blue | 0.62 | 0.38 | 0 |
| green | 0 | 0 | 0 |

**breakdown of clusters by region (IND with q >60%**

| | HF | LF | PNP |
|---|---|---|---|
| dark purple | 0.68 | 0.32 | 0 |
| light purple | 0.21 | 0.79 | 0 |
| Orange | 0.12 | 0.04 | 0.84 |
| Blue | 0.64 | 0.36 | 0 |
| Green | 0.2 | 0.8 | 0 |

**C)** Summary by region - proportion of individuals in each region assigned to each STRUCTURE cluster; left - including only individuals with a very clear major cluster assignment (q>0.75); right - all individuals, including highly admixed individuals with no clear major cluster assignment.

| **breakdown of regions by cluster (including only IND with q >75%)** | | | | | |
|---|---|---|---|---|---|
| | dark purple | light purple | orange | blue | green |
| **PNP** | 0 | 0 | 1 | 0 | 0 |
| **LF** | 0.23 | 0.35 | 0.04 | 0.38 | 0 |
| **HF** | 0.34 | 0.07 | 0.03 | 0.55 | 0 |

| **breakdown of regions by cluster - all IND** | | | | | |
|---|---|---|---|---|---|
| | dark purple | light purple | orange | blue | green |
| **PNP** | 0.05 | 0.09 | 0.76 | 0.03 | 0.08 |
| **LF** | 0.17 | 0.25 | 0.11 | 0.31 | 0.16 |
| **HF** | 0.26 | 0.11 | 0.10 | 0.44 | 0.09 |

**Relatedness analyses**

**Fig. S3. Heatmap of all Lynch-Ritland pairwise relatedness values.** Colors correspond to values in gradient bar on right. Each row/column is a single individual, and they are ordered by social group and then by fragment. Reflecting patterns of higher within-group and fragment relatedness, green, yellow, and red cells (denoting higher pairwise relatedness values), are clustered along the diagonal; a number of sampling locations are marked in yellow boxes and labeled to emphasize these clusters of more highly related individuals. Three examples of highly related pairs sampled in different fragments are denoted with yellow arrows.

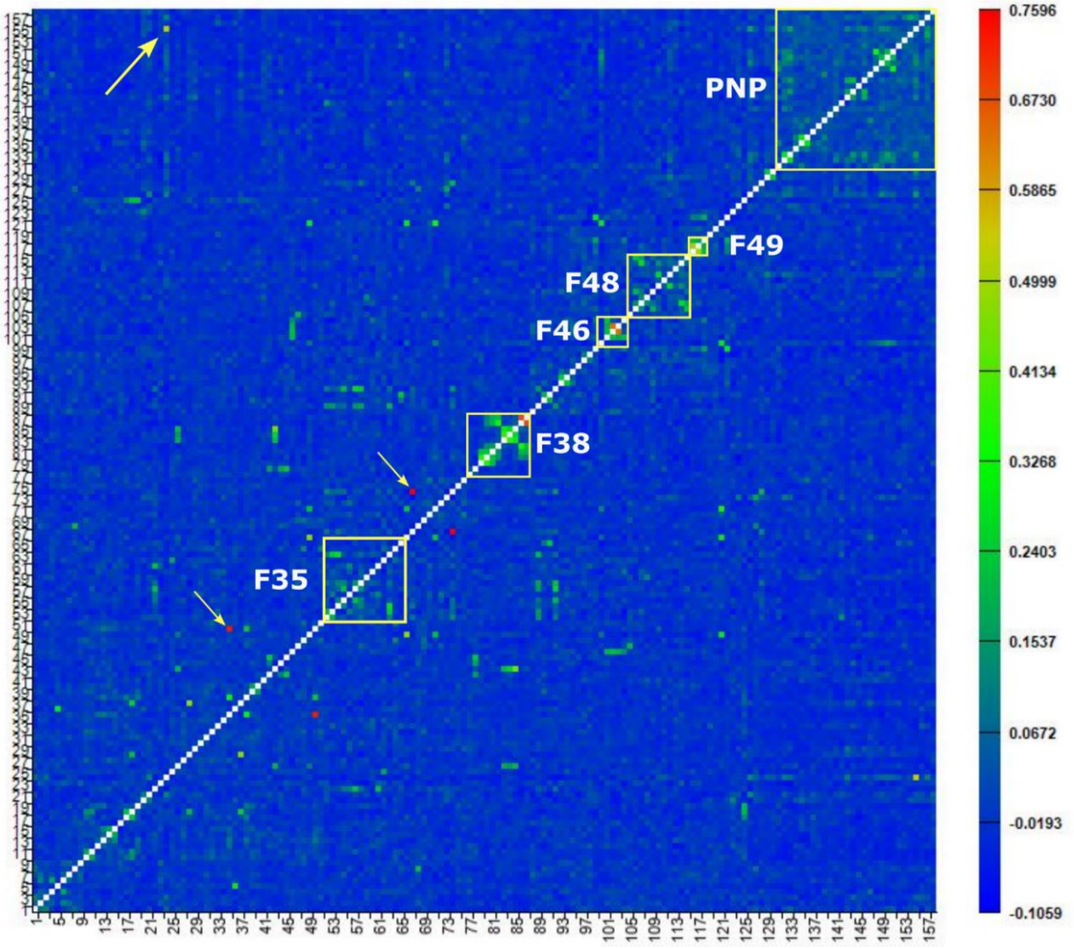

**Table S7. Input data for the Lynch-Ritland by-individual pairwise genetic relatedness network of individuals sampled in different locations.** In orange – top 0.25% of values; in green – bottom 0.25% of values. Acronyms: juvenile (J); adult (A); subadult (SA); male (M); female (F); lone subadult female (LSAF); between fragments (BF); fragment-PNP (FPNP).

| IND1 | Fragment | Group | Age | Sex | IND2 | Fragment | Group | Age | Sex | LR pairwise relatedness | category |
|---|---|---|---|---|---|---|---|---|---|---|---|
| A35 | Chacamax | CM | J | M | K45 | Leon Brindis | LB | A | M | 0.7295 | BF |
| K22 | Artisanias | AR4 | A | F | SVB-202 | PNP | Unites | A | M | 0.5344 | FPNP |
| K6 | Camino Real | CR | A | M | A37 | Chacamax | CM | A | F | 0.4842 | BF |
| K43 | Leon Brindis | LB | A | F | AC7 | 37 | 37D | A | F | 0.4759 | BF |
| K58 | PNP | EP | J | M | SM27 | 38 | 38C | SA | F | 0.454 | BF |
| KK16 | 37 | 37C | SA | M | CA22 | 38NB | 38NB | A | M | 0.3508 | BF |
| AC22 | 13 | 13B | A | F | A36 | Chacamax | CM | J | M | 0.3106 | BF |
| K43 | Leon Brindis | LB | A | F | CA22 | 38NB | 38NB | A | M | 0.3014 | BF |
| AC7 | 37 | 37D | A | F | CA22 | 38NB | 38NB | A | M | 0.3013 | BF |
| DL60 | 45 | 45B | A | F | CA22 | 38NB | 38NB | A | M | 0.299 | BF |
| DL57 | 45 | 45B | A | F | KK100 | 38NB | 38NB | J | M | 0.295 | BF |
| KK94 | 35 | 35A | A | M | KK107 | 23 | 23A | J | F | 0.2669 | BF |
| K58 | PNP | EP | J | M | SM26 | 38 | 38C | A | M | 0.2634 | BF |
| DL37 | 25 | 25A | A | M | A37 | Chacamax | CM | A | F | 0.2554 | BF |
| A38 | Chacamax | CM | A | M | K45 | Leon Brindis | LB | A | M | 0.2475 | BF |
| SM39 | 35 | 35F | A | F | CA26 | 23 | 23A | A | F | 0.2395 | BF |
| K60 | PNP | EP | A | M | CA11 | 38 | 38C | A | F | 0.2366 | BF |
| A13 | Artisanias | A4 | SA | F | SM34 | 35 | 35C | A | F | 0.2325 | BF |
| K43 | Leon Brindis | LB | A | F | KK16 | 37 | 37C | SA/J | M | 0.232 | BF |
| A13 | Artisanias | A4 | SA | F | KK94 | 35 | 35A | A | M | 0.2318 | BF |
| KK96 | 35 | 35B | A | M | KK107 | 23 | 23A | J | F | 0.2295 | BF |
| DL37 | 25 | 25A | A | M | K6 | Camino Real | CR | A | M | 0.2288 | BF |
| A32 | La Mision | LM | J | F | DL55 | 46 | 46A | A | F | 0.2267 | BF |
| KK27 | 25 | 25A | A | F | KK76 | 1 | 1C | A | F | 0.2222 | BF |
| K34 | Artisanias | LSAF | SA | F | SM27 | 38 | 38C | SA | F | 0.2215 | BF |
| DL37 | 25 | 25A | A | M | KK76 | 1 | 1C | A | F | 0.221 | BF |
| K31 | La Mision | LM | A | M | KK37 | 46 | 46A | A | F | 0.2192 | BF |
| DL23 | 16 | 16A | A | F | DL14 | 37 | 37D | A | F | 0.2174 | BF |

| | | | | | | | | | | | |
|---|---|---|---|---|---|---|---|---|---|---|---|
| CA20 | 35 | 35F | A | M | KK107 | 23 | 23A | J | F | 0.2163 | BF |
| DL89 | 48 | 48E | A | F | CA22 | 38NB | 38NB | A | M | -0.1059 | BF |
| DL57 | 45 | 45B | A | F | SVB-103 | PNP | Jaguar1 | J | M | -0.1009 | FPNP |
| CA9 | 38 | 38A | SA | M | DL89 | 48 | 48E | A | F | -0.0976 | BF |
| KK64 | 43 | 43B | SA | M | SVB-198 | PNP | Bolas | A | F | -0.0964 | FPNP |
| CA9 | 38 | 38A | SA | M | KK74 | 1 | 1B | A | F | -0.0961 | BF |
| SM39 | 35 | 35F | A | F | DL60 | 45 | 45B | A | F | -0.0915 | BF |
| TP3 | 29 | 29A | J | M | SVB-112 | PNP | Pigritos | A | F | -0.0912 | FPNP |
| K58 | PNP | EP | J | M | SVB-156 | PNP | T Norte | A | F | -0.0911 | FPNP |
| DL36 | 25 | 25A | A | M | SVB-134 | PNP | Museum | J | F | -0.0894 | FPNP |
| SM39 | 35 | 35F | A | F | SVB-151 | PNP | Bano Reina | J | F | -0.0894 | FPNP |
| CA14 | 35 | 35B | A | F | SVB-196 | PNP | Bolas | A | M | -0.0887 | FPNP |
| CA17 | 35 | 35B | A | F | KK60 | 1 | 1A | SA | M | -0.0886 | BF |
| SM29 | 38 | 38A | A | F | DL89 | 48 | 48E | A | F | -0.0877 | BF |
| CA17 | 35 | 35B | A | F | KK36 | 46 | 46A | A | M | -0.0876 | BF |
| CA17 | 35 | 35B | A | F | SVB-199 | PNP | Unites | A | F | -0.0872 | FPNP |
| K6 | Camino Real | CR | A | M | KK96 | 35 | 35B | A | M | -0.087 | BF |
| A16 | Artisanias | A4 | ? | F | SVB-198 | PNP | Bolas | A | F | -0.087 | FPNP |
| K60 | PNP | EP | A | M | SVB-134 | PNP | Museum | J | F | -0.0863 | FPNP |
| AC44 | 43 | 43B | A | M | CA22 | 38NB | 38NB | A | M | -0.086 | BF |
| DL87 | 48 | 48D | A | F | SVB-151 | PNP | Bano Reina | J | F | -0.0858 | FPNP |
| SM29 | 38 | 38A | A | F | AC44 | 43 | 43B | A | M | -0.0856 | BF |
| DL60 | 45 | 45B | A | F | SVB-62 | PNP | Motiepa | A | F | -0.0854 | FPNP |
| CA22 | 38NB | 38NB | A | M | SVB-62 | PNP | Motiepa | A | F | -0.0852 | FPNP |
| A16 | Artisanias | A4 | ? | F | SVB-156 | PNP | T Norte | A | F | -0.0851 | FPNP |
| LE16 | 25 | 20B | A | M | CA26 | 23 | 23A | A | F | -0.085 | BF |
| LE16 | 25 | 20B | A | M | DL75 | 41 | 41A | A | M | -0.0841 | BF |
| A35 | Chacamax | CM | J | M | SVB-156 | PNP | T Norte | A | F | -0.0841 | FPNP |
| AC30 | 41 | 41A | A | F | SVB-126 | PNP | Tolvies | J | M | -0.0837 | FPNP |
| SM35 | 35 | 35C | J | F | SM9 | 1 | 1C | A | M | -0.0835 | BF |

**Table S8. Demographic composition of top 1% most highly related pairs and comparison to composition of whole dataset.** Abbreviations: adult (A); adult female (AF); adult male (AM), immatures – all subadults, juveniles, and infants (IMM).

A) Number of each type of pair from the whole dataset (N=12403 pairs)

| | Pair type | AF-AF | AF-AM | AM-AM | A-IMM | IMM-IMM | total | | |
|---|---|---|---|---|---|---|---|---|---|
| PNP | Within-group | 3 | 9 | 1 | 10 | 1 | 24 | Total pairs in PNP | 378 |
| | Between-group in PNP | 42 | 61 | 20 | 177 | 54 | 354 | | |
| Fragments | Within-group | 19 | 37 | 6 | 46 | 6 | 114 | Total pairs in fragmented landscape | 8384 |
| | Between-group in fragment | 44 | 53 | 22 | 90 | 14 | 223 | | |
| | Between-fragment | 1769 | 2095 | 602 | 2968 | 613 | 8047 | Total pairs between fragments-PNP | 3641 |
| Between fragment-PNP | | 610 | 781 | 252 | 1611 | 387 | 3641 | | |
| Total | | 2487 | 3036 | 903 | 4902 | 1075 | 12403 | | |

B) Proportion of each type of pair from the whole dataset (N=12403 pairs)

| | Pair type | AF-AF | AF-AM | AM-AM | A-IMM | IMM-IMM | total | | |
|---|---|---|---|---|---|---|---|---|---|
| PNP | Within-group | 0.0002 | 0.0007 | 0.0001 | 0.0008 | 0.0001 | 0.0019 | Total pairs in PNP | 0.0305 |
| | Between-group in PNP | 0.0034 | 0.0049 | 0.0016 | 0.0143 | 0.0044 | 0.0285 | | |
| Fragments | Within-group | 0.0015 | 0.0030 | 0.0005 | 0.0037 | 0.0005 | 0.0092 | Total pairs in fragmented landscape | 0.6760 |
| | Between-group in fragment | 0.0035 | 0.0043 | 0.0018 | 0.0073 | 0.0011 | 0.0180 | | |
| | Between-fragment | 0.1426 | 0.1689 | 0.0485 | 0.2393 | 0.0494 | 0.6488 | Total pairs between fragments-PNP | 0.2936 |
| Between fragment-PNP | | 0.0492 | 0.0630 | 0.0203 | 0.1299 | 0.0312 | 0.2936 | | |
| Total | | 0.2005 | 0.2448 | 0.0728 | 0.3952 | 0.0867 | 1 | | |

C) Proportion of top 1% most related pairs in each category (N=124 pairs)

| | Pair type | AF-AF | AF-AM | AM-AM | A-IMM | IMM-IMM | Total | | |
|---|---|---|---|---|---|---|---|---|---|
| PNP | Within-group | 0.0081 | 0.0161 | 0 | 0.0403 | 0 | 0.0645 | Total pairs in PNP | 0.1371 |
| | Between-group in PNP | 0.0161 | 0.0323 | 0 | 0.0242 | 0 | 0.0726 | | |
| Fragments | Within-group | 0.0645 | 0.0645 | 0.0081 | 0.1210 | 0.0161 | 0.2742 | Total pairs in fragmented landscape | 0.8226 |
| | Between-group in fragment | 0 | 0.0081 | 0.0161 | 0.0726* | 0.0081* | 0.1048 | | |
| | Between-fragment | 0.0565 | 0.1371 | 0.0242 | 0.1694 | 0.0565 | 0.4435 | Total pairs between fragments-PNP | 0.0403 |
| Between fragment-PNP | | 0.0081 | 0.0161 | 0.0081 | 0.0081 | 0 | 0.0403 | | |
| Total | | 0.1532 | 0.2742 | 0.0565 | 0.4355 | 0.0806 | 1 | | |

*All of these pairs include sub-adult males (adult-subadult male or subadult male-subadult male)

**D)** Proportions that differed significantly between whole dataset and top 1%. Proportions that were significantly higher in top 1% - in blue; proportions that were significantly lower in top 1% - in red.

| | **Pair type** | **AF-AF** | **AF-AM** | **AM-AM** | **A-IMM** | **IMM-IMM** | **Total (all pair types)** | | |
|---|---|---|---|---|---|---|---|---|---|
| **PNP** | **Within-group** | **Z= 4.85, p< .001** | **Z=5.76, p<.001** | Z=0.1, p= .920 | **Z=12.66, p<.001** | Z=0.1, p=.920 | **Z=13.74, p<.001** | **Total pairs in PNP** | **Z=6.76, p<.001** |
| | **Between-group in PNP** | **Z= 2.39, p= .017** | **Z=4.22, p< .001** | Z=0.45, p= .655 | Z=0.92, p= .356 | Z=0.74, p=.462 | **Z=2.91, p=.004** | | |
| **Fragments** | **Within-group** | **Z=15.05, p<.001** | **Z=11.4, p< .001** | **Z=3.55, p< .001** | **Z=18.66, p< .001** | **Z=6.86, p<.001** | **Z=27.18, p<.001** | **Total pairs fragmented landscape** | **Z=3.48, p<.001** |
| | **Between-group in fragment** | Z=0.66, p= .506 | Z= 0.64, p= .521 | **Z=3.64, p< .001** | **Z=8.17, p< .001** | **Z=2.22, p=.026** | **Z=7.08, p<.001** | | |
| | **Between-fragment** | **Z= 2.74, p= .006** | Z= 0.94, p =.346 | Z= 1.26, p= .208 | Z=1.82, p=.069 | Z =0.36, p=.720 | **Z=4.76, p<.001** | **Total pairs between fragments-PNP** | **Z=6.18, p<.001** |
| **Between fragment-PNP** | | **Z= 2.12, p= .034** | **Z=2.14, p= .032** | Z= 0.97, p= .334 | **Z=4.03, p< .001** | **Z=2, p =.046** | **Z=6.18, p<.001** | | |

When examining the composition of the top 1% of pairs with the highest relatedness values in terms of demography and category (pairs of individuals from the same social group, from different groups within the same fragment or within PNP, and from different fragments or fragments and PNP), we found that the top 1% had significantly fewer than expected between-fragment and between fragment-PNP pairs, and significantly more within-group and between-group (within the same fragment or within PNP) pairs. There were differences between adult males and females as well: while between-fragment AF-AF pairs were significantly underrepresented in the top 1% of relatedness values, we found a significantly higher than expected proportion of highly related AM-AM between-group pairs in the same fragment in the fragmented landscape; higher relatedness between males within fragments was further supported by significantly higher than expected proportions of highly related adult - subadult male and subadult male - subadult male pairs in different groups in the same fragment among the top 1% most highly related pairs (as dispersal in black howler monkeys typically occurs around the age of sexual maturity or during adulthood [Van Belle and Di Fiore 2022], these subadult males may have been sampled post-natal dispersal).

**Fig. S4. Pairwise relatedness across socio-spatial categories in the fragmented landscape vs. the continuous forest (PNP).** Fragmented landscape on left (figures A,C); PNP on right (figures B,D). E) Comparing within-group relatedness in groups in PNP vs. in fragments (left), and between-group relatedness in PNP vs. between different groups in individual fragments. Note different ranges of Lynch-Ritland (LR) pairwise relatedness values on y-axes. Statistical significance (p-value) of differences in plots A, B, E noted with asterisks: * = 0.01 – 0.05; ** = 0.001 – 0.01; *** = 0 – 0.001.

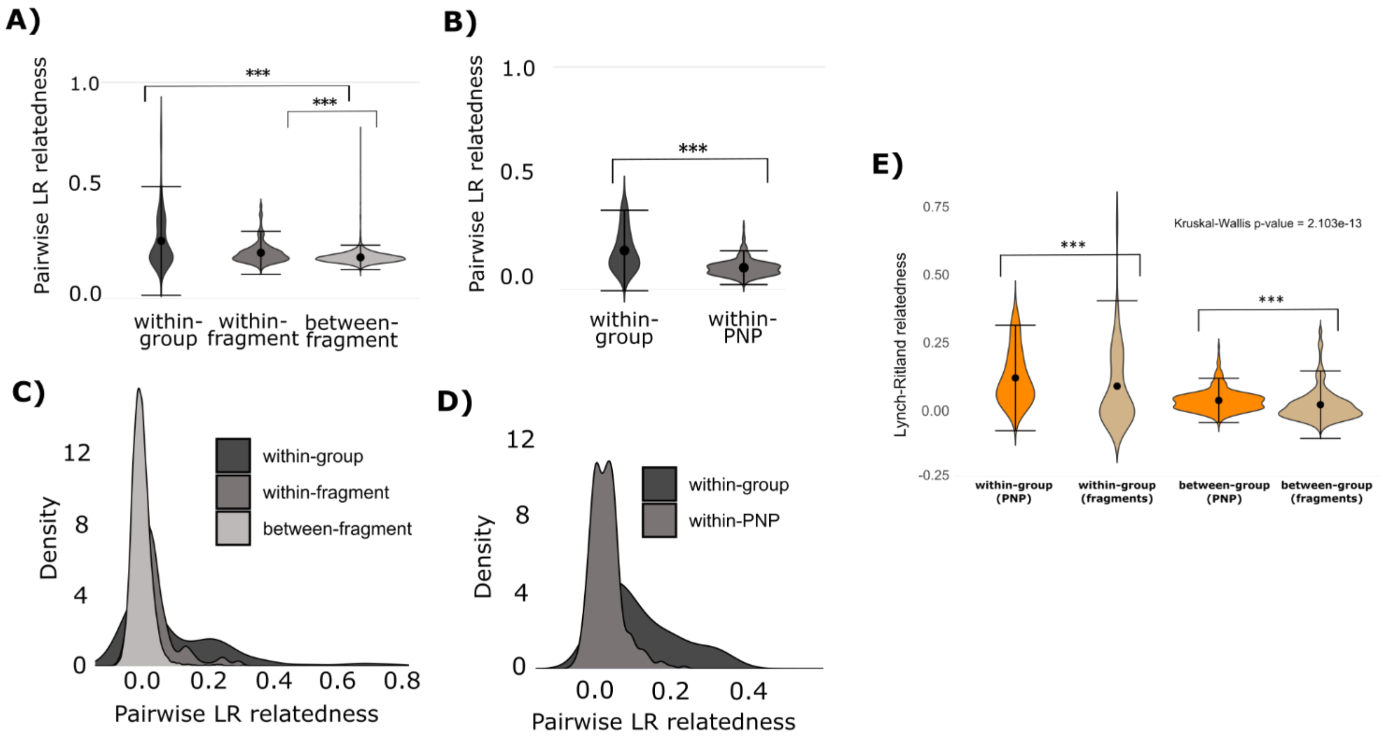

#### Gene flow analyses

**Fig. S5. Supporting graphs for EEMS results for three merged chains of 600 demes.** See SI Methods for detailed descriptions of each graph.

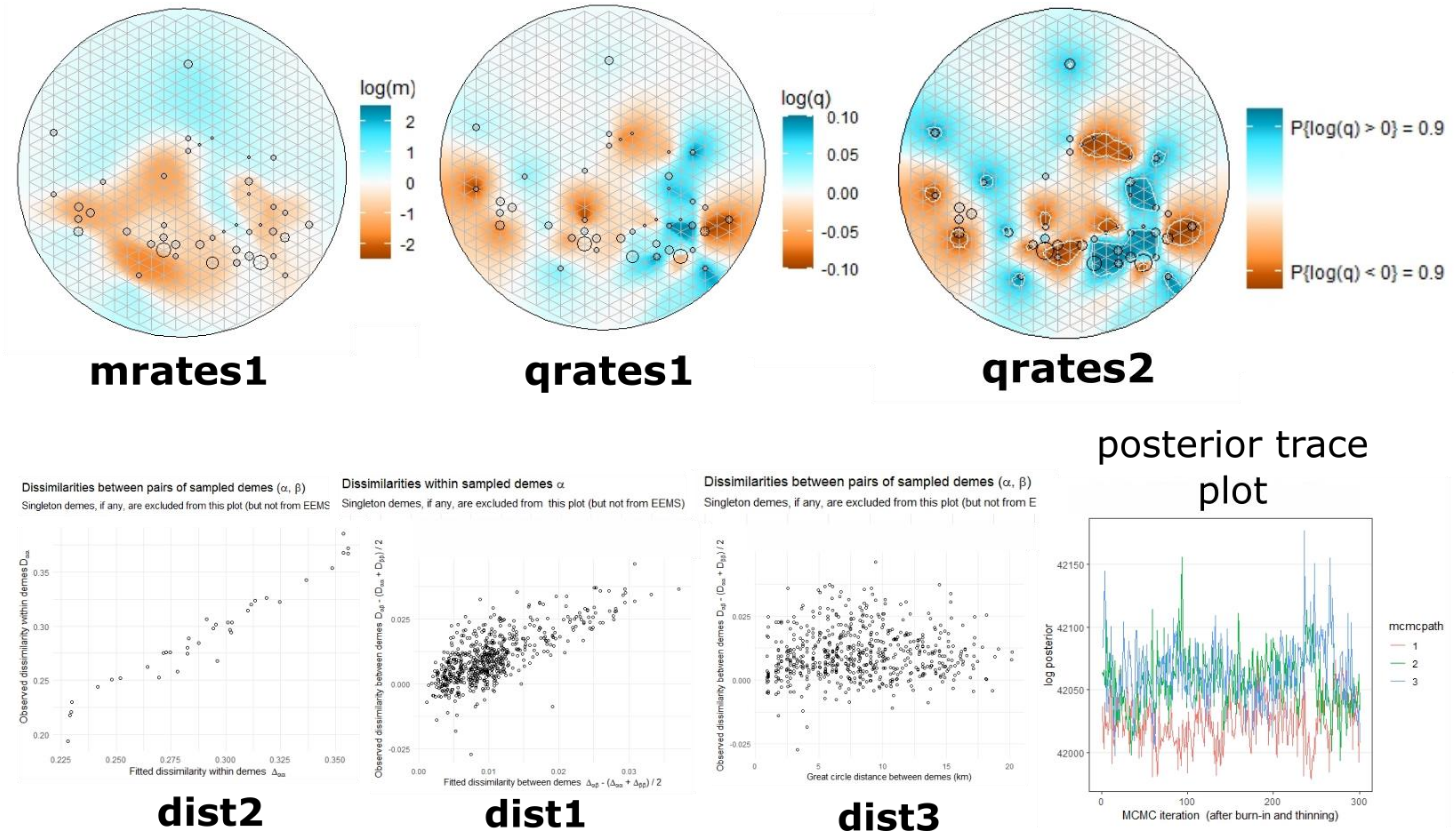

**Fig. S6. EEMS plots of merged results of 100-800 x3 deme chains,** overlaid on the sampling grid for 600 demes to facilitate comparisons to Fig. 2 and Fig. S15. See SI Methods for detailed descriptions of graphs.

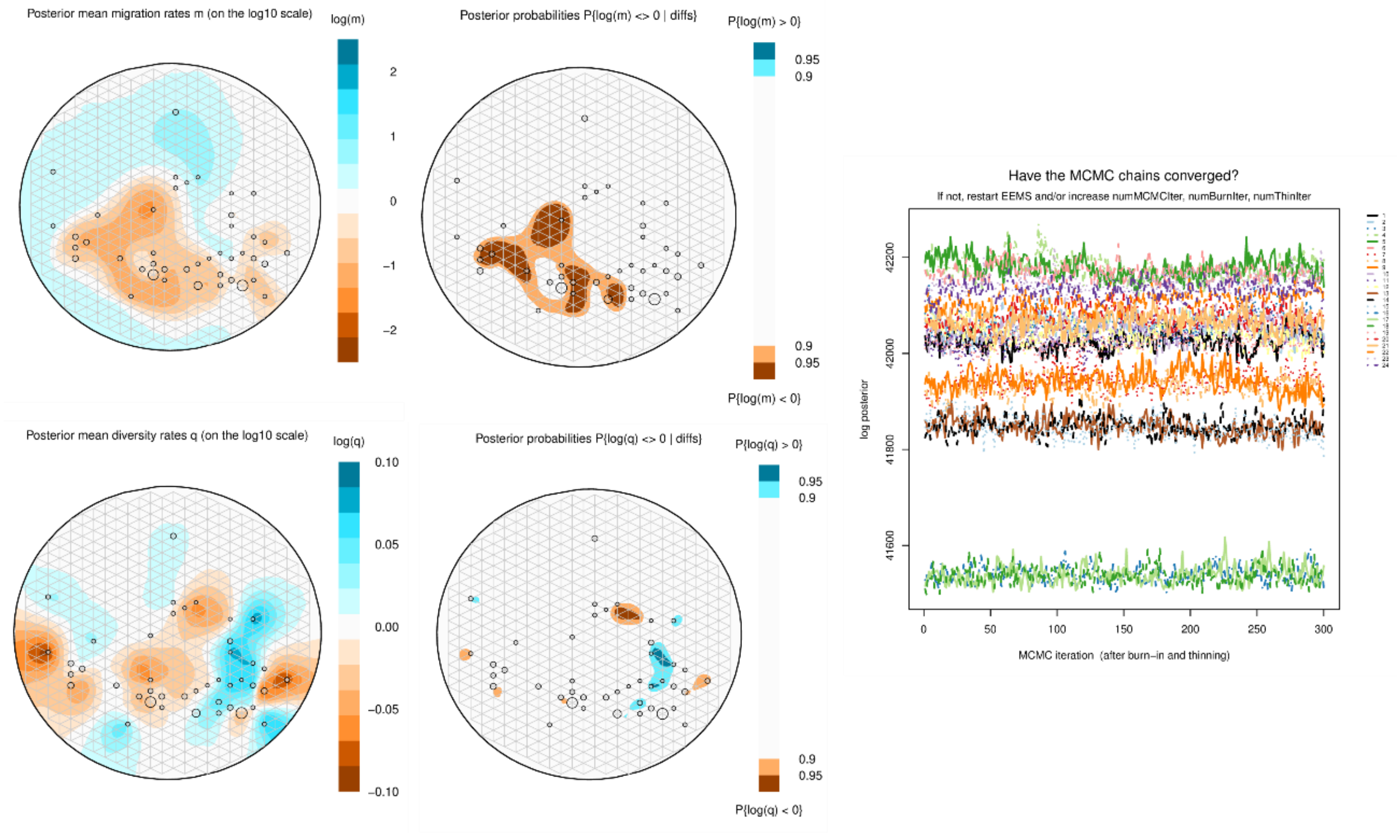

#### Isolation-by-distance (IBD) and spatial autocorrelation analyses

**Table S9. Summary of sample sizes for maximum-likelihood population effects (MLPE) tests of isolation-by-distance (IBD).**

Full spatial extents

| | all adults and subadults | males only | females only |
|---|---|---|---|
| **PNP** | 20 | 8 | 12 |
| **whole fragmented landscape** | 118 | 50 | 68 |
| **LF region** | 58 | 25 | 33 |
| **HF region** | 60 | 25 | 35 |

Subsets of HF, LF regions to match 2km extent of PNP

| | all adults and subadults | | males only | | females only | |
|---|---|---|---|---|---|---|
| | **individuals** | **pairs** | **individuals** | **pairs** | **individuals** | **pairs** |
| **whole fragmented landscape** | 116 | 759 | 46 | 119 | 61 | 256 |
| **LF region** | 58 | 485 | 24 | 74 | 32 | 169 |
| **HF region** | 58 | 127 | 18 | 21 | 28 | 44 |

**Fig. S7. Results of maximum-likelihood population effects (MLPE) models**, implemented as linear mixed-effects models, quantifying isolation-by-distance (IBD) within each region (PNP, LF, HF) for males and females combined. A) At full landscape extent for each region; B) HF, LF data subset to 2km extent, to facilitate direct comparisons to PNP's 2km sampling scale.

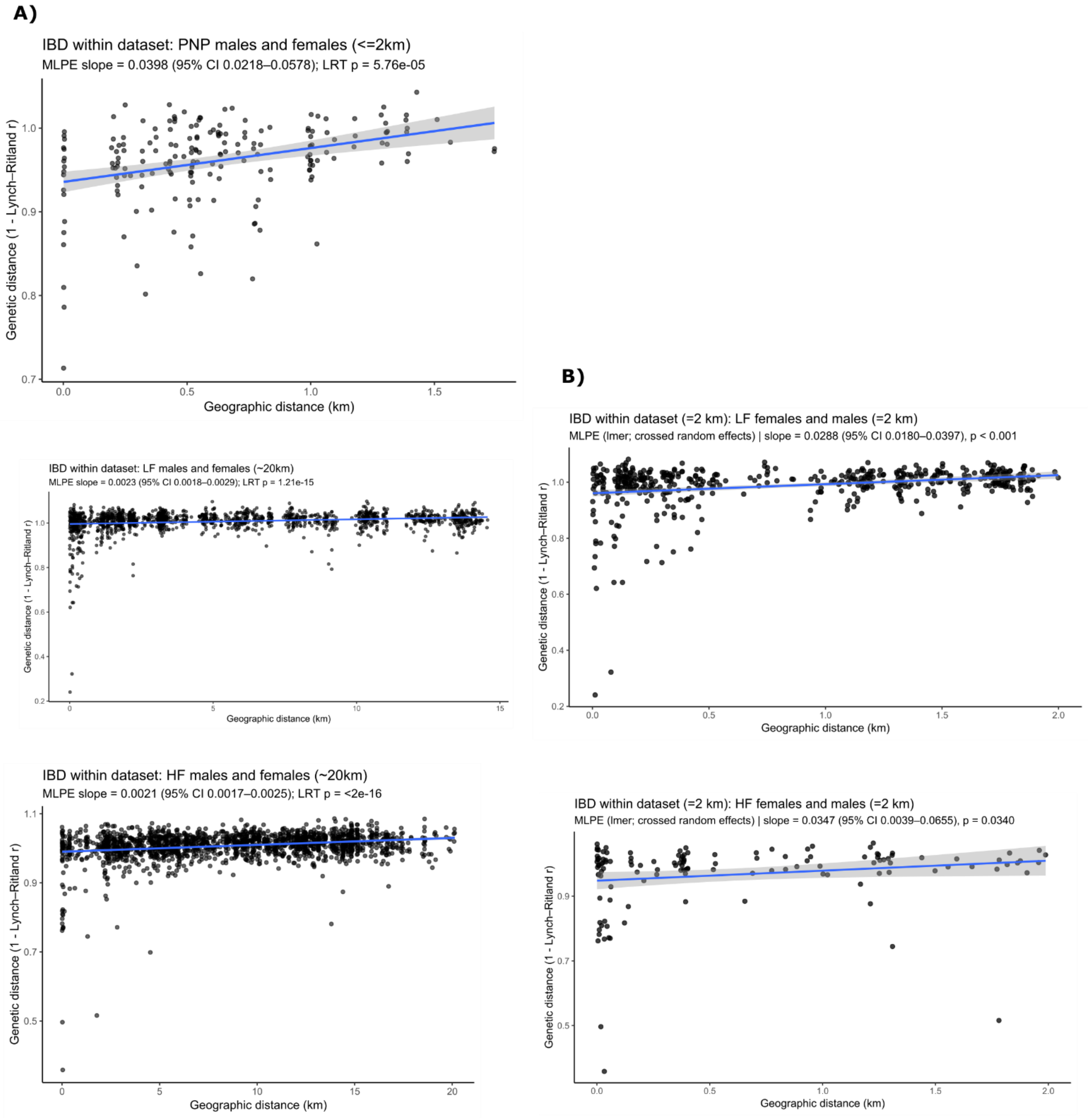

**Table S10. Results of maximum-likelihood population effects (MLPE) analyses for comparing isolation-by-distance (IBD) across regions and by sex.** The sex-, region-, and spatial extent-specific IBD results are presented in main text figure 3B; the results presented here are comparisons of those IBD results to assess if genetic spatial structure differs significantly between regions / sexes; A) MLPE tests for differences in isolation-by-distance slopes among regions; B) estimated regional IBD slopes and pairwise regional comparisons; C) sex-specific IBD slopes and male–female comparisons within regions.

A) MLPE tests for differences in isolation-by-distance slopes among regions. Models were fitted using maximum likelihood.

| Dataset | Comparison | Null model | Alternative model | $\chi^2$ | df | P-value | Interpretation |
|---|---|---|---|---|---|---|---|
| Males | Regional IBD slopes (PNP, LF, HF) | Common slope among regions | Region-specific slopes | 8.3977 | 2 | **0.015** | Regional slopes differ |
| Females | Regional IBD slopes (PNP, LF, HF) | Common slope among regions | Region-specific slopes | 8.9833 | 2 | **0.011** | Regional slopes differ |
| Sexes pooled | Regional IBD slopes (PNP, LF, HF) | Common slope among regions | Region-specific slopes | 20.6660 | 2 | **$3.25 \times 10^{-5}$** | Regional slopes differ |

B) Estimated regional IBD slopes and pairwise regional comparisons. Positive differences indicate a steeper slope in the first region listed. Kenward–Roger inference was used for male and female comparisons; pooled comparisons used asymptotic inference. Pairwise P values and reported contrast CIs were Tukey-adjusted.

<u>Part 1. Estimated IBD slopes by region</u>

| Dataset | Region | Estimated slope | SE | df | 95% CI |
|---|---|---|---|---|---|
| Males | PNP | 0.0353 | 0.0273 | 498 | −0.0184 to 0.0889 |
| Males | LF | 0.0043 | 0.0007 | 417 | 0.0030 to 0.0056 |
| Males | HF | 0.0021 | 0.0005 | 622 | 0.0010 to 0.0031 |
| Females | PNP | 0.0411 | 0.0140 | 1077 | 0.0137 to 0.0685 |
| Females | LF | 0.0015 | 0.0004 | 392 | 0.0006 to 0.0023 |
| Females | HF | 0.0020 | 0.0004 | 1058 | 0.0012 to 0.0028 |
| Sexes pooled | PNP | 0.0405 | 0.0084 | ∞ | 0.0240 to 0.0570 |
| Sexes pooled | LF | 0.0023 | 0.0003 | ∞ | 0.0018 to 0.0028 |
| Sexes pooled | HF | 0.0021 | 0.0002 | ∞ | 0.0017 to 0.0026 |

Part 2. Tukey-adjusted pairwise comparisons of regional IBD slopes

| Dataset | Regional contrast | Difference in slopes | SE | df | Test statistic | 95% CI | Adjusted P-value |
|---|---|---|---|---|---|---|---|
| Males | PNP − LF | 0.0309 | 0.0273 | 498 | t = 1.132 | −0.0333 to 0.0952 | 0.4950 |
| Males | PNP − HF | 0.0332 | 0.0273 | 498 | t = 1.214 | −0.0317 to 0.0974 | 0.4456 |
| Males | LF − HF | 0.0022 | 0.0008 | 537 | t = 2.649 | 0.0003 to 0.0042 | **0.0226** |
| Females | PNP − LF | 0.0397 | 0.0139 | 1172 | t = 2.849 | 0.0070 to 0.07233 | **0.0124** |
| Females | PNP − HF | 0.0391 | 0.0139 | 1172 | t = 2.809 | 0.0064 to 0.07177 | **0.014** |
| Females | LF − HF | -0.0006 | 0.0006 | 1190 | t = −0.978 | -0.0019 to 0.0008 | 0.5909 |
| Sexes pooled | PNP − LF | 0.0382 | 0.0084 | ∞ | z = 4.527 | 0.0184 to 0.0580 | **<0.0001** |
| Sexes pooled | PNP − HF | 0.0384 | 0.0084 | ∞ | z = 4.548 | 0.0186 to 0.0581 | **<0.0001** |
| Sexes pooled | LF − HF | 0.0002 | 0.0003 | ∞ | z = 0.496 | −0.0006 to 0.0010 | 0.8734 |

C) Sex-specific IBD slopes and male–female comparisons within regions. Slopes represent the estimated change in genetic distance per kilometer of geographic separation. A positive M − F value indicates a steeper male slope. Degrees of freedom and confidence intervals were calculated using the Kenward–Roger method.

| **Overall model comparison** | | | | |
|---|---|---|---|---|
| | $\chi^2$ | **df** | **P value** | **Interpretation** |
| Model without geographic-distance interactions versus full geographic distance × region × sex model | 21.02 | 5 | **0.0008** | IBD slopes vary according to region and/or sex |

| **Estimated sex-specific slopes and within-region sex contrasts** | | | | | | | | | | | | |
|---|---|---|---|---|---|---|---|---|---|---|---|---|
| **Region** | **Female slope** | **Female SE** | **Female 95% CI** | **Male slope** | **Male SE** | **Male 95% CI** | **Difference (M − F)** | **SE of difference** | **df** | **95% CI of difference** | **t-ratio** | **P-value** |
| PNP | 0.0408 | 0.0143 | 0.0128-0.0689 | 0.0355 | 0.0261 | −0.0157-0.0867 | -0.0053 | 0.0298 | 1578 | −0.0637-0.0531 | -0.178 | 0.8584 |
| LF | 0.0015 | 0.0004 | 0.0006-0.0024 | 0.0041 | 0.0006 | 0.0029-0.0053 | 0.0026 | 0.0008 | 882 | 0.0011-0.0041 | 3.380 | **0.0008** |
| HF | 0.0021 | 0.0004 | 0.0013-0.0029 | 0.0021 | 0.0005 | 0.0010-0.0031 | -0.00003 | 0.0007 | 1755 | −0.0013-0.0013 | -0.046 | 0.9634 |

To summarize, when the full spatial extents of all datasets were included, male IBD slopes differ across the three regions (global test: p=0.015), and in pairwise comparisons, only the two fragmented regions (high- and low-fragmented) differed significantly in the strength of IBD for males, with IBD being more pronounced in the low-fragmented region. Female IBD slopes differed across the three regions (global test: p=0.011), and in pairwise comparisons, the genetic spatial structure for females in PNP (continuous forest) was significantly stronger than for females in both the high-fragmented and low-fragmented regions. When all individuals (males and females) were pooled together for each region, IBD slopes differed significantly among them (global test: p=3.25e-05), and in pairwise comparisons, the genetic spatial structure in PNP (continuous forest) was significantly stronger than the genetic spatial structure in both the high-fragmented and low-fragmented regions.

S10.C shows results of comparisons of IBD between males and females within each region, at that region's full spatial extent. Including landscape-specific sex differences in IBD significantly improved the model (global test: p=0.0008). The only significant pairwise difference was found between males and females in the low-fragmented region, where males showed significantly higher IBD than females.

We repeated these tests with the datasets for the high and low fragmented regions restricted to the 2km extent, to match the spatial scale of sampling in PNP. Main text figure 3B shows that at this spatial extent, in PNP only females show significant IBD (for males, while the slope is strong, it is non-significant, likely due at least in part to the small sample size of N=8); in the low-fragmented region, both males and females showed significant IBD at the 2km extent; and in the high-fragmented region neither males nor females showed significant IBD at the 2km extent. However, while significant IBD was found at the 2km spatial extent for both sexes in the low-fragmented region only, when comparing spatial genetic structure between regions and sexes at this limited spatial extent, no significant differences were found, either between males and females within the low- or high-fragmented regions, or for males or females between regions (results not shown, available upon request).

**Fig. S8. Spatial autocorrelation plots** to quantify relatedness values that are significantly higher or lower than expected (noted on the plots with black dots) at different distance classes, across levels of fragmentation and sexes. In columns: PNP (continuous forest), whole fragmented landscape combined, low-fragmented region (LF), and high-fragmented region (HF). In rows: all adults and subadults, then females and males separately. Note the different distance classes in PNP and the fragmented landscape, as a result of the latter's larger spatial extent, and the different ranges of relatedness values (y-axes) for the subsets of the data. 95% CI denoted by gray lines in all plots; relatedness values noted with blue lines. The first distance class in all plots denotes within-group relatedness.

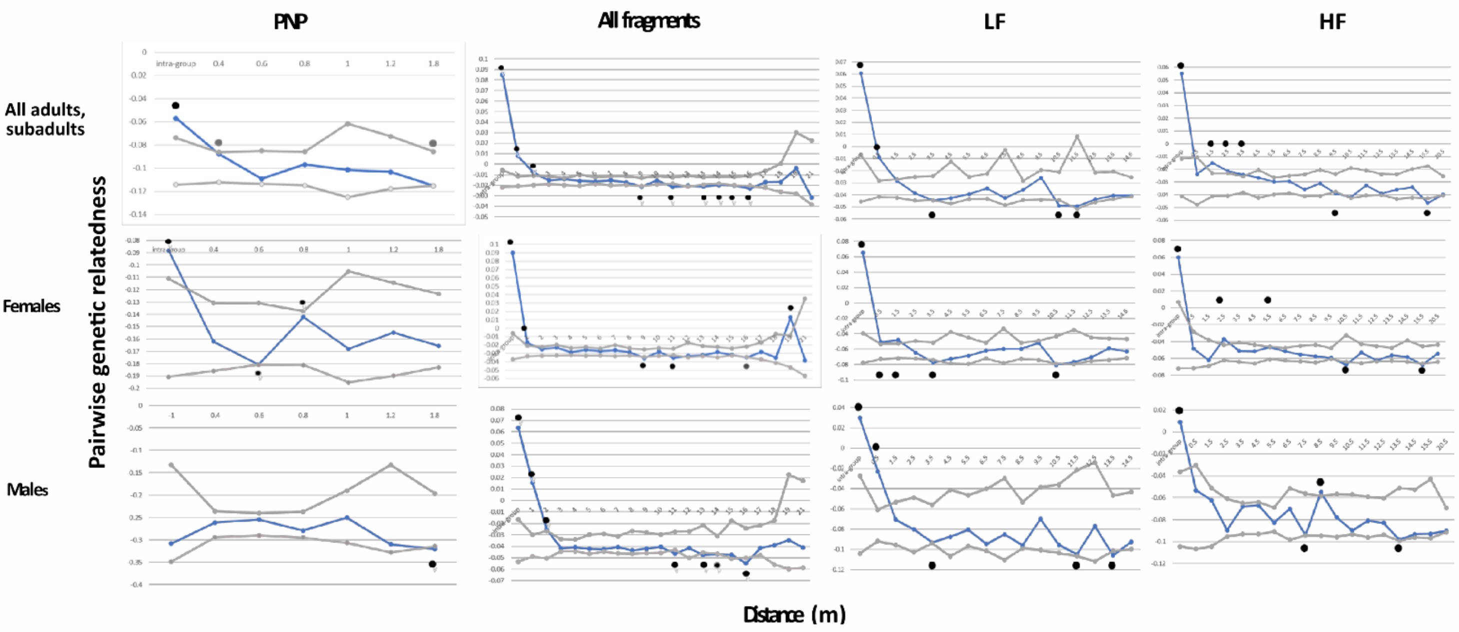

**Table S11. Detailed SPAGeDi (Spatial Pattern Analysis of Genetic Diversity) results.** All datasets include adults and subadults. Distances in km.

| **PNP - all individuals; N = 20; # permutations = 499; # SNPs = 4047** | | | | | | | | | |
|---|---|---|---|---|---|---|---|---|---|
| ALL LOCI | Pairwise RELATIONSHIP coefficients ('r' in Lynch & Ritland, 1999) | | | | | | | Slopes of regression analyses | |
| | intra-group | 2 | 3 | 4 | 5 | 6 | 7 | b-lin (slope | b-log (slope ln(dist)) |
| Object permuted | IaSG | SGLaSG | SGLaSG | SGLaSG | SGLaSG | SGLaSG | SGLaSG | SGLaSG | SGLaSG |
| N valid permut | 499 | 499 | 499 | 499 | 499 | 499 | 499 | 499 | 499 |
| N different permut val | 499 | 499 | 499 | 499 | 482 | 488 | 496 | 499 | 499 |
| Max distance | intra-group | 0.4 | 0.6 | 0.8 | 1 | 1.2 | 1.8 | | |
| Number of pairs | 18 | 37 | 48 | 35 | 21 | 11 | 20 | | |
| % partic | 75 | 85 | 80 | 70 | 65 | 45 | 75 | | |
| CV partic | 0.84 | 0.75 | 0.64 | 0.8 | 0.86 | 1.25 | 1.28 | | |
| Mean distance | 0 | 0.27 | 0.50 | 0.70 | 0.97 | 1.09 | 1.40 | | |
| Obs val | -0.057 | -0.088 | -0.109 | -0.097 | -0.101 | -0.103 | -0.115 | -0.012 | -0.008 |
| Mean permut val | -0.097 | -0.101 | -0.101 | -0.101 | -0.102 | -0.101 | -0.101 | 0.000 | 0.000 |
| SD permut val | 0.010 | 0.006 | 0.007 | 0.007 | 0.015 | 0.011 | 0.007 | 0.006 | 0.005 |
| 95%CI-inf | -0.114 | -0.112 | -0.114 | -0.115 | -0.125 | -0.118 | -0.115 | -0.013 | -0.011 |
| 95%CI-sup | -0.074 | -0.086 | -0.085 | -0.086 | -0.062 | -0.072 | -0.086 | 0.011 | 0.009 |
| 1-sided test, H1: obs<exp | 1.00 | 0.96 | 0.10 | 0.75 | 0.61 | 0.44 | **0.03** | 0.04 | 0.06 |
| 1-sided test, H1: obs>exp | **0.00** | **0.04** | 0.91 | 0.26 | 0.39 | 0.56 | 0.97 | 0.96 | 0.94 |
| 2-sided test, H1: obs<>exp | **0.01** | 0.08 | 0.19 | 0.51 | 0.77 | 0.87 | 0.05 | 0.08 | 0.12 |

| **PNP - females; N = 12; # permutations = 499; # SNPs = 3260** | | | | | | | | | |
|---|---|---|---|---|---|---|---|---|---|
| ALL LOCI | Pairwise RELATIONSHIP coefficients ('r' in Lynch & Ritland, 1999) | | | | | | | Slopes of regression analyses | |
| | intra-group | 2 | 3 | 4 | 5 | 6 | 7 | b-lin (slope | b-log (slope ln(dist)) |
| Object permuted | IaSG | SGLaSG | SGLaSG | SGLaSG | SGLaSG | SGLaSG | SGLaSG | SGLaSG | SGLaSG |
| N valid permut | 499 | 499 | 499 | 499 | 499 | 499 | 499 | 499 | 499 |
| N different permut val | 497 | 377 | 462 | 488 | 192 | 435 | 459 | 497 | 497 |
| Max distance | intra-group | 0.4 | 0.6 | 0.8 | 1 | 1.2 | 1.8 | | |
| Number of pairs | 5 | 10 | 17 | 13 | 8 | 5 | 8 | | |
| % partic | 58.3 | 83.3 | 66.7 | 66.7 | 66.7 | 50 | 75 | | |
| CV partic | 1 | 0.78 | 0.79 | 0.83 | 0.87 | 1.12 | 1.25 | | |
| Mean distance | 0 | 0.26 | 0.49 | 0.70 | 1.00 | 1.05 | 1.39 | | |
| Obs val | -0.088 | -0.162 | -0.181 | -0.142 | -0.168 | -0.155 | -0.166 | 0.012 | 0.007 |
| Mean permut val | -0.159 | -0.164 | -0.162 | -0.162 | -0.164 | -0.162 | -0.162 | 0.001 | 0.000 |
| SD permut val | 0.021 | 0.014 | 0.013 | 0.012 | 0.023 | 0.019 | 0.015 | 0.015 | 0.010 |
| 95%CI-inf | -0.191 | -0.186 | -0.181 | -0.181 | -0.195 | -0.190 | -0.183 | -0.024 | -0.020 |
| 95%CI-sup | -0.111 | -0.131 | -0.131 | -0.137 | -0.105 | -0.114 | -0.123 | 0.032 | 0.019 |
| 1-sided test, H1: obs<exp | 1.00 | 0.60 | **0.03** | 0.96 | 0.52 | 0.69 | 0.52 | 0.76 | 0.75 |
| 1-sided test, H1: obs>exp | **0.00** | 0.40 | 0.97 | **0.05** | 0.49 | 0.31 | 0.48 | 0.24 | 0.25 |
| 2-sided test, H1: obs<>exp | **0.00** | 0.79 | 0.06 | 0.09 | 0.97 | 0.61 | 0.97 | 0.47 | 0.51 |

| **PNP - males; N = 8; # permutations = 499; # SNPs = 2230** | | | | | | | | | |
|---|---|---|---|---|---|---|---|---|---|
| ALL LOCI | Pairwise RELATIONSHIP coefficients ('r' in Lynch & Ritland, 1999) | | | | | | | Slopes of regression analyses | |
| | intra-group | 2 | 3 | 4 | 5 | 6 | 7 | b-lin (slope | b-log (slope ln(dist)) |
| Object permuted | IaSG | SGLaSG | SGLaSG | SGLaSG | SGLaSG | SGLaSG | SGLaSG | SGLaSG | SGLaSG |
| N valid permut | 499 | 499 | 499 | 499 | 499 | 499 | 499 | 499 | 499 |
| N different permut val | 28 | 409 | 465 | 413 | 103 | 21 | 104 | 476 | 476 |
| Max distance | intra-group | 0.4 | 0.6 | 0.8 | 1 | 1.2 | 1.8 | | |
| Number of pairs | 1 | 8 | 8 | 5 | 3 | 1 | 2 | | |
| % partic | 25 | 75 | 87.5 | 62.5 | 62.5 | 25 | 37.5 | | |
| CV partic | 1.85 | 0.76 | 0.53 | 0.93 | 0.94 | 1.85 | 1.51 | | |
| Mean distance | 0.00 | 0.28 | 0.51 | 0.71 | 0.94 | 1.18 | 1.34 | | |
| Obs val | -0.308 | -0.261 | -0.255 | -0.279 | -0.250 | -0.310 | -0.320 | -0.036 | -0.016 |
| Mean permut val | -0.268 | -0.267 | -0.267 | -0.269 | -0.266 | -0.270 | -0.269 | -0.001 | -0.001 |
| SD permut val | 0.047 | 0.015 | 0.013 | 0.014 | 0.027 | 0.037 | 0.027 | 0.025 | 0.016 |
| 95%CI-inf | -0.349 | -0.294 | -0.290 | -0.295 | -0.306 | -0.328 | -0.314 | -0.052 | -0.034 |
| 95%CI-sup | -0.132 | -0.236 | -0.240 | -0.237 | -0.190 | -0.132 | -0.196 | 0.049 | 0.027 |
| 1-sided test, H1: obs<exp | 0.15 | 0.68 | 0.84 | 0.22 | 0.77 | 0.15 | **0.01** | 0.08 | 0.17 |
| 1-sided test, H1: obs>exp | 0.88 | 0.33 | 0.16 | 0.78 | 0.24 | 0.90 | 1.00 | 0.92 | 0.84 |
| 2-sided test, H1: obs<>exp | 0.27 | 0.65 | 0.32 | 0.44 | 0.47 | 0.25 | **0.01** | 0.15 | 0.33 |

**Fragmented landscape - all individuals; N = 118; # permutations = 40; # SNPs = 6824**

| ALL LOCI | Pairwise RELATIONSHIP coefficients ('r' in Lynch & Ritland, 1999) | | | | | | | | | | | | | | | | | | | | | Slopes of regression analyses | |
|---|---|---|---|---|---|---|---|---|---|---|---|---|---|---|---|---|---|---|---|---|---|---|---|
| | intra-group | 2 | 3 | 4 | 5 | 6 | 7 | 8 | 9 | 10 | 11 | 12 | 13 | 14 | 15 | 16 | 17 | 18 | 19 | 20 | 21 | b-lin (slope linear dist) | b-log (slope |
| Object permuted | IaSG | SGLaSG | SGLaSG | SGLaSG | SGLaSG | SGLaSG | SGLaSG | SGLaSG | SGLaSG | SGLaSG | SGLaSG | SGLaSG | SGLaSG | SGLaSG | SGLaSG | SGLaSG | SGLaSG | SGLaSG | SGLaSG | SGLaSG | SGLaSG | SGLaSG | SGLaSG |
| N valid permut | 40 | 40 | 40 | 40 | 40 | 40 | 40 | 40 | 40 | 40 | 40 | 40 | 40 | 40 | 40 | 40 | 40 | 40 | 40 | 40 | 40 | 40 | 40 |
| N different permut val | 40 | 40 | 40 | 40 | 40 | 40 | 40 | 40 | 40 | 40 | 40 | 40 | 40 | 40 | 40 | 40 | 40 | 40 | 40 | 40 | 40 | 40 | 40 |
| Max distance | intra-group | 1 | 2 | 3 | 4 | 5 | 6 | 7 | 8 | 9 | 10 | 11 | 12 | 13 | 14 | 15 | 16 | 17 | 18 | 19 | 21 | | |
| Number of pairs | 94 | 306 | 358 | 596 | 586 | 376 | 519 | 378 | 334 | 383 | 363 | 529 | 219 | 449 | 490 | 429 | 251 | 145 | 54 | 30 | 14 | | |
| % partic | 76.3 | 90.7 | 72.9 | 87.3 | 92.4 | 80.5 | 100 | 91.5 | 88.1 | 91.5 | 96.6 | 83.1 | 72 | 69.5 | 72.9 | 74.6 | 66.1 | 55.9 | 26.3 | 16.1 | 7.6 | | |
| CV partic | 0.93 | 0.9 | 1.06 | 0.81 | 0.68 | 0.96 | 0.76 | 0.96 | 0.99 | 1.11 | 1.07 | 0.86 | 1.1 | 1 | 1.01 | 1.27 | 1.12 | 1.71 | 2.4 | 3.33 | 3.8 | | |
| Mean distance | 0.00 | 0.42 | 1.45 | 2.49 | 3.37 | 4.56 | 5.51 | 6.39 | 7.50 | 8.59 | 9.44 | 10.49 | 11.51 | 12.44 | 13.51 | 14.46 | 15.37 | 16.48 | 17.50 | 18.37 | 19.69 | | |
| Obs val | 0.085 | 0.008 | -0.008 | -0.015 | -0.014 | -0.016 | -0.017 | -0.015 | -0.017 | -0.021 | -0.015 | -0.022 | -0.020 | -0.021 | -0.020 | -0.020 | -0.023 | -0.017 | -0.017 | -0.004 | -0.032 | -0.001 | -0.006 |
| Mean permut val | -0.015 | -0.016 | -0.015 | -0.016 | -0.017 | -0.016 | -0.016 | -0.016 | -0.016 | -0.017 | -0.016 | -0.016 | -0.016 | -0.016 | -0.016 | -0.016 | -0.016 | -0.016 | -0.016 | -0.016 | -0.019 | 0.000 | 0.000 |
| SD permut val | 0.004 | 0.002 | 0.002 | 0.002 | 0.002 | 0.002 | 0.002 | 0.002 | 0.002 | 0.002 | 0.002 | 0.002 | 0.003 | 0.002 | 0.002 | 0.002 | 0.003 | 0.004 | 0.006 | 0.010 | 0.013 | 0.000 | 0.001 |
| 95%CI-inf | -0.022 | -0.021 | -0.020 | -0.019 | -0.020 | -0.020 | -0.019 | -0.020 | -0.020 | -0.021 | -0.019 | -0.019 | -0.021 | -0.020 | -0.019 | -0.020 | -0.020 | -0.023 | -0.026 | -0.028 | -0.039 | 0.000 | -0.001 |
| 95%CI-sup | -0.006 | -0.011 | -0.010 | -0.011 | -0.013 | -0.010 | -0.012 | -0.011 | -0.011 | -0.013 | -0.011 | -0.012 | -0.011 | -0.012 | -0.012 | -0.012 | -0.011 | -0.006 | 0.000 | 0.030 | 0.022 | 0.000 | 0.001 |
| 1-sided test, H1: obs<exp | 1 | 1 | 1 | 0.71 | 0.93 | 0.59 | 0.32 | 0.71 | 0.32 | **0.00** | 0.80 | **0.00** | 0.10 | **0.00** | **0.00** | **0.05** | **0.00** | 0.34 | 0.59 | 0.95 | 0.12 | 0.00 | 0.00 |
| 1-sided test, H1: obs>exp | **0** | **0** | **0** | 0.32 | 0.10 | 0.44 | 0.71 | 0.32 | 0.71 | 1.00 | 0.22 | 1.00 | 0.93 | 1.00 | 1.00 | 0.98 | 1.00 | 0.68 | 0.44 | 0.07 | 0.90 | 1.00 | 1.00 |
| 2-sided test, H1: obs<>exp | **0** | **0** | **0** | 0.61 | 0.17 | 0.85 | 0.61 | 0.61 | 0.61 | **0.00** | 0.41 | **0.00** | 0.17 | **0.00** | **0.00** | 0.07 | **0.00** | 0.66 | 0.85 | 0.12 | 0.22 | 0.00 | 0.00 |

**Fragmented landscape - females; N=68; # permutations = 40; # SNPs = 8454**

| ALL LOCI | Pairwise RELATIONSHIP coefficients ('r' in Lynch & Ritland, 1999) | | | | | | | | | | | | | | | | | | | | | Slopes of regression analyses | |
|---|---|---|---|---|---|---|---|---|---|---|---|---|---|---|---|---|---|---|---|---|---|---|---|
| | intra-group | 2 | 3 | 4 | 5 | 6 | 7 | 8 | 9 | 10 | 11 | 12 | 13 | 14 | 15 | 16 | 17 | 18 | 19 | 20 | 21 | b-lin (slope linear dist) | b-log (slope |
| Object permuted | IaSG | SGLaSG | SGLaSG | SGLaSG | SGLaSG | SGLaSG | SGLaSG | SGLaSG | SGLaSG | SGLaSG | SGLaSG | SGLaSG | SGLaSG | SGLaSG | SGLaSG | SGLaSG | SGLaSG | SGLaSG | SGLaSG | SGLaSG | SGLaSG | SGLaSG | SGLaSG |
| N valid permut | 40 | 40 | 40 | 40 | 40 | 40 | 40 | 40 | 40 | 40 | 40 | 40 | 40 | 40 | 40 | 40 | 40 | 40 | 40 | 40 | 40 | 40 | 40 |
| N different permut val | 40 | 40 | 40 | 40 | 40 | 40 | 40 | 40 | 40 | 40 | 40 | 40 | 40 | 40 | 40 | 40 | 40 | 40 | 40 | 40 | 40 | 40 | 40 |
| Max distance | intra-group | 1 | 2 | 3 | 4 | 5 | 6 | 7 | 8 | 9 | 10 | 11 | 12 | 13 | 14 | 15 | 16 | 17 | 18 | 19 | 21 | | |
| Number of pairs | 24 | 102 | 130 | 173 | 161 | 112 | 158 | 115 | 121 | 133 | 129 | 144 | 75 | 177 | 163 | 193 | 105 | 43 | 11 | 6 | 3 | | |
| % partic | 50 | 85.3 | 69.1 | 82.4 | 82.4 | 73.5 | 97.1 | 85.3 | 85.3 | 88.2 | 98.5 | 79.4 | 70.6 | 73.5 | 63.2 | 73.5 | 63.2 | 48.5 | 19.1 | 10.3 | 5.9 | | |
| CV partic | 1.23 | 0.89 | 1.01 | 0.93 | 0.84 | 1.11 | 0.71 | 1.02 | 1 | 1.11 | 1.07 | 0.99 | 1.11 | 0.9 | 1.05 | 1.15 | 1.04 | 2.01 | 3.31 | 4.37 | 4.69 | | |
| Mean distance | 0.00 | 0.44 | 1.48 | 2.47 | 3.38 | 4.53 | 5.55 | 6.40 | 7.54 | 8.55 | 9.43 | 10.51 | 11.55 | 12.45 | 13.48 | 14.44 | 15.39 | 16.58 | 17.59 | 18.36 | 19.89 | | |
| Obs val | 0.090 | -0.017 | -0.025 | -0.023 | -0.029 | -0.026 | -0.027 | -0.026 | -0.029 | -0.036 | -0.028 | -0.035 | -0.033 | -0.032 | -0.029 | -0.032 | -0.035 | -0.028 | -0.035 | 0.013 | -0.038 | -0.001 | -0.004 |
| Mean permut val | -0.028 | -0.029 | -0.029 | -0.029 | -0.028 | -0.028 | -0.029 | -0.029 | -0.029 | -0.029 | -0.029 | -0.028 | -0.030 | -0.028 | -0.029 | -0.029 | -0.029 | -0.030 | -0.028 | -0.032 | -0.026 | 0.000 | 0.000 |
| SD permut val | 0.007 | 0.003 | 0.003 | 0.003 | 0.002 | 0.003 | 0.002 | 0.003 | 0.002 | 0.002 | 0.002 | 0.002 | 0.004 | 0.003 | 0.003 | 0.002 | 0.004 | 0.005 | 0.008 | 0.009 | 0.019 | 0.000 | 0.001 |
| 95%CI-inf | -0.037 | -0.034 | -0.033 | -0.033 | -0.032 | -0.033 | -0.033 | -0.033 | -0.033 | -0.034 | -0.033 | -0.032 | -0.035 | -0.033 | -0.035 | -0.032 | -0.035 | -0.037 | -0.041 | -0.047 | -0.057 | 0.000 | -0.001 |
| 95%CI-sup | -0.006 | -0.021 | -0.022 | -0.020 | -0.024 | -0.023 | -0.024 | -0.020 | -0.024 | -0.025 | -0.024 | -0.024 | -0.017 | -0.021 | -0.023 | -0.024 | -0.022 | -0.017 | -0.007 | -0.008 | 0.036 | 0.000 | 0.002 |
| 1-sided test, H1: obs<exp | 1 | 1 | 0.88 | 0.95 | 0.46 | 0.88 | 0.83 | 0.90 | 0.51 | **0.00** | 0.71 | **0.00** | 0.17 | 0.12 | 0.56 | 0.10 | **0.05** | 0.63 | 0.20 | 1.00 | 0.29 | 0.00 | 0.00 |
| 1-sided test, H1: obs>exp | **0** | **0** | 0.15 | 0.07 | 0.56 | 0.15 | 0.20 | 0.12 | 0.51 | 1.00 | 0.32 | 1.00 | 0.85 | 0.90 | 0.46 | 0.93 | 0.98 | 0.39 | 0.83 | **0.00** | 0.73 | 1.00 | 1.00 |
| 2-sided test, H1: obs<>exp | **0** | **0** | 0.27 | 0.12 | 0.90 | 0.27 | 0.37 | 0.22 | 1.00 | **0.00** | 0.61 | **0.00** | 0.32 | 0.22 | 0.90 | 0.17 | 0.07 | 0.76 | 0.37 | **0.00** | 0.56 | 0.00 | 0.00 |

**Fragmented landscape - males; N = 50; # permutations = 45; # SNPs = 7516**

| ALL LOCI | Pairwise RELATIONSHIP coefficients ('r' in Lynch & Ritland, 1999) | | | | | | | | | | | | | | | | | | | | | Slopes of regression analyses | |
|---|---|---|---|---|---|---|---|---|---|---|---|---|---|---|---|---|---|---|---|---|---|---|---|
| | intra-group | 2 | 3 | 4 | 5 | 6 | 7 | 8 | 9 | 10 | 11 | 12 | 13 | 14 | 15 | 16 | 17 | 18 | 19 | 20 | 21 | b-lin (slope linear dist) | b-log (slope ln(dist)) |
| Object permuted | IaSG | SGLaSG | SGLaSG | SGLaSG | SGLaSG | SGLaSG | SGLaSG | SGLaSG | SGLaSG | SGLaSG | SGLaSG | SGLaSG | SGLaSG | SGLaSG | SGLaSG | SGLaSG | SGLaSG | SGLaSG | SGLaSG | SGLaSG | SGLaSG | SGLaSG | SGLaSG |
| N valid permut | 45 | 45 | 45 | 45 | 45 | 45 | 45 | 45 | 45 | 45 | 45 | 45 | 45 | 45 | 45 | 45 | 45 | 45 | 45 | 45 | 45 | 45 | 45 |
| N different permut val | 45 | 45 | 45 | 45 | 45 | 45 | 45 | 45 | 45 | 45 | 45 | 45 | 45 | 45 | 45 | 45 | 45 | 45 | 45 | 45 | 45 | 45 | 45 |
| Max distance | intra-group | 1 | 2 | 3 | 4 | 5 | 6 | 7 | 8 | 9 | 10 | 11 | 12 | 13 | 14 | 15 | 16 | 17 | 18 | 19 | 21 | | |
| Number of pairs | 15 | 55 | 49 | 123 | 126 | 72 | 104 | 80 | 63 | 61 | 59 | 108 | 38 | 59 | 92 | 38 | 28 | 29 | 16 | 6 | 4 | | |
| % partic | 42 | 70 | 70 | 88 | 90 | 78 | 82 | 88 | 74 | 74 | 68 | 78 | 66 | 54 | 72 | 46 | 48 | 46 | 34 | 16 | 10 | | |
| CV partic | 1.3 | 0.97 | 1.2 | 0.76 | 0.58 | 0.87 | 0.9 | 0.91 | 1.13 | 1.23 | 1.22 | 0.84 | 1.2 | 1.23 | 0.99 | 1.26 | 1.58 | 1.71 | 1.97 | 2.86 | 3.43 | | |
| Mean distance | 0.00 | 0.39 | 1.38 | 2.49 | 3.38 | 4.60 | 5.45 | 6.37 | 7.45 | 8.66 | 9.50 | 10.46 | 11.45 | 12.44 | 13.54 | 14.63 | 15.37 | 16.33 | 17.52 | 18.43 | 19.50 | | |
| Obs val | 0.063 | 0.015 | -0.025 | -0.042 | -0.041 | -0.042 | -0.042 | -0.041 | -0.044 | -0.042 | -0.040 | -0.046 | -0.041 | -0.047 | -0.047 | -0.047 | -0.055 | -0.041 | -0.039 | -0.035 | -0.041 | -0.001 | -0.011 |
| Mean permut val | -0.041 | -0.040 | -0.042 | -0.040 | -0.040 | -0.039 | -0.039 | -0.041 | -0.040 | -0.039 | -0.039 | -0.039 | -0.041 | -0.039 | -0.040 | -0.041 | -0.041 | -0.038 | -0.041 | -0.038 | -0.040 | 0.000 | 0.000 |
| SD permut val | 0.009 | 0.004 | 0.005 | 0.003 | 0.003 | 0.004 | 0.003 | 0.003 | 0.004 | 0.005 | 0.004 | 0.003 | 0.006 | 0.005 | 0.004 | 0.007 | 0.006 | 0.006 | 0.008 | 0.018 | 0.013 | 0.000 | 0.001 |
| 95%CI-inf | -0.053 | -0.049 | -0.051 | -0.044 | -0.044 | -0.047 | -0.045 | -0.046 | -0.046 | -0.046 | -0.046 | -0.043 | -0.050 | -0.045 | -0.046 | -0.051 | -0.050 | -0.048 | -0.056 | -0.060 | -0.059 | 0.000 | -0.002 |
| 95%CI-sup | -0.017 | -0.030 | -0.027 | -0.034 | -0.034 | -0.030 | -0.029 | -0.031 | -0.027 | -0.028 | -0.030 | -0.027 | -0.027 | -0.022 | -0.031 | -0.018 | -0.024 | -0.022 | -0.017 | 0.022 | 0.017 | 0.000 | 0.003 |
| 1-sided test, H1: obs<exp | 1 | 1 | 1 | 0.26 | 0.50 | 0.24 | 0.15 | 0.57 | 0.20 | 0.35 | 0.46 | **0.00** | 0.57 | **0.00** | **0.04** | 0.13 | **0.00** | 0.30 | 0.67 | 0.65 | 0.52 | 0.00 | 0.00 |
| 1-sided test, H1: obs>exp | **0** | **0** | **0** | 0.76 | 0.52 | 0.78 | 0.87 | 0.46 | 0.83 | 0.67 | 0.57 | 1.00 | 0.46 | 1.00 | 0.98 | 0.89 | 1.00 | 0.72 | 0.35 | 0.37 | 0.50 | 1.00 | 1.00 |
| 2-sided test, H1: obs<>exp | **0** | **0** | **0** | 0.50 | 0.98 | 0.46 | 0.28 | 0.89 | 0.37 | 0.67 | 0.89 | **0.00** | 0.89 | **0.00** | 0.07 | 0.24 | **0.00** | 0.59 | 0.67 | 0.72 | 0.98 | 0.00 | 0.00 |

**LF - all individuals; N = 58; # permutations = 99; # SNPs = 7839**

| ALL LOCI | Pairwise RELATIONSHIP coefficients ('r' in Lynch & Ritland, 1999) | | | | | | | | | | | | | | | | Slopes of regression analyses | |
|---|---|---|---|---|---|---|---|---|---|---|---|---|---|---|---|---|---|---|
| | intra-group | 2 | 3 | 4 | 5 | 6 | 7 | 8 | 9 | 10 | 11 | 12 | 13 | 14 | 15 | 16 | b-lin (slope linear dist) | b-log (slope ln(dist)) |
| Object permuted | IaSG | SGLaSG | SGLaSG | SGLaSG | SGLaSG | SGLaSG | SGLaSG | SGLaSG | SGLaSG | SGLaSG | SGLaSG | SGLaSG | SGLaSG | SGLaSG | SGLaSG | SGLaSG | SGLaSG | SGLaSG |
| N valid permut | 99 | 99 | 99 | 99 | 99 | 99 | 99 | 99 | 99 | 99 | 99 | 99 | 99 | 99 | 99 | 99 | 99 | 99 |
| N different permut val | 99 | 99 | 99 | 99 | 99 | 99 | 99 | 99 | 99 | 99 | 99 | 99 | 99 | 99 | 99 | 99 | 99 | 99 |
| Max distance | intra-group | 0.5 | 1.5 | 2.5 | 3.5 | 4.5 | 5.5 | 6.5 | 7.5 | 8.5 | 9.5 | 10.5 | 11.5 | 12.5 | 13.5 | 14.6 | | |
| Number of pairs | 50 | 167 | 164 | 206 | 196 | 41 | 85 | 110 | 43 | 84 | 68 | 97 | 56 | 70 | 91 | 125 | | |
| % partic | 79.3 | 87.9 | 69 | 67.2 | 77.6 | 58.6 | 67.2 | 62.1 | 48.3 | 58.6 | 69 | 63.8 | 34.5 | 37.9 | 55.2 | 44.8 | | |
| CV partic | 0.9 | 0.67 | 1.05 | 1.01 | 0.82 | 1.45 | 1.42 | 1.43 | 1.56 | 1.37 | 1.4 | 1.19 | 1.68 | 1.53 | 1.16 | 1.25 | | |
| Mean distance | 0.00 | 0.25 | 1.11 | 1.95 | 3.17 | 3.87 | 4.89 | 6.03 | 6.88 | 7.85 | 8.94 | 10.28 | 10.78 | 12.13 | 13.01 | 13.81 | | |
| Obs val | 0.061 | -0.009 | -0.028 | -0.039 | -0.045 | -0.043 | -0.039 | -0.035 | -0.043 | -0.036 | -0.026 | -0.049 | -0.049 | -0.044 | -0.041 | -0.041 | -0.001 | -0.006 |
| Mean permut val | -0.033 | -0.036 | -0.036 | -0.036 | -0.035 | -0.034 | -0.036 | -0.036 | -0.036 | -0.036 | -0.034 | -0.035 | -0.032 | -0.035 | -0.035 | -0.035 | 0.000 | 0.000 |
| SD permut val | 0.008 | 0.003 | 0.004 | 0.005 | 0.005 | 0.009 | 0.005 | 0.005 | 0.009 | 0.004 | 0.006 | 0.005 | 0.013 | 0.007 | 0.005 | 0.004 | 0.000 | 0.001 |
| 95%CI-inf | -0.045 | -0.042 | -0.042 | -0.045 | -0.044 | -0.047 | -0.044 | -0.043 | -0.048 | -0.044 | -0.044 | -0.044 | -0.052 | -0.046 | -0.043 | -0.041 | -0.001 | -0.002 |
| 95%CI-sup | -0.007 | -0.028 | -0.027 | -0.025 | -0.024 | -0.013 | -0.025 | -0.022 | -0.003 | -0.028 | -0.019 | -0.021 | 0.008 | -0.022 | -0.021 | -0.026 | 0.001 | 0.002 |
| 1-sided test, H1: obs<exp | 1 | 1 | 0.97 | 0.31 | **0.03** | 0.16 | 0.26 | 0.63 | 0.24 | 0.53 | 0.89 | **0** | **0.04** | 0.11 | 0.1 | 0.06 | 0 | 0 |
| 1-sided test, H1: obs>exp | **0** | **0** | **0.04** | 0.7 | 0.98 | 0.85 | 0.75 | 0.38 | 0.77 | 0.48 | 0.12 | 1 | 0.97 | 0.9 | 0.91 | 0.95 | 1 | 1 |
| 2-sided test, H1: obs<>exp | **0** | **0** | 0.07 | 0.61 | 0.05 | 0.31 | 0.51 | 0.75 | 0.47 | 0.95 | 0.23 | **0** | 0.07 | 0.21 | 0.19 | 0.11 | 0 | 0 |

**LF - females; N = 33; # permutations = 99; # SNPs = 6773**

| ALL LOCI | Pairwise RELATIONSHIP coefficients ('r' in Lynch & Ritland, 1999) | | | | | | | | | | | | | | | | Slopes of regression analyses | |
|---|---|---|---|---|---|---|---|---|---|---|---|---|---|---|---|---|---|---|
| | intra-group | 2 | 3 | 4 | 5 | 6 | 7 | 8 | 9 | 10 | 11 | 12 | 13 | 14 | 15 | 16 | b-lin (slope linear dist) | b-log (slope ln(dist)) |
| Object permuted | IaSG | SGLaSG | SGLaSG | SGLaSG | SGLaSG | SGLaSG | SGLaSG | SGLaSG | SGLaSG | SGLaSG | SGLaSG | SGLaSG | SGLaSG | SGLaSG | SGLaSG | SGLaSG | SGLaSG | SGLaSG |
| N valid permut | 99 | 99 | 99 | 99 | 99 | 99 | 99 | 99 | 99 | 99 | 99 | 99 | 99 | 99 | 99 | 99 | 99 | 99 |
| N different permut val | 99 | 99 | 99 | 99 | 99 | 99 | 99 | 99 | 99 | 99 | 99 | 99 | 99 | 99 | 99 | 99 | 99 | 99 |
| Max distance | intra-group | 0.5 | 1.5 | 2.5 | 3.5 | 4.5 | 5.5 | 6.5 | 7.5 | 8.5 | 9.5 | 10.5 | 11.5 | 12.5 | 13.5 | 14.6 | | |
| Number of pairs | 11 | 44 | 68 | 75 | 42 | 12 | 23 | 41 | 11 | 31 | 21 | 22 | 15 | 34 | 36 | 42 | | |
| % partic | 57.6 | 81.8 | 75.8 | 66.7 | 60.6 | 45.5 | 63.6 | 66.7 | 33.3 | 60.6 | 51.5 | 36.4 | 24.2 | 45.5 | 54.5 | 45.5 | | |
| CV partic | 0.97 | 0.65 | 0.87 | 0.93 | 1.1 | 1.59 | 1.47 | 1.31 | 1.62 | 1.28 | 1.57 | 1.5 | 1.87 | 1.32 | 1.08 | 1.22 | | |
| Mean distance | 0 | 0.26 | 1.07 | 1.93 | 3.21 | 3.94 | 4.99 | 5.99 | 6.77 | 7.90 | 8.99 | 10.29 | 10.86 | 12.11 | 12.98 | 13.89 | | |
| Obs val | 0.065 | -0.051 | -0.048 | -0.065 | -0.077 | -0.073 | -0.069 | -0.062 | -0.059 | -0.060 | -0.052 | -0.081 | -0.076 | -0.071 | -0.059 | -0.063 | -0.001 | -0.004 |
| Mean permut val | -0.062 | -0.062 | -0.062 | -0.062 | -0.063 | -0.060 | -0.062 | -0.063 | -0.062 | -0.063 | -0.063 | -0.062 | -0.060 | -0.062 | -0.061 | -0.062 | 0.000 | 0.000 |
| SD permut val | 0.009 | 0.005 | 0.005 | 0.006 | 0.006 | 0.010 | 0.008 | 0.006 | 0.010 | 0.005 | 0.006 | 0.009 | 0.012 | 0.007 | 0.007 | 0.005 | 0.000 | 0.002 |
| 95%CI-inf | -0.078 | -0.073 | -0.072 | -0.072 | -0.074 | -0.079 | -0.079 | -0.072 | -0.078 | -0.073 | -0.074 | -0.079 | -0.079 | -0.075 | -0.074 | -0.071 | -0.001 | -0.003 |
| 95%CI-sup | -0.039 | -0.053 | -0.053 | -0.050 | -0.052 | -0.038 | -0.045 | -0.052 | -0.033 | -0.052 | -0.049 | -0.043 | -0.035 | -0.045 | -0.046 | -0.047 | 0.001 | 0.003 |
| 1-sided test, H1: obs<exp | 1 | 1 | 1 | 0.28 | **0** | 0.1 | 0.16 | 0.53 | 0.66 | 0.75 | 0.96 | **0.02** | 0.08 | 0.13 | 0.62 | 0.45 | 0.06 | 0.02 |
| 1-sided test, H1: obs>exp | **0** | **0** | **0** | 0.73 | 1 | 0.91 | 0.85 | 0.48 | 0.35 | 0.26 | 0.05 | 0.99 | 0.93 | 0.88 | 0.39 | 0.56 | 0.95 | 0.99 |
| 2-sided test, H1: obs<>exp | **0** | **0** | **0** | 0.55 | **0** | 0.19 | 0.31 | 0.95 | 0.69 | 0.51 | 0.09 | **0.03** | 0.15 | 0.25 | 0.77 | 0.89 | 0.11 | 0.03 |

**LF - males; N = 25; # permutations = 199; # SNPs = 5772**

| ALL LOCI | Pairwise RELATIONSHIP coefficients ('r' in Lynch & Ritland, 1999) | | | | | | | | | | | | | | | | Slopes of regression analyses | |
|---|---|---|---|---|---|---|---|---|---|---|---|---|---|---|---|---|---|---|
| | intra-group | 2 | 3 | 4 | 5 | 6 | 7 | 8 | 9 | 10 | 11 | 12 | 13 | 14 | 15 | 16 | b-lin (slope linear dist) | b-log (slope ln(dist)) |
| Object permuted | IaSG | SGLaSG | SGLaSG | SGLaSG | SGLaSG | SGLaSG | SGLaSG | SGLaSG | SGLaSG | SGLaSG | SGLaSG | SGLaSG | SGLaSG | SGLaSG | SGLaSG | SGLaSG | SGLaSG | SGLaSG |
| N valid permut | 199 | 199 | 199 | 199 | 199 | 199 | 199 | 199 | 199 | 199 | 199 | 199 | 199 | 199 | 199 | 199 | 199 | 199 |
| N different permut val | 199 | 199 | 199 | 199 | 199 | 199 | 199 | 199 | 199 | 199 | 199 | 199 | 198 | 199 | 199 | 199 | 199 | 199 |
| Max distance | intra-group | 0.5 | 1.5 | 2.5 | 3.5 | 4.5 | 5.5 | 6.5 | 7.5 | 8.5 | 9.5 | 10.5 | 11.5 | 12.5 | 13.5 | 14.5 | | |
| Number of pairs | 9 | 38 | 17 | 28 | 50 | 9 | 21 | 12 | 14 | 14 | 10 | 20 | 18 | 6 | 12 | 22 | | |
| % partic | 48 | 88 | 40 | 56 | 76 | 36 | 72 | 32 | 44 | 36 | 40 | 64 | 36 | 28 | 48 | 44 | | |
| CV partic | 1.17 | 0.65 | 1.53 | 1.23 | 0.7 | 1.53 | 1.31 | 1.82 | 1.53 | 1.77 | 1.49 | 1.3 | 1.49 | 2.09 | 1.49 | 1.38 | | |
| Mean distance | 0 | 0.26 | 1.21 | 1.99 | 3.11 | 3.66 | 4.86 | 6.17 | 6.96 | 7.73 | 8.87 | 10.19 | 10.62 | 12.26 | 13.04 | 13.74 | | |
| Obs val | 0.030 | -0.023 | -0.071 | -0.080 | -0.093 | -0.088 | -0.081 | -0.095 | -0.085 | -0.096 | -0.070 | -0.096 | -0.105 | -0.077 | -0.106 | -0.092 | -0.003 | -0.016 |
| Mean permut val | -0.077 | -0.079 | -0.079 | -0.081 | -0.080 | -0.080 | -0.079 | -0.079 | -0.080 | -0.079 | -0.078 | -0.078 | -0.081 | -0.078 | -0.079 | -0.079 | 0.000 | 0.000 |
| SD permut val | 0.018 | 0.008 | 0.010 | 0.012 | 0.009 | 0.016 | 0.012 | 0.014 | 0.019 | 0.012 | 0.015 | 0.015 | 0.018 | 0.021 | 0.013 | 0.012 | 0.001 | 0.002 |
| 95%CI-inf | -0.104 | -0.092 | -0.096 | -0.103 | -0.094 | -0.107 | -0.097 | -0.101 | -0.110 | -0.099 | -0.101 | -0.104 | -0.107 | -0.112 | -0.102 | -0.100 | -0.002 | -0.006 |
| 95%CI-sup | -0.027 | -0.061 | -0.053 | -0.049 | -0.056 | -0.041 | -0.046 | -0.040 | -0.030 | -0.053 | -0.038 | -0.036 | -0.021 | -0.015 | -0.047 | -0.043 | 0.001 | 0.003 |
| 1-sided test, H1: obs<exp | 1.00 | 1.00 | 0.83 | 0.60 | **0.04** | 0.33 | 0.50 | 0.09 | 0.46 | 0.05 | 0.75 | 0.08 | **0.05** | 0.63 | **0.01** | 0.09 | 0.00 | 0.00 |
| 1-sided test, H1: obs>exp | **0.00** | **0.00** | 0.18 | 0.41 | 0.97 | 0.68 | 0.51 | 0.92 | 0.55 | 0.96 | 0.26 | 0.93 | 0.96 | 0.38 | 1.00 | 0.92 | 1.00 | 1.00 |
| 2-sided test, H1: obs<>exp | **0.00** | **0.00** | 0.35 | 0.82 | 0.08 | 0.65 | 0.99 | 0.18 | 0.92 | 0.10 | 0.51 | 0.16 | 0.09 | 0.75 | **0.02** | 0.17 | 0.00 | 0.00 |

**HF - all individuals; N=60; # permutations = 79; # SNPs = 8166**

| ALL LOCI | Pairwise RELATIONSHIP coefficients ('r' in Lynch & Ritland, 1999) | | | | | | | | | | | | | | | | | | Slopes of regression analyses | |
|---|---|---|---|---|---|---|---|---|---|---|---|---|---|---|---|---|---|---|---|---|
| | intra-group | 2 | 3 | 4 | 5 | 6 | 7 | 8 | 9 | 10 | 11 | 12 | 13 | 14 | 15 | 16 | 17 | 18 | b-lin (slope linear dist) | b-log (slope ln(dist)) |
| Object permuted | IaSG | SGLaSG | SGLaSG | SGLaSG | SGLaSG | SGLaSG | SGLaSG | SGLaSG | SGLaSG | SGLaSG | SGLaSG | SGLaSG | SGLaSG | SGLaSG | SGLaSG | SGLaSG | SGLaSG | SGLaSG | SGLaSG | SGLaSG |
| N valid permut | 79 | 79 | 79 | 79 | 79 | 79 | 79 | 79 | 79 | 79 | 79 | 79 | 79 | 79 | 79 | 79 | 79 | 79 | 79 | 79 |
| N different permut val | 79 | 79 | 79 | 79 | 79 | 79 | 79 | 79 | 79 | 79 | 79 | 79 | 79 | 79 | 79 | 79 | 79 | 79 | 79 | 79 |
| Max distance | intra-group | 0.5 | 1.5 | 2.5 | 3.5 | 4.5 | 5.5 | 6.5 | 7.5 | 8.5 | 9.5 | 10.5 | 11.5 | 12.5 | 13.5 | 14.5 | 15.5 | 20.5 | | |
| Number of pairs | 44 | 31 | 39 | 111 | 126 | 77 | 155 | 118 | 85 | 66 | 121 | 79 | 93 | 149 | 112 | 111 | 113 | 140 | | |
| % partic | 73.3 | 48.3 | 46.7 | 80 | 83.3 | 78.3 | 80 | 93.3 | 76.7 | 58.3 | 86.7 | 71.7 | 83.3 | 95 | 65 | 68.3 | 55 | 68.3 | | |
| CV partic | 0.95 | 1.46 | 1.46 | 0.95 | 0.87 | 0.84 | 0.85 | 0.91 | 1.1 | 1.28 | 0.77 | 1.06 | 0.92 | 0.67 | 1.18 | 1 | 1.08 | 1.28 | | |
| Mean distance | 0.00 | 0.33 | 1.07 | 2.19 | 2.99 | 3.98 | 4.99 | 5.98 | 7.03 | 8.05 | 9.11 | 9.96 | 10.97 | 12.14 | 13.06 | 14.15 | 14.85 | 17.15 | | |
| Obs val | 0.055 | -0.024 | -0.015 | -0.021 | -0.024 | -0.027 | -0.030 | -0.029 | -0.036 | -0.031 | -0.039 | -0.042 | -0.033 | -0.039 | -0.036 | -0.034 | -0.046 | -0.040 | -0.001 | -0.008 |
| Mean permut val | -0.031 | -0.032 | -0.033 | -0.034 | -0.033 | -0.033 | -0.033 | -0.033 | -0.033 | -0.034 | -0.032 | -0.033 | -0.032 | -0.033 | -0.034 | -0.033 | -0.033 | -0.033 | 0.000 | 0.000 |
| SD permut val | 0.006 | 0.008 | 0.005 | 0.004 | 0.003 | 0.004 | 0.003 | 0.003 | 0.004 | 0.005 | 0.003 | 0.006 | 0.004 | 0.004 | 0.004 | 0.004 | 0.005 | 0.003 | 0.000 | 0.001 |
| 95%CI-inf | -0.041 | -0.048 | -0.041 | -0.041 | -0.038 | -0.042 | -0.040 | -0.039 | -0.041 | -0.041 | -0.038 | -0.043 | -0.041 | -0.040 | -0.043 | -0.042 | -0.043 | -0.041 | -0.001 | -0.004 |
| 95%CI-sup | -0.012 | -0.011 | -0.023 | -0.023 | -0.026 | -0.021 | -0.026 | -0.025 | -0.024 | -0.021 | -0.024 | -0.019 | -0.021 | -0.024 | -0.024 | -0.020 | -0.018 | -0.025 | 0.000 | 0.003 |
| 1-sided test, H1: obs<exp | 1.00 | 0.85 | 1.00 | 1.00 | 1.00 | 0.96 | 0.86 | 0.85 | 0.25 | 0.70 | **0.00** | 0.05 | 0.51 | 0.08 | 0.35 | 0.45 | **0.00** | 0.05 | 0.00 | 0.00 |
| 1-sided test, H1: obs>exp | **0.00** | 0.16 | **0.00** | **0.00** | **0.00** | 0.05 | 0.15 | 0.16 | 0.76 | 0.31 | 1.00 | 0.96 | 0.50 | 0.94 | 0.66 | 0.56 | 1.00 | 0.96 | 1.00 | 1.00 |
| 2-sided test, H1: obs<>exp | **0.00** | 0.31 | **0.00** | **0.00** | **0.00** | 0.09 | 0.29 | 0.31 | 0.49 | 0.61 | **0.00** | 0.09 | 0.99 | 0.14 | 0.69 | 0.89 | **0.00** | 0.09 | 0.00 | 0.00 |

**HF - females; N = 35; # permutations = 99; # SNPs = 7227**

| ALL LOCI | Pairwise RELATIONSHIP coefficients ('r' in Lynch & Ritland, 1999) | | | | | | | | | | | | | | | | | | Slopes of regression analyses | |
|---|---|---|---|---|---|---|---|---|---|---|---|---|---|---|---|---|---|---|---|---|
| | intra-group | 2 | 3 | 4 | 5 | 6 | 7 | 8 | 9 | 10 | 11 | 12 | 13 | 14 | 15 | 16 | 17 | 18 | b-lin (slope linear dist) | b-log (slope ln(dist)) |
| Object permuted | IaSG | SGLaSG | SGLaSG | SGLaSG | SGLaSG | SGLaSG | SGLaSG | SGLaSG | SGLaSG | SGLaSG | SGLaSG | SGLaSG | SGLaSG | SGLaSG | SGLaSG | SGLaSG | SGLaSG | SGLaSG | SGLaSG | SGLaSG |
| N valid permut | 99 | 99 | 99 | 99 | 99 | 99 | 99 | 99 | 99 | 99 | 99 | 99 | 99 | 99 | 99 | 99 | 99 | 99 | 99 | 99 |
| N different permut val | 99 | 99 | 99 | 99 | 99 | 99 | 99 | 99 | 99 | 99 | 99 | 99 | 99 | 99 | 99 | 99 | 99 | 99 | 99 | 99 |
| Max distance | intra-group | 0.5 | 1.5 | 2.5 | 3.5 | 4.5 | 5.5 | 6.5 | 7.5 | 8.5 | 9.5 | 10.5 | 11.5 | 12.5 | 13.5 | 14.5 | 15.5 | 20.5 | | |
| Number of pairs | 13 | 14 | 9 | 36 | 27 | 24 | 42 | 50 | 32 | 26 | 51 | 20 | 27 | 55 | 58 | 38 | 37 | 36 | | |
| % partic | 42.9 | 48.6 | 28.6 | 62.9 | 80 | 71.4 | 82.9 | 88.6 | 68.6 | 62.9 | 88.6 | 57.1 | 74.3 | 91.4 | 68.6 | 65.7 | 45.7 | 71.4 | | |
| CV partic | 1.4 | 1.31 | 1.85 | 1.17 | 1.16 | 1 | 0.87 | 0.94 | 1.23 | 1.15 | 0.71 | 1.33 | 1.1 | 0.8 | 1.06 | 1.02 | 1.29 | 1.52 | | |
| Mean distance | 0.00 | 0.29 | 1.09 | 2.13 | 3.00 | 3.93 | 5.11 | 5.99 | 7.08 | 8.02 | 9.09 | 9.99 | 10.96 | 12.21 | 13.07 | 14.25 | 14.90 | 16.96 | | |
| Obs val | 0.060 | -0.049 | -0.062 | -0.037 | -0.051 | -0.052 | -0.047 | -0.052 | -0.056 | -0.058 | -0.059 | -0.068 | -0.053 | -0.063 | -0.056 | -0.059 | -0.068 | -0.054 | -0.001 | -0.005 |
| Mean permut val | -0.053 | -0.055 | -0.055 | -0.056 | -0.054 | -0.055 | -0.055 | -0.055 | -0.055 | -0.056 | -0.056 | -0.053 | -0.056 | -0.055 | -0.056 | -0.055 | -0.055 | -0.055 | 0.000 | 0.000 |
| SD permut val | 0.014 | 0.010 | 0.008 | 0.004 | 0.006 | 0.005 | 0.004 | 0.004 | 0.004 | 0.005 | 0.003 | 0.007 | 0.005 | 0.005 | 0.004 | 0.006 | 0.006 | 0.005 | 0.000 | 0.002 |
| 95%CI-inf | -0.072 | -0.072 | -0.069 | -0.063 | -0.064 | -0.066 | -0.061 | -0.063 | -0.064 | -0.065 | -0.061 | -0.064 | -0.066 | -0.064 | -0.063 | -0.064 | -0.067 | -0.064 | -0.001 | -0.004 |
| 95%CI-sup | 0.007 | -0.029 | -0.038 | -0.044 | -0.041 | -0.044 | -0.046 | -0.048 | -0.045 | -0.044 | -0.048 | -0.033 | -0.043 | -0.046 | -0.048 | -0.039 | -0.046 | -0.044 | 0.000 | 0.003 |
| 1-sided test, H1: obs<exp | 1 | 0.81 | 0.21 | 1 | 0.67 | 0.83 | 0.98 | 0.86 | 0.51 | 0.35 | 0.15 | **0.02** | 0.69 | 0.07 | 0.44 | 0.28 | **0.02** | 0.57 | 0 | 0 |
| 1-sided test, H1: obs>exp | **0** | 0.2 | 0.8 | **0** | 0.34 | 0.18 | **0.03** | 0.15 | 0.5 | 0.66 | 0.86 | 0.99 | 0.32 | 0.94 | 0.57 | 0.73 | 0.99 | 0.44 | 1 | 1 |
| 2-sided test, H1: obs<>exp | **0** | 0.39 | 0.41 | **0** | 0.67 | 0.35 | 0.05 | 0.29 | 0.99 | 0.69 | 0.29 | **0.03** | 0.63 | 0.13 | 0.87 | 0.55 | **0.03** | 0.87 | 0 | 0 |

**HF - males; N=25; # permutations = 149; # SNPs = 6014**

| ALL LOCI | Pairwise RELATIONSHIP coefficients ('r' in Lynch & Ritland, 1999) | | | | | | | | | | | | | | | | | | Slopes of regression analyses | |
|---|---|---|---|---|---|---|---|---|---|---|---|---|---|---|---|---|---|---|---|---|
| | intra-group | 2 | 3 | 4 | 5 | 6 | 7 | 8 | 9 | 10 | 11 | 12 | 13 | 14 | 15 | 16 | 17 | 18 | b-lin (slope linear dist) | b-log (slope ln(dist)) |
| Object permuted | IaSG | SGLaSG | SGLaSG | SGLaSG | SGLaSG | SGLaSG | SGLaSG | SGLaSG | SGLaSG | SGLaSG | SGLaSG | SGLaSG | SGLaSG | SGLaSG | SGLaSG | SGLaSG | SGLaSG | SGLaSG | SGLaSG | SGLaSG |
| N valid permut | 149 | 149 | 149 | 149 | 149 | 149 | 149 | 149 | 149 | 149 | 149 | 149 | 149 | 149 | 149 | 149 | 149 | 149 | 149 | 149 |
| N different permut val | 149 | 149 | 149 | 149 | 149 | 149 | 149 | 149 | 149 | 149 | 149 | 149 | 149 | 149 | 149 | 149 | 149 | 149 | 149 | 149 |
| Max distance | intra-group | 0.5 | 1.5 | 2.5 | 3.5 | 4.5 | 5.5 | 6.5 | 7.5 | 8.5 | 9.5 | 10.5 | 11.5 | 12.5 | 13.5 | 14.5 | 15.5 | 20.5 | | |
| Number of pairs | 6 | 5 | 8 | 21 | 25 | 22 | 34 | 10 | 12 | 10 | 15 | 17 | 16 | 24 | 8 | 12 | 24 | 31 | | |
| % partic | 36 | 28 | 44 | 80 | 80 | 76 | 60 | 56 | 56 | 36 | 56 | 68 | 76 | 84 | 48 | 52 | 60 | 64 | | |
| CV partic | 1.49 | 1.91 | 1.49 | 0.87 | 0.8 | 0.9 | 1 | 1.25 | 1.14 | 1.57 | 1.32 | 1.08 | 0.97 | 0.75 | 1.27 | 1.1 | 0.9 | 1.12 | | |
| Mean distance | 0.00 | 0.34 | 1.00 | 2.26 | 3.01 | 4.02 | 4.91 | 6.01 | 7.03 | 8.13 | 9.04 | 10.01 | 11.07 | 12.08 | 12.85 | 13.99 | 14.76 | 17.27 | | |
| Obs val | 0.009 | -0.053 | -0.062 | -0.090 | -0.068 | -0.067 | -0.083 | -0.070 | -0.094 | -0.055 | -0.078 | -0.090 | -0.081 | -0.083 | -0.098 | -0.093 | -0.093 | -0.090 | -0.001 | -0.008 |
| Mean permut val | -0.080 | -0.080 | -0.082 | -0.082 | -0.082 | -0.081 | -0.081 | -0.082 | -0.081 | -0.081 | -0.081 | -0.080 | -0.082 | -0.081 | -0.080 | -0.081 | -0.080 | -0.081 | 0.000 | 0.000 |
| SD permut val | 0.015 | 0.017 | 0.012 | 0.008 | 0.007 | 0.007 | 0.006 | 0.010 | 0.009 | 0.009 | 0.009 | 0.009 | 0.009 | 0.008 | 0.012 | 0.010 | 0.011 | 0.006 | 0.000 | 0.003 |
| 95%CI-inf | -0.104 | -0.107 | -0.104 | -0.095 | -0.093 | -0.093 | -0.091 | -0.098 | -0.094 | -0.095 | -0.096 | -0.093 | -0.096 | -0.094 | -0.099 | -0.096 | -0.097 | -0.091 | -0.001 | -0.005 |
| 95%CI-sup | -0.036 | -0.030 | -0.051 | -0.061 | -0.065 | -0.064 | -0.069 | -0.052 | -0.056 | -0.058 | -0.057 | -0.057 | -0.059 | -0.060 | -0.051 | -0.053 | -0.043 | -0.069 | 0.001 | 0.005 |
| 1-sided test, H1: obs<exp | 1.000 | 0.947 | 0.940 | 0.187 | 0.953 | 0.953 | 0.427 | 0.887 | **0.040** | 0.993 | 0.673 | 0.087 | 0.600 | 0.460 | **0.040** | 0.080 | 0.073 | 0.053 | 0 | 0 |
| 1-sided test, H1: obs>exp | **0.000** | 0.060 | 0.067 | 0.820 | 0.053 | 0.053 | 0.580 | 0.120 | 0.967 | 0.013 | 0.333 | 0.920 | 0.407 | 0.547 | 0.967 | 0.927 | 0.933 | 0.953 | 1 | 1 |
| 2-sided test, H1: obs<>exp | **0.000** | 0.113 | 0.127 | 0.367 | 0.100 | 0.100 | 0.847 | 0.233 | 0.073 | **0.020** | 0.660 | 0.167 | 0.807 | 0.913 | 0.073 | 0.153 | 0.140 | 0.100 | 0 | 0 |

### Microbiome pipeline and analyses

**Fig. S9. Rarefaction plots.** A. Richness. B. Shannon diversity. C. Faith's phylogenetic diversity.

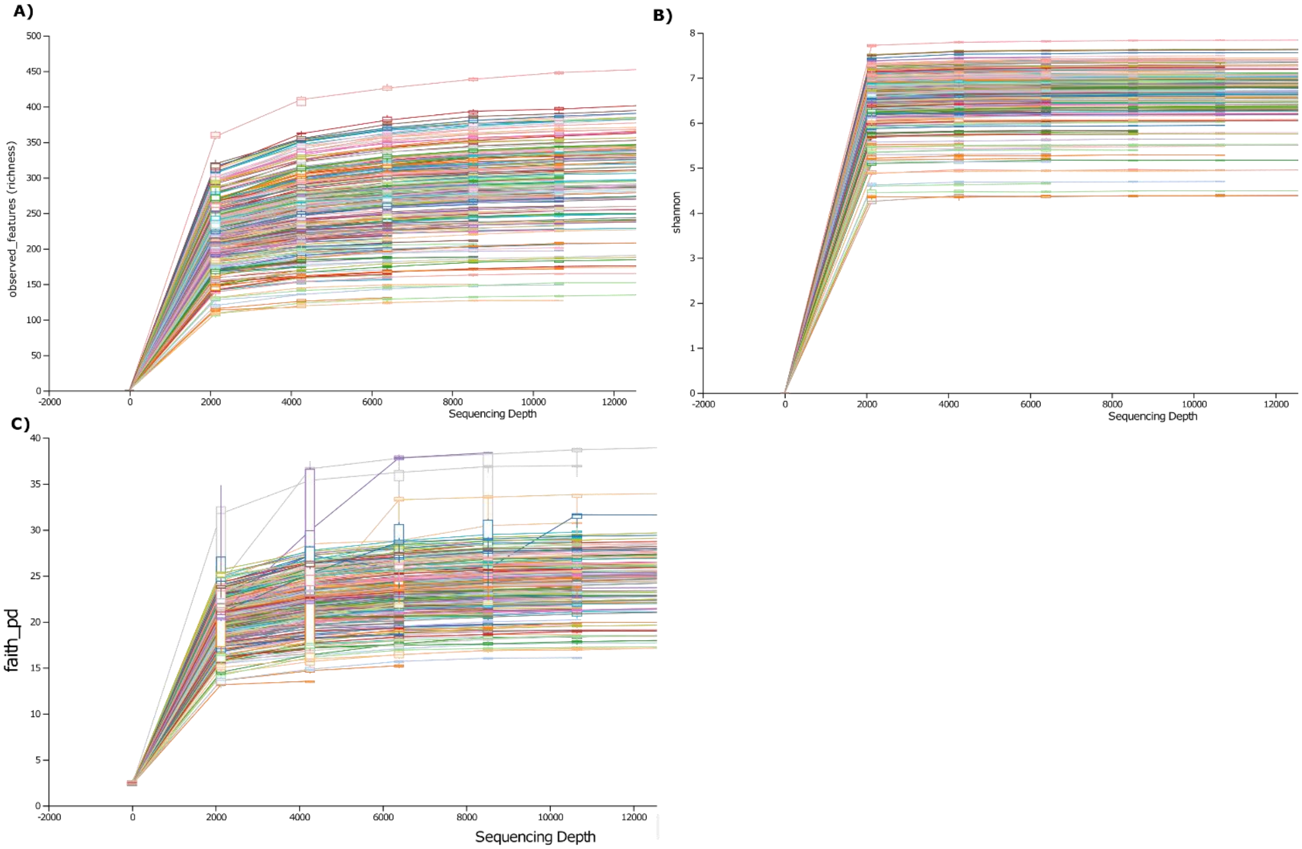

<u>**Microbiome spatial patterns**</u>

**Fig. S10. By-individual PCoA plots showing sorting of microbiomes by different variables (rows) with four dissimilarity indices (columns).** A) sequencing batch (batch 1 – red; batch 2 – blue); B) fragment; C) Age (adults – red; subadults – green; juveniles – orange; infants – blue); D) sex (male – red; female – blue).

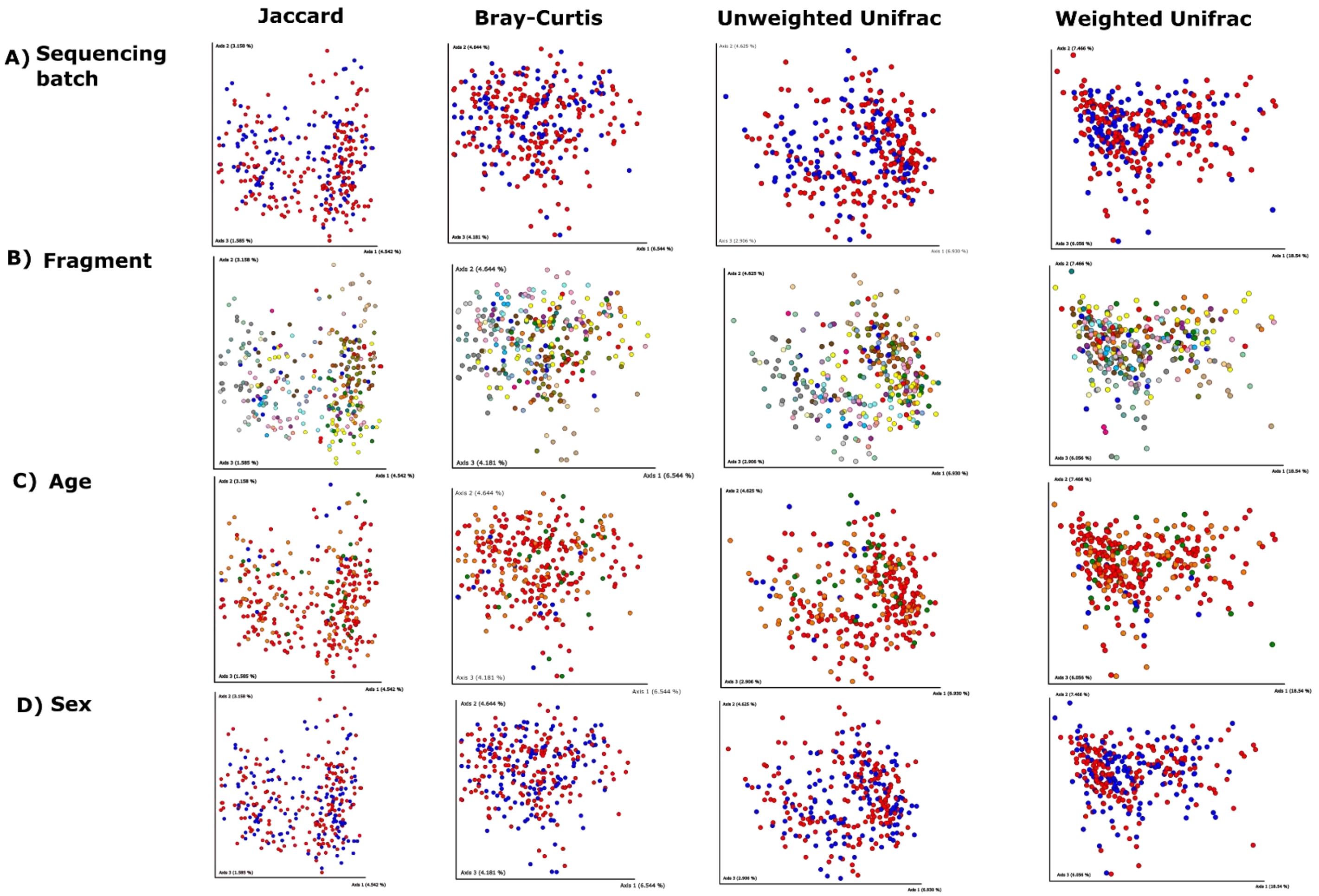

**Fig. S11. Violin plots of the habitat variables that differed significantly between fragments in the high-fragmented (HF) and low-fragmented (LF) regions when fragments were grouped according to the microbiome PCoA plot results**.

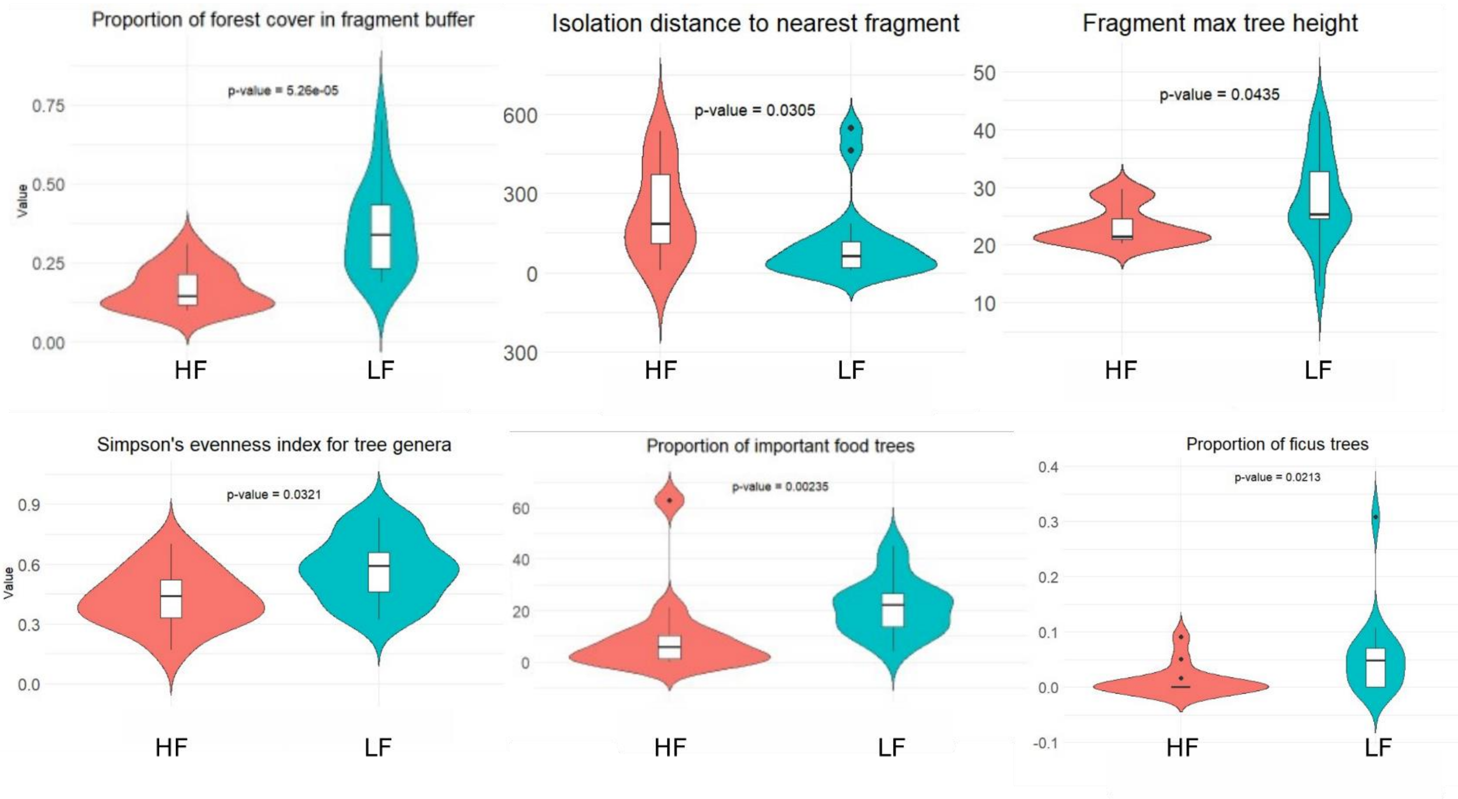

**Fig. S12. Regrouping of study fragments according to microbiome spatial patterns.** A) Map of the study site with the eight high-fragmented (HF) locations with microbiomes more similar to the low-fragmented (LF) region colored in yellow; for all subsequent microbiome analyses, these yellow fragments were grouped with the LF region (blue), not the HF region (red). B) By-individual PCoA plots showing sorting of microbiomes by region, with the individuals from the LF-adjacent fragments grouped with the LF region fragments (in blue), as opposed to with the HF fragments (red) as they were originally grouped. Results shown for: i. Jaccard dissimilarity index; ii. Unweighted Unifrac dissimilarity index; iii. Weighted Unifrac dissimilarity index.

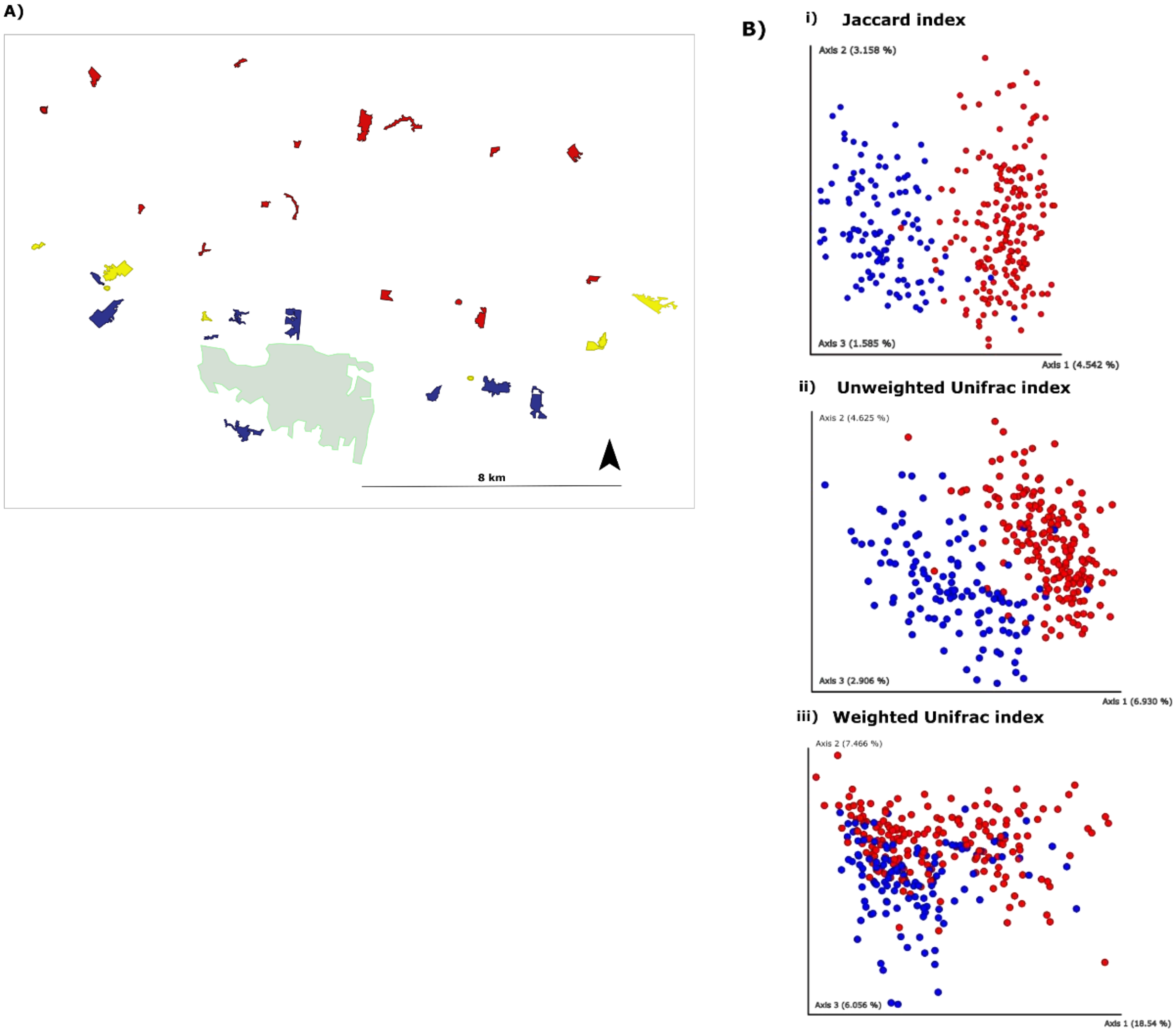

**Types and drivers of variation in microbiomes across fragments and regions**

**Table S12. Summary of SIMPER analysis results** (see below table for additional text regarding the results). Column headers: **Average:** The average contribution of that taxon to the overall dissimilarity between the two groups. Higher values mean greater influence on group separation. **SD**: The standard deviation of that contribution across samples, showing how consistently the taxon contributes to the difference. **Ratio**: The average-to-standard deviation ratio (average / SD), indicating how consistently the taxon contributes — higher ratios mean more consistent contributions. **avHF**: The average abundance of the taxon among individuals in the HF region. **avLF**: The average abundance of the taxon among individuals in the LF region. **P-value**: The p-value from permutation testing, indicating whether the contribution of that taxon to group differences is statistically significant. **Contribution (not cumulative):** The individual percentage contribution of each taxon to the total dissimilarity, not accumulated — shows each taxon's specific impact. Taxa above the bold line in the table contributed significantly ($p<0.05$) to the differences between the two regions, as quantified with the Bray-Curtis dissimilarity index. Taxa colored in pale green were not classified beyond the level of Class, and so were excluded from further analyses. Note that the contribution of some families, such as Lachnospiraceae, to the differentiation between regions was greater than Fig. 4D may indicate, given that some ASVs were identified to the level of genus (and are noted accordingly in Fig. 4D, without specifying family), and some to the level of family.

| Taxon | average | SD | Ratio | avHF | avLF | p-value | contribution to overall BC index | More in HF or LF |
|---|---|---|---|---|---|---|---|---|
| d__Bacteria.p__Firmicutes_A.c__Clostridia_258483.o__Lachnospirales.f__Lachnospiraceae.__ | 0.0335 | 0.028 | 1.197 | 916.99 | 757.18 | 0.001 | 0.0750 | HF |
| d__Bacteria.p__Firmicutes_A.c__Clostridia_258483.o__Oscillospirales.f__Oscillospiraceae_88309.g__Lawsonibacter | 0.0174 | 0.020 | 0.869 | 249.82 | 138.17 | 0.001 | 0.0390 | HF |
| d__Bacteria.p__Firmicutes_A.c__Clostridia_258483.o__Oscillospirales.f__Oscillospiraceae_88309.__ | 0.0145 | 0.018 | 0.798 | 234.97 | 123.10 | 0.001 | 0.0324 | HF |
| d__Bacteria.p__Firmicutes_A.c__Clostridia_258483.o__Oscillospirales.f__Ruminococcaceae.g__Faecalibacterium | 0.0143 | 0.014 | 1.039 | 230.92 | 190.01 | 0.003 | 0.0319 | HF |
| d__Bacteria.p__Firmicutes_A.c__Clostridia_258483.o__Lachnospirales.f__Lachnospiraceae.g__COE1 | 0.0102 | 0.009 | 1.140 | 81.13 | 141.49 | 0.025 | 0.0228 | LF |
| d__Bacteria.p__Firmicutes_A.c__Clostridia_258483.o__Lachnospirales.f__Lachnospiraceae.g__Porcincola | 0.0088 | 0.009 | 1.027 | 117.83 | 90.61 | 0.001 | 0.0197 | HF |
| d__Bacteria.p__Firmicutes_A.c__Clostridia_258483.o__Oscillospirales.f__Acutalibacteraceae.g__Ruminococcus_E | 0.0083 | 0.012 | 0.670 | 90.10 | 48.80 | 0.001 | 0.0186 | HF |
| d__Bacteria.p__Firmicutes_A.c__Clostridia_258483.o__Oscillospirales.f__Ruminococcaceae.g__Ruminococcus_D | 0.0059 | 0.006 | 1.037 | 67.79 | 50.70 | 0.009 | 0.0131 | HF |

| | | | | | | | | |
|---|---|---|---|---|---|---|---|---|
| d__Bacteria.p__Firmicutes_D.c__Bacilli.o__Erysipelotrichales.f__Coprobacillaceae.__ | 0.0057 | 0.007 | 0.820 | 84.73 | 47.02 | 0.001 | 0.0128 | HF |
| d__Bacteria.p__Firmicutes_A.c__Clostridia_258483.o__Oscillospirales.f__Acutalibacteraceae.g__Pseudoruminococcus_A | 0.0048 | 0.007 | 0.724 | 44.28 | 17.96 | 0.001 | 0.0108 | HF |
| d__Bacteria.p__Firmicutes_A.c__Clostridia_258483.o__Oscillospirales.f__Ruminococcaceae.g__Ruminococcus_C_58660 | 0.0043 | 0.004 | 1.134 | 19.50 | 43.05 | 0.001 | 0.0096 | LF |
| d__Bacteria.p__Firmicutes_A.c__Clostridia_258483.o__Lachnospirales.f__Lachnospiraceae.g__Agathobacter_164119 | 0.0035 | 0.005 | 0.742 | 32.40 | 17.79 | 0.004 | 0.0079 | HF |
| d__Bacteria.p__Firmicutes_A.c__Clostridia_258483.o__Oscillospirales.f__Ruminococcaceae.g__Paludicola | 0.0030 | 0.008 | 0.358 | 27.42 | 8.04 | 0.001 | 0.0067 | HF |
| d__Bacteria.p__Bacteroidota.c__Bacteroidia.o__Bacteroidales.f__Muribaculaceae.__ | 0.0030 | 0.006 | 0.503 | 26.49 | 6.14 | 0.001 | 0.0066 | HF |
| d__Bacteria.p__Firmicutes_C.c__Negativicutes.o__Veillonellales.f__Dialisteraceae.g__UBA5809 | 0.0023 | 0.002 | 1.088 | 10.49 | 27.99 | 0.004 | 0.0051 | LF |
| d__Bacteria.p__Firmicutes_A.c__Clostridia_258483.o__Christensenellales.f__CAG.74.g__ | 0.0018 | 0.002 | 0.967 | 6.68 | 22.85 | 0.001 | 0.0041 | LF |
| d__Bacteria.p__Actinobacteriota.c__Coriobacteriia.o__Coriobacteriales.f__Atopobiaceae.__ | 0.0017 | 0.002 | 1.048 | 23.03 | 18.21 | 0.014 | 0.0038 | HF |
| d__Bacteria.p__Firmicutes_A.c__Clostridia_258483.o__Oscillospirales.f__Acutalibacteraceae.g__RUG420 | 0.0014 | 0.001 | 1.033 | 16.24 | 8.61 | 0.001 | 0.0032 | HF |
| d__Bacteria.p__Firmicutes_A.c__Clostridia_258483.o__Oscillospirales.f__Acutalibacteraceae.g__Fimenecus | 0.0014 | 0.002 | 0.662 | 11.57 | 5.77 | 0.003 | 0.0031 | HF |
| d__Bacteria.p__Firmicutes_C.c__Negativicutes.o__Acidaminococcales.f__Acidaminococcaceae.g__Succiniclasticum | 0.0013 | 0.001 | 1.254 | 13.85 | 8.34 | 0.001 | 0.0030 | HF |
| d__Bacteria.p__Firmicutes_A.c__Clostridia_258483.o__Lachnospirales.f__Lachnospiraceae.g__Merdisoma | 0.0012 | 0.001 | 1.099 | 19.90 | 16.98 | 0.046 | 0.0027 | HF |
| d__Bacteria.p__Firmicutes_A.c__Clostridia_258483.o__Oscillospirales.f__Acutalibacteraceae.g__RUG762 | 0.0009 | 0.001 | 0.724 | 7.09 | 4.41 | 0.013 | 0.0020 | HF |
| d__Bacteria.p__Firmicutes_A.c__Clostridia_258483.o__Oscillospirales.f__Acutalibacteraceae.g__Eubacterium_R | 0.0009 | 0.001 | 1.253 | 11.11 | 7.16 | 0.001 | 0.0019 | HF |
| d__Bacteria.p__Firmicutes_D.c__Bacilli.o__RFN20.f__CAG.288.g__Asteroleplasma | 0.0007 | 0.003 | 0.254 | 6.76 | 1.70 | 0.001 | 0.0016 | HF |

| | | | | | | | | |
|---|---|---|---|---|---|---|---|---|
| d__Bacteria.p__Firmicutes_A.c__Clostridia_258483.o__Oscillospirales.f__Oscillospiraceae_88309.g__ | 0.0007 | 0.001 | 1.158 | 8.87 | 5.44 | 0.001 | 0.0015 | HF |
| d__Archaea.p__Thermoplasmatota.c__Thermoplasmata_1773.o__Methanomassiliicoccales.f__Methanomethylophilaceae.g__Methanomethylophilus | 0.0006 | 0.001 | 0.781 | 5.95 | 2.74 | 0.002 | 0.0014 | HF |
| d__Bacteria.p__Firmicutes_A.c__Clostridia_258483.o__Lachnospirales.f__Lachnospiraceae.g__ | 0.0005 | 0.001 | 0.404 | 3.62 | 1.40 | 0.01 | 0.0010 | HF |
| d__Bacteria.p__Synergistota.c__Synergistia.o__Synergistales.f__Synergistaceae.__ | 0.0004 | 0.000 | 0.938 | 4.06 | 2.69 | 0.019 | 0.0010 | HF |
| d__Bacteria.p__Actinobacteriota.c__Coriobacteriia.o__Coriobacteriales.f__Atopobiaceae.g__UBA1367 | 0.0004 | 0.001 | 0.700 | 2.97 | 1.20 | 0.001 | 0.0008 | HF |
| d__Bacteria.p__Bacteroidota.c__Bacteroidia.o__Bacteroidales.f__Marinilabiliaceae.g__JC017 | 0.0003 | 0.001 | 0.507 | 2.86 | 1.30 | 0.004 | 0.0008 | HF |
| d__Bacteria.p__Bacteroidota.c__Bacteroidia.o__Bacteroidales.f__Tannerellaceae.__ | 0.0003 | 0.000 | 0.686 | 2.34 | 1.00 | 0.001 | 0.0006 | HF |
| d__Bacteria.p__Actinobacteriota.c__Coriobacteriia.o__Coriobacteriales.f__QAMH01.g__QAMH01 | 0.0003 | 0.000 | 0.755 | 2.20 | 1.21 | 0.001 | 0.0006 | HF |
| d__Bacteria.p__Firmicutes_A.c__Clostridia_258483.o__Peptostreptococcales.f__Anaerovoracaceae.__ | 0.0002 | 0.000 | 0.766 | 1.97 | 0.96 | 0.001 | 0.0005 | HF |
| d__Bacteria.p__Proteobacteria.c__Alphaproteobacteria.o__Caulobacterales.f__Caulobacteraceae.g__Brevundimonas | 0.0002 | 0.001 | 0.194 | 1.66 | 0.06 | 0.001 | 0.0004 | HF |
| d__Bacteria.p__Cyanobacteria.c__Cyanobacteriia.o__PCC.6307.f__Cyanobiaceae.g__Cyanobium_A_31525 | 0.0001 | 0.000 | 0.182 | 0.80 | 0.02 | 0.002 | 0.0002 | HF |
| d__Bacteria.p__Actinobacteriota.c__Coriobacteriia.o__Coriobacteriales.f__Eggerthellaceae.g__ZJ304 | 0.0001 | 0.000 | 0.418 | 0.72 | 0.11 | 0.001 | 0.0002 | HF |
| d__Bacteria.p__Firmicutes_A.c__Clostridia_258483.o__Lachnospirales.f__Lachnospiraceae.g__Eubacterium_J | 0.0001 | 0.000 | 0.278 | 0.61 | 0.16 | 0.014 | 0.0002 | HF |
| d__Bacteria.p__Proteobacteria.c__Gammaproteobacteria.o__Enterobacterales_A_737866.f__Enterobacteriaceae_A.g__Pantoea_A_680069 | 0.0001 | 0.001 | 0.092 | 0.72 | 0.00 | 0.001 | 0.0002 | HF |
| d__Bacteria.p__Firmicutes_A.c__Clostridia_258483.o__Oscillospirales.f__Oscillospiraceae_88309.g__UBA2658 | 0.0001 | 0.000 | 0.201 | 0.70 | 0.01 | 0.003 | 0.0002 | HF |
| d__Bacteria.p__Firmicutes_A.c__Clostridia_258483.o__Peptostreptococcales.f__Anaerovoracaceae.g__RUG13615 | 0.0001 | 0.000 | 0.440 | 0.49 | 0.27 | 0.044 | 0.0002 | HF |

| | | | | | | | | |
|---|---|---|---|---|---|---|---|---|
| d__Bacteria.p__Proteobacteria.c__Alphaproteobacteria.o__Sphingomonadales.f__Sphingomonadaceae.g__Blastomonas | 0.0000 | 0.000 | 0.213 | 0.41 | 0.07 | 0.034 | 0.0001 | HF |
| d__Bacteria.p__Proteobacteria.c__Alphaproteobacteria.o__Sphingomonadales.f__Sphingomonadaceae.__ | 0.0000 | 0.000 | 0.204 | 0.44 | 0.02 | 0.003 | 0.0001 | HF |
| d__Bacteria.p__Firmicutes_A.c__Clostridia_258483.o__Lachnospirales.f__Lachnospiraceae.g__Butyrivibrio_A_168226 | 0.0000 | 0.000 | 0.319 | 0.29 | 0.04 | 0.001 | 0.0001 | HF |
| d__Bacteria.p__Bacteroidota.c__Bacteroidia.o__Chitinophagales.f__Chitinophagaceae_966727.g__Lacibacter | 0.0000 | 0.000 | 0.155 | 0.23 | 0.02 | 0.027 | 0.0001 | HF |
| d__Bacteria.p__Bacteroidota.c__Bacteroidia.o__Sphingobacteriales.f__Sphingobacteriaceae.g__Pedobacter_887417 | 0.0000 | 0.000 | 0.175 | 0.22 | 0.02 | 0.028 | 0.0001 | HF |
| d__Bacteria.p__Proteobacteria.c__Alphaproteobacteria.o__Reyranellales.f__Reyranellaceae.g__Reyranella | 0.0000 | 0.000 | 0.209 | 0.21 | 0.02 | 0.006 | 0.0001 | HF |
| d__Bacteria.p__Firmicutes_A.c__Clostridia_258483.o__Oscillospirales.f__Oscillospiraceae_88309.g__Sporobacter | 0.0000 | 0.000 | 0.306 | 0.13 | 0.05 | 0.044 | 0.0000 | HF |
| d__Bacteria.p__Proteobacteria.c__Gammaproteobacteria.o__Enterobacterales_A_737866.f__Enterobacteriaceae_A.__ | 0.0000 | 0.000 | 0.125 | 0.14 | 0.00 | 0.001 | 0.0000 | HF |
| d__Bacteria.p__Proteobacteria.c__Gammaproteobacteria.o__Pseudomonadales_650611.f__Pseudomonadaceae.__ | 0.0000 | 0.000 | 0.098 | 0.13 | 0.00 | 0.001 | 0.0000 | HF |
| d__Bacteria.p__Proteobacteria.c__Gammaproteobacteria.o__Pseudomonadales_650611.f__Pseudomonadaceae.g__Pseudomonas_B_650451 | 0.0000 | 0.000 | 0.092 | 0.12 | 0.00 | 0.001 | 0.0000 | HF |
| d__Bacteria.p__Cyanobacteria.c__Cyanobacteriia.o__PCC.6307.f__Cyanobiaceae.g__WH.5701 | 0.0000 | 0.000 | 0.092 | 0.09 | 0.00 | 0.001 | 0.0000 | HF |
| d__Bacteria.p__Cyanobacteria.c__Cyanobacteriia.o__Cyanobacteriales.f__Chroococcidiopsidaceae_29158.g__Chroococcidiopsis_29153 | 0.0000 | 0.000 | 0.110 | 0.09 | 0.00 | 0.001 | 0.0000 | HF |
| d__Bacteria.p__Actinobacteriota.c__Coriobacteriia.o__Coriobacteriales.f__Eggerthellaceae.g__UBA9715 | 0.0000 | 0.000 | 0.092 | 0.08 | 0.00 | 0.001 | 0.0000 | HF |
| d__Bacteria.p__Proteobacteria.c__Alphaproteobacteria.o__Sphingomonadales.f__Sphingomonadaceae.g__Sphingomonas_L_486704 | 0.0000 | 0.000 | 0.092 | 0.08 | 0.00 | 0.001 | 0.0000 | HF |
| d__Bacteria.p__Chlamydiota.c__Chlamydiia.o__Chlamydiales_778124.f__Parachlamydiaceae.g__HS.T3 | 0.0000 | 0.000 | 0.092 | 0.07 | 0.00 | 0.001 | 0.0000 | HF |

| | | | | | | | | |
|---|---|---|---|---|---|---|---|---|
| d__Bacteria.p__Proteobacteria.c__Gammaproteobacteria.o__Burkholderiales_592522.f__Burkholderiaceae_A_592522.g__Polaromonas | 0.0000 | 0.000 | 0.092 | 0.07 | 0.00 | 0.001 | 0.0000 | HF |
| d__Bacteria.p__Proteobacteria.c__Gammaproteobacteria.o__Xanthomonadales_616009.f__Xanthomonadaceae_616009.g__Luteimonas_C_615545 | 0.0000 | 0.000 | 0.120 | 0.06 | 0.00 | 0.001 | 0.0000 | HF |
| d__Bacteria.p__Proteobacteria.c__Gammaproteobacteria.o__Burkholderiales_592524.f__Burkholderiaceae_A_574758.g__Massilia | 0.0000 | 0.000 | 0.124 | 0.05 | 0.00 | 0.001 | 0.0000 | HF |
| d__Bacteria.p__Proteobacteria.c__Gammaproteobacteria.o__Pseudomonadales_650611.f__Pseudomonadaceae.g__Pseudomonas_E_647464 | 0.0000 | 0.000 | 0.092 | 0.05 | 0.00 | 0.001 | 0.0000 | HF |
| d__Bacteria.p__Proteobacteria.c__Gammaproteobacteria.o__Pseudomonadales_650611.f__Pseudomonadaceae.g__Pseudomonas_E_650325 | 0.0000 | 0.000 | 0.140 | 0.04 | 0.00 | 0.001 | 0.0000 | HF |
| d__Bacteria.p__Firmicutes_D.c__Bacilli.o__Lactobacillales.f__Lactobacillaceae.g__Weissella_A_338544 | 0.0000 | 0.000 | 0.092 | 0.04 | 0.00 | 0.001 | 0.0000 | HF |
| d__Bacteria.p__Proteobacteria.c__Alphaproteobacteria.o__Rhizobiales_A_501059.f__Rhizobiaceae_A_501059.g__Mycoplana_499574 | 0.0000 | 0.000 | 0.092 | 0.03 | 0.00 | 0.001 | 0.0000 | HF |
| d__Bacteria.p__Cyanobacteria.c__Cyanobacteriia.o__Elainellales.f__Elainellaceae.g__O.77 | 0.0000 | 0.000 | 0.117 | 0.03 | 0.00 | 0.001 | 0.0000 | HF |
| d__Bacteria.p__Actinobacteriota.c__Actinomycetia.o__Actinomycetales.f__Cellulomonadaceae.g__Actinotalea | 0.0000 | 0.000 | 0.092 | 0.03 | 0.00 | 0.001 | 0.0000 | HF |
| d__Bacteria.p__Actinobacteriota.c__Actinomycetia.o__Actinomycetales.f__Microbacteriaceae.__ | 0.0000 | 0.000 | 0.117 | 0.03 | 0.00 | 0.001 | 0.0000 | HF |
| d__Bacteria.p__Bacteroidota.c__Bacteroidia.o__Cytophagales.f__Spirosomaceae.__ | 0.0000 | 0.000 | 0.117 | 0.03 | 0.00 | 0.001 | 0.0000 | HF |
| d__Bacteria.p__Actinobacteriota.c__Actinomycetia.o__Propionibacteriales.f__Nocardioidaceae.g__Aeromicrobium | 0.0000 | 0.000 | 0.124 | 0.03 | 0.00 | 0.001 | 0.0000 | HF |
| d__Bacteria.p__Firmicutes_A.c__Clostridia_258483.o__Oscillospirales.f__Butyricicoccaceae.g__Pseudobutyricicoccus | 0.0000 | 0.000 | 0.092 | 0.03 | 0.00 | 0.001 | 0.0000 | HF |
| d__Bacteria.p__Proteobacteria.c__Alphaproteobacteria.o__Rhizobiales_A_501396.f__Rhizobiaceae_A_499470.__ | 0.0000 | 0.000 | 0.092 | 0.02 | 0.00 | 0.001 | 0.0000 | HF |
| d__Bacteria.p__Planctomycetota.c__Planctomycetia.o__Gemmatales.f__Gemmataceae.g__Urbifossiella | 0.0000 | 0.000 | 0.092 | 0.02 | 0.00 | 0.001 | 0.0000 | HF |

| | | | | | | | | |
|---|---|---|---|---|---|---|---|---|
| d__Bacteria.p__Bacteroidota.c__Bacteroidia.o__Chitinophagales.f__Chitinophagaceae_929731.__ | 0.0000 | 0.000 | 0.092 | 0.02 | 0.00 | 0.001 | 0.0000 | HF |
| d__Bacteria.p__Bacteroidota.c__Bacteroidia.o__Flavobacteriales_877923.f__Weeksellaceae.g__Chryseobacterium_796614 | 0.0000 | 0.000 | 0.092 | 0.02 | 0.00 | 0.001 | 0.0000 | HF |
| d__Bacteria.p__Proteobacteria.c__Alphaproteobacteria.o__Acetobacterales.f__Acetobacteraceae.g__Roseomonas_A_507058 | 0.0000 | 0.000 | 0.131 | 0.02 | 0.00 | 0.001 | 0.0000 | HF |
| d__Bacteria.p__Proteobacteria.c__Gammaproteobacteria.o__Burkholderiales_592522.f__Burkholderiaceae_A_592522.g__Ramlibacter_588642 | 0.0000 | 0.000 | 0.092 | 0.02 | 0.00 | 0.001 | 0.0000 | HF |
| d__Bacteria.p__Proteobacteria.c__Alphaproteobacteria.o__Rhizobiales_A_504721.f__Methyloligellaceae.g__Methyloceanibacter | 0.0000 | 0.000 | 0.092 | 0.02 | 0.00 | 0.001 | 0.0000 | HF |
| d__Bacteria.p__Proteobacteria.c__Alphaproteobacteria.o__Rhizobiales_A_504705.f__Phreatobacteraceae.g__Phreatobacter | 0.0000 | 0.000 | 0.092 | 0.02 | 0.00 | 0.001 | 0.0000 | HF |
| d__Bacteria.p__Bacteroidota.c__Bacteroidia.o__Bacteroidales.f__Prolixibacteraceae.__ | 0.0000 | 0.000 | 0.092 | 0.01 | 0.00 | 0.001 | 0.0000 | HF |
| d__Bacteria.p__Bacteroidota.c__Bacteroidia.o__AKYH767.f__B.17BO.g__UBA2475 | 0.0000 | 0.000 | 0.092 | 0.01 | 0.00 | 0.001 | 0.0000 | HF |
| d__Bacteria.p__Acidobacteriota.c__Acidobacteriae.o__Acidobacteriales.f__SbA1.g__Gp1.AA122 | 0.0000 | 0.000 | 0.092 | 0.01 | 0.00 | 0.001 | 0.0000 | HF |
| d__Bacteria.p__Verrucomicrobiota.c__Verrucomicrobiae.o__Opitutales.f__Opitutaceae.__ | 0.0000 | 0.000 | 0.092 | 0.01 | 0.00 | 0.001 | 0.0000 | HF |
| d__Bacteria.p__Bacteroidota.c__Bacteroidia.o__Bacteroidales.f__Porphyromonadaceae.g__Porphyromonas_A_859423 | 0.0000 | 0.000 | 0.092 | 0.01 | 0.00 | 0.001 | 0.0000 | HF |
| d__Bacteria.p__Actinobacteriota.c__Actinomycetia.o__Mycobacteriales.f__Mycobacteriaceae.g__Nocardia | 0.0000 | 0.000 | 0.092 | 0.01 | 0.00 | 0.001 | 0.0000 | HF |
| d__Bacteria.p__Actinobacteriota.c__Actinomycetia.o__Mycobacteriales.f__Micromonosporaceae.g__Actinoplanes | 0.0000 | 0.000 | 0.092 | 0.01 | 0.00 | 0.001 | 0.0000 | HF |
| d__Bacteria.p__Proteobacteria.c__Gammaproteobacteria.o__Burkholderiales_592524.f__Burkholderiaceae_A_580492.g__Pandoraea | 0.0000 | 0.000 | 0.092 | 0.01 | 0.00 | 0.001 | 0.0000 | HF |
| d__Bacteria.p__Proteobacteria.c__Gammaproteobacteria.o__Burkholderiales_592522.f__Burkholderiaceae_A_592522.g__Aquabacterium_B_592457 | 0.0000 | 0.000 | 0.092 | 0.01 | 0.00 | 0.001 | 0.0000 | HF |

| | | | | | | | | |
|---|---|---|---|---|---|---|---|---|
| d__Bacteria.p__Actinobacteriota.c__Coriobacteriia.o__Coriobacteriales.f__Eggerthellaceae.g__CAAEEV01 | 0.0000 | 0.000 | 0.092 | 0.01 | 0.00 | 0.001 | 0.0000 | HF |
| d__Bacteria.p__Verrucomicrobiota.c__Verrucomicrobiae.o__Opitutales.f__.g__ | 0.0000 | 0.000 | 0.092 | 0.02 | 0.00 | 0.001 | 0.0000 | HF |
| d__Bacteria.p__Proteobacteria.c__Alphaproteobacteria.o__Caedimonadales.f__.g__ | 0.0000 | 0.000 | 0.092 | 0.03 | 0.00 | 0.001 | 0.0000 | HF |
| d__Bacteria.__.__.__.__.__ | 0.0156 | 0.028 | 0.561 | 151.88 | 96.57 | 0.016 | 0.0349 | HF |
| d__Bacteria.p__Bacteroidota.c__Bacteroidia.__.__.__ | 0.0042 | 0.005 | 0.842 | 43.15 | 29.75 | 0.004 | 0.0095 | HF |
| d__Bacteria.p__Firmicutes_A.c__Clostridia_258483.o__Oscillospirales.f__Ruminococcaceae.g__Ruminiclostridium_E | 0.0025 | 0.006 | 0.394 | 19.33 | 10.37 | 0.05 | 0.0056 | |
| d__Bacteria.p__Firmicutes_D.c__Bacilli.o__Aneurinibacillales.f__RAOX.1.g__YIM.78166 | 0.0000 | 0.000 | 0.141 | 0.03 | 0.01 | 0.051 | 0.0000 | |
| d__Bacteria.p__Firmicutes_A.c__Clostridia_258483.o__Oscillospirales.f__Ruminococcaceae.g__Gemmiger_A_73129 | 0.0062 | 0.007 | 0.919 | 59.01 | 55.95 | 0.052 | 0.0138 | |
| d__Bacteria.p__Proteobacteria.c__Alphaproteobacteria.o__Sphingomonadales.f__Sphingomonadaceae.g__Sphingopyxis | 0.0001 | 0.000 | 0.223 | 0.46 | 0.12 | 0.053 | 0.0001 | |
| d__Bacteria.p__Bacteroidota.c__Bacteroidia.o__Flavobacteriales_877923.f__Flavobacteriaceae.g__Flavobacterium | 0.0000 | 0.000 | 0.125 | 0.22 | 0.02 | 0.053 | 0.0001 | |
| d__Bacteria.p__Firmicutes_A.c__Clostridia_258483.o__Lachnospirales.f__Lachnospiraceae.g__Coprococcus_A_121497 | 0.0016 | 0.005 | 0.356 | 11.88 | 5.88 | 0.055 | 0.0036 | |
| d__Bacteria.p__Firmicutes_D.c__Bacilli.o__Lactobacillales.f__Streptococcaceae.g__Streptococcus | 0.0003 | 0.001 | 0.383 | 2.32 | 1.11 | 0.06 | 0.0007 | |
| d__Bacteria.p__Cyanobacteria.c__Vampirovibrionia.o__Gastranaerophilales.f__Gastranaerophilaceae.g__Zag111 | 0.0066 | 0.013 | 0.518 | 53.06 | 39.85 | 0.061 | 0.0148 | |
| d__Bacteria.p__Proteobacteria.c__Alphaproteobacteria.o__Rhizobiales_A_500471.f__Rhizobiaceae_A_500471.g__Agrobacterium | 0.0000 | 0.000 | 0.138 | 0.08 | 0.01 | 0.061 | 0.0000 | |
| d__Bacteria.p__Cyanobacteria.c__Vampirovibrionia.o__Gastranaerophilales.f__Gastranaerophilaceae.g__CAG.196 | 0.0001 | 0.000 | 0.323 | 0.41 | 0.20 | 0.067 | 0.0001 | |
| d__Bacteria.p__Proteobacteria.c__Gammaproteobacteria.o__Burkholderiales_592524.f__Burkholderiaceae_A_574938.g__Oxalobacter_566322 | 0.0001 | 0.000 | 0.418 | 0.45 | 0.26 | 0.073 | 0.0001 | |

| | | | | | | | |
|---|---|---|---|---|---|---|---|
| d__Bacteria.p__Proteobacteria.c__Alphaproteobacteria.o__Rhizobiales_A_504705.f__Beijerinckiaceae.__ | 0.0000 | 0.000 | 0.146 | 0.03 | 0.01 | 0.076 | 0.0000 |
| d__Bacteria.p__Firmicutes_A.c__Clostridia_258483.o__Lachnospirales.f__Lachnospiraceae.g__Paralachnospira | 0.0000 | 0.000 | 0.165 | 0.05 | 0.02 | 0.077 | 0.0000 |
| d__Bacteria.p__Firmicutes_A.c__Clostridia_258483.o__Lachnospirales.f__Lachnospiraceae.g__1XD42.69 | 0.0001 | 0.000 | 0.211 | 0.71 | 0.19 | 0.079 | 0.0002 |
| d__Bacteria.p__Firmicutes_D.c__Bacilli.o__RF39.f__UBA660.g__UBA5026 | 0.0000 | 0.000 | 0.189 | 0.27 | 0.06 | 0.079 | 0.0001 |
| d__Bacteria.p__Actinobacteriota.c__Coriobacteriia.o__Coriobacteriales.f__Atopobiaceae.g__Thermophilibacter | 0.0008 | 0.002 | 0.491 | 7.29 | 4.85 | 0.088 | 0.0017 |
| d__Bacteria.p__Firmicutes_D.c__Bacilli.o__RF39.f__UBA660.g__Faecimonas | 0.0002 | 0.001 | 0.421 | 1.55 | 0.87 | 0.089 | 0.0005 |
| d__Bacteria.p__Firmicutes_D.c__Bacilli.o__RFN20.f__CAG.826.__ | 0.0000 | 0.000 | 0.131 | 0.14 | 0.02 | 0.09 | 0.0000 |
| d__Archaea.p__Thermoplasmatota.c__Thermoplasmata_1773.o__Methanomassiliicoccales.f__Methanomethylophilaceae.g__UBA71 | 0.0002 | 0.000 | 0.483 | 1.41 | 0.90 | 0.113 | 0.0005 |
| d__Bacteria.p__Firmicutes_A.c__Clostridia_258483.__.__.__ | 0.0118 | 0.014 | 0.856 | 100.97 | 133.84 | 0.128 | 0.0265 |
| d__Bacteria.p__Firmicutes_A.c__Clostridia_258483.o__Lachnospirales.f__Lachnospiraceae.g__Bilifractor | 0.0006 | 0.001 | 0.810 | 5.08 | 4.31 | 0.13 | 0.0013 |
| d__Bacteria.p__Firmicutes_A.c__Clostridia_258483.o__Oscillospirales.f__Ruminococcaceae.g__UBA1394 | 0.0003 | 0.000 | 0.801 | 1.21 | 2.20 | 0.137 | 0.0006 |
| d__Bacteria.p__Proteobacteria.c__Gammaproteobacteria.o__Burkholderiales_592522.f__Burkholderiaceae_A_592522.g__Variovorax | 0.0000 | 0.000 | 0.168 | 0.08 | 0.03 | 0.14 | 0.0000 |
| d__Bacteria.p__Firmicutes_A.c__Clostridia_258483.o__TANB77.f__CAG.508.g__ | 0.0085 | 0.010 | 0.811 | 81.08 | 71.92 | 0.142 | 0.0190 |
| d__Bacteria.p__Proteobacteria.c__Gammaproteobacteria.o__Burkholderiales_592524.__.__ | 0.0000 | 0.000 | 0.118 | 0.02 | 0.01 | 0.147 | 0.0000 |
| d__Bacteria.p__Firmicutes_A.c__Clostridia_258483.o__Peptostreptococcales.f__Anaerovoracaceae.g__ | 0.0023 | 0.004 | 0.595 | 22.12 | 24.26 | 0.151 | 0.0051 |
| d__Bacteria.p__Desulfobacterota_I.c__Desulfovibrionia.o__Desulfovibrionales.f__Desulfovibrionaceae.__ | 0.0000 | 0.000 | 0.118 | 0.02 | 0.01 | 0.151 | 0.0000 |
| d__Archaea.p__Thermoplasmatota.c__Thermoplasmata_1773.o__Methanomassiliicoccales.f__Methanomethylophilaceae.__ | 0.0000 | 0.000 | 0.118 | 0.01 | 0.01 | 0.151 | 0.0000 |

| | | | | | | | |
|---|---|---|---|---|---|---|---|
| d__Bacteria.p__Bacteroidota.c__Bacteroidia.o__Bacteroidales.f__Bacteroidaceae.g__43.108 | 0.0000 | 0.000 | 0.118 | 0.03 | 0.02 | 0.153 | 0.0000 |
| d__Bacteria.p__Firmicutes_A.c__Clostridia_258483.o__Lachnospirales.f__Lachnospiraceae.g__Eubacterium_Q | 0.0000 | 0.000 | 0.118 | 0.01 | 0.01 | 0.153 | 0.0000 |
| d__Bacteria.p__Planctomycetota.c__Planctomycetia.o__Planctomycetales.f__Planctomycetaceae.g__Planctopirus | 0.0000 | 0.000 | 0.116 | 0.03 | 0.01 | 0.154 | 0.0000 |
| d__Bacteria.p__Firmicutes_D.c__Bacilli.o__Bacillales_B_310392.f__Bacillaceae_G_310392.g__Bacillus_A | 0.0000 | 0.000 | 0.116 | 0.03 | 0.01 | 0.158 | 0.0000 |
| d__Bacteria.p__Firmicutes_A.c__Clostridia_258483.o__Oscillospirales.f__Acutalibacteraceae.g__ | 0.0000 | 0.000 | 0.105 | 0.03 | 0.01 | 0.168 | 0.0000 |
| d__Bacteria.p__Firmicutes_D.c__Bacilli.o__RF39.f__UBA660.g__RUG13038 | 0.0007 | 0.001 | 0.781 | 4.60 | 4.53 | 0.176 | 0.0015 |
| d__Bacteria.p__Proteobacteria.c__Gammaproteobacteria.o__Burkholderiales_597441.f__Methylophilaceae.g__Methylophilus | 0.0000 | 0.000 | 0.150 | 0.03 | 0.02 | 0.185 | 0.0000 |
| d__Bacteria.p__Firmicutes_C.c__Negativicutes.o__Acidaminococcales.__.__ | 0.0000 | 0.000 | 0.234 | 0.10 | 0.06 | 0.211 | 0.0000 |
| d__Bacteria.p__Firmicutes_D.c__Bacilli.o__RFN20.f__CAG.826.g__Enteromonas | 0.0020 | 0.007 | 0.314 | 12.95 | 8.36 | 0.213 | 0.0046 |
| d__Bacteria.p__Bacteroidota.c__Bacteroidia.o__Bacteroidales.f__Bacteroidaceae.g__Alloprevotella | 0.0000 | 0.000 | 0.129 | 0.03 | 0.02 | 0.224 | 0.0000 |
| d__Bacteria.p__Firmicutes_D.c__Bacilli.o__Erysipelotrichales.f__Erysipelotrichaceae.g__Bulleidia | 0.0000 | 0.000 | 0.313 | 0.15 | 0.10 | 0.244 | 0.0001 |
| d__Bacteria.p__Proteobacteria.c__Gammaproteobacteria.o__Burkholderiales_592522.f__Burkholderiaceae_A_592522.__ | 0.0000 | 0.000 | 0.183 | 0.09 | 0.05 | 0.246 | 0.0000 |
| d__Bacteria.p__Firmicutes_A.c__Clostridia_258483.o__Peptostreptococcales.f__Anaerovoracaceae.g__RUG754 | 0.0002 | 0.000 | 0.452 | 1.01 | 0.77 | 0.259 | 0.0004 |
| d__Bacteria.p__Firmicutes_D.c__Bacilli.o__RF39.f__UBA660.g__UMGS1449 | 0.0000 | 0.000 | 0.156 | 0.07 | 0.03 | 0.262 | 0.0000 |
| d__Bacteria.p__Firmicutes_A.c__Clostridia_258483.o__UBA1381.f__UBA1381.g__CAG.41 | 0.0040 | 0.004 | 0.978 | 53.17 | 51.81 | 0.269 | 0.0090 |
| d__Bacteria.p__Bacteroidota.c__Bacteroidia.o__Bacteroidales.f__UBA932.g__Merdivivens | 0.0001 | 0.000 | 0.310 | 0.39 | 0.28 | 0.269 | 0.0001 |
| d__Bacteria.p__Firmicutes_A.c__Clostridia_258483.o__Oscillospirales.f_ | 0.0216 | 0.018 | 1.188 | 360.08 | 379.55 | 0.279 | 0.0482 |

| | | | | | | | |
|---|---|---|---|---|---|---|---|
| _Oscillospiraceae_88309.g__Faecousia | | | | | | | |
| d__Bacteria.p__Proteobacteria.c__Gammaproteobacteria.o__Xanthomonadales_616009.f__Xanthomonadaceae_616009.__ | 0.0000 | 0.000 | 0.180 | 0.25 | 0.12 | 0.281 | 0.0001 |
| d__Bacteria.p__Bacteroidota.c__Bacteroidia.o__Bacteroidales.__.__ | 0.0002 | 0.000 | 0.426 | 1.01 | 0.83 | 0.318 | 0.0004 |
| d__Bacteria.p__Bacteroidota.c__Bacteroidia.o__Bacteroidales.f__Bacteroidaceae.g__UBA6398 | 0.0000 | 0.000 | 0.156 | 0.08 | 0.06 | 0.338 | 0.0000 |
| d__Bacteria.p__.c__.o__.f__.g__ | 0.0000 | 0.000 | 0.217 | 0.12 | 0.10 | 0.342 | 0.0000 |
| d__Bacteria.p__Bacteroidota.c__Bacteroidia.o__Bacteroidales.f__Bacteroidaceae.__ | 0.0121 | 0.012 | 0.994 | 161.54 | 147.33 | 0.344 | 0.0270 |
| d__Bacteria.p__Proteobacteria.c__Alphaproteobacteria.o__Rhizobiales_A_501396.f__Rhizobiaceae_A_499470.g__Ochrobactrum_A_499024 | 0.0000 | 0.000 | 0.073 | 0.00 | 0.01 | 0.354 | 0.0000 |
| d__Bacteria.p__Proteobacteria.c__Gammaproteobacteria.o__Pseudomonadales_660879.f__Moraxellaceae.g__Acinetobacter | 0.0000 | 0.000 | 0.073 | 0.00 | 0.01 | 0.357 | 0.0000 |
| d__Bacteria.p__Firmicutes_A.c__Clostridia_258483.o__Lachnospirales.f__Lachnospiraceae.g__AM51.8 | 0.0000 | 0.000 | 0.073 | 0.00 | 0.03 | 0.364 | 0.0000 |
| d__Bacteria.p__Firmicutes_A.c__Clostridia_258483.o__Oscillospirales.f__Ruminococcaceae.g__Soleaferrea | 0.0000 | 0.000 | 0.073 | 0.00 | 0.02 | 0.365 | 0.0000 |
| d__Bacteria.p__Cyanobacteria.c__Vampirovibrionia.o__Gastranaerophilales.f__Gastranaerophilaceae.g__UBA2813 | 0.0024 | 0.008 | 0.291 | 14.36 | 14.55 | 0.367 | 0.0054 |
| d__Bacteria.p__Bacteroidota.c__Bacteroidia.o__Bacteroidales.f__Tannerellaceae.g__Parabacteroides_B_862066 | 0.0000 | 0.000 | 0.073 | 0.00 | 0.12 | 0.367 | 0.0000 |
| d__Bacteria.p__Firmicutes_A.c__Clostridia_258483.o__Lachnospirales.f__Lachnospiraceae.g__UMGS1375 | 0.0000 | 0.000 | 0.073 | 0.00 | 0.06 | 0.367 | 0.0000 |
| d__Bacteria.p__Firmicutes_A.c__Clostridia_258483.o__Oscillospirales.f__Ruminococcaceae.g__HUN007 | 0.0000 | 0.000 | 0.073 | 0.00 | 0.02 | 0.367 | 0.0000 |
| d__Bacteria.p__Proteobacteria.c__Alphaproteobacteria.o__RF32.f__CAG.239.g__CAJLXD01 | 0.0000 | 0.000 | 0.073 | 0.00 | 0.01 | 0.367 | 0.0000 |
| d__Bacteria.p__Firmicutes_A.c__Clostridia_258483.o__Oscillospirales.f__Ruminococcaceae.__ | 0.0074 | 0.008 | 0.931 | 42.35 | 88.59 | 0.368 | 0.0166 |
| d__Bacteria.p__Proteobacteria.c__Gammaproteobacteria.o__Burkholderiales_592522.f__Burkholderiaceae_A_592522.g__Ramlibacter_582307 | 0.0000 | 0.000 | 0.084 | 0.01 | 0.05 | 0.37 | 0.0000 |

| | | | | | | | |
|---|---|---|---|---|---|---|---|
| d__Bacteria.p__Desulfobacterota_I.c__Desulfovibrionia.o__Desulfovibrionales.f__Desulfovibrionaceae.g__Desulfovibrio_R_446353 | 0.0013 | 0.005 | 0.275 | 11.72 | 10.07 | 0.371 | 0.0029 |
| d__Bacteria.p__Firmicutes_A.c__Clostridia_258483.o__Oscillospirales.f__CAG.272.__ | 0.0000 | 0.000 | 0.073 | 0.00 | 0.02 | 0.372 | 0.0000 |
| d__Bacteria.p__Firmicutes_A.c__Clostridia_258483.o__Christensenellales.f__Christensenellaceae.g__ | 0.0000 | 0.000 | 0.073 | 0.00 | 0.01 | 0.372 | 0.0000 |
| d__Bacteria.p__Gemmatimonadota.c__Gemmatimonadetes.o__Gemmatimonadales.f__Gemmatimonadaceae.g__Gemmatimonas | 0.0000 | 0.000 | 0.073 | 0.00 | 0.01 | 0.372 | 0.0000 |
| d__Bacteria.p__Firmicutes_A.c__Clostridia_258483.o__Lachnospirales.f__Lachnospiraceae.g__CAG.45 | 0.0000 | 0.000 | 0.073 | 0.00 | 0.28 | 0.373 | 0.0001 |
| d__Bacteria.p__Firmicutes_A.c__Clostridia_258483.o__Lachnospirales.f__Lachnospiraceae.g__Sellimonas | 0.0000 | 0.000 | 0.073 | 0.00 | 0.24 | 0.373 | 0.0001 |
| d__Bacteria.p__Firmicutes_A.c__Clostridia_258483.o__Lachnospirales.f__Lachnospiraceae.g__Mediterraneibacter_A_155507 | 0.0000 | 0.000 | 0.073 | 0.00 | 0.02 | 0.373 | 0.0000 |
| d__Bacteria.p__Firmicutes_D.c__Bacilli.o__Lactobacillales.f__Lactobacillaceae.__ | 0.0000 | 0.000 | 0.073 | 0.00 | 0.02 | 0.374 | 0.0000 |
| d__Bacteria.p__Firmicutes_A.c__Clostridia_258483.o__Lachnospirales.f__CAG.274.g__CAG.274 | 0.0037 | 0.004 | 0.891 | 26.32 | 49.50 | 0.375 | 0.0082 |
| d__Bacteria.p__Bacteroidota.__.__.__.__ | 0.0000 | 0.000 | 0.171 | 0.06 | 0.05 | 0.376 | 0.0000 |
| d__Bacteria.p__Proteobacteria.c__Gammaproteobacteria.o__UBA9339.f__UBA9339.g__UBA9339 | 0.0000 | 0.000 | 0.073 | 0.00 | 0.01 | 0.376 | 0.0000 |
| d__Bacteria.p__Proteobacteria.c__Gammaproteobacteria.o__Diplorickettsiales.f__Diplorickettsiaceae.g__ | 0.0000 | 0.000 | 0.073 | 0.00 | 0.01 | 0.379 | 0.0000 |
| d__Bacteria.p__Firmicutes_B_370541.c__Syntrophomonadia.o__Syntrophomonadales.f__Syntrophomonadaceae_368540.g__Pelospora | 0.0000 | 0.000 | 0.139 | 0.02 | 0.02 | 0.381 | 0.0000 |
| d__Bacteria.p__Firmicutes_A.c__Clostridia_258483.o__Oscillospirales.f__Oscillospiraceae_88309.g__Marseille.P3106 | 0.0000 | 0.000 | 0.073 | 0.00 | 0.01 | 0.382 | 0.0000 |
| d__Bacteria.p__Firmicutes_A.c__Clostridia_258483.o__Oscillospirales.f__Acutalibacteraceae.g__Scatavimonas | 0.0000 | 0.000 | 0.115 | 0.02 | 0.02 | 0.384 | 0.0000 |
| d__Bacteria.p__Firmicutes_D.c__Bacilli.o__Acholeplasmatales.__.__ | 0.0000 | 0.000 | 0.073 | 0.00 | 0.01 | 0.386 | 0.0000 |
| d__Bacteria.p__Proteobacteria.c__Gammaproteobacteria.o__Xanthomonad | 0.0000 | 0.000 | 0.073 | 0.00 | 0.02 | 0.388 | 0.0000 |

| | | | | | | | |
|---|---|---|---|---|---|---|---|
| ales_616009.f__Xanthomonadaceae_616009.g__Xanthomonas_A_614439 | | | | | | | |
| d__Bacteria.p__Firmicutes_A.c__Clostridia_258483.o__Lachnospirales.f__Lachnospiraceae.g__VUNI01 | 0.0000 | 0.000 | 0.073 | 0.00 | 0.01 | 0.388 | 0.0000 |
| d__Bacteria.p__Gemmatimonadota.c__Gemmatimonadetes.o__Gemmatimonadales.f__Gemmatimonadaceae.__ | 0.0000 | 0.000 | 0.073 | 0.00 | 0.01 | 0.388 | 0.0000 |
| d__Bacteria.p__Firmicutes_D.c__Bacilli.o__Erysipelotrichales.f__Coprobacillaceae.g__Sharpea | 0.0000 | 0.000 | 0.073 | 0.00 | 0.02 | 0.389 | 0.0000 |
| d__Bacteria.p__Proteobacteria.c__Gammaproteobacteria.o__Xanthomonadales_616009.f__Xanthomonadaceae_616009.g__Stenotrophomonas_A_615274 | 0.0000 | 0.000 | 0.073 | 0.00 | 0.01 | 0.389 | 0.0000 |
| d__Bacteria.p__Firmicutes_A.c__Clostridia_258483.o__Lachnospirales.f__Lachnospiraceae.g__Acetitomaculum | 0.0000 | 0.000 | 0.073 | 0.00 | 0.11 | 0.39 | 0.0000 |
| d__Bacteria.p__Proteobacteria.c__Alphaproteobacteria.o__Rhodobacterales.f__Rhodobacteraceae.__ | 0.0000 | 0.000 | 0.126 | 0.06 | 0.04 | 0.39 | 0.0000 |
| d__Bacteria.p__Cyanobacteria.c__Cyanobacteriia.o__Pseudanabaenales.f__Pseudanabaenaceae.g__Pseudanabaena | 0.0000 | 0.000 | 0.073 | 0.00 | 0.02 | 0.39 | 0.0000 |
| d__Bacteria.p__Firmicutes_D.c__Bacilli.o__Paenibacillales.f__Paenibacillaceae_367444.g__Paenibacillus_Z_364416 | 0.0000 | 0.000 | 0.073 | 0.00 | 0.02 | 0.39 | 0.0000 |
| d__Bacteria.p__Acidobacteriota.c__Vicinamibacteria.o__Vicinamibacterales.f__UBA2999.__ | 0.0000 | 0.000 | 0.073 | 0.00 | 0.02 | 0.39 | 0.0000 |
| d__Bacteria.p__Actinobacteriota.c__Thermoleophilia.o__Gaiellales.f__Gaiellaceae.g__AC.16 | 0.0000 | 0.000 | 0.073 | 0.00 | 0.02 | 0.39 | 0.0000 |
| d__Bacteria.p__Proteobacteria.c__Gammaproteobacteria.o__Burkholderiales_597433.f__SG8.39.g__SCGC.AG.212.J23 | 0.0000 | 0.000 | 0.073 | 0.00 | 0.01 | 0.39 | 0.0000 |
| d__Bacteria.p__Actinobacteriota.c__Thermoleophilia.o__Solirubrobacterales.f__Solirubrobacteraceae_405341.g__Solirubrobacter | 0.0000 | 0.000 | 0.073 | 0.00 | 0.01 | 0.39 | 0.0000 |
| d__Bacteria.p__Myxococcota_A_473307.c__Polyangia_463783.o__Polyangiales.f__Polyangiaceae.g__MWCO01 | 0.0000 | 0.000 | 0.073 | 0.00 | 0.01 | 0.39 | 0.0000 |
| d__Bacteria.p__Proteobacteria.c__Gammaproteobacteria.o__GCA.2729495.f__GCA.2729495.g__QUBU01 | 0.0000 | 0.000 | 0.073 | 0.00 | 0.01 | 0.39 | 0.0000 |

| | | | | | | | |
|---|---|---|---|---|---|---|---|
| d__Archaea.p__Thermoproteota.c__Nitrososphaeria_A.o__Nitrososphaerales.f__Nitrososphaeraceae.g__TA.21 | 0.0000 | 0.000 | 0.073 | 0.00 | 0.01 | 0.39 | 0.0000 |
| d__Bacteria.p__Proteobacteria.c__Alphaproteobacteria.o__Rhizobiales_A_504721.f__Hyphomicrobiaceae.g__AWTP1.13 | 0.0000 | 0.000 | 0.073 | 0.00 | 0.01 | 0.39 | 0.0000 |
| d__Bacteria.p__Verrucomicrobiota.c__Verrucomicrobiae.o__Pedosphaerales.f__UBA3939.__ | 0.0000 | 0.000 | 0.073 | 0.00 | 0.01 | 0.39 | 0.0000 |
| d__Bacteria.p__Bacteroidota.c__Bacteroidia.o__Chitinophagales.f__Chitinophagaceae_929731.g__Flavipsychrobacter_929086 | 0.0000 | 0.000 | 0.073 | 0.00 | 0.01 | 0.39 | 0.0000 |
| d__Bacteria.p__Proteobacteria.c__Alphaproteobacteria.o__RF32.f__CAG.239.g__RUG410 | 0.0023 | 0.005 | 0.477 | 5.34 | 19.97 | 0.391 | 0.0052 |
| d__Bacteria.p__Actinobacteriota.c__Actinomycetia.o__Actinomycetales.f__Kineosporiaceae.g__Kineosporia | 0.0000 | 0.000 | 0.073 | 0.00 | 0.01 | 0.391 | 0.0000 |
| d__Bacteria.p__Bacteroidota.c__Bacteroidia.o__Cytophagales.f__Cyclobacteriaceae_900466.g__Ohtaekwangia | 0.0000 | 0.000 | 0.073 | 0.00 | 0.01 | 0.392 | 0.0000 |
| d__Bacteria.p__Proteobacteria.c__Gammaproteobacteria.o__Xanthomonadales_616009.f__Xanthomonadaceae_616009.g__Xanthomonas_A_614442 | 0.0000 | 0.000 | 0.073 | 0.00 | 0.01 | 0.392 | 0.0000 |
| d__Bacteria.p__Bdellovibrionota_E.c__Bdellovibrionia_A_473294.o__Bdellovibrionales.f__Bdellovibrionaceae.g__Bdellovibrio | 0.0000 | 0.000 | 0.073 | 0.00 | 0.01 | 0.392 | 0.0000 |
| d__Bacteria.p__Firmicutes_D.c__Bacilli.o__RF39.f__UBA660.g__Faecenecus | 0.0000 | 0.000 | 0.073 | 0.00 | 0.07 | 0.393 | 0.0000 |
| d__Bacteria.p__Firmicutes_A.c__Clostridia_258483.o__Lachnospirales.f__Anaerotignaceae.g__Anaerotignum_189125 | 0.0000 | 0.000 | 0.073 | 0.00 | 0.01 | 0.393 | 0.0000 |
| d__Bacteria.p__Proteobacteria.c__Alphaproteobacteria.o__Acetobacterales.f__Acetobacteraceae.__ | 0.0000 | 0.000 | 0.073 | 0.00 | 0.01 | 0.394 | 0.0000 |
| d__Bacteria.p__Firmicutes_A.c__Clostridia_258483.o__Lachnospirales.f__Lachnospiraceae.g__Bariatricus | 0.0000 | 0.000 | 0.073 | 0.00 | 0.01 | 0.394 | 0.0000 |
| d__Bacteria.p__Firmicutes_D.c__Bacilli.o__ML615J.28.f__CAG.313.g__CAG.313 | 0.0000 | 0.000 | 0.073 | 0.00 | 0.09 | 0.396 | 0.0000 |
| d__Bacteria.p__Firmicutes_A.c__Clostridia_258483.o__Lachnospirales.f__Lachnospiraceae.g__14.2 | 0.0000 | 0.000 | 0.073 | 0.00 | 0.03 | 0.396 | 0.0000 |
| d__Bacteria.p__Desulfobacterota_I.c__Desulfovibrionia.o__Desulfovibrionales.__.__ | 0.0000 | 0.000 | 0.073 | 0.00 | 0.02 | 0.396 | 0.0000 |

| | | | | | | | |
|---|---|---|---|---|---|---|---|
| d__Bacteria.p__Firmicutes_A.c__Clostridia_258483.o__Oscillospirales.f__Oscillospiraceae_88309.g__Oscillibacter | 0.0000 | 0.000 | 0.073 | 0.00 | 0.02 | 0.396 | 0.0000 |
| d__Bacteria.p__Firmicutes_D.c__Bacilli.o__Lactobacillales.f__Lactobacillaceae.g__Limosilactobacillus | 0.0000 | 0.000 | 0.073 | 0.00 | 0.01 | 0.396 | 0.0000 |
| d__Bacteria.p__Firmicutes_D.c__Bacilli.o__ML615J.28.f__CAG.313.__ | 0.0000 | 0.000 | 0.073 | 0.00 | 0.01 | 0.396 | 0.0000 |
| d__Bacteria.p__Proteobacteria.c__Gammaproteobacteria.o__Burkholderiales_597439.f__Usitatibacteraceae.g__Usitatibacter | 0.0000 | 0.000 | 0.073 | 0.00 | 0.01 | 0.396 | 0.0000 |
| Unassigned.__.__.__.__.__ | 0.0000 | 0.000 | 0.073 | 0.00 | 0.01 | 0.396 | 0.0000 |
| d__Bacteria.p__Firmicutes_A.c__Clostridia_258483.o__Oscillospirales.f__Acutalibacteraceae.g__UBA5905 | 0.0000 | 0.000 | 0.073 | 0.00 | 0.01 | 0.397 | 0.0000 |
| d__Bacteria.p__Firmicutes_A.c__Clostridia_258483.o__Christensenellales.f__CAG.138.g__CAG.1024 | 0.0000 | 0.000 | 0.073 | 0.00 | 0.01 | 0.397 | 0.0000 |
| d__Bacteria.p__Firmicutes_A.c__Clostridia_258483.o__Lachnospirales.f__Lachnospiraceae.g__UBA4285 | 0.0000 | 0.000 | 0.073 | 0.00 | 0.01 | 0.397 | 0.0000 |
| d__Bacteria.p__Firmicutes_D.c__Bacilli.o__Lactobacillales.f__Enterococcaceae.__ | 0.0000 | 0.000 | 0.073 | 0.00 | 0.05 | 0.398 | 0.0000 |
| d__Bacteria.p__Planctomycetota.c__Phycisphaerae.o__Tepidisphaerales.f__Tepidisphaeraceae.g__UBA2421 | 0.0000 | 0.000 | 0.073 | 0.00 | 0.01 | 0.398 | 0.0000 |
| d__Bacteria.p__Proteobacteria.c__Gammaproteobacteria.o__Burkholderiales_592522.f__Burkholderiaceae_A_592522.g__Pseudorhodoferax | 0.0000 | 0.000 | 0.073 | 0.00 | 0.01 | 0.398 | 0.0000 |
| d__Bacteria.p__Proteobacteria.c__Alphaproteobacteria.o__Rhizobiales_A_502138.f__Devosiaceae.g__Devosia_A_501803 | 0.0000 | 0.000 | 0.073 | 0.00 | 0.01 | 0.401 | 0.0000 |
| d__Bacteria.p__Firmicutes_A.c__Clostridia_258483.o__Oscillospirales.f__QAKW01.g__Firm.08 | 0.0000 | 0.000 | 0.073 | 0.00 | 0.02 | 0.404 | 0.0000 |
| d__Bacteria.p__Firmicutes_A.c__Clostridia_258483.o__Lachnospirales.f__Lachnospiraceae.g__Enterocloster | 0.0000 | 0.000 | 0.073 | 0.00 | 0.23 | 0.405 | 0.0001 |
| d__Bacteria.p__Firmicutes_A.c__Clostridia_258483.o__Lachnospirales.f__Lachnospiraceae.g__Anaerostipes | 0.0000 | 0.000 | 0.073 | 0.00 | 0.09 | 0.405 | 0.0000 |
| d__Bacteria.p__Firmicutes_A.c__Clostridia_258483.o__Lachnospirales.f__Lachnospiraceae.g__Fimimorpha | 0.0000 | 0.000 | 0.073 | 0.00 | 0.09 | 0.405 | 0.0000 |
| d__Bacteria.p__Firmicutes_D.c__Bacilli.o__Erysipelotrichales.f__Coprobacillaceae.g__Erysipelatoclostridium | 0.0000 | 0.000 | 0.127 | 0.04 | 0.04 | 0.405 | 0.0000 |
| d__Bacteria.p__Bacteroidota.c__Bacteroidia.o__Chitinophagales.f__Chitin | 0.0000 | 0.000 | 0.111 | 0.03 | 0.03 | 0.405 | 0.0000 |

| | | | | | | | |
|---|---|---|---|---|---|---|---|
| ophagaceae_929731.g__Edaphobaculum | | | | | | | |
| d__Bacteria.p__Firmicutes_A.c__Clostridia_258483.o__Lachnospirales.f__Lachnospiraceae.g__Scatomonas | 0.0000 | 0.000 | 0.114 | 0.03 | 0.03 | 0.405 | 0.0000 |
| d__Bacteria.p__Proteobacteria.c__Alphaproteobacteria.o__Caulobacterales.f__Caulobacteraceae.__ | 0.0000 | 0.000 | 0.073 | 0.00 | 0.04 | 0.405 | 0.0000 |
| d__Bacteria.p__Firmicutes_A.c__Clostridia_258483.o__Oscillospirales.f__Ruminococcaceae.g__Ruthenibacterium | 0.0000 | 0.000 | 0.073 | 0.00 | 0.04 | 0.405 | 0.0000 |
| d__Bacteria.p__Firmicutes_A.c__Clostridia_258483.o__Lachnospirales.f__Lachnospiraceae.g__Hungatella_A_128155 | 0.0000 | 0.000 | 0.073 | 0.00 | 0.03 | 0.405 | 0.0000 |
| d__Bacteria.p__Firmicutes_A.c__Clostridia_258483.o__Oscillospirales.f__UBA929.g__WRAI01 | 0.0000 | 0.000 | 0.073 | 0.00 | 0.02 | 0.405 | 0.0000 |
| d__Bacteria.p__Firmicutes_A.c__Clostridia_258483.o__Oscillospirales.f__Oscillospiraceae_88309.g__Intestinimonas | 0.0000 | 0.000 | 0.073 | 0.00 | 0.02 | 0.405 | 0.0000 |
| d__Bacteria.p__Desulfobacterota_I.c__Desulfovibrionia.o__Desulfovibrionales.f__Desulfovibrionaceae.g__Bilophila | 0.0000 | 0.000 | 0.073 | 0.00 | 0.01 | 0.405 | 0.0000 |
| d__Bacteria.p__Proteobacteria.c__Gammaproteobacteria.o__Enterobacterales_A_737866.f__Enterobacteriaceae_A.g__Morganella | 0.0000 | 0.000 | 0.073 | 0.00 | 0.01 | 0.408 | 0.0000 |
| d__Bacteria.p__Proteobacteria.c__Alphaproteobacteria.o__Rhizobiales_A_504705.f__Xanthobacteraceae_503485.g__VAZQ01 | 0.0000 | 0.000 | 0.073 | 0.00 | 0.05 | 0.41 | 0.0000 |
| d__Bacteria.p__Acidobacteriota.c__Acidobacteriae.o__20CM.2.55.15.f__20CM.2.55.15.g__DSPE01 | 0.0000 | 0.000 | 0.073 | 0.00 | 0.02 | 0.41 | 0.0000 |
| d__Bacteria.p__Verrucomicrobiota.c__Verrucomicrobiae.o__Pedosphaerales.__.__ | 0.0000 | 0.000 | 0.073 | 0.00 | 0.01 | 0.41 | 0.0000 |
| d__Bacteria.p__Desulfobacterota_B.c__Binatia.o__UTPRO1.f__DP.6.g__DP.6 | 0.0000 | 0.000 | 0.073 | 0.00 | 0.01 | 0.41 | 0.0000 |
| d__Bacteria.p__Synergistota.c__Synergistia.o__Synergistales.__.__ | 0.0000 | 0.000 | 0.073 | 0.00 | 0.01 | 0.412 | 0.0000 |
| d__Bacteria.p__Actinobacteriota.c__Actinomycetia.o__Actinomycetales.f__Dermabacteraceae.g__Brachybacterium | 0.0000 | 0.000 | 0.073 | 0.00 | 0.02 | 0.413 | 0.0000 |
| d__Bacteria.p__Firmicutes_A.c__Clostridia_258483.o__Peptostreptococcales.f__Anaerovoracaceae.g__Mogibacterium | 0.0001 | 0.000 | 0.343 | 0.55 | 0.49 | 0.416 | 0.0002 |

| | | | | | | | |
|---|---|---|---|---|---|---|---|
| d__Bacteria.p__Actinobacteriota.c__Actinomycetia.o__Mycobacteriales.f__Mycobacteriaceae.g__Mycobacterium | 0.0000 | 0.000 | 0.073 | 0.00 | 0.01 | 0.419 | 0.0000 |
| d__Bacteria.p__Proteobacteria.c__Alphaproteobacteria.o__Rhizobiales_A_501396.f__Rhizobiaceae_A_499470.g__Mesorhizobium_F_498388 | 0.0000 | 0.000 | 0.073 | 0.00 | 0.01 | 0.419 | 0.0000 |
| d__Bacteria.p__Firmicutes_A.c__Clostridia_258483.o__Oscillospirales.f__Oscillospiraceae_88309.g__WRMH01 | 0.0000 | 0.000 | 0.073 | 0.00 | 0.01 | 0.42 | 0.0000 |
| d__Bacteria.p__Firmicutes_A.c__Clostridia_258483.o__Christensenellales.f__Borkfalkiaceae.g__Borkfalkia | 0.0117 | 0.013 | 0.890 | 112.36 | 105.30 | 0.422 | 0.0263 |
| d__Bacteria.p__Desulfobacterota_B.c__Binatia.o__UBA9968.f__UBA9968.g__WHTF01 | 0.0000 | 0.000 | 0.073 | 0.00 | 0.02 | 0.422 | 0.0000 |
| d__Bacteria.p__Planctomycetota.c__Planctomycetia.o__Pirellulales.f__PALSA.1355.g__PALSA.1355 | 0.0000 | 0.000 | 0.073 | 0.00 | 0.01 | 0.422 | 0.0000 |
| d__Bacteria.p__Planctomycetota.c__Planctomycetia.o__Gemmatales.f__Gemmataceae.g__ | 0.0000 | 0.000 | 0.073 | 0.00 | 0.01 | 0.422 | 0.0000 |
| d__Bacteria.p__Actinobacteriota.c__Actinomycetia.o__Nitriliruptorales.f__Nitriliruptoraceae.g__CSBr16.57R1 | 0.0000 | 0.000 | 0.073 | 0.00 | 0.01 | 0.422 | 0.0000 |
| d__Bacteria.p__Eisenbacteria.c__RBG.16.71.46.o__RBG.16.71.46.f__RBG.16.71.46.g__RBG.16.71.46 | 0.0000 | 0.000 | 0.073 | 0.00 | 0.01 | 0.422 | 0.0000 |
| d__Bacteria.p__Proteobacteria.c__Alphaproteobacteria.o__Geminicoccales.f__Geminicoccaceae.__ | 0.0000 | 0.000 | 0.073 | 0.00 | 0.01 | 0.422 | 0.0000 |
| d__Bacteria.p__Bdellovibrionota_E.c__Bacteriovoracia.o__Bacteriovoracales.f__Peredibacteraceae.g__Peredibacter | 0.0000 | 0.000 | 0.073 | 0.00 | 0.01 | 0.422 | 0.0000 |
| d__Bacteria.p__Actinobacteriota.c__Thermoleophilia.o__Gaiellales.f__Gaiellaceae.g__Palsa.739 | 0.0000 | 0.000 | 0.073 | 0.00 | 0.02 | 0.424 | 0.0000 |
| d__Bacteria.p__Actinobacteriota.c__Actinomycetia.o__Mycobacteriales.f__Micromonosporaceae.g__Micromonospora_H_372070 | 0.0000 | 0.000 | 0.073 | 0.00 | 0.02 | 0.424 | 0.0000 |
| d__Bacteria.p__Actinobacteriota.c__Actinomycetia.o__Mycobacteriales.f__Pseudonocardiaceae.__ | 0.0000 | 0.000 | 0.073 | 0.00 | 0.01 | 0.424 | 0.0000 |
| d__Bacteria.p__Verrucomicrobiota.c__Verrucomicrobiae.o__Chthoniobacterales.f__Chthoniobacteraceae.g__VFJQ01 | 0.0000 | 0.000 | 0.073 | 0.00 | 0.01 | 0.424 | 0.0000 |
| d__Bacteria.p__Firmicutes_D.c__Bacilli.o__Paenibacillales.f__NBRC.103111.g__Paenibacillus_G_363720 | 0.0000 | 0.000 | 0.073 | 0.00 | 0.01 | 0.424 | 0.0000 |

| | | | | | | | |
|---|---|---|---|---|---|---|---|
| d__Bacteria.p__Actinobacteriota.c__Acidimicrobiia_401430.o__Acidimicrobiales.f__JACDCH01.g__VFJN01 | 0.0000 | 0.000 | 0.073 | 0.00 | 0.01 | 0.424 | 0.0000 |
| d__Bacteria.p__Acidobacteriota.c__Vicinamibacteria.o__Vicinamibacterales.f__SCN.69.37.g__SCN.69.37 | 0.0000 | 0.000 | 0.073 | 0.00 | 0.01 | 0.424 | 0.0000 |
| d__Bacteria.p__Bacteroidota.c__Bacteroidia.o__Chitinophagales.f__Chitinophagaceae_966727.g__Flavitalea_936101 | 0.0000 | 0.000 | 0.073 | 0.00 | 0.01 | 0.424 | 0.0000 |
| d__Bacteria.p__Planctomycetota.c__Planctomycetia.o__Pirellulales.f__Pirellulaceae.__ | 0.0000 | 0.000 | 0.073 | 0.00 | 0.01 | 0.424 | 0.0000 |
| d__Bacteria.p__Desulfobacterota_B.c__Binatia.o__Bin18.f__Bin18.g__JABFSC01 | 0.0000 | 0.000 | 0.073 | 0.00 | 0.01 | 0.424 | 0.0000 |
| d__Bacteria.p__Bacteroidota.c__Bacteroidia.o__Bacteroidales.f__Coprobacteraceae.g__Coprobacter | 0.0000 | 0.000 | 0.166 | 0.07 | 0.07 | 0.432 | 0.0000 |
| d__Bacteria.p__Firmicutes_A.c__Clostridia_258483.o__TANB77.f__CAG.508.g__UMGS1994 | 0.0069 | 0.008 | 0.912 | 51.56 | 73.44 | 0.441 | 0.0154 |
| d__Bacteria.p__Firmicutes_A.c__Clostridia_258483.o__TANB77.f__CAG.508.g__Merdicola | 0.0000 | 0.000 | 0.130 | 0.02 | 0.02 | 0.448 | 0.0000 |
| d__Bacteria.p__Cyanobacteria.c__Vampirovibrionia.o__Gastranaerophilales.f__Gastranaerophilaceae.g__QHMH01 | 0.0000 | 0.000 | 0.218 | 0.08 | 0.08 | 0.453 | 0.0000 |
| d__Bacteria.p__Firmicutes_A.c__Clostridia_258483.o__Oscillospirales.f__Oscillospiraceae_88309.g__CAG.83 | 0.0000 | 0.000 | 0.269 | 0.11 | 0.11 | 0.462 | 0.0000 |
| d__Bacteria.p__Firmicutes_D.c__Bacilli.o__Erysipelotrichales.f__Erysipelotrichaceae.__ | 0.0000 | 0.000 | 0.135 | 0.03 | 0.03 | 0.489 | 0.0000 |
| d__Bacteria.p__Bacteroidota.c__Bacteroidia.o__Bacteroidales.f__Bacteroidaceae.g__Bacteroides_H | 0.0001 | 0.000 | 0.299 | 0.61 | 0.88 | 0.496 | 0.0003 |
| d__Bacteria.p__Firmicutes_A.c__Clostridia_258483.o__Clostridiales.f__Clostridiaceae_222000.g__Clostridium_P | 0.0006 | 0.002 | 0.346 | 3.45 | 3.58 | 0.502 | 0.0014 |
| d__Bacteria.p__Proteobacteria.c__Alphaproteobacteria.o__Rhizobiales_A_504705.f__Xanthobacteraceae_503485.__ | 0.0000 | 0.000 | 0.120 | 0.02 | 0.04 | 0.512 | 0.0000 |
| d__Bacteria.p__Firmicutes_A.c__Clostridia_258483.o__Oscillospirales.f__Oscillospiraceae_88309.g__Limivicinus | 0.0028 | 0.003 | 1.029 | 25.79 | 43.88 | 0.525 | 0.0064 |
| d__Bacteria.p__Firmicutes_A.c__Clostridia_258483.o__Lachnospirales.f__Lachnospiraceae.g__Bacteroides_F | 0.0002 | 0.001 | 0.326 | 1.28 | 1.29 | 0.525 | 0.0005 |

| | | | | | | | |
|---|---|---|---|---|---|---|---|
| d__Bacteria.p__Actinobacteriota.c__Coriobacteriia.o__Coriobacteriales.f__Eggerthellaceae.g__CACXMZ01 | 0.0000 | 0.000 | 0.151 | 0.04 | 0.05 | 0.525 | 0.0000 |
| d__Bacteria.p__Firmicutes_A.c__Clostridia_258483.o__Lachnospirales.__.__ | 0.0001 | 0.000 | 0.332 | 0.56 | 0.60 | 0.532 | 0.0002 |
| d__Bacteria.p__Spirochaetota.c__Brachyspirae.o__Brachyspirales.f__Brachyspiraceae.g__Brachyspira | 0.0001 | 0.000 | 0.618 | 0.54 | 0.59 | 0.54 | 0.0002 |
| d__Bacteria.p__Firmicutes_A.c__Clostridia_258483.o__Lachnospirales.f__Lachnospiraceae.g__1XD8.76 | 0.0001 | 0.001 | 0.216 | 0.63 | 0.86 | 0.543 | 0.0003 |
| d__Bacteria.p__Firmicutes_D.c__Bacilli.o__Erysipelotrichales.f__Erysipelotrichaceae.g__UBA636 | 0.0001 | 0.000 | 0.368 | 0.34 | 0.38 | 0.567 | 0.0001 |
| d__Bacteria.p__Firmicutes_D.c__Bacilli.o__RF39.f__UBA660.g__Faecisoma | 0.0000 | 0.000 | 0.177 | 0.10 | 0.14 | 0.57 | 0.0001 |
| d__Bacteria.p__Firmicutes_A.c__Clostridia_258483.o__Oscillospirales.f__Ruminococcaceae.g__ | 0.0000 | 0.000 | 0.173 | 0.03 | 0.04 | 0.57 | 0.0000 |
| d__Bacteria.p__Firmicutes_A.c__Clostridia_258483.o__Christensenellales.f__Borkfalkiaceae.g__UBA1259 | 0.0004 | 0.001 | 0.731 | 2.38 | 2.41 | 0.578 | 0.0008 |
| d__Bacteria.p__Campylobacterota.c__Campylobacteria.o__Campylobacterales.f__Helicobacteraceae.__ | 0.0006 | 0.001 | 0.870 | 5.71 | 5.91 | 0.58 | 0.0014 |
| d__Bacteria.p__Firmicutes_D.c__Bacilli.o__RF39.f__UBA660.g__CAG.605 | 0.0024 | 0.004 | 0.672 | 9.72 | 21.04 | 0.592 | 0.0053 |
| d__Bacteria.p__Bacteroidota.c__Bacteroidia.o__Bacteroidales.f__Bacteroidaceae.g__Paraprevotella | 0.0000 | 0.000 | 0.112 | 0.02 | 0.07 | 0.597 | 0.0000 |
| d__Bacteria.p__Actinobacteriota.c__Coriobacteriia.o__Coriobacteriales.f__Coriobacteriaceae.g__Collinsella | 0.0001 | 0.000 | 0.585 | 0.56 | 0.68 | 0.601 | 0.0002 |
| d__Bacteria.p__Planctomycetota.c__Planctomycetia.o__Pirellulales.__.__ | 0.0000 | 0.000 | 0.098 | 0.00 | 0.02 | 0.604 | 0.0000 |
| d__Bacteria.p__Firmicutes_A.c__Clostridia_258483.o__Christensenellales.f__CAG.74.g__SFMI01 | 0.0004 | 0.000 | 0.866 | 2.92 | 2.95 | 0.605 | 0.0009 |
| d__Bacteria.p__Bacteroidota.c__Bacteroidia.o__Bacteroidales.f__Bacteroidaceae.g__UBA4334 | 0.0000 | 0.000 | 0.095 | 0.00 | 0.04 | 0.607 | 0.0000 |
| d__Bacteria.p__Actinobacteriota.c__Coriobacteriia.o__Coriobacteriales.f__Eggerthellaceae.g__UBA5808 | 0.0000 | 0.000 | 0.104 | 0.00 | 0.01 | 0.609 | 0.0000 |
| d__Bacteria.p__Proteobacteria.c__Gammaproteobacteria.o__Burkholderiales_595427.f__Burkholderiaceae_A_595427.g__Parasutterella | 0.0000 | 0.000 | 0.086 | 0.00 | 0.03 | 0.61 | 0.0000 |
| d__Bacteria.p__Bacteroidota.c__Bacteroidia.o__Bacteroidales.f__Rikenellaceae.g__Alistipes_A_871404 | 0.0000 | 0.000 | 0.093 | 0.00 | 0.02 | 0.61 | 0.0000 |

| | | | | | | | |
|---|---|---|---|---|---|---|---|
| d__Bacteria.p__Cyanobacteria.c__Vampirovibrionia.o__Gastranaerophilales.__.__ | 0.0000 | 0.000 | 0.095 | 0.00 | 0.04 | 0.614 | 0.0000 |
| d__Bacteria.p__Firmicutes_A.c__Clostridia_258483.o__Lachnospirales.f__Lachnospiraceae.g__Marvinbryantia | 0.0028 | 0.003 | 0.953 | 38.95 | 38.73 | 0.616 | 0.0063 |
| d__Bacteria.p__Firmicutes_A.c__Clostridia_258483.o__Christensenellales.f__CAG.138.__ | 0.0000 | 0.000 | 0.098 | 0.00 | 0.02 | 0.617 | 0.0000 |
| d__Bacteria.p__Firmicutes_A.c__Clostridia_258483.o__Lachnospirales.f__Lachnospiraceae.g__UBA4292 | 0.0057 | 0.009 | 0.626 | 32.62 | 38.27 | 0.618 | 0.0128 |
| d__Bacteria.p__Firmicutes_A.c__Clostridia_258483.o__Clostridiales.f__Clostridiaceae_222000.g__Clostridium_T | 0.0000 | 0.000 | 0.101 | 0.00 | 0.12 | 0.619 | 0.0000 |
| d__Bacteria.p__Firmicutes_D.c__Bacilli.o__RF39.f__UBA660.g__Onthocola_B | 0.0003 | 0.000 | 0.736 | 1.68 | 1.75 | 0.621 | 0.0006 |
| d__Bacteria.p__Firmicutes_A.c__Clostridia_258483.o__Christensenellales.f__Borkfalkiaceae.__ | 0.0000 | 0.000 | 0.081 | 0.00 | 0.05 | 0.625 | 0.0000 |
| d__Bacteria.p__Firmicutes_D.c__Bacilli.o__RFN20.f__CAG.826.g__UBA3207 | 0.0000 | 0.000 | 0.103 | 0.00 | 0.05 | 0.625 | 0.0000 |
| d__Bacteria.p__Firmicutes_A.c__Clostridia_258483.o__Christensenellales.f__CAG.552.g__WRAP01 | 0.0000 | 0.000 | 0.104 | 0.00 | 0.01 | 0.625 | 0.0000 |
| d__Bacteria.p__Firmicutes_D.c__Bacilli.o__Acholeplasmatales.f__Anaeroplasmataceae.__ | 0.0000 | 0.000 | 0.094 | 0.00 | 0.20 | 0.626 | 0.0000 |
| d__Bacteria.p__Firmicutes_A.c__Clostridia_258483.o__Oscillospirales.f__Oscillospiraceae_88309.g__Enterenecus | 0.0010 | 0.001 | 0.975 | 8.24 | 11.48 | 0.632 | 0.0022 |
| d__Bacteria.p__Firmicutes_D.c__Bacilli.o__RF39.f__UBA660.__ | 0.0014 | 0.002 | 0.771 | 5.31 | 14.65 | 0.634 | 0.0030 |
| d__Bacteria.p__Firmicutes_A.c__Clostridia_258483.o__Oscillospirales.f__UBA644.g__UBA644 | 0.0000 | 0.000 | 0.104 | 0.00 | 0.01 | 0.637 | 0.0000 |
| d__Bacteria.p__Actinobacteriota.c__Actinomycetia.o__Actinomycetales.f__Brevibacteriaceae.g__Brevibacterium | 0.0000 | 0.000 | 0.095 | 0.00 | 0.13 | 0.638 | 0.0000 |
| d__Bacteria.p__Firmicutes_A.c__Clostridia_258483.o__Lachnospirales.f__Lachnospiraceae.g__Roseburia | 0.0012 | 0.002 | 0.665 | 7.96 | 8.42 | 0.639 | 0.0028 |
| d__Bacteria.p__Acidobacteriota.c__Vicinamibacteria.o__Vicinamibacterales.f__UBA2999.g__ | 0.0000 | 0.000 | 0.104 | 0.02 | 0.09 | 0.639 | 0.0000 |
| d__Bacteria.p__Firmicutes_D.c__Bacilli.o__Bacillales_B_302584.f__DSM.18226_301387.g__Bacillus_BD | 0.0000 | 0.000 | 0.093 | 0.00 | 0.06 | 0.641 | 0.0000 |

| | | | | | | | |
|---|---|---|---|---|---|---|---|
| d__Bacteria.p__Firmicutes_A.c__Clostridia_258483.o__Oscillospirales.f__Acutalibacteraceae.g__Solibaculum | 0.0000 | 0.000 | 0.084 | 0.00 | 0.04 | 0.641 | 0.0000 |
| d__Bacteria.p__Firmicutes_A.c__Clostridia_258483.o__Christensenellales.__.__ | 0.0001 | 0.000 | 0.280 | 0.51 | 0.67 | 0.642 | 0.0003 |
| d__Bacteria.p__Firmicutes_A.c__Clostridia_258483.o__Lachnospirales.f__Lachnospiraceae.g__Lachnospira | 0.0000 | 0.000 | 0.087 | 0.00 | 0.27 | 0.643 | 0.0001 |
| d__Bacteria.p__Firmicutes_C.c__Negativicutes.o__Veillonellales.f__Dialisteraceae.g__UBA1822 | 0.0000 | 0.000 | 0.132 | 0.01 | 0.02 | 0.643 | 0.0000 |
| d__Bacteria.p__Bacteroidota.c__Bacteroidia.o__Bacteroidales.f__Marinifilaceae.g__Butyricimonas | 0.0000 | 0.000 | 0.098 | 0.00 | 0.11 | 0.645 | 0.0000 |
| d__Bacteria.p__Firmicutes_D.c__Bacilli.o__Staphylococcales.f__Staphylococcaceae.__ | 0.0000 | 0.000 | 0.104 | 0.00 | 0.10 | 0.646 | 0.0000 |
| d__Bacteria.p__Firmicutes_A.c__Clostridia_258483.o__Lachnospirales.f__Lachnospiraceae.g__CAG.632 | 0.0047 | 0.008 | 0.588 | 25.96 | 37.43 | 0.647 | 0.0106 |
| d__Bacteria.p__Campylobacterota.c__Campylobacteria.o__Campylobacterales.f__Helicobacteraceae.g__Helicobacter_F | 0.0000 | 0.000 | 0.154 | 0.02 | 0.04 | 0.647 | 0.0000 |
| d__Bacteria.p__Proteobacteria.c__Alphaproteobacteria.o__Geminicoccales.f__Geminicoccaceae.g__HRBIN40 | 0.0000 | 0.000 | 0.102 | 0.00 | 0.03 | 0.651 | 0.0000 |
| d__Bacteria.p__Synergistota.c__Synergistia.o__Synergistales.f__Aminobacteriaceae.__ | 0.0000 | 0.000 | 0.098 | 0.00 | 0.02 | 0.656 | 0.0000 |
| d__Bacteria.p__Firmicutes_A.c__Clostridia_258483.o__TANB77.f__CAG.508.g__CAG.273 | 0.0037 | 0.005 | 0.816 | 12.51 | 39.96 | 0.666 | 0.0082 |
| d__Bacteria.p__Proteobacteria.c__Gammaproteobacteria.__.__.__ | 0.0001 | 0.000 | 0.351 | 0.24 | 0.30 | 0.669 | 0.0001 |
| d__Bacteria.p__Proteobacteria.c__Gammaproteobacteria.o__Enterobacterales_A_737866.f__Succinivibrionaceae.g__Succinivibrio | 0.0000 | 0.000 | 0.161 | 0.06 | 0.12 | 0.669 | 0.0000 |
| d__Bacteria.p__Bacteroidota.c__Bacteroidia.o__Bacteroidales.f__Paludibacteraceae.g__F0058 | 0.0003 | 0.001 | 0.301 | 1.24 | 1.72 | 0.685 | 0.0006 |
| d__Bacteria.p__Firmicutes_A.c__Clostridia_258483.o__Peptostreptococcales.f__Anaerovoracaceae.g__Eubacterium_T | 0.0000 | 0.000 | 0.149 | 0.01 | 0.03 | 0.691 | 0.0000 |
| d__Bacteria.p__Cyanobacteria.c__Vampirovibrionia.o__Gastranaerophilales.f__Gastranaerophilaceae.__ | 0.0056 | 0.006 | 0.927 | 40.26 | 58.88 | 0.698 | 0.0126 |
| d__Bacteria.p__Campylobacterota.c__Campylobacteria.o__Campylobacterales.f__Campylobacteraceae.__ | 0.0000 | 0.000 | 0.119 | 0.01 | 0.05 | 0.704 | 0.0000 |

| | | | | | | | |
|---|---|---|---|---|---|---|---|
| d__Bacteria.p__Firmicutes_A.c__Clostridia_258483.o__Lachnospirales.f__Lachnospiraceae.g__UBA1258 | 0.0000 | 0.000 | 0.175 | 0.03 | 0.06 | 0.712 | 0.0000 |
| d__Bacteria.p__Firmicutes_C.c__Negativicutes.o__Veillonellales.f__Dialisteraceae.__ | 0.0000 | 0.000 | 0.165 | 0.01 | 0.03 | 0.715 | 0.0000 |
| d__Bacteria.p__Verrucomicrobiota.c__Verrucomicrobiae.o__Verrucomicrobiales.f__Akkermansiaceae.g__Akkermansia | 0.0011 | 0.002 | 0.599 | 6.71 | 7.48 | 0.722 | 0.0025 |
| d__Bacteria.p__Cyanobacteria.c__Vampirovibrionia.o__Gastranaerophilales.f__Gastranaerophilaceae.g__ | 0.0008 | 0.001 | 0.638 | 1.95 | 6.61 | 0.723 | 0.0017 |
| d__Bacteria.p__Firmicutes_D.c__Bacilli.__.__.__ | 0.0018 | 0.003 | 0.659 | 9.16 | 13.24 | 0.729 | 0.0040 |
| d__Bacteria.p__Firmicutes_A.c__Clostridia_258483.o__Oscillospirales.f__Butyricicoccaceae.g__Butyricicoccus_A_77030 | 0.0000 | 0.000 | 0.233 | 0.07 | 0.13 | 0.737 | 0.0000 |
| d__Bacteria.p__Firmicutes_A.c__Clostridia_258483.o__Lachnospirales.f__Lachnospiraceae.g__C.53 | 0.0020 | 0.003 | 0.799 | 11.64 | 17.84 | 0.74 | 0.0045 |
| d__Bacteria.p__Firmicutes_A.c__Clostridia_258483.o__.f__.g__ | 0.0020 | 0.003 | 0.811 | 11.29 | 22.94 | 0.744 | 0.0046 |
| d__Bacteria.p__Firmicutes_A.c__Clostridia_258483.o__Oscillospirales.f__Acutalibacteraceae.g__CAG.964 | 0.0000 | 0.000 | 0.171 | 0.03 | 0.06 | 0.745 | 0.0000 |
| d__Bacteria.p__Synergistota.c__Synergistia.o__Synergistales.f__Aminobacteriaceae.g__Fretibacterium | 0.0003 | 0.000 | 0.637 | 1.64 | 2.18 | 0.748 | 0.0007 |
| d__Bacteria.p__Patescibacteria.c__Saccharimonadia.o__Saccharimonadales.f__Nanoperiomorbaceae.g__Nanoperiomorbus | 0.0000 | 0.000 | 0.123 | 0.00 | 0.05 | 0.749 | 0.0000 |
| d__Bacteria.p__Firmicutes_C.c__Negativicutes.o__Acidaminococcales.f__Acidaminococcaceae.__ | 0.0000 | 0.000 | 0.155 | 0.02 | 0.07 | 0.752 | 0.0000 |
| d__Bacteria.p__Firmicutes_A.c__Clostridia_258483.o__Christensenellales.f__CAG.74.g__Onthenecus | 0.0000 | 0.000 | 0.100 | 0.00 | 0.10 | 0.753 | 0.0000 |
| d__Bacteria.p__Firmicutes_A.c__Clostridia_258483.o__TANB77.f__CAG.508.g__CAG.269 | 0.0053 | 0.005 | 1.074 | 61.55 | 66.64 | 0.754 | 0.0119 |
| d__Bacteria.p__Firmicutes_D.c__Bacilli.o__RF39.f__UBA660.g__ | 0.0019 | 0.002 | 0.911 | 11.14 | 20.48 | 0.756 | 0.0043 |
| d__Bacteria.p__Firmicutes_A.c__Clostridia_258483.o__Oscillospirales.f__Acutalibacteraceae.g__Hydrogeniiclostridium | 0.0000 | 0.000 | 0.107 | 0.00 | 0.04 | 0.756 | 0.0000 |
| d__Bacteria.p__Actinobacteriota.c__Coriobacteriia.o__Coriobacteriales.f__Eggerthellaceae.__ | 0.0026 | 0.003 | 0.828 | 25.98 | 35.08 | 0.759 | 0.0058 |
| d__Bacteria.p__Firmicutes_D.c__Bacilli.o__RF39.f__UBA660.g__JAAYOI01 | 0.0000 | 0.000 | 0.118 | 0.00 | 0.03 | 0.76 | 0.0000 |

| | | | | | | | |
|---|---|---|---|---|---|---|---|
| d__Bacteria.p__Firmicutes_A.c__Clostridia_258483.o__Lachnospirales.f__Lachnospiraceae.g__Oribacterium | 0.0001 | 0.000 | 0.354 | 0.54 | 0.76 | 0.762 | 0.0003 |
| d__Bacteria.p__Bacteroidota.c__Bacteroidia.o__Bacteroidales.f__Rikenellaceae.g__Alistipes_A_871400 | 0.0000 | 0.000 | 0.081 | 0.00 | 0.20 | 0.764 | 0.0000 |
| d__Bacteria.p__Proteobacteria.c__Alphaproteobacteria.__.__.__ | 0.0000 | 0.000 | 0.139 | 0.01 | 0.04 | 0.765 | 0.0000 |
| d__Bacteria.p__Bacteroidota.c__Bacteroidia.o__Bacteroidales.f__Bacteroidaceae.g__Phocaeicola_A_858004 | 0.0000 | 0.000 | 0.117 | 0.00 | 0.26 | 0.767 | 0.0001 |
| d__Bacteria.p__Firmicutes_A.c__Clostridia_258483.o__Lachnospirales.f__Lachnospiraceae.g__Eubacterium_G | 0.0000 | 0.000 | 0.108 | 0.00 | 0.05 | 0.773 | 0.0000 |
| d__Bacteria.p__Proteobacteria.c__Alphaproteobacteria.o__UBA3830.f__UBA3830.g__UBA3830 | 0.0000 | 0.000 | 0.183 | 0.03 | 0.07 | 0.776 | 0.0000 |
| d__Bacteria.p__Firmicutes_A.c__Clostridia_258483.o__Lachnospirales.f__Anaerotignaceae.__ | 0.0000 | 0.000 | 0.118 | 0.00 | 0.03 | 0.777 | 0.0000 |
| d__Bacteria.p__Actinobacteriota.c__Coriobacteriia.o__Coriobacteriales.__.__ | 0.0005 | 0.000 | 0.938 | 3.19 | 3.54 | 0.786 | 0.0010 |
| d__Bacteria.p__Firmicutes_A.c__Clostridia_258483.o__Oscillospirales.f__Acutalibacteraceae.g__UBA737 | 0.0006 | 0.001 | 0.767 | 2.13 | 5.30 | 0.801 | 0.0014 |
| d__Bacteria.p__Cyanobacteria.c__Vampirovibrionia.o__Gastranaerophilales.f__Gastranaerophilaceae.g__Stercorousia | 0.0002 | 0.000 | 0.472 | 0.61 | 1.23 | 0.81 | 0.0004 |
| d__Bacteria.p__Campylobacterota.c__Campylobacteria.o__Campylobacterales.f__Campylobacteraceae.g__Campylobacter_D | 0.0001 | 0.000 | 0.362 | 0.58 | 0.89 | 0.818 | 0.0003 |
| d__Bacteria.p__Firmicutes_D.c__Bacilli.o__Erysipelotrichales.f__Coprobacillaceae.g__Catenibacterium | 0.0005 | 0.001 | 0.586 | 2.09 | 3.26 | 0.821 | 0.0010 |
| d__Bacteria.p__Firmicutes_A.c__Clostridia_258483.o__Peptostreptococcales.f__Anaerovoracaceae.g__Copromorpha | 0.0002 | 0.000 | 0.681 | 1.05 | 1.27 | 0.828 | 0.0004 |
| d__Bacteria.p__Actinobacteriota.c__Coriobacteriia.o__Coriobacteriales.f__Coriobacteriaceae.__ | 0.0001 | 0.000 | 0.343 | 0.24 | 0.38 | 0.829 | 0.0001 |
| d__Bacteria.p__Firmicutes_A.c__Clostridia_258483.o__Christensenellales.f__.g__ | 0.0000 | 0.000 | 0.154 | 0.02 | 0.10 | 0.833 | 0.0000 |
| d__Bacteria.p__Cyanobacteria.c__Vampirovibrionia.o__Gastranaerophilales.f__Gastranaerophilaceae.g__UBA6984 | 0.0002 | 0.001 | 0.360 | 0.85 | 1.43 | 0.836 | 0.0005 |
| d__Bacteria.p__Firmicutes_D.c__Bacilli.o__RF39.f__UBA660.g__UBA2730 | 0.0002 | 0.001 | 0.375 | 0.77 | 1.51 | 0.841 | 0.0005 |

| | | | | | | | |
|---|---|---|---|---|---|---|---|
| d__Bacteria.p__Bacteroidota.c__Bacteroidia.o__Bacteroidales.f__UBA932.g__Cryptobacteroides | 0.0077 | 0.013 | 0.617 | 38.66 | 57.47 | 0.848 | 0.0173 |
| d__Bacteria.p__Verrucomicrobiota.c__Lentisphaeria.o__Victivallales.f__Victivallaceae.g__Victivallis | 0.0000 | 0.000 | 0.132 | 0.00 | 0.05 | 0.853 | 0.0000 |
| d__Bacteria.p__Synergistota.c__Synergistia.o__Synergistales.f__Synergistaceae.g__Caccocola | 0.0000 | 0.000 | 0.140 | 0.00 | 0.03 | 0.862 | 0.0000 |
| d__Bacteria.p__Firmicutes_D.c__Bacilli.o__RF39.f__UBA660.g__CAG.533 | 0.0000 | 0.000 | 0.228 | 0.12 | 0.30 | 0.864 | 0.0001 |
| d__Bacteria.p__Actinobacteriota.c__Coriobacteriia.o__Coriobacteriales.f__Eggerthellaceae.g__Slackia_A | 0.0000 | 0.000 | 0.119 | 0.00 | 0.05 | 0.865 | 0.0000 |
| d__Bacteria.p__Spirochaetota.c__Spirochaetia.o__Treponematales.f__Treponemataceae.g__Treponema_D | 0.0008 | 0.008 | 0.094 | 0.00 | 7.69 | 0.869 | 0.0017 |
| d__Bacteria.p__Firmicutes_D.c__Bacilli.o__Staphylococcales.f__Staphylococcaceae.g__Staphylococcus | 0.0001 | 0.001 | 0.124 | 0.03 | 1.27 | 0.878 | 0.0003 |
| d__Bacteria.p__Proteobacteria.c__Gammaproteobacteria.o__Burkholderiales_595427.f__Burkholderiaceae_A_595427.g__Mesosutterella | 0.0009 | 0.001 | 0.993 | 7.23 | 12.11 | 0.879 | 0.0020 |
| d__Bacteria.p__Bacteroidota.c__Bacteroidia.o__Bacteroidales.f__Muribaculaceae.g__SFTJ01 | 0.0001 | 0.000 | 0.283 | 0.40 | 0.94 | 0.882 | 0.0003 |
| d__Bacteria.p__Firmicutes_D.c__Bacilli.o__Acholeplasmatales.f__Anaeroplasmataceae.g__ | 0.0007 | 0.003 | 0.247 | 1.82 | 6.19 | 0.894 | 0.0016 |
| d__Bacteria.p__Firmicutes_C.c__Negativicutes.__.__.__ | 0.0000 | 0.000 | 0.146 | 0.00 | 0.07 | 0.898 | 0.0000 |
| d__Bacteria.p__Firmicutes_D.c__Bacilli.o__RF39.f__UBA660.g__UBA3789 | 0.0020 | 0.003 | 0.736 | 5.39 | 19.11 | 0.899 | 0.0044 |
| d__Bacteria.p__Firmicutes_A.c__Clostridia_258483.o__TANB77.f__CAG.508.__ | 0.0040 | 0.004 | 1.063 | 41.22 | 44.91 | 0.901 | 0.0088 |
| d__Bacteria.p__Firmicutes_A.c__Clostridia_258483.o__Lachnospirales.f__Lachnospiraceae.g__Lacrimispora | 0.0000 | 0.000 | 0.165 | 0.00 | 0.05 | 0.908 | 0.0000 |
| d__Bacteria.p__Firmicutes_D.c__Bacilli.o__RF39.f__UBA660.g__RUG705 | 0.0011 | 0.001 | 0.854 | 5.78 | 9.89 | 0.909 | 0.0024 |
| d__Bacteria.p__Firmicutes_A.c__Clostridia_258483.o__Oscillospirales.f__CAG.272.g__Avispirillum | 0.0000 | 0.000 | 0.193 | 0.04 | 0.17 | 0.914 | 0.0000 |
| d__Bacteria.p__Firmicutes_A.c__Clostridia_258483.o__Oscillospirales.f__Ruminococcaceae.g__CAG.353 | 0.0050 | 0.008 | 0.644 | 23.70 | 45.06 | 0.916 | 0.0111 |
| d__Bacteria.p__Firmicutes_A.c__Clostridia_258483.o__Lachnospirales.f__.g__ | 0.0004 | 0.001 | 0.681 | 1.96 | 2.71 | 0.917 | 0.0008 |

| | | | | | | | |
|---|---|---|---|---|---|---|---|
| d__Bacteria.p__Firmicutes_D.c__Bacilli.o__RF39.f__UBA660.g__RUG12438 | 0.0006 | 0.001 | 0.522 | 2.48 | 4.84 | 0.928 | 0.0014 |
| d__Bacteria.p__Patescibacteria.c__Saccharimonadia.o__Saccharimonadales.f__Nanosyncoccaceae.g__Nanosyncoccus | 0.0000 | 0.000 | 0.278 | 0.04 | 0.12 | 0.929 | 0.0000 |
| d__Bacteria.p__Firmicutes_A.c__Clostridia_258483.o__Lachnospirales.f__Lachnospiraceae.g__Blautia_A_141780 | 0.0001 | 0.001 | 0.084 | 0.02 | 0.59 | 0.93 | 0.0001 |
| d__Bacteria.p__Bacteroidota.c__Bacteroidia.o__Bacteroidales.f__Bacteroidaceae.g__Prevotella | 0.0464 | 0.042 | 1.101 | 544.99 | 708.63 | 0.939 | 0.1038 |
| d__Bacteria.p__Proteobacteria.c__Alphaproteobacteria.o__Rs.D84_512864.f__Rs.D84.g__ | 0.0000 | 0.000 | 0.255 | 0.07 | 0.24 | 0.939 | 0.0001 |
| d__Bacteria.p__Firmicutes_D.c__Bacilli.o__RF39.f__UBA660.g__CAG.302 | 0.0000 | 0.000 | 0.156 | 0.00 | 0.18 | 0.94 | 0.0000 |
| d__Bacteria.p__Firmicutes_A.c__Clostridia_258483.o__Lachnospirales.f__Lachnospiraceae.g__Schaedlerella | 0.0001 | 0.000 | 0.342 | 0.18 | 0.46 | 0.948 | 0.0001 |
| d__Bacteria.p__Firmicutes_A.c__Clostridia_258483.o__Oscillospirales.f__CAG.272.g__QALR01 | 0.0000 | 0.000 | 0.153 | 0.00 | 0.14 | 0.959 | 0.0000 |
| d__Bacteria.p__Firmicutes_A.c__Clostridia_258483.o__Oscillospirales.f__Butyricicoccaceae.g__ | 0.0046 | 0.005 | 0.899 | 50.48 | 54.39 | 0.96 | 0.0104 |
| d__Bacteria.p__Verrucomicrobiota.c__Lentisphaeria.o__Victivallales.f__UBA1829.g__UBA1732 | 0.0000 | 0.000 | 0.176 | 0.00 | 0.14 | 0.967 | 0.0000 |
| d__Bacteria.p__Proteobacteria.__.__.__.__ | 0.0000 | 0.000 | 0.358 | 0.08 | 0.23 | 0.968 | 0.0001 |
| d__Bacteria.p__Proteobacteria.c__Gammaproteobacteria.o__Burkholderiales_597441.f__Neisseriaceae_563222.g__ | 0.0000 | 0.000 | 0.178 | 0.00 | 0.16 | 0.969 | 0.0000 |
| d__Bacteria.p__Firmicutes_A.c__Clostridia_258483.o__Oscillospirales.f__CAG.272.g__RUG13077 | 0.0000 | 0.000 | 0.164 | 0.00 | 0.07 | 0.969 | 0.0000 |
| d__Bacteria.p__Firmicutes_A.c__Clostridia_258483.o__Christensenellales.f__UBA1242.g__ | 0.0013 | 0.003 | 0.427 | 4.94 | 11.42 | 0.972 | 0.0029 |
| d__Bacteria.p__Firmicutes_A.c__Clostridia_258483.o__Christensenellales.f__CAG.552.g__WRAY01 | 0.0003 | 0.001 | 0.445 | 1.18 | 2.72 | 0.972 | 0.0008 |
| d__Bacteria.p__Firmicutes_A.c__Clostridia_258483.o__Lachnospirales.f__Lachnospiraceae.g__CAG.603 | 0.0010 | 0.002 | 0.488 | 3.45 | 8.10 | 0.975 | 0.0023 |
| d__Bacteria.p__Firmicutes_A.c__Clostridia_258483.o__Lachnospirales.f__Lachnospiraceae.g__Butyribacter | 0.0017 | 0.002 | 0.865 | 9.48 | 16.16 | 0.979 | 0.0037 |

| | | | | | | | |
|---|---|---|---|---|---|---|---|
| d__Bacteria.p__Firmicutes_A.c__Clostridia_258483.o__Oscillospirales.f__CAG.382.g__SFLA01 | 0.0000 | 0.000 | 0.264 | 0.03 | 0.16 | 0.981 | 0.0000 |
| d__Bacteria.p__Firmicutes_D.c__Bacilli.o__RF39.f__UBA660.g__Scybalousia | 0.0011 | 0.002 | 0.549 | 3.25 | 9.94 | 0.982 | 0.0024 |
| d__Bacteria.p__Firmicutes_D.c__Bacilli.o__RF39.f__UBA660.g__RUG12783 | 0.0000 | 0.000 | 0.200 | 0.00 | 0.08 | 0.983 | 0.0000 |
| d__Bacteria.p__Verrucomicrobiota.c__Verrucomicrobiae.o__Opitutales.f__UBA953.g__Spyradosoma | 0.0027 | 0.007 | 0.383 | 2.76 | 24.99 | 0.984 | 0.0060 |
| d__Bacteria.p__Firmicutes_D.c__Bacilli.o__Erysipelotrichales.f__Erysipelotrichaceae.g__Holdemanella | 0.0001 | 0.000 | 0.192 | 0.00 | 0.51 | 0.985 | 0.0001 |
| d__Bacteria.p__Firmicutes_A.c__Clostridia_258483.o__Lachnospirales.f__Anaerotignaceae.g__Fimicola | 0.0001 | 0.000 | 0.469 | 0.23 | 0.54 | 0.986 | 0.0002 |
| d__Bacteria.p__Firmicutes_D.c__Bacilli.o__RF39.f__UBA660.g__UBA6985 | 0.0019 | 0.003 | 0.703 | 7.57 | 18.45 | 0.987 | 0.0043 |
| d__Bacteria.p__Firmicutes_A.c__Clostridia_258483.o__Oscillospirales.f__Acutalibacteraceae.__ | 0.0009 | 0.002 | 0.381 | 3.53 | 8.00 | 0.988 | 0.0020 |
| d__Bacteria.p__Actinobacteriota.__.__.__.__ | 0.0000 | 0.000 | 0.331 | 0.05 | 0.21 | 0.988 | 0.0001 |
| d__Bacteria.p__Firmicutes_A.c__Clostridia_258483.o__Christensenellales.f__Borkfalkiaceae.g__ | 0.0000 | 0.000 | 0.198 | 0.00 | 0.17 | 0.992 | 0.0000 |
| d__Bacteria.p__Elusimicrobiota.c__Elusimicrobia_984208.o__Elusimicrobiales.f__Elusimicrobiaceae.g__UBA1436 | 0.0000 | 0.000 | 0.204 | 0.00 | 0.14 | 0.994 | 0.0000 |
| d__Bacteria.p__Firmicutes_D.c__Bacilli.o__RFN20.__.__ | 0.0011 | 0.002 | 0.648 | 2.76 | 10.22 | 0.996 | 0.0024 |
| d__Bacteria.p__Verrucomicrobiota.c__Verrucomicrobiae.o__Opitutales.__.__ | 0.0000 | 0.000 | 0.337 | 0.08 | 0.37 | 0.997 | 0.0001 |
| d__Bacteria.p__Bacteroidota.c__Bacteroidia.o__Bacteroidales.f__Muribaculaceae.g__Sodaliphilus | 0.0036 | 0.004 | 0.841 | 16.92 | 35.85 | 0.998 | 0.0081 |
| d__Bacteria.p__Proteobacteria.c__Alphaproteobacteria.o__RUG11792.f__RUG11792.g__RUG11420 | 0.0002 | 0.000 | 0.733 | 1.00 | 1.66 | 0.998 | 0.0005 |
| d__Bacteria.p__Firmicutes_D.c__Bacilli.o__RF39.f__UBA660.g__Coprosoma | 0.0002 | 0.000 | 0.544 | 0.50 | 1.63 | 0.998 | 0.0004 |
| d__Bacteria.p__Firmicutes_D.c__Bacilli.o__RF39.f__UBA660.g__CAG.914 | 0.0002 | 0.001 | 0.460 | 0.39 | 2.22 | 0.999 | 0.0006 |
| d__Bacteria.p__Firmicutes_A.c__Clostridia_258483.o__Lachnospirales.f__Lachnospiraceae.g__CAG.194 | 0.0002 | 0.000 | 0.515 | 0.31 | 1.87 | 0.999 | 0.0005 |

| | | | | | | | |
|---|---|---|---|---|---|---|---|
| d__Bacteria.p__Firmicutes_D.c__Bacilli.o__RF39.f__UBA660.g__CAG.1000 | 0.0002 | 0.000 | 0.587 | 0.29 | 1.65 | 0.999 | 0.0004 |
| d__Bacteria.p__Cyanobacteria.c__Vampirovibrionia.o__Gastranaerophilales.f__Gastranaerophilaceae.g__Limenecus | 0.0002 | 0.000 | 0.428 | 0.30 | 1.52 | 0.999 | 0.0004 |
| d__Bacteria.p__Firmicutes_A.c__Clostridia_258483.o__Oscillospirales.f__Oscillospiraceae_88309.g__Dysosmobacter | 0.0009 | 0.001 | 0.985 | 7.00 | 10.26 | 1 | 0.0020 |
| d__Bacteria.p__Firmicutes_A.c__Clostridia_258483.o__Oscillospirales.__.__ | 0.0008 | 0.002 | 0.381 | 1.29 | 7.91 | 1 | 0.0019 |
| d__Bacteria.p__Proteobacteria.c__Alphaproteobacteria.o__RF32.f__CAG.239.__ | 0.0006 | 0.001 | 0.519 | 0.18 | 6.38 | 1 | 0.0014 |
| d__Archaea.p__Methanobacteriota_A_1229.c__Methanobacteria.o__Methanobacteriales.f__Methanobacteriaceae.g__Methanobrevibacter_A | 0.0004 | 0.001 | 0.449 | 0.87 | 4.07 | 1 | 0.0010 |
| d__Archaea.p__Methanobacteriota_A_1229.c__Methanobacteria.o__Methanobacteriales.f__Methanobacteriaceae.g__Methanosphaera | 0.0003 | 0.000 | 0.801 | 1.80 | 3.13 | 1 | 0.0008 |
| d__Bacteria.p__Firmicutes_A.c__Clostridia_258483.o__Lachnospirales.f__Lachnospiraceae.g__Eubacterium_F | 0.0003 | 0.001 | 0.557 | 0.47 | 2.95 | 1 | 0.0007 |
| d__Bacteria.p__Firmicutes_A.__.__.__.__ | 0.0003 | 0.001 | 0.356 | 0.22 | 2.38 | 1 | 0.0006 |
| d__Bacteria.p__Proteobacteria.c__Alphaproteobacteria.o__RF32.f__CAG.239.g__51.20 | 0.0002 | 0.001 | 0.300 | 0.13 | 2.14 | 1 | 0.0005 |
| d__Bacteria.p__Firmicutes_D.c__Bacilli.o__RFN20.f__CAG.288.g__Scatoplasma | 0.0002 | 0.000 | 0.370 | 0.03 | 1.77 | 1 | 0.0004 |
| d__Bacteria.p__Proteobacteria.c__Alphaproteobacteria.o__RF32.f__CAG.977.g__UBA2903 | 0.0001 | 0.000 | 0.274 | 0.07 | 1.13 | 1 | 0.0003 |
| d__Bacteria.p__Firmicutes_A.c__Clostridia_258483.o__Oscillospirales.f__Butyricicoccaceae.__ | 0.0001 | 0.000 | 0.487 | 0.08 | 0.90 | 1 | 0.0002 |
| d__Bacteria.p__Proteobacteria.c__Alphaproteobacteria.o__RF32.f__CAG.239.g__UBA1254 | 0.0001 | 0.000 | 0.221 | 0.04 | 0.87 | 1 | 0.0002 |
| d__Bacteria.p__Bacteroidota.c__Bacteroidia.o__Bacteroidales.f__Barnesiellaceae.g__Barnesiella | 0.0001 | 0.000 | 0.184 | 0.00 | 0.82 | 1 | 0.0002 |
| d__Bacteria.p__Bacteroidota.c__Bacteroidia.o__Bacteroidales.f__Tannerellaceae.g__Tannerella | 0.0001 | 0.000 | 0.387 | 0.08 | 0.68 | 1 | 0.0002 |
| d__Bacteria.p__Firmicutes_A.c__Clostridia_258483.o__Lachnospirales.f__Anaerotignaceae.g__ | 0.0000 | 0.000 | 0.332 | 0.02 | 0.48 | 1 | 0.0001 |

| | | | | | | | |
|---|---|---|---|---|---|---|---|
| d__Bacteria.p__Firmicutes_A.c__Clostridia_258483.o__Christensenellales.f__UBA1242.g__WRCD01 | 0.0000 | 0.000 | 0.272 | 0.00 | 0.39 | 1 | 0.0001 |
| d__Bacteria.p__Firmicutes_A.c__Clostridia_258483.o__Oscillospirales.f__Oscillospiraceae_88309.g__Onthomonas | 0.0000 | 0.000 | 0.233 | 0.00 | 0.37 | 1 | 0.0001 |
| d__Bacteria.p__Firmicutes_D.c__Bacilli.o__Erysipelotrichales.__.__ | 0.0000 | 0.000 | 0.310 | 0.01 | 0.21 | 1 | 0.0000 |

Overall Bray Curtis dissimilarity index between HF and LF regions at level of genus: 0.4472. Of the 391 microbial taxa classified to the level of genus (or family, when genus could not be identified), 86 (22%; taxa above bold line in table) contributed significantly ($p<0.05$) to the differences between the two regions, as quantified with the Bray-Curtis dissimilarity index (the taxa colored in pale green were not classified beyond the level of Class, and so were excluded from further analyses). Of these 86 taxa, 40 were not present at all in the LF region and were very rare in the HF region, further supporting the conclusion that the differences between the regions were in large part driven by rarer taxa. Of the remaining 46 taxa, only four were more abundant in the LF region, and 42 were more abundant in the HF region.

**Table S13. Input data for the by-fragment pairwise microbiome Bray-Curtis dissimilarity network.** In orange – top 5% most similar pairs of fragment microbiomes; in green – bottom 5% most dissimilar pairs of fragment microbiomes. No color – pairs with intermediate values not included in the dissimilarity network in Fig. 4E. Fragment 28 was excluded from this analysis as a highly dissimilar outlier; its pairwise dissimilarity values are included at the bottom of the table.

| fragment A | fragment B | Bray-Curtis dissimilarity value | fragment A | fragment B | Bray-Curtis dissimilarity value |
|---|---|---|---|---|---|
| 1 | 48 | 0.445 | 24 | 25 | 0.785 |
| 19 | 37 | 0.443 | 25 | 26 | 0.747 |
| 1 | 37 | 0.441 | 1 | 24 | 0.734 |
| 39 | 48 | 0.439 | 14 | 25 | 0.730 |
| 45 | 50 | 0.437 | 23 | 25 | 0.715 |
| 19 | 48 | 0.437 | 21 | 51 | 0.714 |
| 12 | 16 | 0.434 | 20 | 24 | 0.712 |
| 35 | 45 | 0.423 | 24 | 49 | 0.710 |
| 12 | 48 | 0.421 | 24 | 51 | 0.710 |
| 38 | 45 | 0.418 | 25 | 29 | 0.709 |
| 37 | 38 | 0.417 | 18 | 25 | 0.705 |
| 12 | 13 | 0.415 | 13 | 25 | 0.695 |
| 9 | 16 | 0.411 | 24 | 43 | 0.693 |
| 43 | 50 | 0.407 | 25 | 49 | 0.687 |
| 13 | 18 | 0.404 | 6 | 25 | 0.687 |
| 35 | 38 | 0.403 | 24 | 50 | 0.686 |
| 35 | 48 | 0.400 | 21 | 25 | 0.681 |
| 13 | 16 | 0.400 | 25 | 51 | 0.678 |
| 46 | 48 | 0.390 | 10 | 25 | 0.677 |
| 35 | 37 | 0.387 | 6 | 51 | 0.677 |
| 38 | 48 | 0.384 | 20 | 26 | 0.675 |
| 45 | 46 | 0.379 | 25 | 39 | 0.675 |
| 48 | 50 | 0.374 | 20 | 49 | 0.673 |
| 37 | 48 | 0.369 | 25 | 41 | 0.673 |
| 43 | 48 | 0.362 | 1 | 26 | 0.668 |
| 37 | 45 | 0.360 | 2 | 29 | 0.668 |
| 45 | 48 | 0.334 | 25 | 52 | 0.668 |
| **Intermediate values not included in network:** | | | | | |
| **fragment A** | **fragment B** | **Bray-Curtis dissimilarity value** | **fragment A** | **fragment B** | **Bray-Curtis dissimilarity value** |
| 26 | 51 | 0.667 | 21 | 41 | 0.560 |
| 24 | 45 | 0.663 | 3 | 51 | 0.559 |
| 20 | 29 | 0.661 | 10 | 29 | 0.559 |
| 41 | 51 | 0.661 | 4 | 38 | 0.559 |
| 14 | 20 | 0.661 | 6 | 35 | 0.559 |

| | | | | | |
|---|---|---|---|---|---|
| 2 | 24 | 0.661 | 10 | 39 | 0.559 |
| 4 | 25 | 0.657 | 6 | 52 | 0.559 |
| 12 | 25 | 0.654 | 4 | 46 | 0.559 |
| 1 | 29 | 0.653 | 29 | 45 | 0.559 |
| 24 | 35 | 0.653 | 10 | 26 | 0.558 |
| 24 | 29 | 0.653 | 38 | 52 | 0.558 |
| 24 | 46 | 0.653 | 19 | 41 | 0.558 |
| 14 | 21 | 0.653 | 2 | 18 | 0.558 |
| 20 | 23 | 0.652 | 10 | 41 | 0.558 |
| 24 | 48 | 0.649 | 2 | 3 | 0.558 |
| 2 | 26 | 0.649 | 2 | 4 | 0.557 |
| 1 | 14 | 0.648 | 18 | 29 | 0.557 |
| 4 | 43 | 0.647 | 21 | 48 | 0.557 |
| 26 | 43 | 0.647 | 9 | 20 | 0.556 |
| 3 | 25 | 0.647 | 19 | 26 | 0.556 |
| 24 | 37 | 0.646 | 25 | 48 | 0.556 |
| 29 | 51 | 0.645 | 21 | 23 | 0.556 |
| 43 | 51 | 0.645 | 23 | 46 | 0.556 |
| 25 | 38 | 0.645 | 16 | 29 | 0.556 |
| 3 | 21 | 0.644 | 9 | 43 | 0.556 |
| 1 | 18 | 0.644 | 10 | 23 | 0.555 |
| 16 | 25 | 0.643 | 10 | 50 | 0.555 |
| 23 | 29 | 0.643 | 29 | 35 | 0.555 |
| 2 | 49 | 0.642 | 21 | 37 | 0.554 |
| 4 | 24 | 0.640 | 10 | 19 | 0.554 |
| 19 | 24 | 0.638 | 3 | 12 | 0.552 |
| 6 | 24 | 0.637 | 45 | 51 | 0.552 |
| 39 | 51 | 0.636 | 6 | 38 | 0.552 |
| 3 | 24 | 0.635 | 14 | 38 | 0.552 |
| 49 | 51 | 0.634 | 6 | 41 | 0.551 |
| 21 | 24 | 0.634 | 1 | 49 | 0.551 |
| 10 | 51 | 0.633 | 35 | 52 | 0.551 |
| 16 | 51 | 0.633 | 49 | 52 | 0.550 |
| 6 | 20 | 0.632 | 23 | 48 | 0.550 |
| 10 | 24 | 0.631 | 41 | 46 | 0.550 |
| 24 | 52 | 0.630 | 4 | 48 | 0.550 |
| 20 | 21 | 0.630 | 9 | 26 | 0.550 |
| 14 | 29 | 0.630 | 6 | 19 | 0.549 |
| 14 | 43 | 0.630 | 6 | 37 | 0.549 |
| 26 | 50 | 0.630 | 35 | 51 | 0.549 |
| 29 | 43 | 0.630 | 46 | 52 | 0.549 |
| 25 | 43 | 0.629 | 19 | 20 | 0.549 |
| 4 | 49 | 0.628 | 50 | 52 | 0.548 |

| | | | | | |
|---|---|---|---|---|---|
| 1 | 51 | 0.628 | 23 | 38 | 0.548 |
| 18 | 51 | 0.627 | 4 | 12 | 0.548 |
| 1 | 41 | 0.627 | 14 | 18 | 0.547 |
| 9 | 25 | 0.627 | 35 | 49 | 0.547 |
| 18 | 20 | 0.625 | 18 | 37 | 0.547 |
| 24 | 39 | 0.624 | 6 | 10 | 0.546 |
| 1 | 13 | 0.624 | 6 | 26 | 0.546 |
| 23 | 51 | 0.624 | 10 | 16 | 0.545 |
| 1 | 23 | 0.624 | 19 | 39 | 0.545 |
| 4 | 6 | 0.624 | 4 | 13 | 0.545 |
| 14 | 51 | 0.623 | 2 | 9 | 0.544 |
| 25 | 46 | 0.622 | 37 | 41 | 0.544 |
| 4 | 41 | 0.622 | 39 | 43 | 0.544 |
| 2 | 51 | 0.622 | 13 | 14 | 0.542 |
| 4 | 29 | 0.620 | 23 | 37 | 0.542 |
| 19 | 23 | 0.620 | 10 | 49 | 0.542 |
| 20 | 51 | 0.620 | 39 | 41 | 0.542 |
| 4 | 51 | 0.619 | 37 | 51 | 0.542 |
| 20 | 39 | 0.619 | 4 | 39 | 0.541 |
| 1 | 16 | 0.619 | 43 | 49 | 0.541 |
| 24 | 38 | 0.619 | 2 | 13 | 0.541 |
| 16 | 49 | 0.619 | 25 | 45 | 0.540 |
| 2 | 14 | 0.617 | 6 | 9 | 0.540 |
| 12 | 51 | 0.617 | 2 | 12 | 0.540 |
| 25 | 50 | 0.617 | 19 | 52 | 0.539 |
| 14 | 50 | 0.617 | 38 | 49 | 0.539 |
| 6 | 23 | 0.617 | 39 | 52 | 0.538 |
| 9 | 24 | 0.616 | 16 | 35 | 0.537 |
| 21 | 49 | 0.615 | 37 | 49 | 0.537 |
| 23 | 24 | 0.615 | 18 | 52 | 0.537 |
| 21 | 46 | 0.615 | 9 | 10 | 0.536 |
| 13 | 51 | 0.615 | 19 | 43 | 0.536 |
| 21 | 50 | 0.614 | 9 | 29 | 0.534 |
| 23 | 26 | 0.614 | 12 | 14 | 0.534 |
| 2 | 41 | 0.614 | 6 | 16 | 0.534 |
| 14 | 24 | 0.613 | 18 | 38 | 0.533 |
| 51 | 52 | 0.613 | 19 | 46 | 0.533 |
| 21 | 29 | 0.613 | 23 | 45 | 0.532 |
| 6 | 14 | 0.612 | 14 | 19 | 0.532 |
| 1 | 4 | 0.612 | 6 | 18 | 0.531 |
| 4 | 14 | 0.612 | 13 | 38 | 0.531 |
| 29 | 37 | 0.612 | 4 | 37 | 0.530 |
| 3 | 26 | 0.612 | 3 | 10 | 0.530 |

| | | | | | |
|---|---|---|---|---|---|
| 4 | 21 | 0.611 | 39 | 46 | 0.530 |
| 1 | 21 | 0.611 | 3 | 13 | 0.530 |
| 4 | 26 | 0.610 | 16 | 39 | 0.529 |
| 13 | 20 | 0.609 | 9 | 38 | 0.529 |
| 14 | 49 | 0.609 | 9 | 46 | 0.528 |
| 1 | 52 | 0.609 | 13 | 24 | 0.528 |
| 10 | 20 | 0.609 | 2 | 20 | 0.528 |
| 26 | 37 | 0.608 | 20 | 48 | 0.528 |
| 41 | 49 | 0.608 | 41 | 52 | 0.528 |
| 23 | 50 | 0.608 | 4 | 9 | 0.527 |
| 29 | 52 | 0.608 | 2 | 19 | 0.527 |
| 4 | 52 | 0.606 | 13 | 23 | 0.527 |
| 3 | 41 | 0.605 | 3 | 9 | 0.526 |
| 1 | 6 | 0.605 | 26 | 29 | 0.526 |
| 21 | 43 | 0.605 | 1 | 19 | 0.526 |
| 9 | 23 | 0.605 | 25 | 37 | 0.526 |
| 10 | 21 | 0.604 | 3 | 19 | 0.525 |
| 21 | 52 | 0.603 | 9 | 50 | 0.524 |
| 2 | 21 | 0.603 | 39 | 50 | 0.524 |
| 3 | 14 | 0.602 | 6 | 43 | 0.524 |
| 18 | 49 | 0.602 | 20 | 35 | 0.523 |
| 26 | 46 | 0.602 | 10 | 38 | 0.523 |
| 16 | 43 | 0.601 | 13 | 35 | 0.523 |
| 41 | 43 | 0.601 | 10 | 18 | 0.523 |
| 4 | 50 | 0.601 | 3 | 4 | 0.522 |
| 14 | 52 | 0.599 | 4 | 45 | 0.522 |
| 14 | 46 | 0.599 | 2 | 46 | 0.522 |
| 1 | 10 | 0.598 | 18 | 45 | 0.521 |
| 6 | 29 | 0.598 | 18 | 19 | 0.521 |
| 2 | 25 | 0.598 | 19 | 50 | 0.521 |
| 9 | 51 | 0.597 | 9 | 52 | 0.520 |
| 18 | 24 | 0.597 | 16 | 52 | 0.520 |
| 1 | 9 | 0.597 | 2 | 38 | 0.520 |
| 19 | 49 | 0.596 | 9 | 39 | 0.520 |
| 20 | 43 | 0.596 | 3 | 38 | 0.519 |
| 12 | 20 | 0.596 | 26 | 41 | 0.519 |
| 2 | 23 | 0.595 | 9 | 21 | 0.519 |
| 26 | 49 | 0.595 | 6 | 13 | 0.518 |
| 14 | 39 | 0.594 | 10 | 52 | 0.517 |
| 16 | 50 | 0.594 | 6 | 39 | 0.517 |
| 26 | 38 | 0.594 | 6 | 45 | 0.516 |
| 21 | 39 | 0.593 | 37 | 52 | 0.516 |
| 24 | 26 | 0.592 | 3 | 37 | 0.515 |

| | | | | | |
|---|---|---|---|---|---|
| 10 | 43 | 0.592 | 10 | 37 | 0.515 |
| 23 | 43 | 0.591 | 2 | 35 | 0.515 |
| 13 | 49 | 0.590 | 1 | 46 | 0.514 |
| 10 | 14 | 0.590 | 13 | 37 | 0.514 |
| 23 | 49 | 0.590 | 18 | 39 | 0.514 |
| 19 | 29 | 0.590 | 16 | 41 | 0.513 |
| 29 | 39 | 0.590 | 18 | 48 | 0.513 |
| 18 | 43 | 0.589 | 41 | 45 | 0.513 |
| 29 | 50 | 0.589 | 13 | 21 | 0.512 |
| 23 | 52 | 0.589 | 35 | 39 | 0.511 |
| 26 | 45 | 0.588 | 16 | 37 | 0.511 |
| 3 | 6 | 0.587 | 41 | 48 | 0.511 |
| 1 | 25 | 0.586 | 10 | 46 | 0.510 |
| 26 | 35 | 0.586 | 12 | 21 | 0.510 |
| 4 | 23 | 0.586 | 12 | 23 | 0.510 |
| 13 | 50 | 0.586 | 10 | 13 | 0.509 |
| 43 | 52 | 0.586 | 13 | 48 | 0.509 |
| 4 | 10 | 0.585 | 4 | 16 | 0.508 |
| 1 | 3 | 0.585 | 48 | 52 | 0.508 |
| 3 | 49 | 0.585 | 9 | 41 | 0.508 |
| 14 | 23 | 0.584 | 1 | 2 | 0.507 |
| 21 | 26 | 0.584 | 3 | 48 | 0.507 |
| 20 | 46 | 0.584 | 13 | 45 | 0.507 |
| 4 | 20 | 0.584 | 13 | 19 | 0.507 |
| 23 | 41 | 0.584 | 45 | 52 | 0.505 |
| 38 | 51 | 0.584 | 18 | 26 | 0.505 |
| 21 | 35 | 0.583 | 18 | 35 | 0.503 |
| 20 | 41 | 0.583 | 16 | 48 | 0.502 |
| 18 | 21 | 0.583 | 12 | 43 | 0.502 |
| 18 | 50 | 0.583 | 16 | 26 | 0.500 |
| 26 | 52 | 0.582 | 1 | 38 | 0.500 |
| 24 | 41 | 0.582 | 19 | 38 | 0.499 |
| 14 | 45 | 0.582 | 1 | 35 | 0.498 |
| 3 | 52 | 0.581 | 16 | 45 | 0.496 |
| 1 | 39 | 0.581 | 49 | 50 | 0.496 |
| 18 | 46 | 0.581 | 12 | 50 | 0.496 |
| 46 | 51 | 0.580 | 13 | 52 | 0.496 |
| 29 | 49 | 0.580 | 10 | 35 | 0.495 |
| 3 | 20 | 0.580 | 1 | 43 | 0.495 |
| 4 | 18 | 0.580 | 20 | 45 | 0.495 |
| 3 | 23 | 0.579 | 12 | 29 | 0.495 |
| 20 | 52 | 0.578 | 13 | 39 | 0.494 |
| 2 | 52 | 0.578 | 3 | 46 | 0.492 |

| | | | | | |
|---|---|---|---|---|---|
| 6 | 50 | 0.578 | 10 | 48 | 0.492 |
| 20 | 50 | 0.578 | 35 | 46 | 0.492 |
| 4 | 19 | 0.577 | 12 | 52 | 0.491 |
| 29 | 38 | 0.577 | 20 | 37 | 0.490 |
| 20 | 38 | 0.577 | 10 | 12 | 0.490 |
| 2 | 50 | 0.576 | 37 | 39 | 0.489 |
| 26 | 48 | 0.576 | 35 | 43 | 0.489 |
| 9 | 14 | 0.576 | 6 | 48 | 0.489 |
| 12 | 24 | 0.575 | 9 | 48 | 0.488 |
| 6 | 49 | 0.574 | 6 | 12 | 0.488 |
| 14 | 37 | 0.574 | 10 | 45 | 0.488 |
| 13 | 43 | 0.573 | 18 | 41 | 0.486 |
| 3 | 43 | 0.573 | 12 | 41 | 0.486 |
| 16 | 46 | 0.573 | 12 | 19 | 0.486 |
| 1 | 20 | 0.573 | 48 | 49 | 0.486 |
| 16 | 24 | 0.572 | 12 | 35 | 0.485 |
| 23 | 39 | 0.572 | 38 | 39 | 0.485 |
| 50 | 51 | 0.572 | 3 | 45 | 0.485 |
| 9 | 49 | 0.572 | 9 | 35 | 0.484 |
| 26 | 39 | 0.572 | 9 | 18 | 0.483 |
| 29 | 41 | 0.572 | 12 | 46 | 0.482 |
| 18 | 23 | 0.571 | 38 | 46 | 0.482 |
| 19 | 25 | 0.571 | 13 | 26 | 0.482 |
| 13 | 29 | 0.570 | 9 | 12 | 0.481 |
| 29 | 48 | 0.570 | 12 | 26 | 0.481 |
| 41 | 50 | 0.570 | 2 | 45 | 0.481 |
| 39 | 49 | 0.570 | 39 | 45 | 0.480 |
| 4 | 35 | 0.569 | 1 | 50 | 0.480 |
| 14 | 26 | 0.569 | 16 | 18 | 0.478 |
| 14 | 48 | 0.569 | 13 | 41 | 0.477 |
| 14 | 35 | 0.568 | 37 | 43 | 0.477 |
| 19 | 51 | 0.568 | 46 | 49 | 0.476 |
| 13 | 46 | 0.568 | 9 | 13 | 0.475 |
| 6 | 21 | 0.567 | 19 | 45 | 0.471 |
| 14 | 41 | 0.567 | 9 | 19 | 0.470 |
| 6 | 46 | 0.566 | 12 | 39 | 0.469 |
| 21 | 38 | 0.566 | 1 | 45 | 0.469 |
| 16 | 23 | 0.566 | 20 | 25 | 0.468 |
| 19 | 21 | 0.565 | 2 | 48 | 0.468 |
| 2 | 6 | 0.565 | 3 | 35 | 0.467 |
| 25 | 35 | 0.565 | 12 | 18 | 0.467 |
| 35 | 41 | 0.565 | 2 | 37 | 0.466 |
| 2 | 39 | 0.565 | 38 | 43 | 0.466 |

| | | | | | |
|---|---|---|---|---|---|
| 48 | 51 | 0.564 | 46 | 50 | 0.465 |
| 21 | 45 | 0.564 | 9 | 37 | 0.464 |
| 23 | 35 | 0.564 | 43 | 46 | 0.463 |
| 3 | 39 | 0.563 | 16 | 19 | 0.463 |
| 3 | 29 | 0.563 | 12 | 37 | 0.463 |
| 14 | 16 | 0.562 | 45 | 49 | 0.461 |
| 38 | 41 | 0.562 | 12 | 38 | 0.458 |
| 3 | 18 | 0.561 | 35 | 50 | 0.458 |
| 16 | 38 | 0.561 | 16 | 21 | 0.456 |
| 16 | 20 | 0.561 | 37 | 46 | 0.453 |
| 2 | 43 | 0.561 | 9 | 45 | 0.452 |
| 3 | 50 | 0.561 | 43 | 45 | 0.452 |
| 2 | 16 | 0.561 | 12 | 49 | 0.451 |
| 2 | 10 | 0.561 | 12 | 45 | 0.450 |
| 3 | 16 | 0.560 | 38 | 50 | 0.446 |
| 1 | 12 | 0.560 | 37 | 50 | 0.446 |
| 29 | 46 | 0.560 | 19 | 35 | 0.445 |
| **Excluded fragment 28 as highly dissimilar outlier:** | | | | | |
| **fragment A** | **fragment B** | **Bray-Curtis dissimilarity value** | **fragment A** | **fragment B** | **Bray-Curtis dissimilarity value** |
| 28 | 49 | 0.802 | 28 | 29 | 0.741 |
| 10 | 28 | 0.785 | 28 | 50 | 0.740 |
| 14 | 28 | 0.775 | 28 | 45 | 0.736 |
| 3 | 28 | 0.768 | 28 | 46 | 0.735 |
| 28 | 52 | 0.764 | 2 | 28 | 0.730 |
| 25 | 28 | 0.762 | 12 | 28 | 0.728 |
| 6 | 28 | 0.761 | 28 | 48 | 0.725 |
| 24 | 28 | 0.761 | 28 | 43 | 0.717 |
| 26 | 28 | 0.757 | 28 | 35 | 0.717 |
| 28 | 39 | 0.751 | 13 | 28 | 0.717 |
| 4 | 28 | 0.748 | 28 | 37 | 0.706 |
| 28 | 51 | 0.746 | 19 | 28 | 0.703 |
| 20 | 28 | 0.746 | 28 | 38 | 0.685 |
| 23 | 28 | 0.745 | 16 | 28 | 0.678 |
| 18 | 28 | 0.743 | 9 | 28 | 0.670 |
| 1 | 28 | 0.742 | 21 | 28 | 0.620 |
| 28 | 41 | 0.742 | | | |

**Fig. S13. Bootstrap plot of FAVA results to test for significant difference in microbiome heterogeneity across fragments within each region** (high-fragmentation – HF; low-fragmentation – LF).

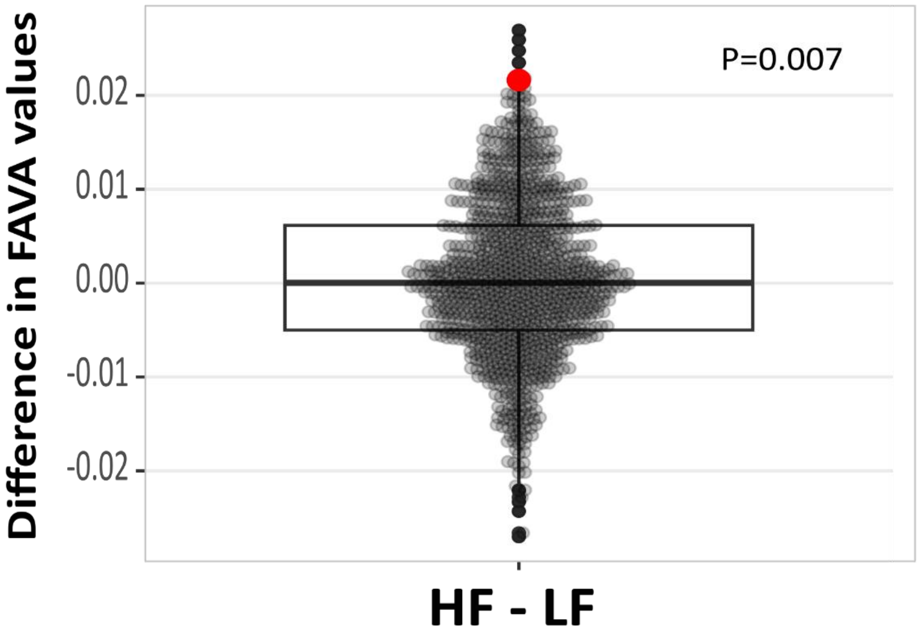

**Table S14. Full results of all nested PERMANOVA models.** $R^2$ values represent the proportion of total variation attributed to each sequentially nested term. Colons denote nested terms. Bold P-value cells indicate $P < 0.05$. A) Whole fragmented landscape, model: north_south_MB/fragment/group/age/sex; 999 permutations; B) high-fragmented region, model: fragment/group/age/sex; 999 permutations; C) low-fragmented region, model: fragment/group/age/sex; 999 permutations.

A) Whole fragmented landscape

| **Distance metric** | **Model term** | **df** | **Sum of squares** | **$R^2$** | **Pseudo-F** | **P value** |
|---|---|---|---|---|---|---|
| **Bray–Curtis** | **north_south_MB** | **1** | **3.875** | **0.046** | **22.593** | **0.001** |
| | north_south_MB:fragment | 1 | 0.924 | 0.011 | 5.386 | **0.001** |
| | north_south_MB:fragment:group | 65 | 32.729 | 0.385 | 2.936 | **0.001** |
| | north_south_MB:fragment:group:age | 76 | 16.879 | 0.199 | 1.295 | **0.001** |
| | north_south_MB:fragment:group:age:sex | 74 | 15.246 | 0.180 | 1.201 | **0.001** |
| | Residual | 89 | 15.266 | 0.180 | | |
| | Total | 306 | 84.920 | 1.000 | | |
| **Jaccard** | **north_south_MB** | **1** | **3.138** | **0.028** | **11.740** | **0.001** |
| | north_south_MB:fragment | 1 | 0.904 | 0.008 | 3.383 | **0.001** |
| | north_south_MB:fragment:group | 65 | 36.732 | 0.331 | 2.114 | **0.001** |
| | north_south_MB:fragment:group:age | 76 | 24.083 | 0.217 | 1.186 | **0.001** |
| | north_south_MB:fragment:group:age:sex | 74 | 22.259 | 0.201 | 1.125 | **0.001** |
| | Residual | 89 | 23.790 | 0.215 | | |
| | Total | 306 | 110.907 | 1.000 | | |
| **Unweighted UniFrac** | **north_south_MB** | **1** | **1.875** | **0.043** | **18.766** | **0.001** |
| | north_south_MB:fragment | 1 | 0.324 | 0.007 | 3.237 | **0.001** |
| | north_south_MB:fragment:group | 65 | 14.241 | 0.329 | 2.192 | **0.001** |
| | north_south_MB:fragment:group:age | 76 | 9.811 | 0.226 | 1.292 | **0.001** |
| | north_south_MB:fragment:group:age:sex | 74 | 8.191 | 0.189 | 1.108 | **0.004** |
| | Residual | 89 | 8.895 | 0.205 | | |
| | Total | 306 | 43.337 | 1.000 | | |
| **Weighted UniFrac** | **north_south_MB** | **1** | **0.020** | **0.012** | **7.387** | **0.003** |
| | north_south_MB:fragment | 1 | 0.025 | 0.015 | 9.120 | **0.001** |
| | north_south_MB:fragment:group | 65 | 0.673 | 0.407 | 3.765 | **0.001** |
| | north_south_MB:fragment:group:age | 76 | 0.387 | 0.234 | 1.851 | **0.001** |
| | north_south_MB:fragment:group:age:sex | 74 | 0.303 | 0.183 | 1.488 | **0.014** |
| | Residual | 89 | 0.245 | 0.148 | | |
| | Total | 306 | 1.652 | 1.000 | | |

B) High-fragmented region

| Distance metric | Model term | df | Sum of squares | $R^2$ | Pseudo-F | P value |
|---|---|---|---|---|---|---|
| **Bray–Curtis** | **Fragment** | **1** | **0.626** | **0.020** | **3.725** | **0.001** |
| | fragment:group | 23 | 12.299 | 0.391 | 3.180 | **0.001** |
| | fragment:group:age | 30 | 6.712 | 0.213 | 1.331 | **0.001** |
| | fragment:group:age:sex | 28 | 5.777 | 0.184 | 1.227 | **0.001** |
| | Residual | 36 | 6.054 | 0.192 | | |
| | Total | 118 | 31.468 | 1.000 | | |
| **Jaccard** | **Fragment** | **1** | **0.674** | **0.016** | **2.544** | **0.001** |
| | fragment:group | 23 | 13.632 | 0.326 | 2.235 | **0.001** |
| | fragment:group:age | 30 | 9.542 | 0.228 | 1.200 | **0.001** |
| | fragment:group:age:sex | 28 | 8.377 | 0.201 | 1.128 | **0.001** |
| | Residual | 36 | 9.546 | 0.229 | | |
| | Total | 118 | 41.772 | 1.000 | | |
| **Unweighted UniFrac** | **Fragment** | **1** | **0.315** | **0.020** | **3.375** | **0.001** |
| | fragment:group | 23 | 5.339 | 0.339 | 2.491 | **0.001** |
| | fragment:group:age | 30 | 3.554 | 0.226 | 1.271 | **0.001** |
| | fragment:group:age:sex | 28 | 3.176 | 0.202 | 1.217 | **0.001** |
| | Residual | 36 | 3.356 | 0.213 | | |
| | Total | 118 | 15.739 | 1.000 | | |
| **Weighted UniFrac** | **Fragment** | **1** | **0.027** | **0.042** | **11.580** | **0.001** |
| | fragment:group | 23 | 0.211 | 0.324 | 3.897 | **0.001** |
| | fragment:group:age | 30 | 0.208 | 0.321 | 2.953 | **0.001** |
| | fragment:group:age:sex | 28 | 0.119 | 0.183 | 1.808 | **0.013** |
| | Residual | 36 | 0.085 | 0.130 | | |
| | Total | 118 | 0.650 | 1.000 | | |

C) Low-fragmented region

| Distance metric | Model term | df | Sum of squares | $R^2$ | Pseudo-F | P value |
|---|---|---|---|---|---|---|
| **Bray–Curtis** | **Fragment** | **1** | **1.194** | **0.024** | **6.870** | **0.001** |
| | fragment:group | 41 | 19.534 | 0.394 | 2.741 | **0.001** |
| | fragment:group:age | 46 | 10.167 | 0.205 | 1.272 | **0.001** |
| | fragment:group:age:sex | 46 | 9.469 | 0.191 | 1.184 | **0.002** |
| | Residual | 53 | 9.212 | 0.186 | | |
| | Total | 187 | 49.577 | 1.000 | | |
| **Jaccard** | **Fragment** | **1** | **1.137** | **0.017** | **4.230** | **0.001** |
| | fragment:group | 41 | 22.194 | 0.336 | 2.014 | **0.001** |
| | fragment:group:age | 46 | 14.541 | 0.220 | 1.176 | **0.001** |
| | fragment:group:age:sex | 46 | 13.882 | 0.210 | 1.123 | **0.001** |
| | Residual | 53 | 14.244 | 0.216 | | |
| | Total | 187 | 65.997 | 1.000 | | |
| **Unweighted UniFrac** | **Fragment** | **1** | **0.415** | **0.016** | **3.972** | **0.001** |
| | fragment:group | 41 | 8.495 | 0.330 | 1.983 | **0.001** |
| | fragment:group:age | 46 | 6.257 | 0.243 | 1.302 | **0.001** |
| | fragment:group:age:sex | 46 | 5.015 | 0.195 | 1.043 | 0.207 |
| | Residual | 53 | 5.539 | 0.215 | | |
| | Total | 187 | 25.722 | 1.000 | | |
| **Weighted UniFrac** | **Fragment** | **1** | **0.008** | **0.008** | **2.518** | **0.092** |
| | fragment:group | 41 | 0.452 | 0.461 | 3.653 | **0.001** |
| | fragment:group:age | 46 | 0.178 | 0.182 | 1.284 | 0.129 |
| | fragment:group:age:sex | 46 | 0.184 | 0.187 | 1.322 | 0.110 |
| | Residual | 53 | 0.160 | 0.163 | | |
| | Total | 187 | 0.982 | 1.000 | | |

**Fig. S14. Nested PERMANOVA results for all dissimilarity indices.** Top – for whole dataset (both fragmented regions) together. Bottom – For the northern, high-fragmented region (left) and southern, low-fragmented region (right) separately. Global model significance denoted by asterisks at top; significance of each variable and its proportional contribution to explained variance noted in the plots. * = 0.01 – 0.05; ** = 0.001 – 0.01; *** = 0 – 0.001.

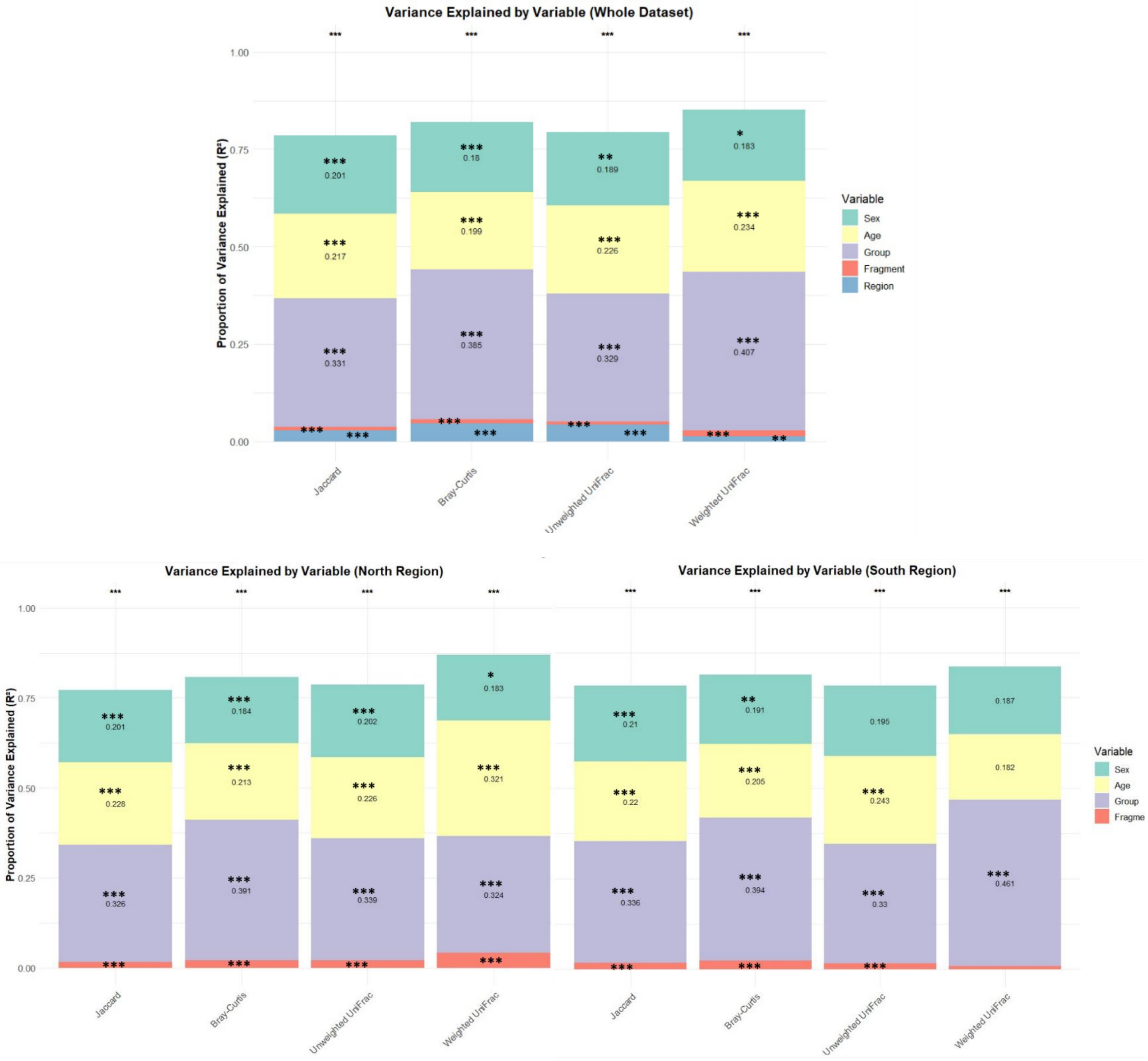

**Table S15. Results of distance-to-centroid and permutation tests** to determine if the microbiomes of likely long-distance dispersers were more similar to the microbiomes in their sampling location or their inferred location of origin**.** Column headers: LR: Lynch-Ritland pairwise relatedness coefficient; Origin: putative group/ fragment of origin; Sample size - origin: number of samples in putative group of origin, when fragment contained only one group, or number of samples in putative fragment of origin, when fragment contained multiple groups; Sample size - sampling group: number of samples in sampling group, excluding disperser; observed delta: negative - sample closer to putative origin; positive - sample closer sampling location; 2-sided p-value: permutation test to determine if the sample is significantly unequally distant to the centroids of the 2 groups; 1-sided p-value: permutation test to determine if sample is closer to likely group of origin; 1-sided p-value: permutation test to determine if sample is closer to sampling location. In red – significant results ($p < 0.05$).

| **Individual 1** | | | | | **Individual 2** | | | | | | | | | | | **results of distance to centroid test based on PCoA coordinates of Bray-Curtis dissimilarity.** | | | |
|---|---|---|---|---|---|---|---|---|---|---|---|---|---|---|---|---|---|---|---|
| **Sample ID** | **Frag-ment** | **Group** | **Age** | **Sex** | **Sample ID** | **Frag-ment** | **Group** | **Age** | **Sex** | **LR** | **likely disperser** | **Origin** | **sampling group** | **sample size - origin** | **sample size - sampling group** | **observed delta** | **2-sided p-value** | **1 sided p-value: sample closer to origin** | **1 sided p-value: sample closer to sampling group** |
| LE5 | 37 | 37A | A | F | KK107 | 23 | 23A | J | F | 0.15 | **KK107** | 37 | 23A | 23 | 3 | 0.167 | 0.001 | 1 | 0 |
| CA14 | 35 | 35B | A | F | KK3 | 10 | 10B | A | F | 0.15 | **KK3** | 35 | 10B | 25 | 4 | 0.154 | 0 | 1 | 0 |
| AC14 | 18 | 18A | A | F | DL37 | 25 | 25A | A | M | 0.14 | **DL37** | 18A | 25A | 4 | 8 | 0.2169 | 0.001 | 1 | 0.001 |
| DL14 | 37 | 37D | A | F | DL7 | 3 | GHA | SA | M | 0.13 | **DL7** | 37 | 3A | 23 | 5 | 0.1074 | 0.006 | 0.998 | 0.002 |
| CA14 | 35 | 35B | A | F | KK107 | 23 | 23A | J | F | 0.11 | **KK107** | 35 | 23A | 25 | 3 | 0.1317 | 0.006 | 0.998 | 0.002 |
| LE16 | 25 | 25A | A | M | SM9 | 1 | 1C | A | M | 0.14 | **LE16** | 1C | 25A | 5 | 8 | 0.239 | 0.004 | 0.998 | 0.004 |
| DL37 | 25 | 25A | A | M | KK76 | 1 | 1C | A | F | 0.22 | **KK76** | 25A | 1C | 8 | 5 | 0.226 | 0.006 | 0.996 | 0.006 |
| DL12 | 37 | 37A | A | M | DL9 | 3 | GHA | A | M | 0.12 | **DL9** | 37A | 3A | 5 | 5 | 0.106 | 0.089 | 0.965 | 0.04 |
| SM39 | 35 | 35F | A | F | CA26 | 23 | 23A | A | F | 0.24 | **CA26** | 35F | 23A | 4 | 3 | 0.047 | 0.272 | 0.903 | 0.122 |
| SM39 | 35 | 35F | A | F | CA25 | 23 | 23A | SA | M | 0.11 | **CA25** | 35F | 23A | 4 | 3 | 0.089 | 0.277 | 0.898 | 0.127 |
| KK3 | 10 | 10B | A | F | KK107 | 23 | 23A | J | F | 0.16 | **KK107** | 10B | 23A | 5 | 3 | 0.0157 | 0.589 | 0.746 | 0.275 |

**Fig. S15. Selected examples of plotted distance-to-centroid test results** assessing if likely long-distance, between-fragment dispersers' gut microbiomes are more similar to those in their sampling group or putative group of origin, as measured by the distance between the disperser's Bray-Curtis dissimilarity index PCoA coordinates (red dot) and the coordinates of the centroids of the inferred origin (green) and sampling (purple) groups.

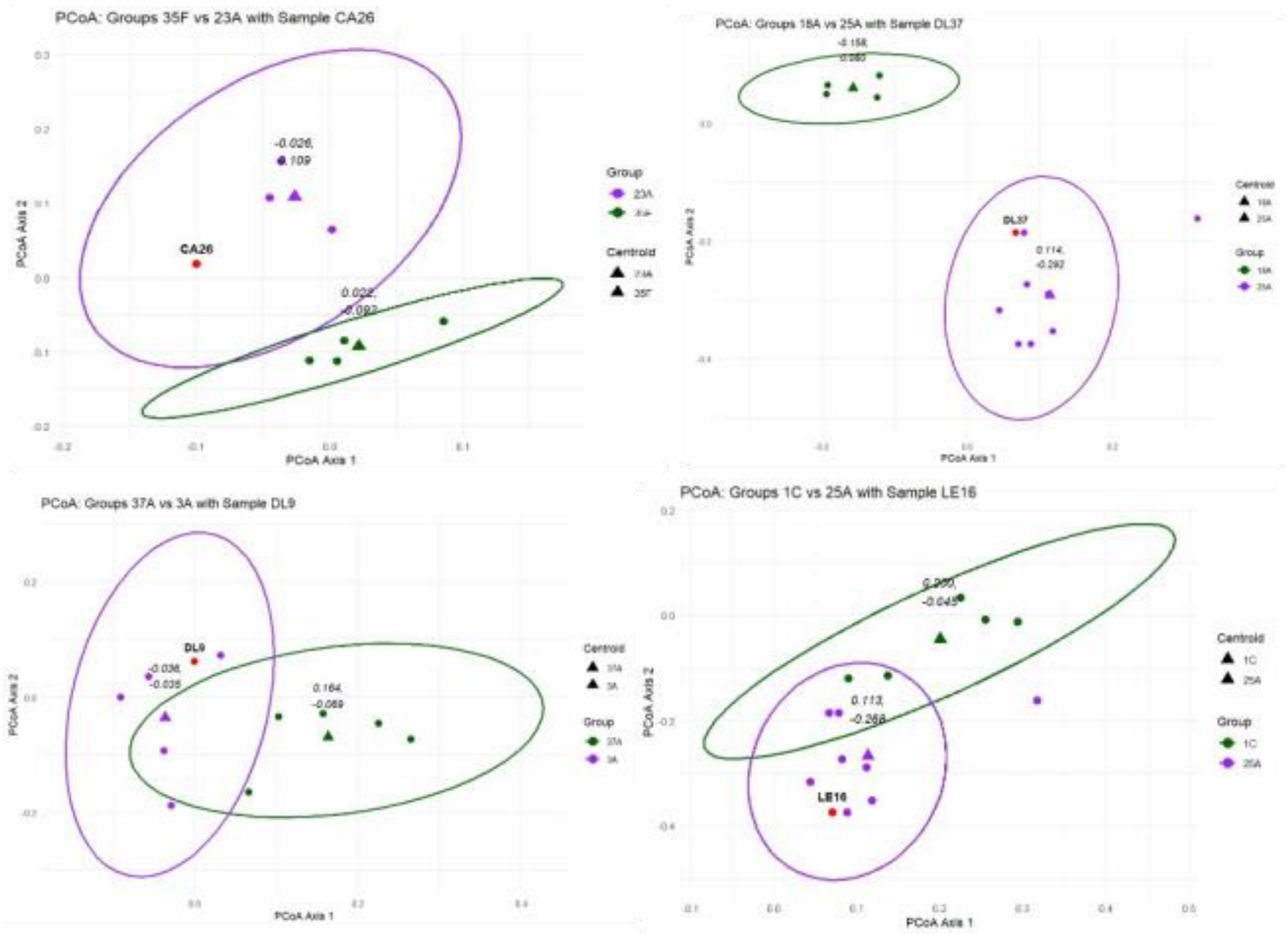

**Fig. S16. Mantel tests for the correlation of microbiome dissimilarity with Euclidean distance, for four dissimilarity indices**. A) Whole dataset (N = 307); B) subset of data including solitary individuals and one individual per social group (N=68); C) subset of data including all individuals in the HF region (N=119); D) subset of data including all individuals in the LF region (N=188); E) subset of HF region including all solitary individuals and one individual per social group (N=25); F) subset of LF region including all solitary individuals and one individual per social group (N=43).

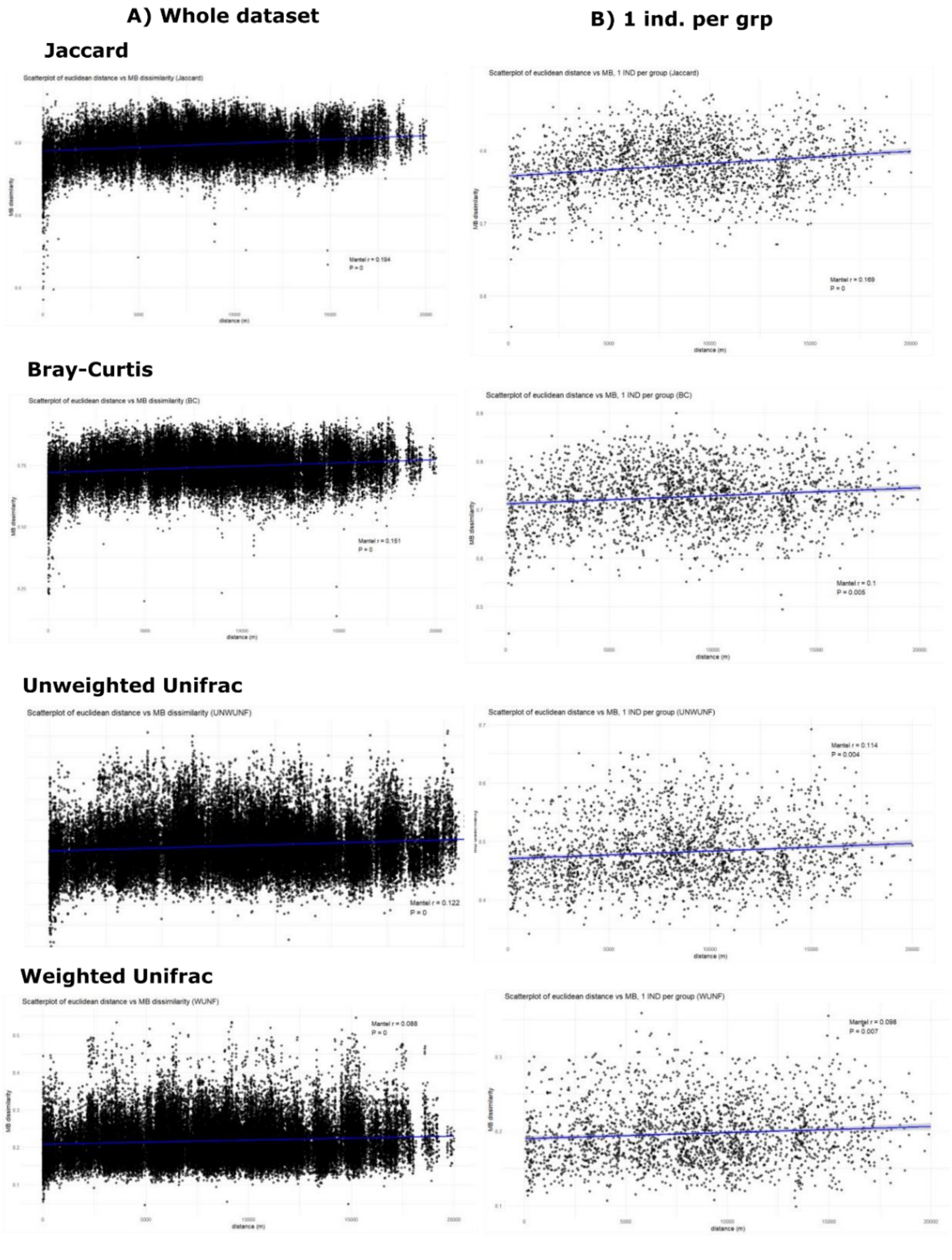

**C) HF region whole dataset** **D) LF region whole dataset**

**Jaccard**

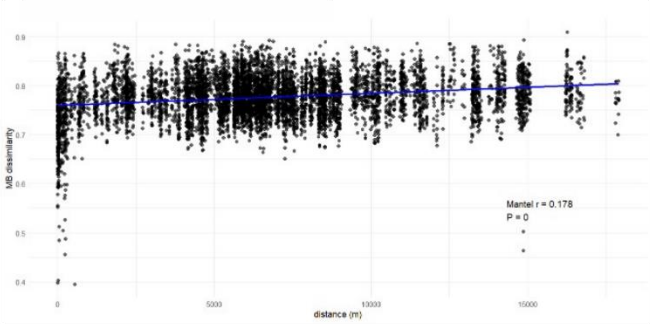

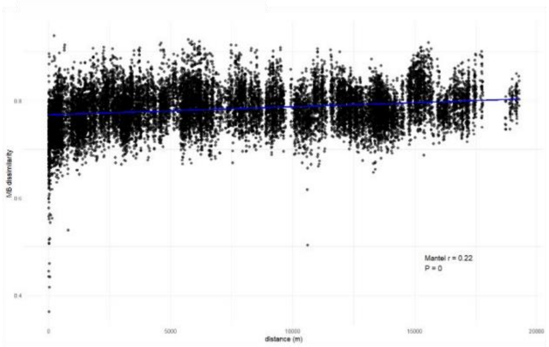

**Bray-Curtis**

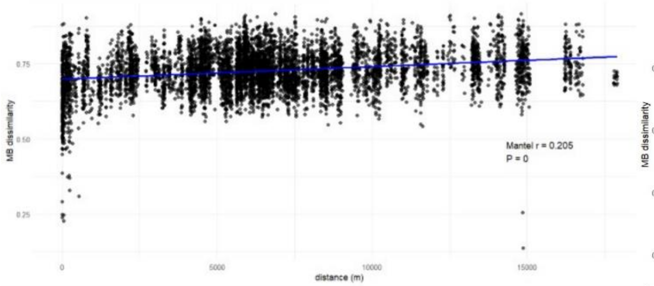

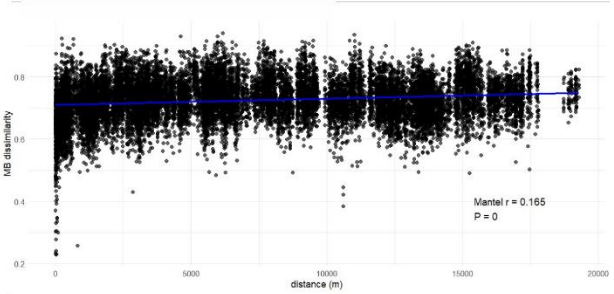

**Unweighted Unifrac**

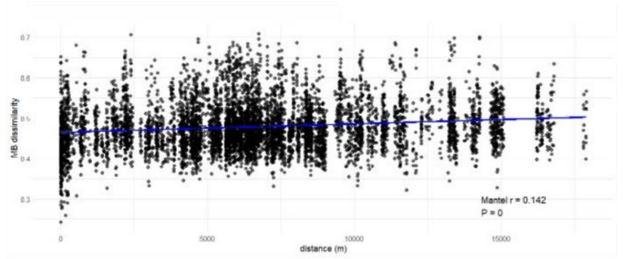

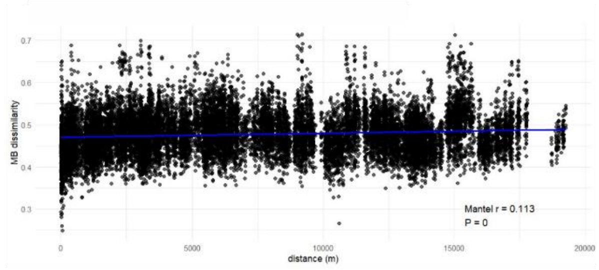

**Weighted Unifrac**

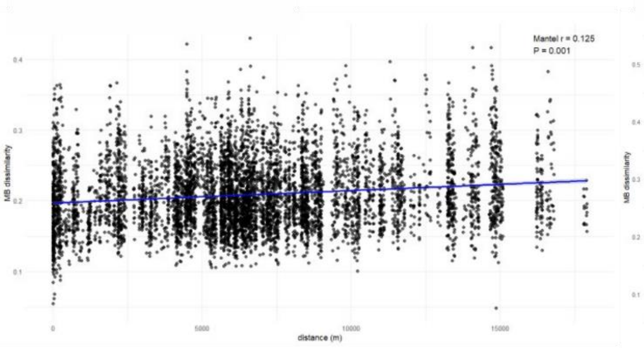

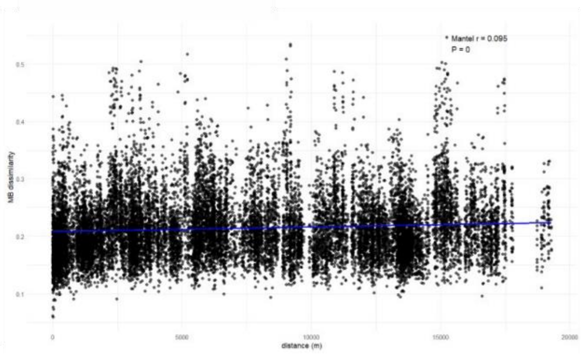

#### E) HF - 1 ind. per grp

#### F) LF - 1 ind. per grp

**Jaccard**

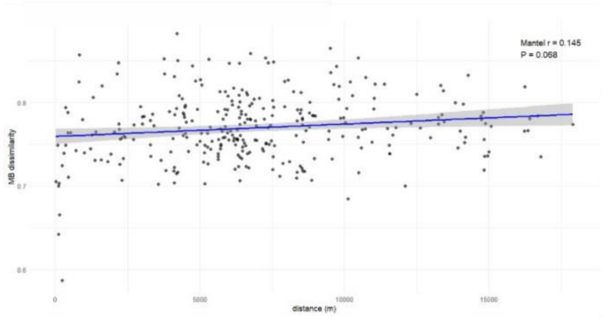

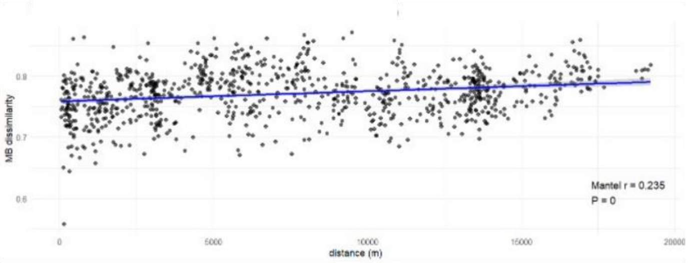

**Bray-Curtis**

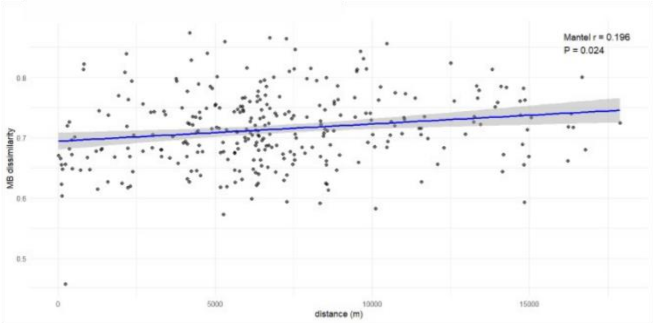

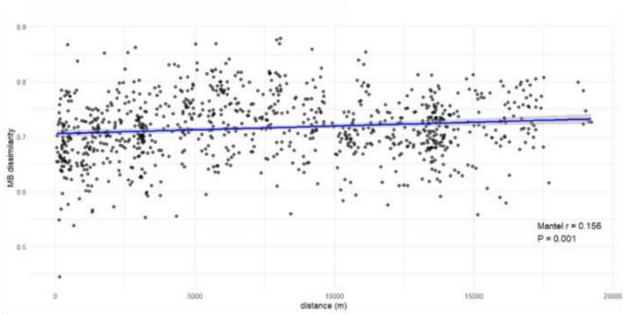

**Unweighted Unifrac**

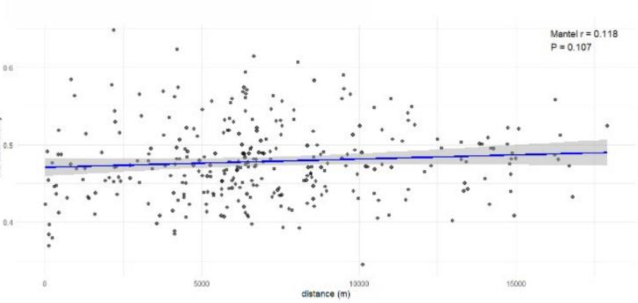

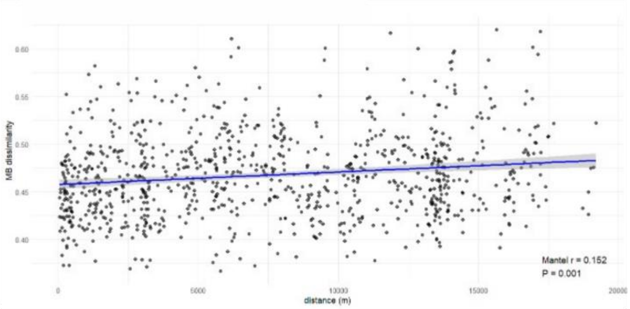

**Weighted Unifrac**

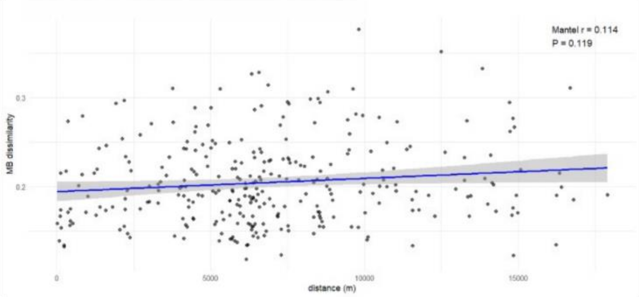

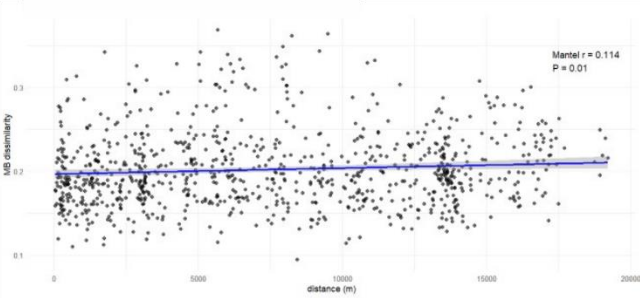

**Fig. S17. Mantel tests for the correlation of fragment microbiome dissimilarity with four dissimilarity indices, with the Bray-Curtis dissimilarity index for fragment tree composition, by region.** Left – HF region (N=13 fragments); right – LF region (N=15 fragments).

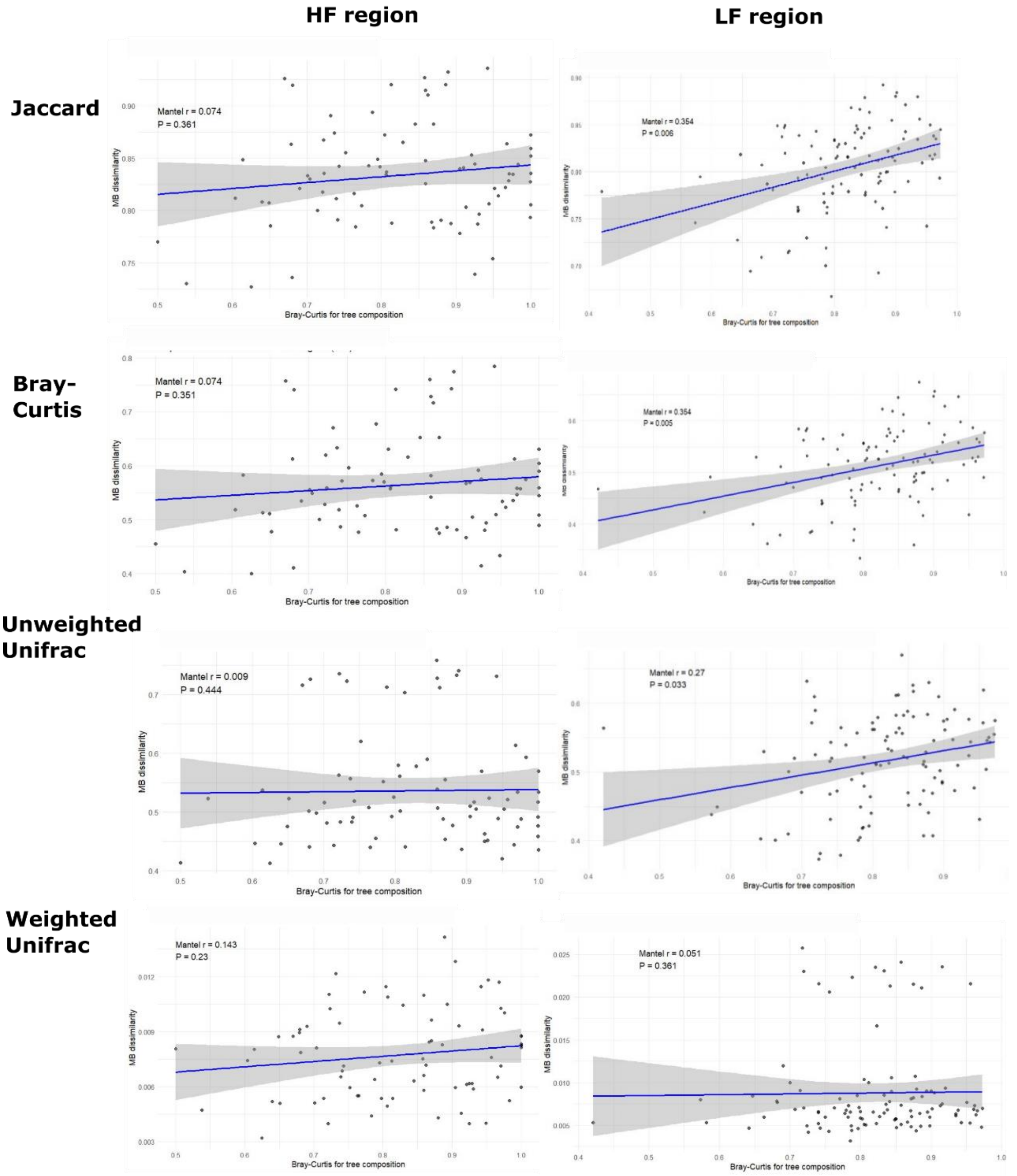

### Ecological and demographic drivers of patterns in black howler population genetics and microbiomes

#### Data summaries and variable selection

**Table S16. Data for all by-fragment explanatory and response variables used in analyses, primarily in the LASSO and multiple regression analyses**. NA – indicates that that variable was unavailable for that fragment, for example, fragments in which vegetation transect data were not collected will have "NA" noted for variables such as stem density and tree genus richness, and fragments from which no samples made it through the quality control and filtering stages of the bioinformatic pipeline for genetic data will have "NA" noted for all population genetic response variables.

| | Explanatory variables | | | | | | | | | | | | | | | | | | | |
|---|---|---|---|---|---|---|---|---|---|---|---|---|---|---|---|---|---|---|---|---|
| | Habitat quality | | | | | | | | | | | | | | | | | Isolation | | |
| fragment | size (ha) | core area | core area index | shape index | stem density | Bray Curtis dissimilarity - trees | proportion of ficus trees | proportion known important food trees | genus richness - trees | shannon diversity - trees | simpson's evenness - trees | mean tree DBH | max. tree DBH | mean tree height | max tree height | number tree stumps | number treelines | proportion of forest cover in buffer | distance to PNP (m) | distance to nearest fragment (m) |
| **1** | 28.7 | 148905900 | 0.490 | 2.79 | 0.048 | 0.867 | 0.07 | 27.08 | 25 | 3.03 | 0.69 | 45.6 | 326 | 15.5 | 40.1 | 6 | 6 | 0.2 | 335 | 20 |
| **2** | 8.1 | 148905900 | 0.490 | 4.08 | 0.02 | 0.888 | 0 | 45 | 14 | 2.45 | 0.63 | 55.2 | 219 | 13.8 | 32.6 | 2 | 3 | 0.39 | 812 | 549 |
| **3** | 2.61 | 148905900 | 0.490 | 2.39 | 0.018 | 0.882 | 0.071 | 22.22 | 13 | 2.48 | 0.82 | 55.4 | 202 | 14.8 | 37.5 | 3 | 4 | 0.43 | 1521 | 185 |
| **4** | 3.67 | 5400 | 0.097 | 2.26 | 0.013 | 0.882 | 0.308 | 38.5 | 7 | 1.78 | 0.83 | 80.6 | 350 | 12.1 | 24.9 | 3 | 3 | 0.34 | 1798 | 463 |
| **9** | 8.26 | 54900 | 0.550 | 1.52 | 0.064 | 0.863 | 0 | 9.52 | 21 | 2.48 | 0.3 | 19.8 | 61.4 | 10.9 | 23.1 | 12 | 4 | 0.31 | 2086 | 15 |
| **10** | 12.4 | 109800 | 0.387 | 1.84 | 0.024 | 0.993 | 0 | 0 | 4 | 0.51 | 0.32 | 42 | 89.1 | 20.8 | 29.6 | 1 | 0 | 0.21 | 4175 | 10 |
| **12** | 13.3 | 24300 | 0.092 | 3.71 | 0.046 | 0.929 | 0 | 63.04 | 7 | 1.24 | 0.33 | 30.9 | 87.7 | 13.1 | 21.3 | 1 | 4 | 0.12 | 7470 | 285 |
| **13** | 26.1 | 174600 | 0.474 | 2.05 | 0.048 | 0.835 | 0 | 6.25 | 18 | 2.44 | 0.44 | 26.7 | 60.7 | 14 | 22.7 | 7 | 6 | 0.12 | 6807 | 107 |
| **14** | 4.54 | 15300 | 0.354 | 1.68 | 0.045 | 0.866 | 0 | 2.22 | 11 | 1.81 | 0.37 | 35.9 | 145 | 11.4 | 24.6 | 3 | 2 | 0.12 | 7690 | 142 |
| **16** | 12.1 | 31500 | 0.330 | 2.22 | 0.051 | 0.807 | 0.051 | 5.88 | 16 | 2.50 | 0.61 | 22.6 | 116.5 | 9.7 | 21.2 | 5 | 1 | 0.22 | 9296 | 68 |
| **18** | 5.47 | 18900 | 0.339 | 1.73 | 0.04 | 0.856 | 0 | 10.26 | 18 | 2.43 | 0.35 | 27.9 | 128.9 | 13.1 | 28.4 | 5 | 1 | 0.25 | 7807 | 123 |
| **19** | 27 | 247500 | 0.222 | 3.46 | 0.036 | 0.809 | 0 | 11.11 | 14 | 2.28 | 0.51 | 25.1 | 66 | 11 | 20.9 | 0 | 8 | 0.19 | 9045 | 63 |
| **20** | 6.98 | 247500 | 0.222 | 1.92 | 0.03 | 0.864 | 0.107 | 10 | 8 | 1.49 | 0.36 | 32.8 | 80 | 11.6 | 20.2 | 2 | 4 | 0.22 | 7657 | 9 |
| **21** | 11.7 | 37800 | 0.167 | 1.77 | 0.033 | 0.805 | 0.091 | 21.21 | 18 | 2.61 | 0.57 | 35 | 116 | 10 | 20.4 | 4 | 2 | 0.25 | 9618 | 508 |
| **24** | 3.6 | 8100 | 0.225 | 2.28 | 0.082 | 0.913 | 0 | 1.22 | 8 | 0.70 | 0.17 | 21 | 69.7 | 12.8 | 21.5 | 1 | 0 | 0.16 | 8605 | 163 |
| **25** | 8.88 | 42300 | 0.435 | 1.54 | 0.048 | 0.86 | 0 | 4.17 | 13 | 2.07 | 0.4 | 32.6 | 170 | 10.3 | 25.5 | 1 | 2 | 0.23 | 7224 | 16 |
| **26** | 3.17 | 8100 | 0.243 | 1.6 | 0.053 | 0.851 | 0 | 0 | 20 | 2.65 | 0.52 | 20.6 | 131 | 8 | 28.4 | 2 | 1 | 0.1 | 6477 | 536 |
| **28** | 3.33 | 0 | 0.000 | 1.71 | 0.048 | 0.846 | 0 | 0 | 14 | 2.23 | 0.45 | 26.7 | 95 | 10.2 | 21 | 7 | 0 | 0.13 | 4129 | 209 |
| **29** | 4.99 | 3150 | 0.042 | 3.13 | 0.055 | 0.825 | 0 | 13.46 | 18 | 2.68 | 0.7 | 25.2 | 92 | 7.4 | 20.2 | 8 | 6 | 0.17 | 4131 | 417 |
| **35** | 21.2 | 148905900 | 0.490 | 2.06 | 0.073 | 0.808 | 0.038 | 13.24 | 30 | 3.14 | 0.62 | 21.4 | 51 | 11.3 | 24.8 | 8 | 3 | 0.49 | 5541 | 57 |
| **37** | 27.5 | 148905900 | 0.490 | 2.15 | 0.051 | 0.87 | 0 | 12 | 28 | 2.93 | 0.41 | 24.6 | 73.6 | 12.5 | 25.2 | 6 | 0 | 0.44 | 4125 | 90 |
| **38** | 10.6 | 148905900 | 0.490 | 1.65 | 0.055 | 0.898 | 0 | 27.45 | 18 | 2.34 | 0.32 | 22.1 | 70.1 | 12.9 | 28.4 | 2 | 0 | 0.58 | 2346 | 118 |
| **39** | 2.24 | 148905900 | 0.490 | 2.85 | 0.047 | 0.843 | 0.086 | 13.04 | 28 | 3.23 | 0.78 | 30.9 | 134 | 11.7 | 32.6 | 6 | 1 | 0.43 | 5627 | 8 |
| **41** | 3.29 | 2700 | 0.103 | 1.48 | 0.068 | 0.84 | 0.016 | 1.47 | 20 | 2.55 | 0.45 | 24.9 | 85.7 | 10.2 | 21 | 3 | 3 | 0.1 | 5981 | 215 |
| **43** | 4.12 | 3600 | 0.082 | 1.88 | 0.055 | 0.857 | 0.064 | 25.93 | 29 | 3.09 | 0.53 | 24.5 | 78.3 | 10.6 | 43 | 5 | 2 | 0.31 | 7848 | 145 |
| **45** | 5.26 | 148905900 | 0.490 | 1.75 | 0.041 | 0.841 | 0.056 | 17.95 | 19 | 2.69 | 0.6 | 22.8 | 58.9 | 9.6 | 23.7 | 5 | 1 | 0.27 | 5363 | 99 |
| **46** | 29.2 | 195300 | 0.410 | 2.35 | 0.041 | 0.836 | 0.03 | 24.39 | 14 | 2.35 | 0.59 | 21.2 | 48 | 13.3 | 33 | 12 | 7 | 0.21 | 4681 | 99 |
| **48** | 36.2 | 148905900 | 0.490 | 2.04 | 0.058 | 0.834 | 0.048 | 25 | 29 | 3.06 | 0.51 | 19.6 | 89.4 | 10.4 | 25 | 8 | 3 | 0.38 | 4840 | 27 |
| **49** | 0.19 | NA | NA | 1.53 | NA | NA | NA | NA | NA | NA | NA | NA | NA | NA | NA | NA | 1 | 0.28 | 5297 | 8 |
| **50** | 21 | NA | NA | 3.39 | NA | NA | NA | NA | NA | NA | NA | NA | NA | NA | NA | NA | 2 | 0.7 | 337 | 19 |
| **Artisanias** | 13.76 | NA | NA | 2.1 | NA | NA | NA | NA | NA | NA | NA | NA | NA | NA | NA | NA | 4 | NA | 12351 | 98 |
| **CBTA** | 26 | NA | NA | 3.21 | NA | NA | NA | NA | NA | NA | NA | NA | NA | NA | NA | NA | 4 | NA | 5937 | 47 |
| **Camino Real** | 6.84 | NA | NA | 1.35 | NA | NA | NA | NA | NA | NA | NA | NA | NA | NA | NA | NA | 2 | NA | 4152 | 31 |
| **Chacamax** | 11.7 | NA | NA | 4.34 | NA | NA | NA | NA | NA | NA | NA | NA | NA | NA | NA | NA | 10 | NA | 5456 | 78 |
| **Leon Brindis** | 5.66 | NA | NA | 2.51 | NA | NA | NA | NA | NA | NA | NA | NA | NA | NA | NA | NA | 3 | NA | 6949 | 103 |
| **La Mision** | 15.4 | NA | NA | 2.41 | NA | NA | NA | NA | NA | NA | NA | NA | NA | NA | NA | NA | 7 | NA | 6112 | 567 |
| **Nututun** | 15.21 | NA | NA | 3.57 | NA | NA | NA | NA | NA | NA | NA | NA | NA | NA | NA | NA | 9 | NA | 5663 | 58 |
| **Quiloma** | 5.25 | NA | NA | 1.73 | NA | NA | NA | NA | NA | NA | NA | NA | NA | NA | NA | NA | 8 | NA | 2362 | 272 |

| | Explanatory variables | | | | | | | | Response variables | | | | | | |
|---|---|---|---|---|---|---|---|---|---|---|---|---|---|---|---|
| | Demography | | | | | | Sample size | | Population genetics | | | | Microbiome | | |
| fragment | total fragment population size | population density | number groups in fragment | proportion AM | proportion AF | ratio of INF:AF | number of samples - genetics | number of samples - microbiome | nucleotide diversity ($\pi$) | inbreeding | mean within-fragment relatednes | mean relatedness to other fragments | Shannon diversity | Faith's PD | mean Bray-Curtis dissimilarity to other fragments |
| **1** | 18 | 0.63 | 3 | 0.17 | 0.39 | 0.43 | 4 | 12 | 0.209 | 0.058 | -0.019 | -0.014 | 5.61 | 36.90 | 0.58 |
| **2** | 7 | 0.86 | 1 | 0.29 | 0.43 | 0 | NA | 7 | NA | NA | NA | NA | 5.31 | 30.33 | 0.57 |
| **3** | 8 | 3.07 | 1 | 0.13 | 0.25 | 1 | 2 | 6 | 0.204 | 0.009 | 0.024 | -0.016 | 5.42 | 32.83 | 0.57 |
| **4** | 3 | 0.82 | 1 | 0.33 | 0.33 | 0 | 2 | 2 | 0.161 | 0.007 | 0.238 | -0.019 | 4.99 | 27.41 | 0.59 |
| **9** | 7 | 0.85 | 1 | 0.14 | 0.29 | 0.5 | NA | 7 | NA | NA | NA | NA | 5.22 | 29.48 | 0.53 |
| **10** | 8 | 0.65 | 2 | 0.13 | 0.75 | 0 | 1 | 8 | 0.207 | NA | NA | -0.012 | 5.43 | 31.06 | 0.56 |
| **12** | 15 | 1.13 | 2 | 0.2 | 0.33 | 0.2 | 3 | 14 | 0.209 | 0.027 | 0.072 | -0.01 | 5.36 | 30.13 | 0.51 |
| **13** | 17 | 0.65 | 3 | 0.24 | 0.29 | 0.25 | 3 | 12 | 0.205 | 0.037 | 0.047 | -0.01 | 5.41 | 30.14 | 0.53 |
| **14** | 5 | 1.1 | 1 | 0.2 | 0.4 | 0.5 | 1 | 3 | 0.18 | NA | NA | -0.003 | 5.39 | 29.86 | 0.60 |
| **16** | 24 | 1.98 | 4 | 0.17 | 0.38 | 0.56 | 2 | 21 | 0.194 | 0.031 | 0.005 | -0.005 | 5.20 | 28.07 | 0.54 |
| **18** | 8 | 1.46 | 1 | 0.13 | 0.38 | 0.33 | 2 | 4 | 0.203 | 0.002 | 0.183 | -0.006 | 5.39 | 31.58 | 0.56 |
| **19** | 18 | 0.67 | 2 | 0.17 | 0.28 | 0.8 | 4 | 10 | 0.205 | 0.028 | 0.08 | -0.008 | 5.60 | 32.18 | 0.54 |
| **20** | 8 | 1.15 | 1 | 0.13 | 0.38 | 1 | 1 | 6 | 0.184 | NA | NA | -0.008 | 5.37 | 32.10 | 0.60 |
| **21** | 22 | 1.88 | 2 | 0.14 | 0.32 | 0.5 | NA | 10 | NA | NA | NA | NA | 5.13 | 30.58 | 0.59 |
| **24** | 3 | 0.83 | 1 | 0.33 | 0.33 | 1 | NA | 3 | NA | NA | NA | NA | 5.01 | 25.26 | 0.65 |
| **25** | 11 | 1.24 | 1 | 0.27 | 0.18 | 1 | 5 | 9 | 0.228 | 0.051 | 0.03 | -0.012 | 4.90 | 28.11 | 0.65 |
| **26** | 7 | 2.21 | 1 | 0.29 | 0.43 | 0 | NA | 7 | NA | NA | NA | NA | 5.20 | 29.89 | 0.59 |
| **28** | 5 | 1.5 | 1 | 0.2 | 0.4 | 0.5 | NA | 2 | NA | NA | NA | NA | 3.88 | 26.80 | 0.73 |
| **29** | 6 | 1.2 | 1 | 0.33 | 0.33 | 0 | 2 | 6 | 0.192 | 0.004 | 0.246 | -0.01 | 5.10 | 28.52 | 0.60 |
| **35** | 27 | 1.27 | 6 | 0.26 | 0.41 | 0.27 | 14 | 25 | 0.235 | 0.167 | 0.018 | -0.009 | 5.66 | 30.27 | 0.52 |
| **37** | 28 | 1.02 | 4 | 0.21 | 0.29 | 0.63 | 11 | 23 | 0.224 | 0.156 | 0.011 | -0.011 | 5.50 | 30.82 | 0.51 |
| **38** | 20 | 1.89 | 2 | 0.25 | 0.25 | 0.2 | 11 | 10 | 0.222 | 0.148 | 0.065 | -0.019 | 5.62 | 33.50 | 0.53 |
| **39** | 5 | 2.23 | 1 | 0.2 | 0.2 | 1 | 1 | 4 | 0.121 | NA | NA | -0.004 | 5.35 | 34.10 | 0.55 |
| **41** | 7 | 2.13 | 1 | 0.29 | 0.29 | 0 | 2 | 7 | 0.189 | 0.019 | 0.004 | -0.006 | 5.10 | 28.14 | 0.57 |
| **43** | 14 | 3.4 | 2 | 0.14 | 0.29 | 0.75 | 2 | 10 | 0.192 | 0.004 | 0.112 | -0.01 | 5.23 | 33.42 | 0.56 |
| **45** | 14 | 2.66 | 2 | 0.14 | 0.36 | 0.2 | 2 | 10 | 0.185 | 0.024 | -0.02 | -0.013 | 5.51 | 33.20 | 0.50 |
| **46** | 8 | 0.27 | 1 | 0.13 | 0.38 | 0.67 | 5 | 7 | 0.202 | 0.051 | 0.121 | -0.008 | 5.29 | 29.70 | 0.54 |
| **48** | 38 | 1.05 | 6 | 0.18 | 0.34 | 0.46 | 10 | 31 | 0.217 | 0.147 | 0.054 | -0.014 | 5.54 | 32.26 | 0.49 |
| **49** | 8 | 42.11 | 1 | 0.13 | 0.38 | 0.67 | 3 | NA | 0.183 | 0.004 | 0.301 | -0.011 | NA | NA | NA |
| **50** | 11 | 0.52 | 2 | 0.18 | 0.36 | 0.25 | 2 | NA | 0.191 | 0.02 | 0.008 | -0.01 | NA | NA | NA |
| **Artisanias** | NA | NA | 6 | NA | NA | 0.2 | 5 | NA | 0.241 | 0.096 | -0.021 | -0.011 | NA | NA | NA |
| **CBTA** | NA | NA | 3 | NA | NA | 0 | 5 | NA | 0.239 | 0.085 | 0.004 | -0.008 | NA | NA | NA |
| **Camino Real** | NA | NA | 1 | NA | NA | 0 | 2 | NA | 0.224 | 0.02 | -0.007 | -0.013 | NA | NA | NA |
| **Chacamax** | 7 | 0.6 | 1 | 0.14 | 0.08 | 0.5 | 4 | NA | 0.223 | 0.066 | 0.04 | -0.005 | NA | NA | NA |
| **Leon Brindis** | 10 | 1.77 | 1 | 0.2 | 0.1 | 1 | 2 | NA | 0.202 | 0.03 | -0.035 | -0.01 | NA | NA | NA |
| **La Mision** | NA | NA | 1 | NA | NA | 0.33 | 2 | NA | 0.2 | 0.039 | -0.032 | -0.012 | NA | NA | NA |
| **Nututun** | NA | NA | 3 | NA | NA | 0.86 | 1 | NA | 0.217 | NA | NA | -0.006 | NA | NA | NA |
| **Quiloma** | 5 | 0.95 | 1 | 0.4 | 0.2 | 1 | 1 | NA | 0.193 | NA | NA | -0.006 | NA | NA | NA |

**Fig. S18. Heatmap of Spearman correlation coefficients for all pairs of explanatory and response variables** (population genetic and microbiome) included in LASSO and multiple regression pipeline. Significant correlations are marked with an asterisk, and the Spearman's *r* value is denoted by color according to the scale to the right of the plot.

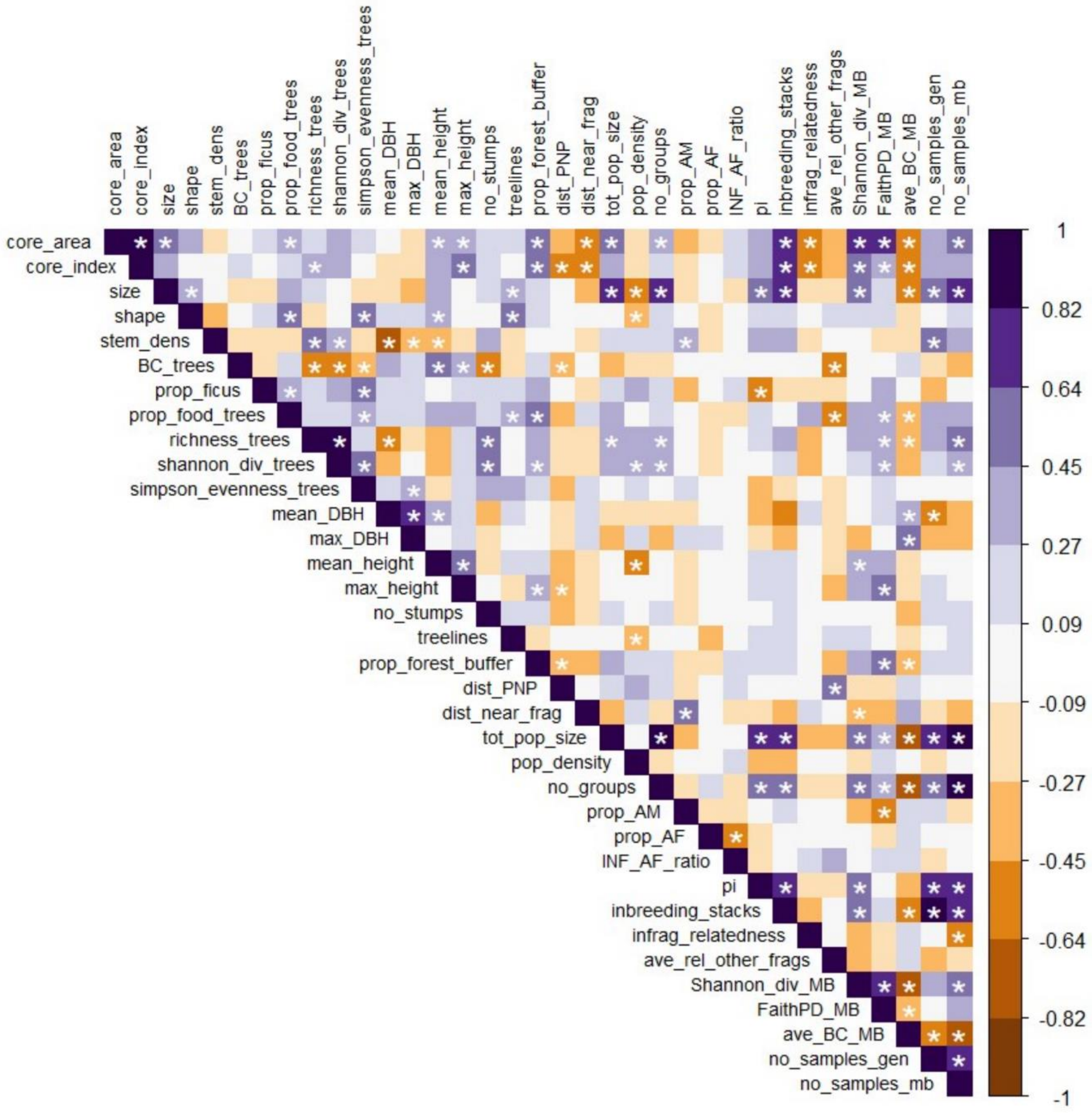

**Fig. S19. Results of combined exhaustive fragment-deletion, cross-validation, and predictor-perturbation sensitivity analysis with LASSO and AICc:** Frequency of explanatory variables (y-axis) selected in ≥33% of reduced datasets (323 for population genetic variables and 1218 for microbiome variables; these thresholds are denoted with dashed vertical lines in plots), with (red) and without (blue) up to 5% randomized error rates in the explanatory variables, for each response variable. Top – population genetic response variables. Bottom – gut microbiome response variables.

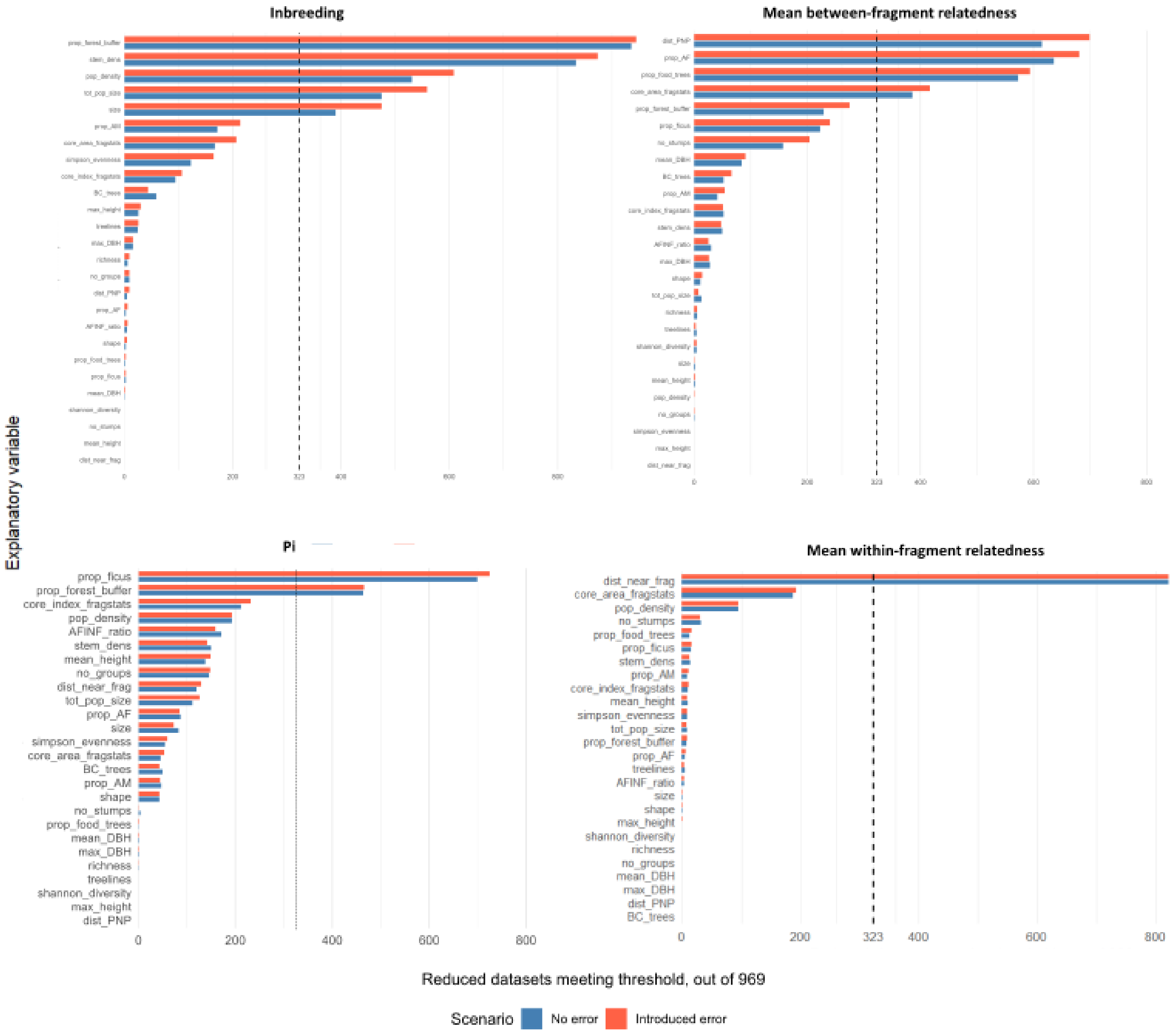

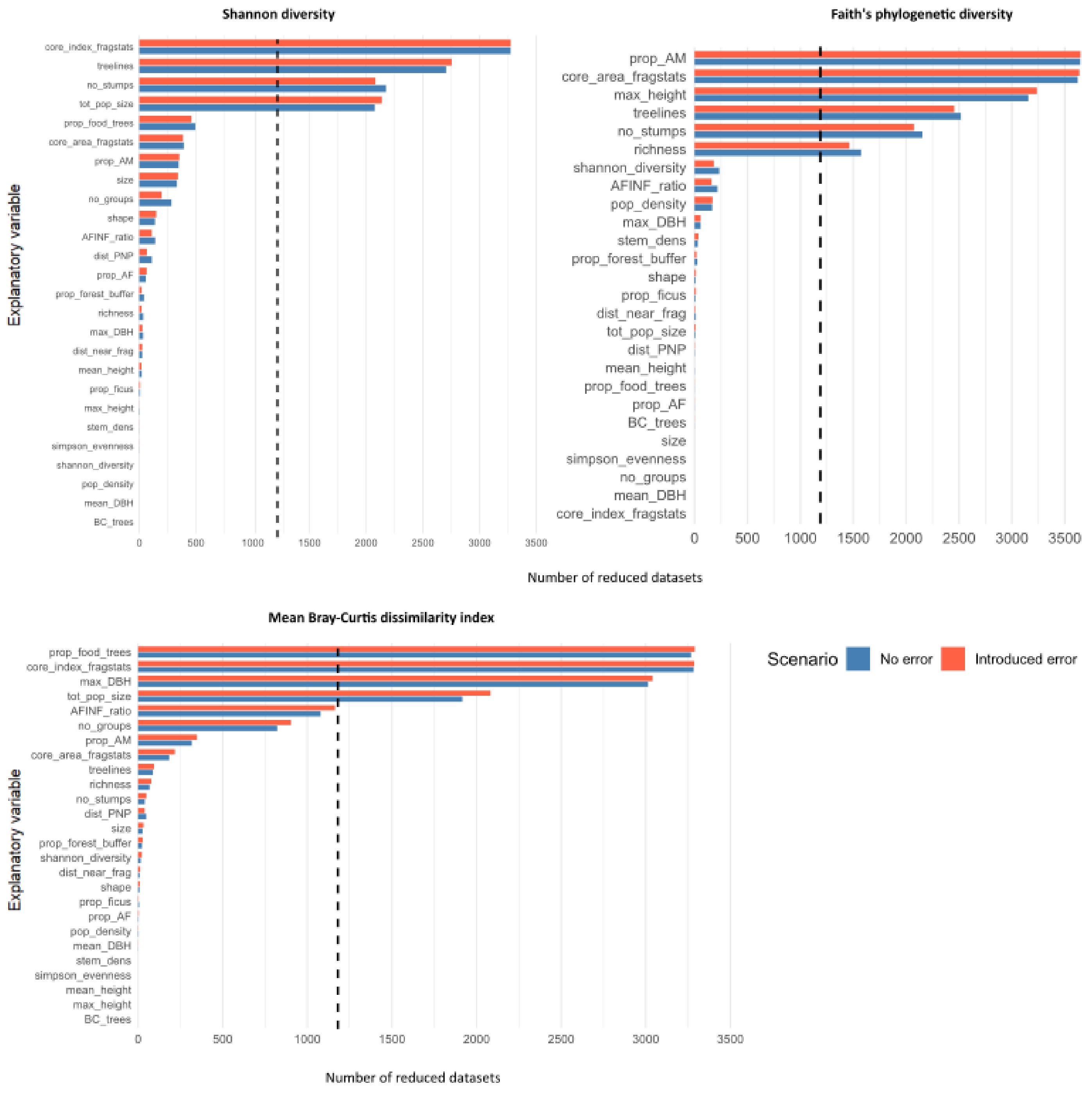
Shannon diversity
Explanatory variable
core_index_fragstats
treelines
no_stumps
tot_pop_size
prop_food_trees
core_area_fragstats
prop_AM
size
no_groups
shape
AFINF_ratio
dist_PNP
prop_AF
prop_forest_buffer
richness
max_DBH
dist_near_frag
mean_height
prop_ficus
max_height
stem_dens
simpson_evenness
shannon_diversity
pop_density
mean_DBH
BC_trees
0
500
1000
1500
2000
2500
3000
3500
Faith's phylogenetic diversity
prop_AM
core_area_fragstats
max_height
treelines
no_stumps
richness
shannon_diversity
AFINF_ratio
pop_density
max_DBH
stem_dens
prop_forest_buffer
shape
prop_ficus
dist_near_frag
tot_pop_size
dist_PNP
mean_height
prop_food_trees
prop_AF
BC_trees
size
simpson_evenness
no_groups
mean_DBH
core_index_fragstats
0
500
1000
1500
2000
2500
3000
3500
Number of reduced datasets
Mean Bray-Curtis dissimilarity index
Explanatory variable
prop_food_trees
core_index_fragstats
max_DBH
tot_pop_size
AFINF_ratio
no_groups
prop_AM
core_area_fragstats
treelines
richness
no_stumps
dist_PNP
size
prop_forest_buffer
shannon_diversity
dist_near_frag
shape
prop_ficus
prop_AF
pop_density
mean_DBH
stem_dens
simpson_evenness
mean_height
max_height
BC_trees
0
500
1000
1500
2000
2500
3000
3500
Number of reduced datasets
Scenario
No error
Introduced error

#### Results of multiple regression analyses

**Table S17. Full output of final multiple regression models for all response variables.**

| Response variable | Predictor variable | B | 95% CI | P-value | $R^2$ | Adjusted $R^2$ | F statistic | N |
|---|---|---|---|---|---|---|---|---|
| **Population genetics** | | | | | | | | |
| **Inbreeding** | | | | | 0.953 | 0.933 | $F_{5,12} = 13.26$, $p < 0.001$ | 18 |
| | proportion of forest in fragment buffer | 0.0385 | 0.0286 to 0.0483 | <0.001 | | | | |
| | stem density | 0.0214 | 0.0116 to 0.0312 | <0.001 | | | | |
| | population density | -0.0183 | -0.0318 to -0.0048 | 0.012 | | | | |
| | population size | 0.0124 | -0.0048 to 0.0296 | 0.141 | | | | |
| | fragment size | 0.0071 | -0.0118 to 0.0259 | 0.429 | | | | |
| **Nucleotide diversity (π)** | | | | | 0.667 | 0.622 | $F_{2,15} = 15.0$, $p < 0.001$ | 18 |
| | proportion of forest in fragment buffer | 0.0090 | 0.0033 to 0.0147 | 0.004 | | | | |
| | proportion of ficus trees | -0.0129 | -0.0186 to -0.0072 | <0.001 | | | | |
| **Mean within-fragment relatedness** | | | | | 0.509 | 0.478 | $F_{1,16} = 15.0$, $p < 0.001$ | 18 |
| | distance to nearest fragment | 0.0578 | 0.0277 to 0.0879 | <0.001 | | | | |
| **mean relatedness to other fragments** | | | | | 0.803 | 0.743 | $F_{4,13} = 13.26$, $p < 0.001$ | 18 |
| | proportion important food trees | -0.0014 | -0.0025 to -0.0002 | 0.025 | | | | |
| | proportion of adult females | 0.0017 | 0.0005 to 0.0028 | 0.007 | | | | |
| | distance to PNP | 0.0023 | 0.0008 to 0.0037 | 0.005 | | | | |
| | fragment core area | -0.0009 | -0.0023 to 0.0005 | 0.197 | | | | |
| **Gut microbiomes** | | | | | | | | |
| **Shannon diversity** | | | | | 0.594 | 0.523 | $F_{4,23} = 8.408$, $p < 0.001$ | 28 |
| | proportion of core area in fragment | 0.1980 | 0.0963 to 0.2997 | < 0.001 | | | | |
| | number of treelines | 0.1226 | 0.0255 to 0.2197 | 0.016 | | | | |
| | number of tree stumps | -0.1192 | -0.2203 to -0.0181 | 0.023 | | | | |
| | population size | 0.1048 | 0.0054 to 0.2042 | 0.04 | | | | |

| | | | | | | | | |
|---|---|---|---|---|---|---|---|---|
| **Faith's phylogenetic diversity** | | | | | 0.834 | 0.786 | $F_{6,21} = 17.52$, $p < 0.001$ | 28 |
| | fragment core area | 0.9388 | 0.3573 to 1.5203 | 0.003 | | | | |
| | number of tree stumps | -1.0391 | -1.6428 to -0.4354 | 0.002 | | | | |
| | number of treelines | 0.7343 | 0.2414 to 1.2272 | 0.005 | | | | |
| | proportion of adult males | -1.2698 | -1.7677 to -0.7719 | < 0.001 | | | | |
| | tree genera richness | 0.9030 | 0.2377 to 1.5683 | 0.01 | | | | |
| | maximum tree height | 0.5809 | 0.0469 to 1.1149 | 0.035 | | | | |
| **Mean Bray-Curtis dissimilarity index** | | | | | 0.63 | 0.566 | $F_{4,23} = 9.796$, $p < 0.001$ | 28 |
| | proportion of food trees | -0.0238 | -0.0389 to -0.0087 | 0.003 | | | | |
| | proportion of core area in fragment | -0.0219 | -0.0366 to -0.0072 | 0.002 | | | | |
| | maximum tree DBH | 0.0180 | 0.0027 to 0.0333 | 0.023 | | | | |
| | total population size | -0.0142 | -0.0297 to 0.0013 | 0.071 | | | | |

**Fig. S20. Linear regression plots for all pairs of explanatory and response variables found to have a significant relationship**, from among the variables found to contribute significantly to each variable's multiple regression model. Top – population genetic response variables. Bottom – gut microbiome response variables.

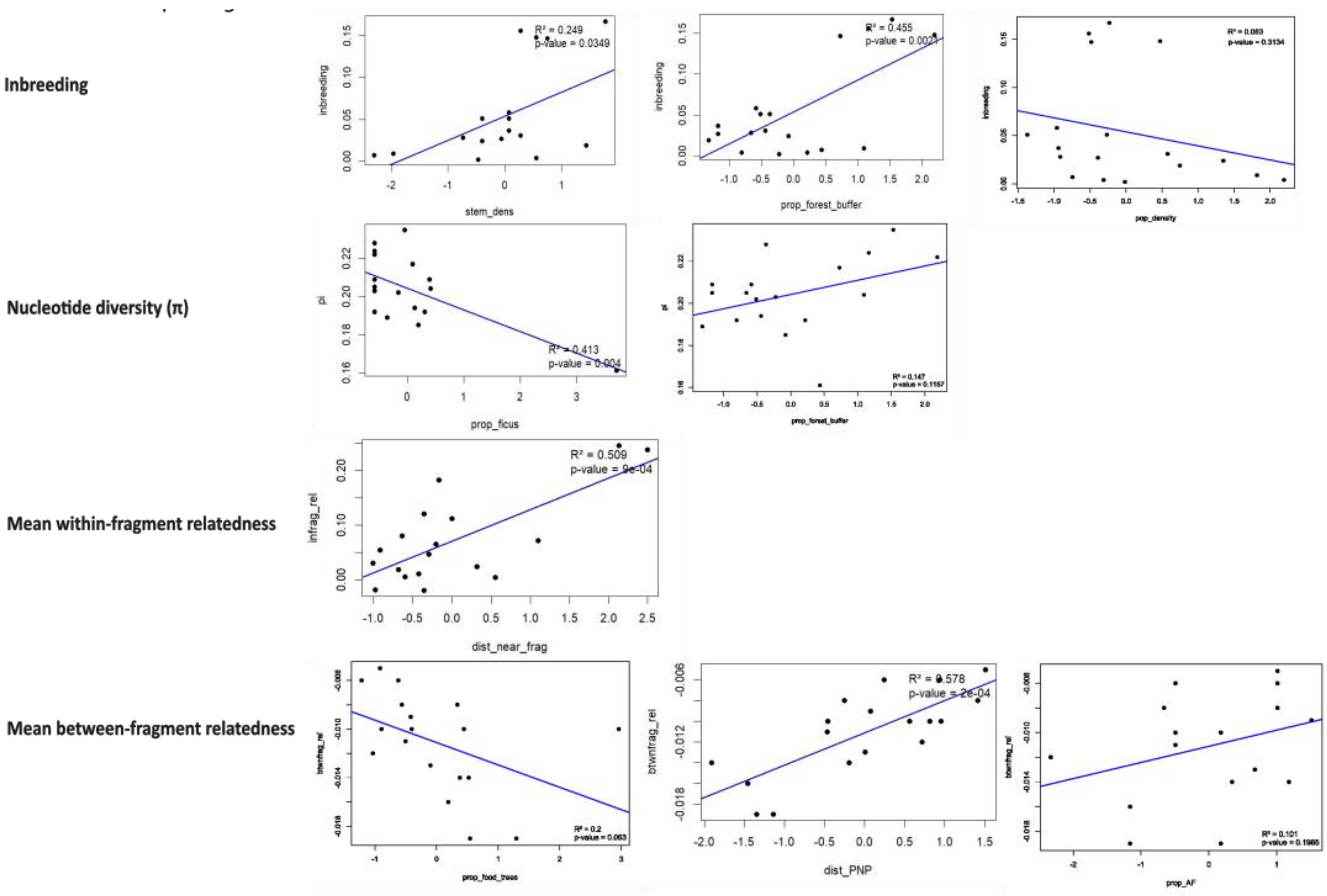

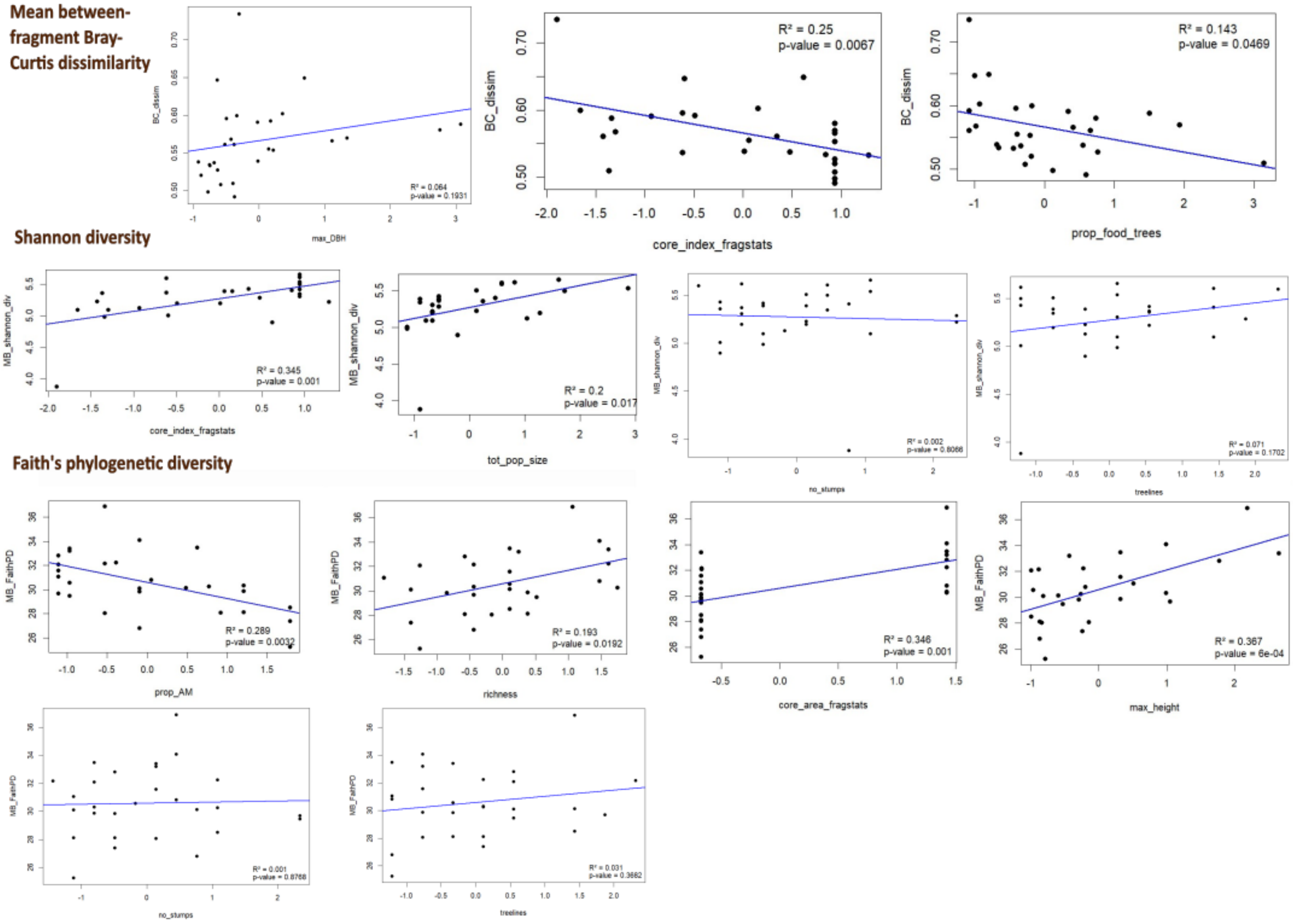

Mean between-fragment Bray-Curtis dissimilarity
BC_dissim
max_DBH
R² = 0.064
p-value = 0.1931
BC_dissim
core_index_fragstats
R² = 0.25
p-value = 0.0067
BC_dissim
prop_food_trees
R² = 0.143
p-value = 0.0469
Shannon diversity
MB_shannon_div
core_index_fragstats
R² = 0.345
p-value = 0.001
MB_shannon_div
tot_pop_size
R² = 0.2
p-value = 0.017
MB_shannon_div
no_stumps
R² = 0.002
p-value = 0.8066
MB_shannon_div
treelines
R² = 0.071
p-value = 0.1702
Faith's phylogenetic diversity
MB_FaithPD
prop_AM
R² = 0.289
p-value = 0.0032
MB_FaithPD
richness
R² = 0.193
p-value = 0.0192
core_area_fragstats
R² = 0.346
p-value = 0.001
MB_FaithPD
max_height
R² = 0.367
p-value = 6e-04
MB_FaithPD
no_stumps
R² = 0.001
p-value = 0.8768
MB_FaithPD
treelines
R² = 0.031
p-value = 0.3662

#### Additional sources found only in Supplementary Information
